# ABEL: an active-learning behavior estimation and labeling platform

**DOI:** 10.64898/2026.08.30.748115

**Authors:** Jobe L. Ritchie, Brianna E. George, Alison V. Roland, Caroline G. Krieman, Brooke N. Bender, Maya R. Eberle, Grace A. Stys, William C. Der, Lili S. Kooyman, Adriane M. Lawes, Rajon T. Scott, Todd K. O’Buckley, Marissa M. R. McLean, Chloe J. Gallagher, Joyce Besheer, Thomas L. Kash

## Abstract

Detailed behavior analysis is essential for thorough characterization of ethologically relevant behaviors in model organisms, yet manual annotation of the full behavioral repertoire remains subjective, time intensive, and susceptible to observer error. Advances in machine learning have enabled high-throughput pose estimation on recorded video, but tools for behavior classification from pose and video data are still developing. Instead of hand-scoring every frame of video, experimenters can instead label a small subset of video frames and software trained through machine learning makes predictions on the rest. Here, we present an Active-learning Behavior Estimation and Labeling (ABEL) platform that uses clip-level active learning (i.e., human labeling of short video snippets) with multimodal features (pose, video, context/ROI) to train robust behavior classifiers. We rigorously validated ABEL-derived behavior predictions against expert human observers and field-standard automated software, across diverse rodent behavioral assays. Across eight assays and 45 behaviors, model training required 19.5 hours of human annotation in total, with the reviewer scoring ∼8% of available video. Models trained in ABEL achieved a mean precision-recall area under the curve (PR-AUC – a 0-1 score of how well a model balances missed detections against false alarms, with 1 being perfect) of 0.90 (SD 0.09, range 0.60-0.99), with no association between performance and behavior prevalence (r = 0.20). This was aided by custom tools, Essence Extractor and UMAP Interactive Selection, for targeted discovery of high probability clips which reduce the clip review needed to find a rare behavior 6-fold relative to random sampling and 10-fold relative to labeling whole videos. As a biological validation, we assessed how ABEL-derived behaviors relate to underlying neuronal calcium dynamics. Behavior labels were tightly synced with neuronal signatures distinct from ambiguous behavior and randomly chosen, behavior-unrelated time windows (shuffle control). Together, these data indicate that ABEL provides an efficient platform for frame-precise classification of distinct ethologically relevant behaviors.

## Introduction

The impact of an experimental manipulation on an animal’s behavioral state is often inferred from a small number of summary measures derived from the animal’s location, such as time in a zone or distance traveled. Though descriptive in their own right, these measures overlook a rich repertoire of potentially informative behavioral data. Traditional methods of labeling discrete behaviors (e.g., manual scoring from video recordings) are prohibitively time-consuming. As such, most analyses stop short of detailed characterization of ethologically relevant behaviors. Modern machine learning approaches address this deficit by enabling rigorous behavior analysis with high throughput.

Keypoint tracking of freely-behaving organisms is largely a solved problem. Advances in convolutional neural networks (CNNs) have enabled near-human precision tracking of keypoints using minimal user-labeled data. Multiple freely-available pipelines for generating accurate keypoint tracking data from raw video are available, including DeepLabCut (DLC)^1^ and SLEAP^2^. Yet, keypoint tracking alone provides limited insight into behavior without additional analysis. Thus, translating pose estimates into biologically meaningful behavior classification is the critical next step.

Computational approaches to behavior classification fall into two main categories, each with their own tradeoffs. Unsupervised methods, such as motion sequencing (MoSeq^3^) or keypoint-informed MoSeq^4^, discover recurring sub-second movement motifs directly from raw data without human labeling. While unsupervised approaches are powerful for unbiased discovery of novel patterns of motion, they can lack face validity and interpretability, as discovered motifs may not necessarily map onto ethologically relevant behaviors. Supervised methods, such as Simple Behavioral Analysis (SimBA)^5^, bridge this gap by introducing annotation of video with specific, named behaviors to train behavior classifiers. Human-provided behavior labels strengthen face validity but reintroduce the potential efficiency bottleneck of human labeling. An active-learning approach, introduced by A-SOiD^6^, addresses the issue of labeling burden by surfacing uncertain frames for review rather than requiring exhaustive annotation, reducing labeling required by 85% relative to alternative supervised approaches, while achieving similar classification performance.

We present a graphical user interface (GUI)-based, no-coding required platform for human-in-the-loop annotation and training of predictive models for behavior analysis. ABEL (Active-learning Behavior Estimation and Labeling) adapts the frame-level active-learning approach^6^ to a clip-level pipeline with a built-in annotation platform. Temporally- and contextually-aware feature extraction adds pose, video, and region of interest (ROI)-context features that pose alone does not provide. This yields models with high precision on held-out subjects and improved differentiation of like behaviors. Additionally, ABEL is equipped with a suite of data analysis, visualization, and validation tools to provide data insights including group-based analyses, graphing, statistics, and export of bout start and end frames for use with aligning fiber photometry data. Here, we validated ABEL against human labels and a selection of field-standard automated approaches across a diverse range of common rodent behavior assays. Custom tools in ABEL, including Uniform Manifold Approximation and Projection (UMAP) Interactive Selection and Essence Extractor, surface 1.3-1.5x more positive clips than active-learning at low clip budgets and reduced human labeling effort to discover low-abundance behavior positive clips by 1.8x. We next provide proof-of-principle biological validation through streamlined integration of ABEL behavior predictions with underlying neuronal activity, using calcium dynamics in the central nucleus of the amygdala (CeA) and bed nucleus of the stria terminalis (BNST). ABEL behavior predictions were distinct in neuronal activity signature relative to random noise or ambiguous behavior, and behavior start frames had sub-second precision with peak calcium dynamics. The accuracy, precision, and efficiency of ABEL make it a valuable tool for thorough behavior analysis and integration with *in vivo* measures such as fiber photometry. ABEL is available free for academic use on GitHub (https://github.com/JobeRitchie/ABEL.git).

## Results

ABEL is a human-in-the-loop pipeline for annotation and training of machine learning models that estimates behavior from keypoint labeling and raw video data (**Fig. 1**). At the core of ABEL is the active-learning approach, which selectively extracts high-impact uncertain examples for review. ABEL builds on this approach, adapting it to a clip-based data structure with added multi-modal features. Additionally, ABEL provides two specialized tools for model refinement, Essence Extractor and UMAP Interactive Selection, through a GUI. ABEL takes the user from raw pose-tracked data to refined exports in a single pipeline (**Fig. 1**).

**Figure 1.**
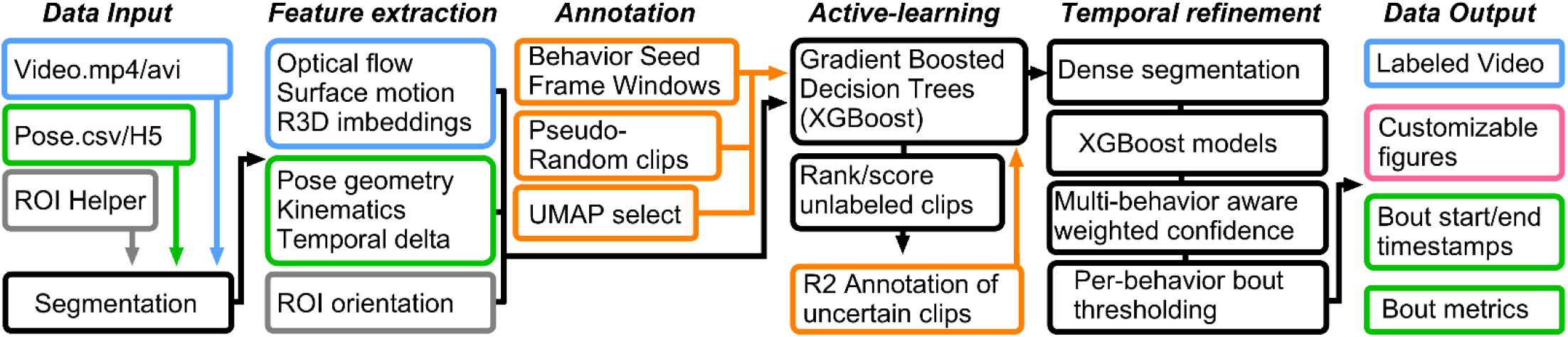
ABEL is a human-in-the-loop GUI-based pipeline for annotation and iterative refinement of behavior models. Flowchart schematic of data flow from raw video and keypoints to refined behavior exports. Data input in the form of video (blue) can be in mp4 or avi format. Keypoint tracking (green) from either DeepLabCut or SLEAP can be in .csv or H5 format. Region of interest (ROI; grey) data is defined within an interactive tool in ABEL. Data is then segmented, and a robust feature set is extracted per segment. The user then annotates clips (orange) using ABEL’s built-in annotation suite. Annotations and extracted features are then used to train gradient-boosted decision trees, which can then extract additional clips for iterative model refinement. Whole sessions are then densely segmented and analyzed by refined XGBoost models. Per-behavior confidence thresholds and between-behavior confidence suppression (mutual inhibition) is then applied. Data can then be visualized in ABEL and exported in several formats tailored for ease of analysis in graphing and statistics software.

ABEL accepts input of raw video in .mp4/.avi format and is compatible with keypoint tracking from either DeepLabCut or SLEAP platforms. Incorporated in ABEL is an ROI helper for defining any number of zones for context-level feature extraction and providing time in ROI as a basic endpoint. This feature supports multi-ROI labeling in a session-specific or project-wide manner. The raw input data are then segmented into user-defined clip lengths, depending on the length of behaviors of interest. Features across video, pose, and context/ROI domains are then extracted from data segments. Feature count scales with number of body parts tracked and ROI count (∼2080 on average, full per-project feature set available in **Supplement Table 1**). The user then provides a first round of annotations via seed clips, through dispersed pseudorandom labeling (i.e. Random Low-Prob (absent)), or by zone selection in an unsupervised UMAP. Notably, the annotation process is highly streamlined, allowing experienced users to annotate clips in as little as 1-5 seconds per clip and has built-in features for review of scored clips. Clip features and annotations are then used to train gradient-boosted decision trees (XGBoost), which score uncertain clips for further annotation and training through an active-learning process. Confidence scores from a second dense segmentation step are then adjusted via mutual inhibition (default weight = 0.20) which can be tailored by behavior pair to disambiguate similar competing behaviors. Finally, per-behavior confidence thresholds, minimum bout length, and temporal merging are assigned.

### ABEL agrees with expert human observation across a wide range of assays and diverse behaviors

We first sought to compare ABEL-derived behavior estimations against a well characterized reference dataset, our previously published home cage behavior data set that we have rigorously validated^7^. Our published data used DeepLabCut and SimBA to assess rearing, grooming, and digging behaviors in a home cage environment from overhead videos. Thus, models were created in ABEL to detect these same behaviors using the same videos as input data. Note that all models presented here were applied to data not used for model training. Lin’s concordance correlation coefficient (CCC) is reported as the primary metric of observer agreement due to its sensitivity to differences in magnitude, with Pearson correlation provided secondarily. In the home cage behavior assay (**Fig. 2A**), ABEL models reached F1 = 0.861-0.914 and PR-AUC = 0.862-0.932. Time spent rearing, grooming, and digging agreed with expert human scoring and matched or exceeded SimBA on concordance (**Fig. 2B-D**; CCC, ABEL vs SimBA: 0.92 (95% CI 0.77–0.97) vs. 0.81 (0.54–0.93); grooming 0.97 (0.90-0.99) vs. 0.98 (0.95-0.99); digging 0.98 (0.91-0.99) vs. 0.87 (0.62-0.96); n = 10 mice). Pearson *r* was 0.946, 0.987 and 0.979 for ABEL and 0.904, 0.994 and 0.901 for SimBA (all p < 0.001). Bland-Altman analysis, which assesses agreement between quantitative measures, showed a small negative offset for grooming (−30.3 s, −55.3 to −5.3; 15% of the mean), indicating that ABEL slightly under-reports long grooming bouts possibly due to oversegmentation.

**Figure 2.**
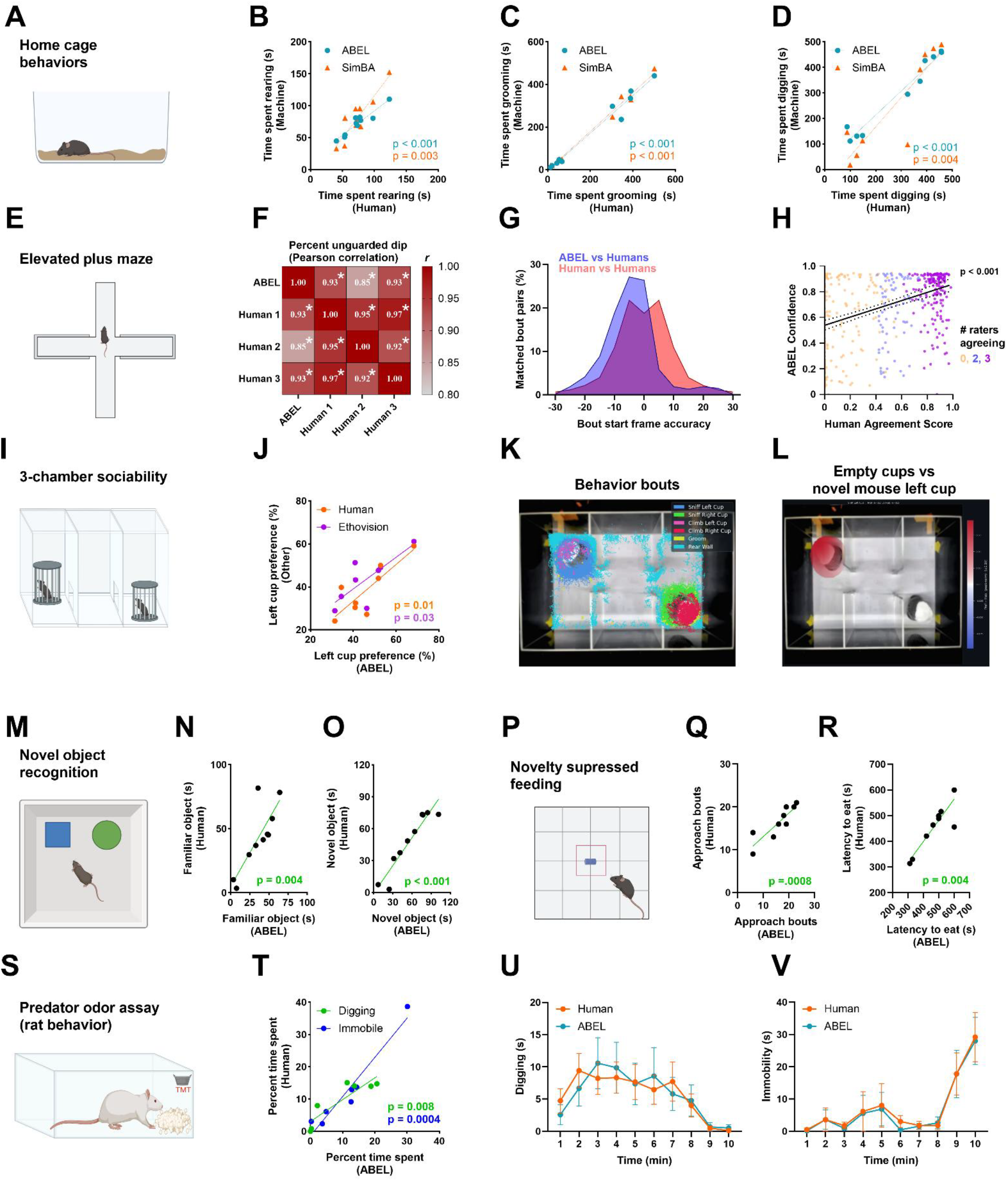
ABEL aligns with expert human observation and other field-standard tools across diverse assays and behaviors. **(A)** Illustration of home cage setup used for validation of ABEL-derived behavior labels against expert human observation and Simple Behavioral Analysis (SimBA). Home cage validation (n = 10 mice) was conducted for **(B)** rearing, **(C)** grooming, and **(D)** digging. **(E)** Illustration of elevated plus maze. **(F)** Pearson correlation matrix of ABEL-derived predictions (n = 10 mice) of pDips compared with three human observers. *p < 0.05. **(G)** Bout start frame accuracy of ABEL vs. all human observers and human observer. vs rest. **(H)** ABEL prediction confidence compared to human observer agreement score. Line represents mean ± SEM. Color coding determined by number of raters agreeing on bout start within 30 frames. **(I)** Illustration of 3-chamber social assay. **(J)** Comparison of ABEL, human, and Ethovision prediction (n = 8 mice) of left cup preference. **(K)** Composite overlay of 3-chamber social assay and ABEL-predicted behaviors. **(L)** Difference in left cup sniffing between acclimation day and social preference day. Novel mouse in left cup and no mouse in right cup. **(M)** Illustration of novel object recognition assay. **(N)** Correlation of novel object investigation time from human scoring and ABEL predictions (n = 10 mice). **(O) C**orrelation of familiar object investigation time from human scoring and ABEL predictions. **(P)** Illustration of novelty suppressed feeding assay. **(Q)** Correlation of approach behavior from human scoring and ABEL predictions (n = 10 mice). **(R)** Correlation of latency to eat behavior for human scoring and ABEL predictions. **(S)** Illustration of predator odor assay in rats. **(T)** Correlation of percent time digging and immobile from human scoring and ABEL predictions (n = 7 rats). **(U)** Time spent digging in one-minute bins. **(V)** Time spent immobile in one-minute bins.

We next assessed the consistency of ABEL behavior estimation in the elevated plus maze (EPM), compared with manual scoring by a set of three human observers (**Fig. 2E**). For analysis, the four ‘observers’ (ABEL + three humans) were compared on head dip and protected head dip (pDip), operationally defined as lowering the head off the open arm while the hind section remains outside or inside of the closed arm, respectively^8^. ABEL classified dips and pDips with F1 = 0.91 and 0.89 (PR-AUC = 0.94 and 0.91). The proportion of pDips out of total dips (pDip/Dip+pDip), an established measure of anxiety-like behavior^9^, was calculated and compared between observers. ABEL tracked with human consensus in rank order (**Fig. 2F**; ABEL vs. human mean r = 0.93, 0.85, 0.93, 95% CI 0.68-0.98, n=10, p=0.0002) but returned lower absolute values (Lin’s CCC = 0.76, 95% CI 0.43-0.91; Bland-Altman bias -10.99%) while three human observers had little offset (CCC 0.90-0.95; bias -0.6 to -4.7%). Exact behavior start frame accuracy of ABEL compared to the human average start frame (Machine vs. humans) was highly overlapping with human vs. humans accuracy, though ABEL labels were shifted slightly earlier by a mean of 4.4 frames (0.15 s at 30 Hz; SD=9.0; **Fig. 2G**). A challenge that we encountered while validating EPM behavior was inter-rater reliability, especially when comparing frame-specific accuracy. Considering this, we next assessed how confident ABEL was at predicting dips from the entire human-scored data set and plotted this against a human consensus score. We found that model confidence in dip behaviors scales up with human consensus (**Fig. 2H**; r = 0.42, CI 0.33-0.49, n = 414 bouts, p < 0.001; mean confidence 0.56, 0.66, 0.79 for 0, 2, 3 raters in agreement; Kruskal-Wallis H = 46.88, p < 0.0001). This suggests that ABEL performs similarly to humans at predicting behavior across a spectrum of behavior ambiguity.

The 3-chamber social assay can be scored based on proximity tracking using commercially available software, but expert human scoring remains the gold standard due to the challenge of automated differentiation of social investigation from general exploratory behaviors^10,11^. Using multi-ROI feature extraction in ABEL, we achieved preference scores in line with both an expert human observer and time spent in zone extracted using Ethovision XT software by Noldus (**Fig. 2I-J**; ABEL vs. human r = 0.83, 95% CI 0.31–0.97, CCC = 0.68, p = 0.010; ABEL vs. Ethovision r = 0.78, 95% CI 0.16–0.96, CCC = 0.75, p = 0.024; n = 8 mice). The built-in behavior overlay visualization demonstrates clear differentiation of ROI-specific and non-specific behaviors (**Fig. 2K**). This is exemplified further by the change in left cup exploration on the habituation day (empty cups) versus the novel social preference day (**Fig. 2L**; novel mouse in left cup, faux mouse in right cup).

Multi-ROI features similarly helped achieve human-like performance in the novel object recognition task (**Fig. 2M-O**; novel object r = 0.95, 95% CI 0.79–0.99, CCC = 0.91; familiar object r = 0.81, 95% CI 0.38–0.95, CCC = 0.75; n = 10 mice, both p < 0.005). Using a single ROI, we also achieved reliable prediction of distinct food-directed behaviors in the novelty suppressed feeding task (**Fig. 2P-R**; approach count r = 0.88, 95% CI 0.56–0.97, CCC = 0.79; latency to eat r = 0.90, 95% CI 0.63–0.98, CCC = 0.89; n = 10 mice, both p < 0.001).

Finally, we assessed the efficacy of ABEL with a side-viewing angle of rats in the predator odor assay (2,3,5-Trimethyl-3-Thiazoline; TMT; **Fig. 2S**). Overall percent time spent digging and immobile agreed with expert human observation (**Fig. 2T**; immobility r = 0.97, 95% CI 0.78–0.99, CCC = 0.94; digging r = 0.89, 95% CI 0.40–0.98, CCC = 0.86; n = 7 rats, both p < 0.01). Plotting the data across time in 1-minute bins revealed close numerical agreement between ABEL and expert observation (**Fig. 2U-V**; per-rat mean difference, digging +0.03 s/bin, 95% CI −2.29 to +2.36; immobility −0.62 s/bin, 95% CI −2.76 to +1.52), though the low sample size prohibited formal testing of equivalence. In sum, these data indicate that ABEL estimates behavior at a level comparable to expert human observation across the assays tested.

### ABEL is data-efficient and provides advanced tools for model refinement

ABEL uses a clip-based adaptation of the active-learning approach^6^. Active-learning intentionally extracts uncertain, low-confidence clips for review for subsequent training. This enables more data-efficient model refinement as compared to reviewing randomly selected clips or entire videos, thus reducing the number of clips required to reach sufficient model quality. On average across 45 behavior models and eight distinct assays, a clip budget of 250 in ABEL yields a mean F1 of 0.70 and PR-AUC of 0.80 (**Fig. 3A**; average of five random seed replicates trained on positive clips with 5x negative clip class imbalance). Notably, performance showed no detected association with behavior prevalence spanning 0.26-44.39% of reviewed segments (**Fig 3B**; Pearson r = 0.20, 95% CI -0.10 to 0.46, n = 45 behaviors, p = 0.19 on log10 prevalence). Across all behaviors assessed, agreement between ABEL and expert annotation averaged k = 0.79 (95% CI 0.76-0.82, n = 45 behaviors) and 96% of behaviors scored above k = 0.6 (**Fig. 3C**). Performance in Fig. 3 is reported at the clip level, before dense temporal refinement and should be taken as a conservative lower bound. The bouts that ABEL exports are further refined by smoothing, thresholding, and bout duration filtering.

**Figure 3.**
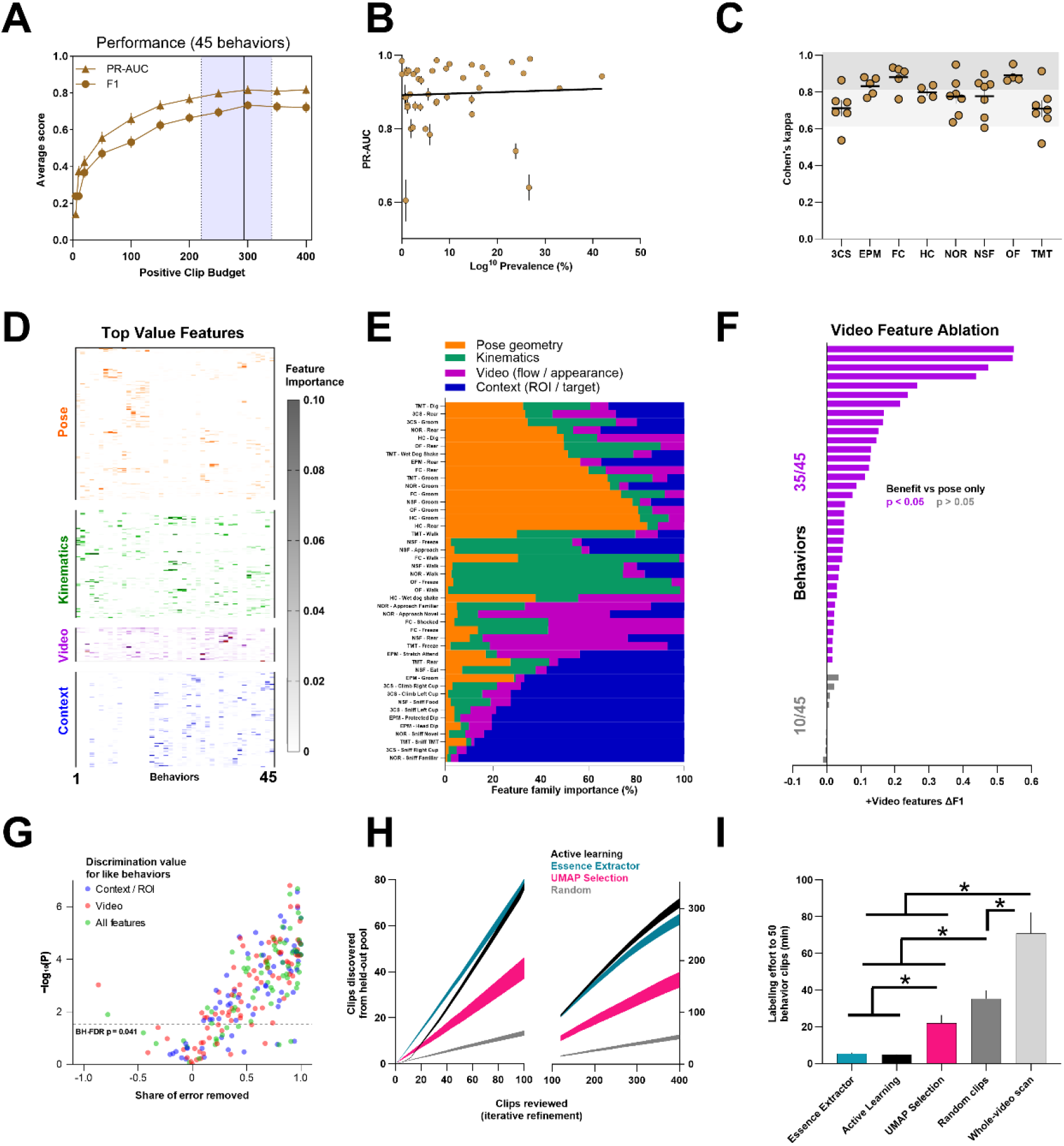
Predictive models developed in ABEL are precise and accurate due to diverse feature extraction and tools for refinement. **(A)** Training metrics (mean ± SEM) given different positive clip budgets (45 behaviors at ≤100 clips falling to 30 at 400). **(B)** Precision-recall area under the curve across varying behavior prevalence. **(C)** Cohen’s Kappa reliability (mean ± SEM) for eight assays. Data points represent behaviors analyzed. **(D)** Top-importance features as determined by normalized model gain across 3 seeds, ordered by likeness using hierarchical clustering. **(E)** Percent share of feature family importance per behavior, ordered by most important feature family. **(F)** F1 gain (mean ± SD) from inclusion of video feature family. Behaviors labeled with purple represent significant gain in F1 across 5 seeds. **(G)** Share of behavior pair differentiation error removed by inclusion of context and/or video features. Dotted line indicates BH-FDR corrected alpha. **(H)** Number of clips surfaced (mean ± SEM) at different clip budgets using different clip surfacing strategies across 45 models. *Posthoc* comparisons were made at 25 and 300 clip budgets. **(I)** Minutes of review time to confirm 50 positive clips (mean ± SEM) based on clip generation approach. Clip review is charged at 4 s per 0.5 s clip and 1x real time for whole video scan. Brackets indicate two-tailed t-tests with Holm correction. *p<0.05.

Two primary strengths of ABEL are robust feature extraction through a combination of video, pose, and ROI-associated datapoints, and an extensive toolset for model refinement. ABEL extracts 2080 features on average that vary in predictive importance by behavior (**Fig. 3D**; Feature importance is based on XGBoost gain normalized within model). This extensive feature space enables identification of diverse behaviors that rely on distinct features or feature families (**Fig 3E**). Although video features accounted for a relatively small portion of total model gain across projects, withholding the video feature family significantly reduced held-out ΔF1 in 35 of 45 behaviors (**Fig 3F**; 2-tailed paired t-tests on per-seed F1 gains vs. pose only, BH-FDR corrected q < 0.05). Context/ROI features provide even more value than video features due to ROI-directed behaviors in several assays. Including context/ROI features improved held-out F1 by a mean of 0.13 across 32 behaviors in projects with defined ROIs (28 of 32 behaviors improved; not shown). Differentiating like behaviors is challenging from pose alone. To assess how well ABEL can differentiate similar behaviors, we next generated models for all behaviors with and without each feature family. Inclusion of context and/or video features improves pairwise discrimination in 70 of 82 behavior pairs (**Fig. 3G**; median 69.3% reduction in remaining pose-only model error, 2-tailed paired t-tests with BH-FDR correction on per-seed ΔROC-AUC, ps < 0.041).

ABEL provides intuitive tools for surfacing positive clips during early model refinement and for low-abundance behaviors. The first tool, the Essence Extractor (within ABEL’s Targeted Clip Mining dialog), surveys a subset of user selected clips for shared features which differ from the total clip population. The second tool, UMAP Interactive Selection, allows for manual selection based on proximity in a reduced-dimension feature space. The relative value of selection tool choice when identifying positive clips depends on clip budget (**Fig 3H**; two-way repeated-measures ANOVA on discovered clips, 45 behaviors; strategy x budget interaction F_(3,126)_ = 162.46, p < 0.001, main effect of strategy, F_(3,126)_ = 183.41, p < 0.001). At a clip budget of 25, Essence Extractor was more effective than either UMAP Interactive Selection (1.89x, p = 0.0005) or active-learning (1.55x, p < 0.001) making it the most effective tool during early model refinement. Both targeted strategies extracted ∼5-6x more positives than random selection at every tested clip budget. However, at a clip budget of 300, active-learning overtakes Essence Extractor (1.36x, p < 0.001) and UMAP Interactive Selection (1.71x, p < 0.001), suggesting active-learning is better suited to later model refinement. This relationship shifts when searching for low-prevalence behaviors (**Fig. 3I;** prevalence of rarest behavior from 8 assays = 1.23 ± 0.41% SEM of reviewed segments). Across the eight rarest behaviors (one per assay), all targeted strategies reduce human labeling effort to discover 25 rare behavior positive clips relative to random selection and whole-video scan (minutes to label 25 positive clips for the eight rarest behaviors, assuming 4 s per clip; one-way repeated-measures ANOVA on log10 minutes with behavior as the repeating unit, F_(4,28)_ = 40.40, p < 0.001). Essence Extractor and active learning performed similarly for rare-behavior discovery (p=0.76) and both substantially outperformed alternatives (All p < 0.001). These data indicate that Essence Extractor and active-learning are strong tools for early model refinement.

### ABEL extracts neurophysiologically distinct behaviors with tight frame accuracy

One of the primary benefits of a detailed characterization of ethologically relevant behaviors is linking behavior to underlying neuronal activity. As a biological validation, we used fiber photometry recordings of calcium signal in the BNST and CeA recorded during the Novelty Suppressed Feeding assay (NSF; **Fig. 4A**). These structures have previously been implicated in approach and consummatory behaviors, including during approaches to high-fat diet (HFD) pellets during NSF, making them a strong positive biological control for ABEL-derived behavior labels^13,14^.

**Figure 4.**
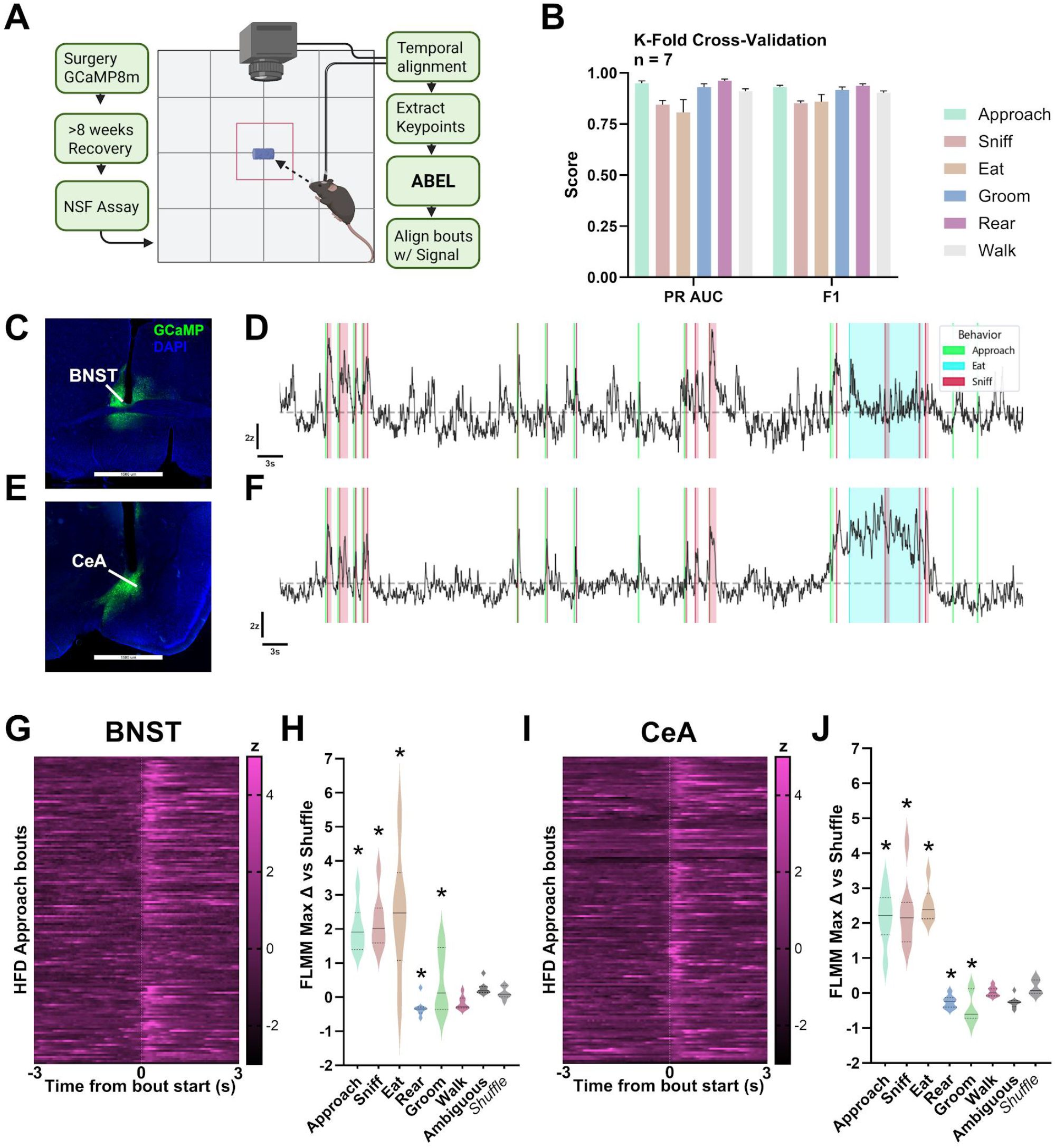
ABEL streamlines linking brain function with behavior. **(A)** Experiment timeline. Mice (n = 7 males) received unilateral GCaMP and fiber optic implants in the bed nucleus of the stria terminalis (BNST) and central nucleus of the amygdala (CeA) to enable simultaneous recording of calcium activity in two brain structures. After 8 weeks of recovery, mice were administered the novelty suppressed feeding (NSF) assay with fiber photometry recordings synced to an overhead camera. Keypoints were extracted using DeepLabCut (DLC). Video and keypoint data were used to train models in ABEL and bout start frames were exported for alignment with photometry signal. **(B)** Leave-one-subject-out (i.e., k-fold cross-validation) validation of model precision and recall across an array of behaviors including approach, sniff, eat, groom, rear, and walk. **(C, E)** GCaMP expression and fiber placement in the BNST and CeA. **(D, F)** Representative z-scored calcium signal in BNST and CeA overlaid with ABEL-derived bout frame data. Bout labels were restricted to approach, sniff, and eat for visual clarity. **(G)** BNST calcium signal aligned to HFD approach bouts. **(H)** Functional Linear Mixed Model (FLMM) analysis of BNST bout traces compared to pseudorandom shuffle bout start frames. Plotted data represent approximate GLS contrasts for per-subject peaks. *Significantly different from shuffle. **(I)** CeA calcium signal aligned to HFD approach bouts. **(J)** FLMM analysis of CeA bout traces compared to pseudorandom shuffle bout start frames.

Using a small NSF dataset and ∼2 hours of human effort training time, ABEL achieved high-precision models for a range of ethologically relevant behaviors, including approach, sniff, eat, rear, groom, and walk (**Fig. 4B**; F1 = 0.90 ± 0.03, PR-AUC, 0.90, ± 0.02). Bout start and end frames for food pellet approaches were then exported from ABEL and aligned with fiber photometry recordings from the BNST and CeA using open-source software (**Fig. 4C-F**; https://github.com/JobeRitchie/TRACY-Photometry-Processing-Suite). Calcium signals in the BNST and CeA were tightly associated with the start of approach behavior (**Fig. 4G, 4I**; Approach onset vs peak delta z-score/s, p < 0.001, r = 0.48). To assess if the behavior frame estimates generated with ABEL represent neurologically distinct event windows, we conducted functional linear mixed model analysis^15^ (FLMM). This analysis revealed that ABEL detects behaviors that correspond to distinct neural states. Neuronal dynamics in both the BNST and CeA during approach, sniff, groom, eat, and rear significantly differed from dynamics during randomly drawn frame windows (**Fig. 4H, 4J**; approximated peak difference in *z* is shown, all p < 0.01). In contrast, periods of ambiguous behavior were not separable from random windows in either region (ambiguous vs random, p > 0.05). Notably, the significance windows in CeA for both rear and approach begin within one to two frames from behavior bout start frame (33-67 ms at 30Hz) while the eat significance window precedes the start of eating for both regions by around 30-40 frames. Together, these data suggest that behaviors characterized by ABEL are both representative of underlying neuronal dynamics and highly frame-precise.

## Discussion

Advances in machine learning have enabled deep characterization of behavior with higher throughput than previously possible. There are many effective tools available for training predictive models for detecting behavior from video, spanning a range of unsupervised and supervised approaches. Here, we present ABEL (Active-learning Behavior Estimation and Labeling) which uses a supervised clip-based active-learning^6^ loop and an array of user-friendly tools for model refinement and visualization (**Fig. 1**). We rigorously validated ABEL against prior human labeling of published data, human labeling of novel data, and field-standard tools. ABEL predictions on held-out data were strongly correlated with human predictions and select analyses by both SimBA and Ethovision across an eclectic range of rodent behaviors spanning eight included assays (**Fig. 2**). Next, we demonstrate that the precision and accuracy of ABEL is made possible by its robust set of multi-modal features. Specifically, ABEL differentiates similar behaviors where pose-only behavior estimation would fail by incorporating contextual and temporal information (**Fig. 3**). Lastly, we investigated if ABEL behavior estimations are related to underlying neuronal dynamics. Calcium dynamics in the BNST and CeA temporally paired with behaviors in the NSF assay was used as an exemplar case based on the established role of the extended amygdala in approach and consummatory behaviors.^13,14^ ABEL behavior predictions were highly frame-precise and represented distinct neuronal signatures for approach, sniffing, eating, and grooming relative to random shuffle control (**Fig. 4**). These data support the conclusion that ABEL is an effective tool for estimating behavior at a level comparable to expert human observation for the behaviors tested.

ABEL builds on existing tools through implementation of clip-level annotation and analysis, mutual inhibition, and multi-modal features. At the core of ABEL is the data-efficient active-learning approach demonstrated by A-SOiD, which surfaces uncertain frames for review to drive rapid improvement in model strength^6^. We adapted this frame-level approach to a clip-level structure that includes within-clip pose and video features that provide additional value for solving behaviors that share otherwise similar pose characteristics. Unsupervised approaches, such as MoSeq, provide deep characterization of behavior at the level of sub-second behavior motifs. Where the supervised approach of ABEL excels is in labeling ethologically relevant outputs with clear behavior labels, aiding in interpretability and face validity. To facilitate this labeling effort without the use of additional software, ABEL includes a built-in interface that enables rapid annotation and clip review. An important caveat is that supervised approaches cannot detect novel behaviors not anticipated by the user. ABEL therefore provides built-in tools for unsupervised analysis in parallel to the supervised workflow: an unsupervised UMAP Selection interface, transition-probability and motif-discovery analytics. These tools address the limitation of the supervised loop by surfacing unanticipated candidates that can then be named and analyzed.

ABEL’s clearest advantage over existing tools is the efficient discovery of low-prevalence behaviors. Existing tools that require annotating entire videos necessitate an intensive human labeling effort to develop strong predictive models^5^. The tools built into ABEL – namely UMAP Interactive Selection and Essence Extractor – reduce human effort by several-fold relative to annotation of random clips or whole videos. This is achieved by surfacing clips from the unlabeled population that align with a selection of positive exemplar clips in feature space. To our knowledge, Essence Extractor has no direct equivalent in existing platforms and reduces labeling effort for low-prevalence behaviors by 1.85x relative to active-learning. However, these tools are meant to function as supplements to the already efficient active-learning approach^6^. The net result is the ability to rapidly develop strong models independent of behavior scarcity.

Behavior estimation in ABEL maps to underlying neuronal dynamics in the CeA and BNST. Calcium dynamics in the CeA and BNST are well-characterized for their role in approach and consummatory behaviors across numerous publications^13,14^, making their signal a strong positive biological validation for behavior labels in the

NSF assay. We observed that behavior bout frame estimation on held-out data yielded biologically distinct calcium dynamics relative to randomized bout traces. Specifically, food directed behaviors (approach, sniff, eat) were associated with a robust increase in CeA and BNST calcium dynamics relative to random labeling or ambiguous behavior. Interestingly, the calcium shift from bouts of eating preceded the start of eating behavior by over a second, while approach and sniff behaviors were temporally aligned to underlying neuronal activity with frame-level precision. This is not surprising given the increased calcium signal associated with other food-directed behaviors that typically precede eating, such as sniffing, as we have shown here and in prior reports^13,14^. As such, the inability to differentiate signals specific to eating is a limitation of our analysis. By contrast, rearing and grooming behaviors were associated with decreased calcium signal depending on brain region, suggesting a brain region- and behavior-specific activity profile.

## Limitations

Our validation and implementation of ABEL has limitations that should be addressed. Inherent in the use of supervised machine learning is a limitation in the discovery of novel behaviors. By design, supervised approaches cannot detect behaviors that the user did not anticipate. ABEL mitigates this by providing unsupervised tools that run parallel to the supervised workflow, but behavior labeling burden still lies on the individual scorer. Ground truth observation for model development was provided by a single annotator (JLR) which could limit validity or generalizability of the models as presented. However, rigorous validation of ABEL against other expert observers indicated a high degree of reliability. Another potential limitation is the fixed 0.5-second segmentation used for all models. The single time frame for segmentation was used to maintain consistency for validation while also being sufficient to accommodate the length of behaviors presented here. This introduces the caveat that a single time frame could miss features that occur on longer time scales. Another limitation is our assessment of human labeling effort. Effort comparisons were based on approximate human labeling time and this may be biased in favor of demonstrating a reduction in labeling time with ABEL. The 4 s per clip and 1x real-time for whole-video scan assumptions were chosen conservatively and meant to provide a point of reference as opposed to representing real-world annotation time. However, the fold differences in labeling effort scale with these assumptions, limiting any overt bias. In practice, timestamps from the review queue across projects show a median of 1 s per clip decision (43431 decisions). Finally, our biological validation has a limited scope. Calcium dynamics were assessed in only two structures and a limited sample size. It is possible that investigation of other brain circuits or behaviors using ABEL would produce less tightly aligned bout metrics.

## Methods

### Hardware

ABEL is designed to be run locally on a Windows operating system. All computation and analysis presented here was performed on an ASUS laptop with Intel Core i9-13980HX CPU, 64GB physical memory, and an NVIDIA 4070 laptop GPU. CPU fallback is available but will require considerably longer time for computation.

### Subjects

Adult male and female C57BL6/J mice (Stock #: 000664, Jackson Laboratories), Wistar rats (Charles River), and Long-Evans rats (Inotiv/Envigo) were housed in polycarbonate cages (GM500, Techniplast) under a 12:12 reverse dark-light cycle (lights off at 7:00 A.M.). Housing rooms in the vivarium were temperature- and humidity-controlled. Rodents had *ad libitum* access to food (Prolab Isopro RMH 3000, LabDiet) and water unless otherwise stated. All experiments were approved by the UNC School of Medicine Institutional Animal Care and Use Committee (IACUC) and in accordance with the NIH guidelines for the care and use of laboratory animals.

### Surgery

Mice received stereotaxic surgery under isoflurane anesthesia. Unilateral injections of AAV8-hSyn-GCaMP8f were administered to two opposite-hemisphere target structures consecutively (BNST: 0.95 ML, 0.3 AP, -4.3 DV; CeA: 2.8 ML, -1.2 AP, -4.7 DV). Optic fibers were then lowered to the same coordinates and fixed in place with metabond. Mice were provided post-operative carprofen (5 mg/kg) and allowed to recover from surgery for >8 weeks prior to testing.

**Table 1.** Included Behavioral Assays. Detailed assay protocols are included in Supplemental Methods.

| Assay | Subjects | Behaviors | Video | Duration | PR-AUC mean | Kappa mean |
| --- | --- | --- | --- | --- | --- | --- |
| Home Cage | 31 mice | Rear, groom, dig, wet dog shake | Logitech 640x480 | 30 min | 0.923 | 0.805 |
| Elevated Plus Maze | 56 mice | Head dip, protected head dip, rear, groom, stretch attend | Logitech 640x480 | 10 min | 0.926 | 0.834 |
| 3-chamber Social | 11 mice | Left cup: sniff, climb<br>Right cup: sniff, climb<br>Else: rear, groom | Logitech 640x480 | 30 min | 0.852 | 0.694 |
| Novel Object Recognition | 20 mice | Familiar: approach, sniff<br>Novel: approach, sniff<br>Else: rear, groom, walk | Logitech 640x480 | 30 min | 0.871 | 0.777 |
| Novelty Suppressed Feeding | 76 mice | Approach, sniff, eat, rear, groom, freeze, walk | Logitech 640x480 | 10 min | 0.894 | 0.787 |
| Open Field Test | 40 mice | Rear, groom, walk, freeze | Logitech 640x480 | 30 min | 0.970 | 0.891 |
| Predator odor assay | 29 rats | Dig, rear, groom, immobility, explore, wet dog shake, sniff tmt | Panasonic 1200x530 | 10 min | 0.842 | 0.714 |
| Fear conditioning | 58 mice | Freeze, shocked, rear, groom, explore | MedAssoc. 320x240 | 23 min | 0.956 | 0.881 |
| All assays | 321 animals | 45 behaviors |  |  | 0.896 | 0.788 |

### Fiber Photometry

Fiber photometry analysis was conducted in freely-available software using field-standard normalization and motion-correction techniques (Tracy; available at https://github.com/JobeRitchie/TRACY-Photometry-Processing-Suite). Photometry was acquired with alternating 470 nm and 415 nm channels at a sampling rate of 30 Hz per channel. Traces were deinterleaved and corrected for photobleaching by fitting a biexponential function. Motion artifacts were removed by regressing the 415 nm ΔF/F onto the 470 nm ΔF/F using robust Huber regression and subtracting the fitted isosbestic component. Corrected traces were z-scored across session then lowpass filtered with zero-phase second-order Butterworth filter (6 Hz cutoff). Photometry was aligned to behavior using nearest-match computer time stamp at start of recording. Behavior-aligned traces were based on bout start frames exported from ABEL and the 5 seconds before and after the bout start (± 150 frames) were used for bout analysis. Overlapping bouts were excluded. Session length was truncated to 10 minutes.

### Pose prediction

Pose tracking was conducted in DeepLabCut^1^ using custom-trained models per assay. Multiple models were trained to account for assay-specific camera angle and lighting. All models were trained on a DEKR_w18 backbone through 200 epochs (best snapshot used for analysis) and validated by visualizing adequate body-part tracking with confidence above 0.60 on labeled videos. Some videos were tracked using SLEAP^2^ for the purpose of developing a seamless integration workflow, but they were not used for ABEL model generation other than internal testing.

### Clip Segmentation and Feature Extraction

Raw data were segmented using single fixed-scale windowing of 0.5 seconds with a stride of either 0.1 or 0.5 seconds. Pose data was filtered to a minimum confidence threshold of 0.2, gaps of 10 frames or less were filled using linear interpolation, then a 3-frame rolling average was applied to remove jitter. Per-frame features extracted differed between projects (1952-2374 feature range, of which 512 are R3D-18 appearance embeddings) depending on available keypoint annotation and user-defined ROIs. For a complete list of features, see **Supplemental Table 1**.

### Annotation

The three seeding modes include dispersed pseudorandom, UMAP Interactive Selection, and Essence Extractor. Pseudorandom clip generation provides clips dispersed across subjects and sessions through seeded random number generation with optional per-session cap. For UMAP Interactive Selection, a UMAP is fit on labeled and predicted points and default settings of 2 components, an adaptive neighborhood size of max(5, min(25, √n)), minimum distance of 0.15, and a default to principal component analysis when a UMAP cannot be fit. Essence Extractor first surfaces features that separate user-defined exemplar clips from the unlabeled population using a rank-based statistical equivalent to absolute value of AUC-0.5. Features falling below 0.10 are discarded, and remaining candidate features are filtered to a candidate set (default: 5 criterion features, 8 ranking features). The displayed criterion features summarize what separates exemplar clips. Clip ranking is performed by a sparse linear model. Features are ranked by |AUC – 0.5|, the top 300 are retained, then an L1-penalized logistic regression (liblinear, C = 0.1, class balanced, on median-imputed and standardized features) is fit to discriminate the exemplar clips from a background sample of 1500 unlabeled clips.

### Active-learning machine learning

Active-learning candidates are scored by a single blended priority. Each unlabeled clip receives a rank score combining its behavior-aware uncertainty, predicted probability for target behavior (weight 0.5), reviewer feedback (0.2), and exclusivity uncertainty (0.2). Clips previously reviewed as negatives that score above 0.6 receive an additional +0.3 hard-negative bonus, surfacing most confident errors. No automatic stopping criterion is set. Batch size is defaulted to 20 but is adjusted per project. Positive augmentation is on by default and includes: Gaussian jitter at σ = 0.05 of each feature’s standard deviation, 10% feature dropout, 3 synthetic copies per positive clip. Model generation required approximately 2-5 hours of human effort time per assay (3-7 behaviors per assay). For comparing clip discovery strategies, active-learning ranked candidate clips by target-class alone so that all four strategies were compared evenly.

### Classifier

Behavior classifiers were trained using the Extreme Gradient Boosted (XGBoost) machine learning library. One binary model was trained per behavior (one-vs-rest) with Platt-scaled calibration. Models were trained on CUDA with a CPU fallback. Base hyperparameters: tree_method “hist”, max_depth 6, learning rate 0.1, n_estimators 300, subsample 0.8, colsample_bytree 0.8, reg_alpha 0.1, reg_lambda 1.0, min_child_weight 5, random_state 42. Adaptive complexity is disabled by default, but will further scale n_estimators and max_depth by the ratio of positive clips to features. Versions used: xgboost 3.2.0 and scikit-learn 1.7.2. ABEL additionally extracts deep video embeddings using pretrained R3D-18 3D convolutional network and includes these 512-dimensional appearance features in the XGBoost feature matrix as ordinary numeric features.

### Temporal refinement

Once satisfactory models have been generated, they are applied to a newly segmented dataset with high degree of clip overlap (default set to 0.1s stride). Per-behavior frame-level confidence traces are generated by finding the per-frame average of the densely overlapping clip-level predictions. At this stage, mutual inhibition can be applied to specific behavior pairs or with a global setting (default of 0.2) to aid in differentiating confounding behaviors. Next, per-behavior minimum confidence and bout length thresholds are set to establish positive bout for subsequent analysis. Default settings: onset threshold 0.50, minimum bout duration 6 frames, merge gap 3 frames, smoothing window 5, and a 1.5 s warmup (excludes unreliable frames at start of estimation). Tools for flagging false positives and false negatives are available for further refinement.

### Functional linear mixed model analysis

Neural correlates of behavior were assessed using calcium dynamics time-locked to behavior onset ± 5 seconds. Statistical comparison of bout associated traces was conducted using functional linear mixed model fitted by fast univariate inference^15,18^. A linear mixed model with a random intercept per subject was fitted independently at each timepoint. Behaviors were fitted in a single factor model per brain region (BNST, CeA). The reference point was calcium dynamics during random frame windows using pseudorandom bout start frames. Significance over a window was defined by a joint 95% confidence band. Approximate per-subject peak effects were computed from a subset-clustered generalized-least-squares covariance of the coefficient function. Models were fitted with fastFMM 1.0.1 (fui in R 4.6.0).

### Statistical Analysis

All analyses were conducted in GraphPad Prism (11.01) or with statistical libraries in Python 3.11 (NumPy 1.26, SciPy 1.11, pandas 2.1). Model performance is reported as the mean across five random seeds trained on a 75% training split with a 25% held-out test split (video session as unit). The training pool was rebalanced to a 5:1 negative-to-positive class ratio; held-out sessions were evaluated on all reviewed labeled clips without rebalancing. The primary model performance metric is precision-recall area under the curve (PR-AUC) with F1 score as a secondary metric. Agreement between observers was quantified with Lin’s concordance correlation coefficient (CCC) and Bland-Altman limits of agreement, in addition to Pearson correlation. Equivalence was tested using two one-sided tests against a specified margin (TOST). Feature family ablation (**Fig. 3F**) was assessed per behavior averaged across random seeds with two-tailed paired t-tests on ΔPR-AUC between the full feature set and the ablated feature set (pose only). Benjamini-Hochberg (BH-FDR) corrected p-values were used to control false discovery rate (q < 0.05 across 45 behaviors). Behavior pair discrimination (**Fig. 3G**) was assessed using two-tailed paired t-tests with BH-FDR correction. Clip discovery strategies were analyzed by two-way repeated-measures ANOVA on log2-transformed discovery-clip counts with strategy and clip budget as within-subjects factors and behavior as the repeating unit; *post hoc* comparisons were conducted at pre-determined clip budgets (25 and 300). Annotation effort in human labeling time was derived by conservatively assuming 4 seconds per clip and analyzed using one-way repeated-measures ANOVA on log10-transformed minutes with strategy as the within-behavior factor (n = 45 behaviors), Greenhouse-Geisser corrected with Holm-corrected paired *post hoc* contrasts. The whole-video comparator was charged the mean of real time video on randomly selected videos simulated across 4000 video orders. Prevalence-performance associations (**Fig. 3B**) were tested by Pearson correlation on log10 prevalence. Effects are reported as fold differences after back transformation. Alpha was set to 0.05.

## Code and Data Availability

ABEL is open-source and freely available for academic use at https://github.com/JobeRitchie/ABEL.git. The version used for analyses reported here is v0.11.0. Data is available from the corresponding author upon reasonable request.

## Large Language Model (LLM) Disclosure

Claude Code (Opus 5) was used to aid in coding and to verify statistical reporting. All manuscript drafts were written by the authors and contain no LLM-derived text. The authors assume full responsibility for the accuracy of data presented here.

## Acknowledgements

This work was supported by National Institutes of Health’s (NIH) R01AA022048, T32AA007573, T32NS007431, K12GM000678 and AA026537.

