## Supplemental Methods for "ABEL: an active-learning behavior estimation and labeling platform"

### Home cage behavior assay

Mice were recorded in the home cage in a well-lit sound attenuated box (300 lux) for 30 minutes. Nesting materials were removed from the home cage prior to testing. Behavior was recorded from above with a Logitech webcam.

### Elevated plus maze

Mice were placed on a standard elevated plus maze with their head oriented towards the closed arms. The test continued for 10 minutes. Behavior was recorded from above with a Logitech webcam.

### 3-chamber social novelty

The 3-chamber social novelty assay took place in a chambered arena with one wire containment cup in each of the side chambers. Testing consisted of a 30-minute session broken up into 10-minute blocks. For the first block, the wire cups were empty (habituation). For the second block, one same-sex adolescent (5-6 weeks) mouse and one 3D printed faux mouse were added to separate containment cups. For the third block, the faux mouse was removed and a second, novel, same sex adolescent mouse was added in its place. Behavior was recorded from above with a Logitech webcam.

### Novel object recognition

Novel object recognition testing consisted of two 30-minute sessions split 3 hours apart. During the first session, mice were placed in an arena with two objects and allowed to explore. During the second session, 24 hrs later, mice were placed in the same arena but one of the objects has been swapped for a novel object. Mice are scored based on time spent with the novel object or familiar object. Behavior was recorded from above with a Logitech webcam.

### Novelty suppressed feeding

The novelty suppressed feeding assay was conducted in an open arena with a high fat diet (HFD) pellet taped in the center of the arena. Lux was set at 400 and behaviors were recorded from overhead. Mice were habituated to HFD two days prior to testing. Food restriction was applied for two hours prior to testing. Mice were allowed to explore the arena for 10 minutes. Behavior was recorded from above with a Logitech webcam.

### Open Field Test

Mice were placed in the corner of an empty open arena (51 cm x 51 cm) and allowed to explore for 30 minutes. Behavior was recorded from above with a Logitech webcam.

### Predator odor assay

TMT exposures were similar to previously established methods[16,17]. Chambers contained ~130 g of ALPHA-dri ® bedding that was evenly dispersed along the bottom of the chamber. A metal basket containing a piece of filter paper was hung on the top of the right side of the wall, and the filter paper was not accessible to the rats at any point during the experiment. Rats were placed in the chamber and TMT (10 μLL; 2,3,5-Trimethyl-3-Thiazoline, purity ≥97.0%, BioSRQ) was applied to the filter paper. The 10-minute exposures were video recorded, and behaviors were analyzed by expert observers using ANY-maze Video Tracking Software, with the primary two behaviors evaluated being digging and immobility. Digging was defined as rapid movement of the two front paws to push the bedding towards the metal basket or burrowing of the head under the bedding. Immobility was quantified as a lack of movement for greater than 2 seconds.

**Fear Conditioning**

Mice received trace fear conditioning in a sound attenuated chamber (Med Associates; grid floor, cleaned with 20% ethanol +1% vanilla solution). Conditioning consisted of five tone shock pairings with a 20-second trace separating the tone from shock. Behavior was recorded from above.
