## Supplemental Table for "ABEL: an active-learning behavior estimation and labeling platform"

| feature | modality | 3CS | EPM | FC |
|---|---|---|---|---|
| back_left_paw_acceleration_energy | kinematics | 0 | 0 | 0 |
| back_left_paw_acceleration_max | kinematics | 0 | 0 | 0 |
| back_left_paw_acceleration_mean | kinematics | 0 | 0 | 0 |
| back_left_paw_acceleration_median | kinematics | 0 | 0 | 0 |
| back_left_paw_acceleration_p10 | kinematics | 0 | 0 | 0 |
| back_left_paw_acceleration_p90 | kinematics | 0 | 0 | 0 |
| back_left_paw_acceleration_periodicity | kinematics | 0 | 0 | 0 |
| back_left_paw_acceleration_std | kinematics | 0 | 0 | 0 |
| back_left_paw_forward_velocity_energy | kinematics | 0 | 0 | 0 |
| back_left_paw_forward_velocity_max | kinematics | 0 | 0 | 0 |
| back_left_paw_forward_velocity_mean | kinematics | 0 | 0 | 0 |
| back_left_paw_forward_velocity_median | kinematics | 0 | 0 | 0 |
| back_left_paw_forward_velocity_p10 | kinematics | 0 | 0 | 0 |
| back_left_paw_forward_velocity_p90 | kinematics | 0 | 0 | 0 |
| back_left_paw_forward_velocity_periodicity | kinematics | 0 | 0 | 0 |
| back_left_paw_forward_velocity_std | kinematics | 0 | 0 | 0 |
| back_left_paw_jerk_energy | kinematics | 0 | 0 | 0 |
| back_left_paw_jerk_max | kinematics | 0 | 0 | 0 |
| back_left_paw_jerk_mean | kinematics | 0 | 0 | 0 |
| back_left_paw_jerk_median | kinematics | 0 | 0 | 0 |
| back_left_paw_jerk_p10 | kinematics | 0 | 0 | 0 |
| back_left_paw_jerk_p90 | kinematics | 0 | 0 | 0 |
| back_left_paw_jerk_periodicity | kinematics | 0 | 0 | 0 |
| back_left_paw_jerk_std | kinematics | 0 | 0 | 0 |
| back_left_paw_lateral_velocity_energy | kinematics | 0 | 0 | 0 |
| back_left_paw_lateral_velocity_max | kinematics | 0 | 0 | 0 |
| back_left_paw_lateral_velocity_mean | kinematics | 0 | 0 | 0 |
| back_left_paw_lateral_velocity_median | kinematics | 0 | 0 | 0 |
| back_left_paw_lateral_velocity_p10 | kinematics | 0 | 0 | 0 |
| back_left_paw_lateral_velocity_p90 | kinematics | 0 | 0 | 0 |
| back_left_paw_lateral_velocity_periodicity | kinematics | 0 | 0 | 0 |
| back_left_paw_lateral_velocity_std | kinematics | 0 | 0 | 0 |
| back_left_paw_speed_energy | kinematics | 0 | 0 | 0 |
| back_left_paw_speed_max | kinematics | 0 | 0 | 0 |
| back_left_paw_speed_mean | kinematics | 0 | 0 | 0 |
| back_left_paw_speed_median | kinematics | 0 | 0 | 0 |
| back_left_paw_speed_p10 | kinematics | 0 | 0 | 0 |
| back_left_paw_speed_p90 | kinematics | 0 | 0 | 0 |
| back_left_paw_speed_periodicity | kinematics | 0 | 0 | 0 |
| back_left_paw_speed_std | kinematics | 0 | 0 | 0 |
| back_left_paw_velocity_x_energy | kinematics | 0 | 0 | 0 |
| back_left_paw_velocity_x_max | kinematics | 0 | 0 | 0 |
| back_left_paw_velocity_x_mean | kinematics | 0 | 0 | 0 |

| | | | | |
|---|---|---|---|---|
| back_left_paw_velocity_x_median | kinematics | 0 | 0 | 0 |
| back_left_paw_velocity_x_p10 | kinematics | 0 | 0 | 0 |
| back_left_paw_velocity_x_p90 | kinematics | 0 | 0 | 0 |
| back_left_paw_velocity_x_periodicity | kinematics | 0 | 0 | 0 |
| back_left_paw_velocity_x_std | kinematics | 0 | 0 | 0 |
| back_left_paw_velocity_y_energy | kinematics | 0 | 0 | 0 |
| back_left_paw_velocity_y_max | kinematics | 0 | 0 | 0 |
| back_left_paw_velocity_y_mean | kinematics | 0 | 0 | 0 |
| back_left_paw_velocity_y_median | kinematics | 0 | 0 | 0 |
| back_left_paw_velocity_y_p10 | kinematics | 0 | 0 | 0 |
| back_left_paw_velocity_y_p90 | kinematics | 0 | 0 | 0 |
| back_left_paw_velocity_y_periodicity | kinematics | 0 | 0 | 0 |
| back_left_paw_velocity_y_std | kinematics | 0 | 0 | 0 |
| back_right_paw_acceleration_energy | kinematics | 0 | 0 | 0 |
| back_right_paw_acceleration_max | kinematics | 0 | 0 | 0 |
| back_right_paw_acceleration_mean | kinematics | 0 | 0 | 0 |
| back_right_paw_acceleration_median | kinematics | 0 | 0 | 0 |
| back_right_paw_acceleration_p10 | kinematics | 0 | 0 | 0 |
| back_right_paw_acceleration_p90 | kinematics | 0 | 0 | 0 |
| back_right_paw_acceleration_periodicity | kinematics | 0 | 0 | 0 |
| back_right_paw_acceleration_std | kinematics | 0 | 0 | 0 |
| back_right_paw_forward_velocity_energy | kinematics | 0 | 0 | 0 |
| back_right_paw_forward_velocity_max | kinematics | 0 | 0 | 0 |
| back_right_paw_forward_velocity_mean | kinematics | 0 | 0 | 0 |
| back_right_paw_forward_velocity_median | kinematics | 0 | 0 | 0 |
| back_right_paw_forward_velocity_p10 | kinematics | 0 | 0 | 0 |
| back_right_paw_forward_velocity_p90 | kinematics | 0 | 0 | 0 |
| back_right_paw_forward_velocity_periodicity | kinematics | 0 | 0 | 0 |
| back_right_paw_forward_velocity_std | kinematics | 0 | 0 | 0 |
| back_right_paw_jerk_energy | kinematics | 0 | 0 | 0 |
| back_right_paw_jerk_max | kinematics | 0 | 0 | 0 |
| back_right_paw_jerk_mean | kinematics | 0 | 0 | 0 |
| back_right_paw_jerk_median | kinematics | 0 | 0 | 0 |
| back_right_paw_jerk_p10 | kinematics | 0 | 0 | 0 |
| back_right_paw_jerk_p90 | kinematics | 0 | 0 | 0 |
| back_right_paw_jerk_periodicity | kinematics | 0 | 0 | 0 |
| back_right_paw_jerk_std | kinematics | 0 | 0 | 0 |
| back_right_paw_lateral_velocity_energy | kinematics | 0 | 0 | 0 |
| back_right_paw_lateral_velocity_max | kinematics | 0 | 0 | 0 |
| back_right_paw_lateral_velocity_mean | kinematics | 0 | 0 | 0 |
| back_right_paw_lateral_velocity_median | kinematics | 0 | 0 | 0 |
| back_right_paw_lateral_velocity_p10 | kinematics | 0 | 0 | 0 |
| back_right_paw_lateral_velocity_p90 | kinematics | 0 | 0 | 0 |
| back_right_paw_lateral_velocity_periodicity | kinematics | 0 | 0 | 0 |

| | | | | |
|---|---|---|---|---|
| back_right_paw_lateral_velocity_std | kinematics | 0 | 0 | 0 |
| back_right_paw_speed_energy | kinematics | 0 | 0 | 0 |
| back_right_paw_speed_max | kinematics | 0 | 0 | 0 |
| back_right_paw_speed_mean | kinematics | 0 | 0 | 0 |
| back_right_paw_speed_median | kinematics | 0 | 0 | 0 |
| back_right_paw_speed_p10 | kinematics | 0 | 0 | 0 |
| back_right_paw_speed_p90 | kinematics | 0 | 0 | 0 |
| back_right_paw_speed_periodicity | kinematics | 0 | 0 | 0 |
| back_right_paw_speed_std | kinematics | 0 | 0 | 0 |
| back_right_paw_velocity_x_energy | kinematics | 0 | 0 | 0 |
| back_right_paw_velocity_x_max | kinematics | 0 | 0 | 0 |
| back_right_paw_velocity_x_mean | kinematics | 0 | 0 | 0 |
| back_right_paw_velocity_x_median | kinematics | 0 | 0 | 0 |
| back_right_paw_velocity_x_p10 | kinematics | 0 | 0 | 0 |
| back_right_paw_velocity_x_p90 | kinematics | 0 | 0 | 0 |
| back_right_paw_velocity_x_periodicity | kinematics | 0 | 0 | 0 |
| back_right_paw_velocity_x_std | kinematics | 0 | 0 | 0 |
| back_right_paw_velocity_y_energy | kinematics | 0 | 0 | 0 |
| back_right_paw_velocity_y_max | kinematics | 0 | 0 | 0 |
| back_right_paw_velocity_y_mean | kinematics | 0 | 0 | 0 |
| back_right_paw_velocity_y_median | kinematics | 0 | 0 | 0 |
| back_right_paw_velocity_y_p10 | kinematics | 0 | 0 | 0 |
| back_right_paw_velocity_y_p90 | kinematics | 0 | 0 | 0 |
| back_right_paw_velocity_y_periodicity | kinematics | 0 | 0 | 0 |
| back_right_paw_velocity_y_std | kinematics | 0 | 0 | 0 |
| body_angle_to_roi_1_delta | context (ROI/target) | 1 | 1 | 1 |
| body_angle_to_roi_1_energy | context (ROI/target) | 1 | 1 | 1 |
| body_angle_to_roi_1_max | context (ROI/target) | 1 | 1 | 1 |
| body_angle_to_roi_1_mean | context (ROI/target) | 1 | 1 | 1 |
| body_angle_to_roi_1_median | context (ROI/target) | 1 | 1 | 1 |
| body_angle_to_roi_1_p10 | context (ROI/target) | 1 | 1 | 1 |
| body_angle_to_roi_1_p90 | context (ROI/target) | 1 | 1 | 1 |
| body_angle_to_roi_1_periodicity | context (ROI/target) | 1 | 1 | 1 |
| body_angle_to_roi_1_std | context (ROI/target) | 1 | 1 | 1 |
| body_angle_to_roi_1_trend | context (ROI/target) | 1 | 1 | 1 |
| body_angle_to_roi_2_delta | context (ROI/target) | 1 | 0 | 0 |
| body_angle_to_roi_2_energy | context (ROI/target) | 1 | 0 | 0 |
| body_angle_to_roi_2_max | context (ROI/target) | 1 | 0 | 0 |
| body_angle_to_roi_2_mean | context (ROI/target) | 1 | 0 | 0 |
| body_angle_to_roi_2_median | context (ROI/target) | 1 | 0 | 0 |
| body_angle_to_roi_2_p10 | context (ROI/target) | 1 | 0 | 0 |
| body_angle_to_roi_2_p90 | context (ROI/target) | 1 | 0 | 0 |
| body_angle_to_roi_2_periodicity | context (ROI/target) | 1 | 0 | 0 |
| body_angle_to_roi_2_std | context (ROI/target) | 1 | 0 | 0 |

| | | | | |
|---|---|---|---|---|
| body_angle_to_roi_2_trend | context (ROI/target) | 1 | 0 | 0 |
| body_angle_to_target_delta | context (ROI/target) | 1 | 1 | 1 |
| body_angle_to_target_energy | context (ROI/target) | 1 | 1 | 1 |
| body_angle_to_target_max | context (ROI/target) | 1 | 1 | 1 |
| body_angle_to_target_mean | context (ROI/target) | 1 | 1 | 1 |
| body_angle_to_target_median | context (ROI/target) | 1 | 1 | 1 |
| body_angle_to_target_p10 | context (ROI/target) | 1 | 1 | 1 |
| body_angle_to_target_p90 | context (ROI/target) | 1 | 1 | 1 |
| body_angle_to_target_periodicity | context (ROI/target) | 1 | 1 | 1 |
| body_angle_to_target_std | context (ROI/target) | 1 | 1 | 1 |
| body_angle_to_target_trend | context (ROI/target) | 1 | 1 | 1 |
| body_centroid_roi_1_axial_abs_energy | context (ROI/target) | 1 | 1 | 1 |
| body_centroid_roi_1_axial_abs_max | context (ROI/target) | 1 | 1 | 1 |
| body_centroid_roi_1_axial_abs_mean | context (ROI/target) | 1 | 1 | 1 |
| body_centroid_roi_1_axial_abs_median | context (ROI/target) | 1 | 1 | 1 |
| body_centroid_roi_1_axial_abs_p10 | context (ROI/target) | 1 | 1 | 1 |
| body_centroid_roi_1_axial_abs_p90 | context (ROI/target) | 1 | 1 | 1 |
| body_centroid_roi_1_axial_abs_periodicity | context (ROI/target) | 1 | 1 | 1 |
| body_centroid_roi_1_axial_abs_std | context (ROI/target) | 1 | 1 | 1 |
| body_centroid_roi_1_axial_energy | context (ROI/target) | 1 | 1 | 1 |
| body_centroid_roi_1_axial_max | context (ROI/target) | 1 | 1 | 1 |
| body_centroid_roi_1_axial_mean | context (ROI/target) | 1 | 1 | 1 |
| body_centroid_roi_1_axial_median | context (ROI/target) | 1 | 1 | 1 |
| body_centroid_roi_1_axial_p10 | context (ROI/target) | 1 | 1 | 1 |
| body_centroid_roi_1_axial_p90 | context (ROI/target) | 1 | 1 | 1 |
| body_centroid_roi_1_axial_periodicity | context (ROI/target) | 1 | 1 | 1 |
| body_centroid_roi_1_axial_std | context (ROI/target) | 1 | 1 | 1 |
| body_centroid_roi_1_lateral_abs_energy | context (ROI/target) | 1 | 1 | 1 |
| body_centroid_roi_1_lateral_abs_max | context (ROI/target) | 1 | 1 | 1 |
| body_centroid_roi_1_lateral_abs_mean | context (ROI/target) | 1 | 1 | 1 |
| body_centroid_roi_1_lateral_abs_median | context (ROI/target) | 1 | 1 | 1 |
| body_centroid_roi_1_lateral_abs_p10 | context (ROI/target) | 1 | 1 | 1 |
| body_centroid_roi_1_lateral_abs_p90 | context (ROI/target) | 1 | 1 | 1 |
| body_centroid_roi_1_lateral_abs_periodicity | context (ROI/target) | 1 | 1 | 1 |
| body_centroid_roi_1_lateral_abs_std | context (ROI/target) | 1 | 1 | 1 |
| body_centroid_roi_1_lateral_energy | context (ROI/target) | 1 | 1 | 1 |
| body_centroid_roi_1_lateral_max | context (ROI/target) | 1 | 1 | 1 |
| body_centroid_roi_1_lateral_mean | context (ROI/target) | 1 | 1 | 1 |
| body_centroid_roi_1_lateral_median | context (ROI/target) | 1 | 1 | 1 |
| body_centroid_roi_1_lateral_p10 | context (ROI/target) | 1 | 1 | 1 |
| body_centroid_roi_1_lateral_p90 | context (ROI/target) | 1 | 1 | 1 |
| body_centroid_roi_1_lateral_periodicity | context (ROI/target) | 1 | 1 | 1 |
| body_centroid_roi_1_lateral_std | context (ROI/target) | 1 | 1 | 1 |
| body_centroid_roi_2_axial_abs_energy | context (ROI/target) | 1 | 0 | 0 |

| | | | | |
|---|---|---|---|---|
| body_centroid_roi_2_axial_abs_max | context (ROI/target) | 1 | 0 | 0 |
| body_centroid_roi_2_axial_abs_mean | context (ROI/target) | 1 | 0 | 0 |
| body_centroid_roi_2_axial_abs_median | context (ROI/target) | 1 | 0 | 0 |
| body_centroid_roi_2_axial_abs_p10 | context (ROI/target) | 1 | 0 | 0 |
| body_centroid_roi_2_axial_abs_p90 | context (ROI/target) | 1 | 0 | 0 |
| body_centroid_roi_2_axial_abs_periodicity | context (ROI/target) | 1 | 0 | 0 |
| body_centroid_roi_2_axial_abs_std | context (ROI/target) | 1 | 0 | 0 |
| body_centroid_roi_2_axial_energy | context (ROI/target) | 1 | 0 | 0 |
| body_centroid_roi_2_axial_max | context (ROI/target) | 1 | 0 | 0 |
| body_centroid_roi_2_axial_mean | context (ROI/target) | 1 | 0 | 0 |
| body_centroid_roi_2_axial_median | context (ROI/target) | 1 | 0 | 0 |
| body_centroid_roi_2_axial_p10 | context (ROI/target) | 1 | 0 | 0 |
| body_centroid_roi_2_axial_p90 | context (ROI/target) | 1 | 0 | 0 |
| body_centroid_roi_2_axial_periodicity | context (ROI/target) | 1 | 0 | 0 |
| body_centroid_roi_2_axial_std | context (ROI/target) | 1 | 0 | 0 |
| body_centroid_roi_2_lateral_abs_energy | context (ROI/target) | 1 | 0 | 0 |
| body_centroid_roi_2_lateral_abs_max | context (ROI/target) | 1 | 0 | 0 |
| body_centroid_roi_2_lateral_abs_mean | context (ROI/target) | 1 | 0 | 0 |
| body_centroid_roi_2_lateral_abs_median | context (ROI/target) | 1 | 0 | 0 |
| body_centroid_roi_2_lateral_abs_p10 | context (ROI/target) | 1 | 0 | 0 |
| body_centroid_roi_2_lateral_abs_p90 | context (ROI/target) | 1 | 0 | 0 |
| body_centroid_roi_2_lateral_abs_periodicity | context (ROI/target) | 1 | 0 | 0 |
| body_centroid_roi_2_lateral_abs_std | context (ROI/target) | 1 | 0 | 0 |
| body_centroid_roi_2_lateral_energy | context (ROI/target) | 1 | 0 | 0 |
| body_centroid_roi_2_lateral_max | context (ROI/target) | 1 | 0 | 0 |
| body_centroid_roi_2_lateral_mean | context (ROI/target) | 1 | 0 | 0 |
| body_centroid_roi_2_lateral_median | context (ROI/target) | 1 | 0 | 0 |
| body_centroid_roi_2_lateral_p10 | context (ROI/target) | 1 | 0 | 0 |
| body_centroid_roi_2_lateral_p90 | context (ROI/target) | 1 | 0 | 0 |
| body_centroid_roi_2_lateral_periodicity | context (ROI/target) | 1 | 0 | 0 |
| body_centroid_roi_2_lateral_std | context (ROI/target) | 1 | 0 | 0 |
| body_centroid_to_roi_1_corner_dist_delta | context (ROI/target) | 1 | 1 | 1 |
| body_centroid_to_roi_1_corner_dist_energy | context (ROI/target) | 1 | 1 | 1 |
| body_centroid_to_roi_1_corner_dist_max | context (ROI/target) | 1 | 1 | 1 |
| body_centroid_to_roi_1_corner_dist_mean | context (ROI/target) | 1 | 1 | 1 |
| body_centroid_to_roi_1_corner_dist_median | context (ROI/target) | 1 | 1 | 1 |
| body_centroid_to_roi_1_corner_dist_p10 | context (ROI/target) | 1 | 1 | 1 |
| body_centroid_to_roi_1_corner_dist_p90 | context (ROI/target) | 1 | 1 | 1 |
| body_centroid_to_roi_1_corner_dist_periodicity | context (ROI/target) | 1 | 1 | 1 |
| body_centroid_to_roi_1_corner_dist_std | context (ROI/target) | 1 | 1 | 1 |
| body_centroid_to_roi_1_corner_dist_trend | context (ROI/target) | 1 | 1 | 1 |
| body_centroid_to_roi_1_dist_delta | context (ROI/target) | 1 | 1 | 1 |
| body_centroid_to_roi_1_dist_energy | context (ROI/target) | 1 | 1 | 1 |
| body_centroid_to_roi_1_dist_max | context (ROI/target) | 1 | 1 | 1 |

| | | | | |
|---|---|---|---|---|
| body_centroid_to_roi_1_dist_mean | context (ROI/target) | 1 | 1 | 1 |
| body_centroid_to_roi_1_dist_median | context (ROI/target) | 1 | 1 | 1 |
| body_centroid_to_roi_1_dist_p10 | context (ROI/target) | 1 | 1 | 1 |
| body_centroid_to_roi_1_dist_p90 | context (ROI/target) | 1 | 1 | 1 |
| body_centroid_to_roi_1_dist_periodicity | context (ROI/target) | 1 | 1 | 1 |
| body_centroid_to_roi_1_dist_std | context (ROI/target) | 1 | 1 | 1 |
| body_centroid_to_roi_1_dist_trend | context (ROI/target) | 1 | 1 | 1 |
| body_centroid_to_roi_1_edge_dist_delta | context (ROI/target) | 1 | 1 | 1 |
| body_centroid_to_roi_1_edge_dist_energy | context (ROI/target) | 1 | 1 | 1 |
| body_centroid_to_roi_1_edge_dist_max | context (ROI/target) | 1 | 1 | 1 |
| body_centroid_to_roi_1_edge_dist_mean | context (ROI/target) | 1 | 1 | 1 |
| body_centroid_to_roi_1_edge_dist_median | context (ROI/target) | 1 | 1 | 1 |
| body_centroid_to_roi_1_edge_dist_p10 | context (ROI/target) | 1 | 1 | 1 |
| body_centroid_to_roi_1_edge_dist_p90 | context (ROI/target) | 1 | 1 | 1 |
| body_centroid_to_roi_1_edge_dist_periodicity | context (ROI/target) | 1 | 1 | 1 |
| body_centroid_to_roi_1_edge_dist_std | context (ROI/target) | 1 | 1 | 1 |
| body_centroid_to_roi_1_edge_dist_trend | context (ROI/target) | 1 | 1 | 1 |
| body_centroid_to_roi_1_signed_dist_delta | context (ROI/target) | 1 | 1 | 1 |
| body_centroid_to_roi_1_signed_dist_energy | context (ROI/target) | 1 | 1 | 1 |
| body_centroid_to_roi_1_signed_dist_max | context (ROI/target) | 1 | 1 | 1 |
| body_centroid_to_roi_1_signed_dist_mean | context (ROI/target) | 1 | 1 | 1 |
| body_centroid_to_roi_1_signed_dist_median | context (ROI/target) | 1 | 1 | 1 |
| body_centroid_to_roi_1_signed_dist_p10 | context (ROI/target) | 1 | 1 | 1 |
| body_centroid_to_roi_1_signed_dist_p90 | context (ROI/target) | 1 | 1 | 1 |
| body_centroid_to_roi_1_signed_dist_periodicity | context (ROI/target) | 1 | 1 | 1 |
| body_centroid_to_roi_1_signed_dist_std | context (ROI/target) | 1 | 1 | 1 |
| body_centroid_to_roi_1_signed_dist_trend | context (ROI/target) | 1 | 1 | 1 |
| body_centroid_to_roi_2_corner_dist_delta | context (ROI/target) | 1 | 0 | 0 |
| body_centroid_to_roi_2_corner_dist_energy | context (ROI/target) | 1 | 0 | 0 |
| body_centroid_to_roi_2_corner_dist_max | context (ROI/target) | 1 | 0 | 0 |
| body_centroid_to_roi_2_corner_dist_mean | context (ROI/target) | 1 | 0 | 0 |
| body_centroid_to_roi_2_corner_dist_median | context (ROI/target) | 1 | 0 | 0 |
| body_centroid_to_roi_2_corner_dist_p10 | context (ROI/target) | 1 | 0 | 0 |
| body_centroid_to_roi_2_corner_dist_p90 | context (ROI/target) | 1 | 0 | 0 |
| body_centroid_to_roi_2_corner_dist_periodicity | context (ROI/target) | 1 | 0 | 0 |
| body_centroid_to_roi_2_corner_dist_std | context (ROI/target) | 1 | 0 | 0 |
| body_centroid_to_roi_2_corner_dist_trend | context (ROI/target) | 1 | 0 | 0 |
| body_centroid_to_roi_2_dist_delta | context (ROI/target) | 1 | 0 | 0 |
| body_centroid_to_roi_2_dist_energy | context (ROI/target) | 1 | 0 | 0 |
| body_centroid_to_roi_2_dist_max | context (ROI/target) | 1 | 0 | 0 |
| body_centroid_to_roi_2_dist_mean | context (ROI/target) | 1 | 0 | 0 |
| body_centroid_to_roi_2_dist_median | context (ROI/target) | 1 | 0 | 0 |
| body_centroid_to_roi_2_dist_p10 | context (ROI/target) | 1 | 0 | 0 |
| body_centroid_to_roi_2_dist_p90 | context (ROI/target) | 1 | 0 | 0 |

| | | | | |
|---|---|---|---|---|
| body_centroid_to_roi_2_dist_periodicity | context (ROI/target) | 1 | 0 | 0 |
| body_centroid_to_roi_2_dist_std | context (ROI/target) | 1 | 0 | 0 |
| body_centroid_to_roi_2_dist_trend | context (ROI/target) | 1 | 0 | 0 |
| body_centroid_to_roi_2_edge_dist_delta | context (ROI/target) | 1 | 0 | 0 |
| body_centroid_to_roi_2_edge_dist_energy | context (ROI/target) | 1 | 0 | 0 |
| body_centroid_to_roi_2_edge_dist_max | context (ROI/target) | 1 | 0 | 0 |
| body_centroid_to_roi_2_edge_dist_mean | context (ROI/target) | 1 | 0 | 0 |
| body_centroid_to_roi_2_edge_dist_median | context (ROI/target) | 1 | 0 | 0 |
| body_centroid_to_roi_2_edge_dist_p10 | context (ROI/target) | 1 | 0 | 0 |
| body_centroid_to_roi_2_edge_dist_p90 | context (ROI/target) | 1 | 0 | 0 |
| body_centroid_to_roi_2_edge_dist_periodicity | context (ROI/target) | 1 | 0 | 0 |
| body_centroid_to_roi_2_edge_dist_std | context (ROI/target) | 1 | 0 | 0 |
| body_centroid_to_roi_2_edge_dist_trend | context (ROI/target) | 1 | 0 | 0 |
| body_centroid_to_roi_2_signed_dist_delta | context (ROI/target) | 1 | 0 | 0 |
| body_centroid_to_roi_2_signed_dist_energy | context (ROI/target) | 1 | 0 | 0 |
| body_centroid_to_roi_2_signed_dist_max | context (ROI/target) | 1 | 0 | 0 |
| body_centroid_to_roi_2_signed_dist_mean | context (ROI/target) | 1 | 0 | 0 |
| body_centroid_to_roi_2_signed_dist_median | context (ROI/target) | 1 | 0 | 0 |
| body_centroid_to_roi_2_signed_dist_p10 | context (ROI/target) | 1 | 0 | 0 |
| body_centroid_to_roi_2_signed_dist_p90 | context (ROI/target) | 1 | 0 | 0 |
| body_centroid_to_roi_2_signed_dist_periodicity | context (ROI/target) | 1 | 0 | 0 |
| body_centroid_to_roi_2_signed_dist_std | context (ROI/target) | 1 | 0 | 0 |
| body_centroid_to_roi_2_signed_dist_trend | context (ROI/target) | 1 | 0 | 0 |
| body_centroid_to_target_dist_delta | context (ROI/target) | 1 | 1 | 1 |
| body_centroid_to_target_dist_energy | context (ROI/target) | 1 | 1 | 1 |
| body_centroid_to_target_dist_max | context (ROI/target) | 1 | 1 | 1 |
| body_centroid_to_target_dist_mean | context (ROI/target) | 1 | 1 | 1 |
| body_centroid_to_target_dist_median | context (ROI/target) | 1 | 1 | 1 |
| body_centroid_to_target_dist_p10 | context (ROI/target) | 1 | 1 | 1 |
| body_centroid_to_target_dist_p90 | context (ROI/target) | 1 | 1 | 1 |
| body_centroid_to_target_dist_periodicity | context (ROI/target) | 1 | 1 | 1 |
| body_centroid_to_target_dist_std | context (ROI/target) | 1 | 1 | 1 |
| body_centroid_to_target_dist_trend | context (ROI/target) | 1 | 1 | 1 |
| body_left_acceleration_energy | kinematics | 0 | 1 | 0 |
| body_left_acceleration_max | kinematics | 0 | 1 | 0 |
| body_left_acceleration_mean | kinematics | 0 | 1 | 0 |
| body_left_acceleration_median | kinematics | 0 | 1 | 0 |
| body_left_acceleration_p10 | kinematics | 0 | 1 | 0 |
| body_left_acceleration_p90 | kinematics | 0 | 1 | 0 |
| body_left_acceleration_periodicity | kinematics | 0 | 1 | 0 |
| body_left_acceleration_std | kinematics | 0 | 1 | 0 |
| body_left_forward_velocity_energy | kinematics | 0 | 1 | 0 |
| body_left_forward_velocity_max | kinematics | 0 | 1 | 0 |
| body_left_forward_velocity_mean | kinematics | 0 | 1 | 0 |

| | | | | |
|---|---|---|---|---|
| body_left_forward_velocity_median | kinematics | 0 | 1 | 0 |
| body_left_forward_velocity_p10 | kinematics | 0 | 1 | 0 |
| body_left_forward_velocity_p90 | kinematics | 0 | 1 | 0 |
| body_left_forward_velocity_periodicity | kinematics | 0 | 1 | 0 |
| body_left_forward_velocity_std | kinematics | 0 | 1 | 0 |
| body_left_jerk_energy | kinematics | 0 | 1 | 0 |
| body_left_jerk_max | kinematics | 0 | 1 | 0 |
| body_left_jerk_mean | kinematics | 0 | 1 | 0 |
| body_left_jerk_median | kinematics | 0 | 1 | 0 |
| body_left_jerk_p10 | kinematics | 0 | 1 | 0 |
| body_left_jerk_p90 | kinematics | 0 | 1 | 0 |
| body_left_jerk_periodicity | kinematics | 0 | 1 | 0 |
| body_left_jerk_std | kinematics | 0 | 1 | 0 |
| body_left_lateral_velocity_energy | kinematics | 0 | 1 | 0 |
| body_left_lateral_velocity_max | kinematics | 0 | 1 | 0 |
| body_left_lateral_velocity_mean | kinematics | 0 | 1 | 0 |
| body_left_lateral_velocity_median | kinematics | 0 | 1 | 0 |
| body_left_lateral_velocity_p10 | kinematics | 0 | 1 | 0 |
| body_left_lateral_velocity_p90 | kinematics | 0 | 1 | 0 |
| body_left_lateral_velocity_periodicity | kinematics | 0 | 1 | 0 |
| body_left_lateral_velocity_std | kinematics | 0 | 1 | 0 |
| body_left_speed_energy | kinematics | 0 | 1 | 0 |
| body_left_speed_max | kinematics | 0 | 1 | 0 |
| body_left_speed_mean | kinematics | 0 | 1 | 0 |
| body_left_speed_median | kinematics | 0 | 1 | 0 |
| body_left_speed_p10 | kinematics | 0 | 1 | 0 |
| body_left_speed_p90 | kinematics | 0 | 1 | 0 |
| body_left_speed_periodicity | kinematics | 0 | 1 | 0 |
| body_left_speed_std | kinematics | 0 | 1 | 0 |
| body_left_velocity_x_energy | kinematics | 0 | 1 | 0 |
| body_left_velocity_x_max | kinematics | 0 | 1 | 0 |
| body_left_velocity_x_mean | kinematics | 0 | 1 | 0 |
| body_left_velocity_x_median | kinematics | 0 | 1 | 0 |
| body_left_velocity_x_p10 | kinematics | 0 | 1 | 0 |
| body_left_velocity_x_p90 | kinematics | 0 | 1 | 0 |
| body_left_velocity_x_periodicity | kinematics | 0 | 1 | 0 |
| body_left_velocity_x_std | kinematics | 0 | 1 | 0 |
| body_left_velocity_y_energy | kinematics | 0 | 1 | 0 |
| body_left_velocity_y_max | kinematics | 0 | 1 | 0 |
| body_left_velocity_y_mean | kinematics | 0 | 1 | 0 |
| body_left_velocity_y_median | kinematics | 0 | 1 | 0 |
| body_left_velocity_y_p10 | kinematics | 0 | 1 | 0 |
| body_left_velocity_y_p90 | kinematics | 0 | 1 | 0 |
| body_left_velocity_y_periodicity | kinematics | 0 | 1 | 0 |

| | | | | |
|---|---|---|---|---|
| body_left_velocity_y_std | kinematics | 0 | 1 | 0 |
| body_length_px_energy | kinematics | 1 | 1 | 1 |
| body_length_px_max | pose geometry | 1 | 1 | 1 |
| body_length_px_mean | pose geometry | 1 | 1 | 1 |
| body_length_px_median | pose geometry | 1 | 1 | 1 |
| body_length_px_p10 | pose geometry | 1 | 1 | 1 |
| body_length_px_p90 | pose geometry | 1 | 1 | 1 |
| body_length_px_periodicity | pose geometry | 1 | 1 | 1 |
| body_length_px_std | kinematics | 1 | 1 | 1 |
| body_mid_acceleration_energy | kinematics | 0 | 1 | 0 |
| body_mid_acceleration_max | kinematics | 0 | 1 | 0 |
| body_mid_acceleration_mean | kinematics | 0 | 1 | 0 |
| body_mid_acceleration_median | kinematics | 0 | 1 | 0 |
| body_mid_acceleration_p10 | kinematics | 0 | 1 | 0 |
| body_mid_acceleration_p90 | kinematics | 0 | 1 | 0 |
| body_mid_acceleration_periodicity | kinematics | 0 | 1 | 0 |
| body_mid_acceleration_std | kinematics | 0 | 1 | 0 |
| body_mid_forward_velocity_energy | kinematics | 0 | 1 | 0 |
| body_mid_forward_velocity_max | kinematics | 0 | 1 | 0 |
| body_mid_forward_velocity_mean | kinematics | 0 | 1 | 0 |
| body_mid_forward_velocity_median | kinematics | 0 | 1 | 0 |
| body_mid_forward_velocity_p10 | kinematics | 0 | 1 | 0 |
| body_mid_forward_velocity_p90 | kinematics | 0 | 1 | 0 |
| body_mid_forward_velocity_periodicity | kinematics | 0 | 1 | 0 |
| body_mid_forward_velocity_std | kinematics | 0 | 1 | 0 |
| body_mid_jerk_energy | kinematics | 0 | 1 | 0 |
| body_mid_jerk_max | kinematics | 0 | 1 | 0 |
| body_mid_jerk_mean | kinematics | 0 | 1 | 0 |
| body_mid_jerk_median | kinematics | 0 | 1 | 0 |
| body_mid_jerk_p10 | kinematics | 0 | 1 | 0 |
| body_mid_jerk_p90 | kinematics | 0 | 1 | 0 |
| body_mid_jerk_periodicity | kinematics | 0 | 1 | 0 |
| body_mid_jerk_std | kinematics | 0 | 1 | 0 |
| body_mid_lateral_velocity_energy | kinematics | 0 | 1 | 0 |
| body_mid_lateral_velocity_max | kinematics | 0 | 1 | 0 |
| body_mid_lateral_velocity_mean | kinematics | 0 | 1 | 0 |
| body_mid_lateral_velocity_median | kinematics | 0 | 1 | 0 |
| body_mid_lateral_velocity_p10 | kinematics | 0 | 1 | 0 |
| body_mid_lateral_velocity_p90 | kinematics | 0 | 1 | 0 |
| body_mid_lateral_velocity_periodicity | kinematics | 0 | 1 | 0 |
| body_mid_lateral_velocity_std | kinematics | 0 | 1 | 0 |
| body_mid_speed_energy | kinematics | 0 | 1 | 0 |
| body_mid_speed_max | kinematics | 0 | 1 | 0 |
| body_mid_speed_mean | kinematics | 0 | 1 | 0 |

| | | | | |
|---|---|---|---|---|
| body_mid_speed_median | kinematics | 0 | 1 | 0 |
| body_mid_speed_p10 | kinematics | 0 | 1 | 0 |
| body_mid_speed_p90 | kinematics | 0 | 1 | 0 |
| body_mid_speed_periodicity | kinematics | 0 | 1 | 0 |
| body_mid_speed_std | kinematics | 0 | 1 | 0 |
| body_mid_velocity_x_energy | kinematics | 0 | 1 | 0 |
| body_mid_velocity_x_max | kinematics | 0 | 1 | 0 |
| body_mid_velocity_x_mean | kinematics | 0 | 1 | 0 |
| body_mid_velocity_x_median | kinematics | 0 | 1 | 0 |
| body_mid_velocity_x_p10 | kinematics | 0 | 1 | 0 |
| body_mid_velocity_x_p90 | kinematics | 0 | 1 | 0 |
| body_mid_velocity_x_periodicity | kinematics | 0 | 1 | 0 |
| body_mid_velocity_x_std | kinematics | 0 | 1 | 0 |
| body_mid_velocity_y_energy | kinematics | 0 | 1 | 0 |
| body_mid_velocity_y_max | kinematics | 0 | 1 | 0 |
| body_mid_velocity_y_mean | kinematics | 0 | 1 | 0 |
| body_mid_velocity_y_median | kinematics | 0 | 1 | 0 |
| body_mid_velocity_y_p10 | kinematics | 0 | 1 | 0 |
| body_mid_velocity_y_p90 | kinematics | 0 | 1 | 0 |
| body_mid_velocity_y_periodicity | kinematics | 0 | 1 | 0 |
| body_mid_velocity_y_std | kinematics | 0 | 1 | 0 |
| body_orientation_delta | kinematics | 1 | 1 | 1 |
| body_orientation_energy | kinematics | 1 | 1 | 1 |
| body_orientation_max | pose geometry | 1 | 1 | 1 |
| body_orientation_mean | pose geometry | 1 | 1 | 1 |
| body_orientation_median | pose geometry | 1 | 1 | 1 |
| body_orientation_p10 | pose geometry | 1 | 1 | 1 |
| body_orientation_p90 | pose geometry | 1 | 1 | 1 |
| body_orientation_periodicity | pose geometry | 1 | 1 | 1 |
| body_orientation_std | kinematics | 1 | 1 | 1 |
| body_orientation_trend | pose geometry | 1 | 1 | 1 |
| body_right_acceleration_energy | kinematics | 0 | 1 | 0 |
| body_right_acceleration_max | kinematics | 0 | 1 | 0 |
| body_right_acceleration_mean | kinematics | 0 | 1 | 0 |
| body_right_acceleration_median | kinematics | 0 | 1 | 0 |
| body_right_acceleration_p10 | kinematics | 0 | 1 | 0 |
| body_right_acceleration_p90 | kinematics | 0 | 1 | 0 |
| body_right_acceleration_periodicity | kinematics | 0 | 1 | 0 |
| body_right_acceleration_std | kinematics | 0 | 1 | 0 |
| body_right_forward_velocity_energy | kinematics | 0 | 1 | 0 |
| body_right_forward_velocity_max | kinematics | 0 | 1 | 0 |
| body_right_forward_velocity_mean | kinematics | 0 | 1 | 0 |
| body_right_forward_velocity_median | kinematics | 0 | 1 | 0 |
| body_right_forward_velocity_p10 | kinematics | 0 | 1 | 0 |

| | | | | |
|---|---|---|---|---|
| body_right_forward_velocity_p90 | kinematics | 0 | 1 | 0 |
| body_right_forward_velocity_periodicity | kinematics | 0 | 1 | 0 |
| body_right_forward_velocity_std | kinematics | 0 | 1 | 0 |
| body_right_jerk_energy | kinematics | 0 | 1 | 0 |
| body_right_jerk_max | kinematics | 0 | 1 | 0 |
| body_right_jerk_mean | kinematics | 0 | 1 | 0 |
| body_right_jerk_median | kinematics | 0 | 1 | 0 |
| body_right_jerk_p10 | kinematics | 0 | 1 | 0 |
| body_right_jerk_p90 | kinematics | 0 | 1 | 0 |
| body_right_jerk_periodicity | kinematics | 0 | 1 | 0 |
| body_right_jerk_std | kinematics | 0 | 1 | 0 |
| body_right_lateral_velocity_energy | kinematics | 0 | 1 | 0 |
| body_right_lateral_velocity_max | kinematics | 0 | 1 | 0 |
| body_right_lateral_velocity_mean | kinematics | 0 | 1 | 0 |
| body_right_lateral_velocity_median | kinematics | 0 | 1 | 0 |
| body_right_lateral_velocity_p10 | kinematics | 0 | 1 | 0 |
| body_right_lateral_velocity_p90 | kinematics | 0 | 1 | 0 |
| body_right_lateral_velocity_periodicity | kinematics | 0 | 1 | 0 |
| body_right_lateral_velocity_std | kinematics | 0 | 1 | 0 |
| body_right_speed_energy | kinematics | 0 | 1 | 0 |
| body_right_speed_max | kinematics | 0 | 1 | 0 |
| body_right_speed_mean | kinematics | 0 | 1 | 0 |
| body_right_speed_median | kinematics | 0 | 1 | 0 |
| body_right_speed_p10 | kinematics | 0 | 1 | 0 |
| body_right_speed_p90 | kinematics | 0 | 1 | 0 |
| body_right_speed_periodicity | kinematics | 0 | 1 | 0 |
| body_right_speed_std | kinematics | 0 | 1 | 0 |
| body_right_velocity_x_energy | kinematics | 0 | 1 | 0 |
| body_right_velocity_x_max | kinematics | 0 | 1 | 0 |
| body_right_velocity_x_mean | kinematics | 0 | 1 | 0 |
| body_right_velocity_x_median | kinematics | 0 | 1 | 0 |
| body_right_velocity_x_p10 | kinematics | 0 | 1 | 0 |
| body_right_velocity_x_p90 | kinematics | 0 | 1 | 0 |
| body_right_velocity_x_periodicity | kinematics | 0 | 1 | 0 |
| body_right_velocity_x_std | kinematics | 0 | 1 | 0 |
| body_right_velocity_y_energy | kinematics | 0 | 1 | 0 |
| body_right_velocity_y_max | kinematics | 0 | 1 | 0 |
| body_right_velocity_y_mean | kinematics | 0 | 1 | 0 |
| body_right_velocity_y_median | kinematics | 0 | 1 | 0 |
| body_right_velocity_y_p10 | kinematics | 0 | 1 | 0 |
| body_right_velocity_y_p90 | kinematics | 0 | 1 | 0 |
| body_right_velocity_y_periodicity | kinematics | 0 | 1 | 0 |
| body_right_velocity_y_std | kinematics | 0 | 1 | 0 |
| center_acceleration_energy | context (ROI/target) | 0 | 0 | 1 |

| | | | | |
|---|---|---|---|---|
| center_acceleration_max | context (ROI/target) | 0 | 0 | 1 |
| center_acceleration_mean | context (ROI/target) | 0 | 0 | 1 |
| center_acceleration_median | context (ROI/target) | 0 | 0 | 1 |
| center_acceleration_p10 | context (ROI/target) | 0 | 0 | 1 |
| center_acceleration_p90 | context (ROI/target) | 0 | 0 | 1 |
| center_acceleration_periodicity | context (ROI/target) | 0 | 0 | 1 |
| center_acceleration_std | context (ROI/target) | 0 | 0 | 1 |
| center_body_acceleration_energy | context (ROI/target) | 1 | 0 | 0 |
| center_body_acceleration_max | context (ROI/target) | 1 | 0 | 0 |
| center_body_acceleration_mean | context (ROI/target) | 1 | 0 | 0 |
| center_body_acceleration_median | context (ROI/target) | 1 | 0 | 0 |
| center_body_acceleration_p10 | context (ROI/target) | 1 | 0 | 0 |
| center_body_acceleration_p90 | context (ROI/target) | 1 | 0 | 0 |
| center_body_acceleration_periodicity | context (ROI/target) | 1 | 0 | 0 |
| center_body_acceleration_std | context (ROI/target) | 1 | 0 | 0 |
| center_body_forward_velocity_energy | context (ROI/target) | 1 | 0 | 0 |
| center_body_forward_velocity_max | context (ROI/target) | 1 | 0 | 0 |
| center_body_forward_velocity_mean | context (ROI/target) | 1 | 0 | 0 |
| center_body_forward_velocity_median | context (ROI/target) | 1 | 0 | 0 |
| center_body_forward_velocity_p10 | context (ROI/target) | 1 | 0 | 0 |
| center_body_forward_velocity_p90 | context (ROI/target) | 1 | 0 | 0 |
| center_body_forward_velocity_periodicity | context (ROI/target) | 1 | 0 | 0 |
| center_body_forward_velocity_std | context (ROI/target) | 1 | 0 | 0 |
| center_body_jerk_energy | context (ROI/target) | 1 | 0 | 0 |
| center_body_jerk_max | context (ROI/target) | 1 | 0 | 0 |
| center_body_jerk_mean | context (ROI/target) | 1 | 0 | 0 |
| center_body_jerk_median | context (ROI/target) | 1 | 0 | 0 |
| center_body_jerk_p10 | context (ROI/target) | 1 | 0 | 0 |
| center_body_jerk_p90 | context (ROI/target) | 1 | 0 | 0 |
| center_body_jerk_periodicity | context (ROI/target) | 1 | 0 | 0 |
| center_body_jerk_std | context (ROI/target) | 1 | 0 | 0 |
| center_body_lateral_velocity_energy | context (ROI/target) | 1 | 0 | 0 |
| center_body_lateral_velocity_max | context (ROI/target) | 1 | 0 | 0 |
| center_body_lateral_velocity_mean | context (ROI/target) | 1 | 0 | 0 |
| center_body_lateral_velocity_median | context (ROI/target) | 1 | 0 | 0 |
| center_body_lateral_velocity_p10 | context (ROI/target) | 1 | 0 | 0 |
| center_body_lateral_velocity_p90 | context (ROI/target) | 1 | 0 | 0 |
| center_body_lateral_velocity_periodicity | context (ROI/target) | 1 | 0 | 0 |
| center_body_lateral_velocity_std | context (ROI/target) | 1 | 0 | 0 |
| center_body_speed_energy | context (ROI/target) | 1 | 0 | 0 |
| center_body_speed_max | context (ROI/target) | 1 | 0 | 0 |
| center_body_speed_mean | context (ROI/target) | 1 | 0 | 0 |
| center_body_speed_median | context (ROI/target) | 1 | 0 | 0 |
| center_body_speed_p10 | context (ROI/target) | 1 | 0 | 0 |

| | | | | |
|---|---|---|---|---|
| center_body_speed_p90 | context (ROI/target) | 1 | 0 | 0 |
| center_body_speed_periodicity | context (ROI/target) | 1 | 0 | 0 |
| center_body_speed_std | context (ROI/target) | 1 | 0 | 0 |
| center_body_velocity_x_energy | context (ROI/target) | 1 | 0 | 0 |
| center_body_velocity_x_max | context (ROI/target) | 1 | 0 | 0 |
| center_body_velocity_x_mean | context (ROI/target) | 1 | 0 | 0 |
| center_body_velocity_x_median | context (ROI/target) | 1 | 0 | 0 |
| center_body_velocity_x_p10 | context (ROI/target) | 1 | 0 | 0 |
| center_body_velocity_x_p90 | context (ROI/target) | 1 | 0 | 0 |
| center_body_velocity_x_periodicity | context (ROI/target) | 1 | 0 | 0 |
| center_body_velocity_x_std | context (ROI/target) | 1 | 0 | 0 |
| center_body_velocity_y_energy | context (ROI/target) | 1 | 0 | 0 |
| center_body_velocity_y_max | context (ROI/target) | 1 | 0 | 0 |
| center_body_velocity_y_mean | context (ROI/target) | 1 | 0 | 0 |
| center_body_velocity_y_median | context (ROI/target) | 1 | 0 | 0 |
| center_body_velocity_y_p10 | context (ROI/target) | 1 | 0 | 0 |
| center_body_velocity_y_p90 | context (ROI/target) | 1 | 0 | 0 |
| center_body_velocity_y_periodicity | context (ROI/target) | 1 | 0 | 0 |
| center_body_velocity_y_std | context (ROI/target) | 1 | 0 | 0 |
| center_forward_velocity_energy | context (ROI/target) | 0 | 0 | 1 |
| center_forward_velocity_max | context (ROI/target) | 0 | 0 | 1 |
| center_forward_velocity_mean | context (ROI/target) | 0 | 0 | 1 |
| center_forward_velocity_median | context (ROI/target) | 0 | 0 | 1 |
| center_forward_velocity_p10 | context (ROI/target) | 0 | 0 | 1 |
| center_forward_velocity_p90 | context (ROI/target) | 0 | 0 | 1 |
| center_forward_velocity_periodicity | context (ROI/target) | 0 | 0 | 1 |
| center_forward_velocity_std | context (ROI/target) | 0 | 0 | 1 |
| center_jerk_energy | context (ROI/target) | 0 | 0 | 1 |
| center_jerk_max | context (ROI/target) | 0 | 0 | 1 |
| center_jerk_mean | context (ROI/target) | 0 | 0 | 1 |
| center_jerk_median | context (ROI/target) | 0 | 0 | 1 |
| center_jerk_p10 | context (ROI/target) | 0 | 0 | 1 |
| center_jerk_p90 | context (ROI/target) | 0 | 0 | 1 |
| center_jerk_periodicity | context (ROI/target) | 0 | 0 | 1 |
| center_jerk_std | context (ROI/target) | 0 | 0 | 1 |
| center_lateral_velocity_energy | context (ROI/target) | 0 | 0 | 1 |
| center_lateral_velocity_max | context (ROI/target) | 0 | 0 | 1 |
| center_lateral_velocity_mean | context (ROI/target) | 0 | 0 | 1 |
| center_lateral_velocity_median | context (ROI/target) | 0 | 0 | 1 |
| center_lateral_velocity_p10 | context (ROI/target) | 0 | 0 | 1 |
| center_lateral_velocity_p90 | context (ROI/target) | 0 | 0 | 1 |
| center_lateral_velocity_periodicity | context (ROI/target) | 0 | 0 | 1 |
| center_lateral_velocity_std | context (ROI/target) | 0 | 0 | 1 |
| center_speed_energy | context (ROI/target) | 0 | 0 | 1 |

| | | | | |
|---|---|---|---|---|
| center_speed_max | context (ROI/target) | 0 | 0 | 1 |
| center_speed_mean | context (ROI/target) | 0 | 0 | 1 |
| center_speed_median | context (ROI/target) | 0 | 0 | 1 |
| center_speed_p10 | context (ROI/target) | 0 | 0 | 1 |
| center_speed_p90 | context (ROI/target) | 0 | 0 | 1 |
| center_speed_periodicity | context (ROI/target) | 0 | 0 | 1 |
| center_speed_std | context (ROI/target) | 0 | 0 | 1 |
| center_velocity_x_energy | context (ROI/target) | 0 | 0 | 1 |
| center_velocity_x_max | context (ROI/target) | 0 | 0 | 1 |
| center_velocity_x_mean | context (ROI/target) | 0 | 0 | 1 |
| center_velocity_x_median | context (ROI/target) | 0 | 0 | 1 |
| center_velocity_x_p10 | context (ROI/target) | 0 | 0 | 1 |
| center_velocity_x_p90 | context (ROI/target) | 0 | 0 | 1 |
| center_velocity_x_periodicity | context (ROI/target) | 0 | 0 | 1 |
| center_velocity_x_std | context (ROI/target) | 0 | 0 | 1 |
| center_velocity_y_energy | context (ROI/target) | 0 | 0 | 1 |
| center_velocity_y_max | context (ROI/target) | 0 | 0 | 1 |
| center_velocity_y_mean | context (ROI/target) | 0 | 0 | 1 |
| center_velocity_y_median | context (ROI/target) | 0 | 0 | 1 |
| center_velocity_y_p10 | context (ROI/target) | 0 | 0 | 1 |
| center_velocity_y_p90 | context (ROI/target) | 0 | 0 | 1 |
| center_velocity_y_periodicity | context (ROI/target) | 0 | 0 | 1 |
| center_velocity_y_std | context (ROI/target) | 0 | 0 | 1 |
| centroid_velocity_energy | context (ROI/target) | 1 | 1 | 1 |
| centroid_velocity_max | context (ROI/target) | 1 | 1 | 1 |
| centroid_velocity_mean | context (ROI/target) | 1 | 1 | 1 |
| centroid_velocity_median | context (ROI/target) | 1 | 1 | 1 |
| centroid_velocity_p10 | context (ROI/target) | 1 | 1 | 1 |
| centroid_velocity_p90 | context (ROI/target) | 1 | 1 | 1 |
| centroid_velocity_periodicity | context (ROI/target) | 1 | 1 | 1 |
| centroid_velocity_std | context (ROI/target) | 1 | 1 | 1 |
| density_outlier_score | pose geometry | 1 | 1 | 1 |
| dist_back_left_paw_to_back_right_paw_delta | kinematics | 0 | 0 | 0 |
| dist_back_left_paw_to_back_right_paw_energy | kinematics | 0 | 0 | 0 |
| dist_back_left_paw_to_back_right_paw_max | pose geometry | 0 | 0 | 0 |
| dist_back_left_paw_to_back_right_paw_mean | pose geometry | 0 | 0 | 0 |
| dist_back_left_paw_to_back_right_paw_median | pose geometry | 0 | 0 | 0 |
| dist_back_left_paw_to_back_right_paw_norm_delta | kinematics | 0 | 0 | 0 |
| dist_back_left_paw_to_back_right_paw_norm_energy | kinematics | 0 | 0 | 0 |
| dist_back_left_paw_to_back_right_paw_norm_max | pose geometry | 0 | 0 | 0 |
| dist_back_left_paw_to_back_right_paw_norm_mean | pose geometry | 0 | 0 | 0 |
| dist_back_left_paw_to_back_right_paw_norm_median | pose geometry | 0 | 0 | 0 |
| dist_back_left_paw_to_back_right_paw_norm_p10 | pose geometry | 0 | 0 | 0 |
| dist_back_left_paw_to_back_right_paw_norm_p90 | pose geometry | 0 | 0 | 0 |

| | | | | |
|---|---|---|---|---|
| dist_back_left_paw_to_back_right_paw_norm_periodic | pose geometry | 0 | 0 | 0 |
| dist_back_left_paw_to_back_right_paw_norm_std | kinematics | 0 | 0 | 0 |
| dist_back_left_paw_to_back_right_paw_norm_trend | pose geometry | 0 | 0 | 0 |
| dist_back_left_paw_to_back_right_paw_p10 | pose geometry | 0 | 0 | 0 |
| dist_back_left_paw_to_back_right_paw_p90 | pose geometry | 0 | 0 | 0 |
| dist_back_left_paw_to_back_right_paw_periodicity | pose geometry | 0 | 0 | 0 |
| dist_back_left_paw_to_back_right_paw_std | kinematics | 0 | 0 | 0 |
| dist_back_left_paw_to_back_right_paw_trend | pose geometry | 0 | 0 | 0 |
| dist_back_left_paw_to_front_left_paw_delta | kinematics | 0 | 0 | 0 |
| dist_back_left_paw_to_front_left_paw_energy | kinematics | 0 | 0 | 0 |
| dist_back_left_paw_to_front_left_paw_max | pose geometry | 0 | 0 | 0 |
| dist_back_left_paw_to_front_left_paw_mean | pose geometry | 0 | 0 | 0 |
| dist_back_left_paw_to_front_left_paw_median | pose geometry | 0 | 0 | 0 |
| dist_back_left_paw_to_front_left_paw_norm_delta | kinematics | 0 | 0 | 0 |
| dist_back_left_paw_to_front_left_paw_norm_energy | kinematics | 0 | 0 | 0 |
| dist_back_left_paw_to_front_left_paw_norm_max | pose geometry | 0 | 0 | 0 |
| dist_back_left_paw_to_front_left_paw_norm_mean | pose geometry | 0 | 0 | 0 |
| dist_back_left_paw_to_front_left_paw_norm_median | pose geometry | 0 | 0 | 0 |
| dist_back_left_paw_to_front_left_paw_norm_p10 | pose geometry | 0 | 0 | 0 |
| dist_back_left_paw_to_front_left_paw_norm_p90 | pose geometry | 0 | 0 | 0 |
| dist_back_left_paw_to_front_left_paw_norm_periodicit | pose geometry | 0 | 0 | 0 |
| dist_back_left_paw_to_front_left_paw_norm_std | kinematics | 0 | 0 | 0 |
| dist_back_left_paw_to_front_left_paw_norm_trend | pose geometry | 0 | 0 | 0 |
| dist_back_left_paw_to_front_left_paw_p10 | pose geometry | 0 | 0 | 0 |
| dist_back_left_paw_to_front_left_paw_p90 | pose geometry | 0 | 0 | 0 |
| dist_back_left_paw_to_front_left_paw_periodicity | pose geometry | 0 | 0 | 0 |
| dist_back_left_paw_to_front_left_paw_std | kinematics | 0 | 0 | 0 |
| dist_back_left_paw_to_front_left_paw_trend | pose geometry | 0 | 0 | 0 |
| dist_back_left_paw_to_front_right_paw_delta | kinematics | 0 | 0 | 0 |
| dist_back_left_paw_to_front_right_paw_energy | kinematics | 0 | 0 | 0 |
| dist_back_left_paw_to_front_right_paw_max | pose geometry | 0 | 0 | 0 |
| dist_back_left_paw_to_front_right_paw_mean | pose geometry | 0 | 0 | 0 |
| dist_back_left_paw_to_front_right_paw_median | pose geometry | 0 | 0 | 0 |
| dist_back_left_paw_to_front_right_paw_norm_delta | kinematics | 0 | 0 | 0 |
| dist_back_left_paw_to_front_right_paw_norm_energy | kinematics | 0 | 0 | 0 |
| dist_back_left_paw_to_front_right_paw_norm_max | pose geometry | 0 | 0 | 0 |
| dist_back_left_paw_to_front_right_paw_norm_mean | pose geometry | 0 | 0 | 0 |
| dist_back_left_paw_to_front_right_paw_norm_median | pose geometry | 0 | 0 | 0 |
| dist_back_left_paw_to_front_right_paw_norm_p10 | pose geometry | 0 | 0 | 0 |
| dist_back_left_paw_to_front_right_paw_norm_p90 | pose geometry | 0 | 0 | 0 |
| dist_back_left_paw_to_front_right_paw_norm_periodic | pose geometry | 0 | 0 | 0 |
| dist_back_left_paw_to_front_right_paw_norm_std | kinematics | 0 | 0 | 0 |
| dist_back_left_paw_to_front_right_paw_norm_trend | pose geometry | 0 | 0 | 0 |
| dist_back_left_paw_to_front_right_paw_p10 | pose geometry | 0 | 0 | 0 |

| | | | | |
|---|---|---|---|---|
| dist_back_left_paw_to_front_right_paw_p90 | pose geometry | 0 | 0 | 0 |
| dist_back_left_paw_to_front_right_paw_periodicity | pose geometry | 0 | 0 | 0 |
| dist_back_left_paw_to_front_right_paw_std | kinematics | 0 | 0 | 0 |
| dist_back_left_paw_to_front_right_paw_trend | pose geometry | 0 | 0 | 0 |
| dist_back_left_paw_to_left_ear_delta | kinematics | 0 | 0 | 0 |
| dist_back_left_paw_to_left_ear_energy | kinematics | 0 | 0 | 0 |
| dist_back_left_paw_to_left_ear_max | pose geometry | 0 | 0 | 0 |
| dist_back_left_paw_to_left_ear_mean | pose geometry | 0 | 0 | 0 |
| dist_back_left_paw_to_left_ear_median | pose geometry | 0 | 0 | 0 |
| dist_back_left_paw_to_left_ear_norm_delta | kinematics | 0 | 0 | 0 |
| dist_back_left_paw_to_left_ear_norm_energy | kinematics | 0 | 0 | 0 |
| dist_back_left_paw_to_left_ear_norm_max | pose geometry | 0 | 0 | 0 |
| dist_back_left_paw_to_left_ear_norm_mean | pose geometry | 0 | 0 | 0 |
| dist_back_left_paw_to_left_ear_norm_median | pose geometry | 0 | 0 | 0 |
| dist_back_left_paw_to_left_ear_norm_p10 | pose geometry | 0 | 0 | 0 |
| dist_back_left_paw_to_left_ear_norm_p90 | pose geometry | 0 | 0 | 0 |
| dist_back_left_paw_to_left_ear_norm_periodicity | pose geometry | 0 | 0 | 0 |
| dist_back_left_paw_to_left_ear_norm_std | kinematics | 0 | 0 | 0 |
| dist_back_left_paw_to_left_ear_norm_trend | pose geometry | 0 | 0 | 0 |
| dist_back_left_paw_to_left_ear_p10 | pose geometry | 0 | 0 | 0 |
| dist_back_left_paw_to_left_ear_p90 | pose geometry | 0 | 0 | 0 |
| dist_back_left_paw_to_left_ear_periodicity | pose geometry | 0 | 0 | 0 |
| dist_back_left_paw_to_left_ear_std | kinematics | 0 | 0 | 0 |
| dist_back_left_paw_to_left_ear_trend | pose geometry | 0 | 0 | 0 |
| dist_back_left_paw_to_mid_back_delta | kinematics | 0 | 0 | 0 |
| dist_back_left_paw_to_mid_back_energy | kinematics | 0 | 0 | 0 |
| dist_back_left_paw_to_mid_back_max | pose geometry | 0 | 0 | 0 |
| dist_back_left_paw_to_mid_back_mean | pose geometry | 0 | 0 | 0 |
| dist_back_left_paw_to_mid_back_median | pose geometry | 0 | 0 | 0 |
| dist_back_left_paw_to_mid_back_norm_delta | kinematics | 0 | 0 | 0 |
| dist_back_left_paw_to_mid_back_norm_energy | kinematics | 0 | 0 | 0 |
| dist_back_left_paw_to_mid_back_norm_max | pose geometry | 0 | 0 | 0 |
| dist_back_left_paw_to_mid_back_norm_mean | pose geometry | 0 | 0 | 0 |
| dist_back_left_paw_to_mid_back_norm_median | pose geometry | 0 | 0 | 0 |
| dist_back_left_paw_to_mid_back_norm_p10 | pose geometry | 0 | 0 | 0 |
| dist_back_left_paw_to_mid_back_norm_p90 | pose geometry | 0 | 0 | 0 |
| dist_back_left_paw_to_mid_back_norm_periodicity | pose geometry | 0 | 0 | 0 |
| dist_back_left_paw_to_mid_back_norm_std | kinematics | 0 | 0 | 0 |
| dist_back_left_paw_to_mid_back_norm_trend | pose geometry | 0 | 0 | 0 |
| dist_back_left_paw_to_mid_back_p10 | pose geometry | 0 | 0 | 0 |
| dist_back_left_paw_to_mid_back_p90 | pose geometry | 0 | 0 | 0 |
| dist_back_left_paw_to_mid_back_periodicity | pose geometry | 0 | 0 | 0 |
| dist_back_left_paw_to_mid_back_std | kinematics | 0 | 0 | 0 |
| dist_back_left_paw_to_mid_back_trend | pose geometry | 0 | 0 | 0 |

| | | | | |
|---|---|---|---|---|
| dist_back_left_paw_to_nose_delta | kinematics | 0 | 0 | 0 |
| dist_back_left_paw_to_nose_energy | kinematics | 0 | 0 | 0 |
| dist_back_left_paw_to_nose_max | pose geometry | 0 | 0 | 0 |
| dist_back_left_paw_to_nose_mean | pose geometry | 0 | 0 | 0 |
| dist_back_left_paw_to_nose_median | pose geometry | 0 | 0 | 0 |
| dist_back_left_paw_to_nose_norm_delta | kinematics | 0 | 0 | 0 |
| dist_back_left_paw_to_nose_norm_energy | kinematics | 0 | 0 | 0 |
| dist_back_left_paw_to_nose_norm_max | pose geometry | 0 | 0 | 0 |
| dist_back_left_paw_to_nose_norm_mean | pose geometry | 0 | 0 | 0 |
| dist_back_left_paw_to_nose_norm_median | pose geometry | 0 | 0 | 0 |
| dist_back_left_paw_to_nose_norm_p10 | pose geometry | 0 | 0 | 0 |
| dist_back_left_paw_to_nose_norm_p90 | pose geometry | 0 | 0 | 0 |
| dist_back_left_paw_to_nose_norm_periodicity | pose geometry | 0 | 0 | 0 |
| dist_back_left_paw_to_nose_norm_std | kinematics | 0 | 0 | 0 |
| dist_back_left_paw_to_nose_norm_trend | pose geometry | 0 | 0 | 0 |
| dist_back_left_paw_to_nose_p10 | pose geometry | 0 | 0 | 0 |
| dist_back_left_paw_to_nose_p90 | pose geometry | 0 | 0 | 0 |
| dist_back_left_paw_to_nose_periodicity | pose geometry | 0 | 0 | 0 |
| dist_back_left_paw_to_nose_std | kinematics | 0 | 0 | 0 |
| dist_back_left_paw_to_nose_trend | pose geometry | 0 | 0 | 0 |
| dist_back_left_paw_to_right_ear_delta | kinematics | 0 | 0 | 0 |
| dist_back_left_paw_to_right_ear_energy | kinematics | 0 | 0 | 0 |
| dist_back_left_paw_to_right_ear_max | pose geometry | 0 | 0 | 0 |
| dist_back_left_paw_to_right_ear_mean | pose geometry | 0 | 0 | 0 |
| dist_back_left_paw_to_right_ear_median | pose geometry | 0 | 0 | 0 |
| dist_back_left_paw_to_right_ear_norm_delta | kinematics | 0 | 0 | 0 |
| dist_back_left_paw_to_right_ear_norm_energy | kinematics | 0 | 0 | 0 |
| dist_back_left_paw_to_right_ear_norm_max | pose geometry | 0 | 0 | 0 |
| dist_back_left_paw_to_right_ear_norm_mean | pose geometry | 0 | 0 | 0 |
| dist_back_left_paw_to_right_ear_norm_median | pose geometry | 0 | 0 | 0 |
| dist_back_left_paw_to_right_ear_norm_p10 | pose geometry | 0 | 0 | 0 |
| dist_back_left_paw_to_right_ear_norm_p90 | pose geometry | 0 | 0 | 0 |
| dist_back_left_paw_to_right_ear_norm_periodicity | pose geometry | 0 | 0 | 0 |
| dist_back_left_paw_to_right_ear_norm_std | kinematics | 0 | 0 | 0 |
| dist_back_left_paw_to_right_ear_norm_trend | pose geometry | 0 | 0 | 0 |
| dist_back_left_paw_to_right_ear_p10 | pose geometry | 0 | 0 | 0 |
| dist_back_left_paw_to_right_ear_p90 | pose geometry | 0 | 0 | 0 |
| dist_back_left_paw_to_right_ear_periodicity | pose geometry | 0 | 0 | 0 |
| dist_back_left_paw_to_right_ear_std | kinematics | 0 | 0 | 0 |
| dist_back_left_paw_to_right_ear_trend | pose geometry | 0 | 0 | 0 |
| dist_back_left_paw_to_tail_base_delta | kinematics | 0 | 0 | 0 |
| dist_back_left_paw_to_tail_base_energy | kinematics | 0 | 0 | 0 |
| dist_back_left_paw_to_tail_base_max | pose geometry | 0 | 0 | 0 |
| dist_back_left_paw_to_tail_base_mean | pose geometry | 0 | 0 | 0 |

| | | | | |
|---|---|---|---|---|
| dist_back_left_paw_to_tail_base_median | pose geometry | 0 | 0 | 0 |
| dist_back_left_paw_to_tail_base_norm_delta | kinematics | 0 | 0 | 0 |
| dist_back_left_paw_to_tail_base_norm_energy | kinematics | 0 | 0 | 0 |
| dist_back_left_paw_to_tail_base_norm_max | pose geometry | 0 | 0 | 0 |
| dist_back_left_paw_to_tail_base_norm_mean | pose geometry | 0 | 0 | 0 |
| dist_back_left_paw_to_tail_base_norm_median | pose geometry | 0 | 0 | 0 |
| dist_back_left_paw_to_tail_base_norm_p10 | pose geometry | 0 | 0 | 0 |
| dist_back_left_paw_to_tail_base_norm_p90 | pose geometry | 0 | 0 | 0 |
| dist_back_left_paw_to_tail_base_norm_periodicity | pose geometry | 0 | 0 | 0 |
| dist_back_left_paw_to_tail_base_norm_std | kinematics | 0 | 0 | 0 |
| dist_back_left_paw_to_tail_base_norm_trend | pose geometry | 0 | 0 | 0 |
| dist_back_left_paw_to_tail_base_p10 | pose geometry | 0 | 0 | 0 |
| dist_back_left_paw_to_tail_base_p90 | pose geometry | 0 | 0 | 0 |
| dist_back_left_paw_to_tail_base_periodicity | pose geometry | 0 | 0 | 0 |
| dist_back_left_paw_to_tail_base_std | kinematics | 0 | 0 | 0 |
| dist_back_left_paw_to_tail_base_trend | pose geometry | 0 | 0 | 0 |
| dist_back_right_paw_to_front_left_paw_delta | kinematics | 0 | 0 | 0 |
| dist_back_right_paw_to_front_left_paw_energy | kinematics | 0 | 0 | 0 |
| dist_back_right_paw_to_front_left_paw_max | pose geometry | 0 | 0 | 0 |
| dist_back_right_paw_to_front_left_paw_mean | pose geometry | 0 | 0 | 0 |
| dist_back_right_paw_to_front_left_paw_median | pose geometry | 0 | 0 | 0 |
| dist_back_right_paw_to_front_left_paw_norm_delta | kinematics | 0 | 0 | 0 |
| dist_back_right_paw_to_front_left_paw_norm_energy | kinematics | 0 | 0 | 0 |
| dist_back_right_paw_to_front_left_paw_norm_max | pose geometry | 0 | 0 | 0 |
| dist_back_right_paw_to_front_left_paw_norm_mean | pose geometry | 0 | 0 | 0 |
| dist_back_right_paw_to_front_left_paw_norm_median | pose geometry | 0 | 0 | 0 |
| dist_back_right_paw_to_front_left_paw_norm_p10 | pose geometry | 0 | 0 | 0 |
| dist_back_right_paw_to_front_left_paw_norm_p90 | pose geometry | 0 | 0 | 0 |
| dist_back_right_paw_to_front_left_paw_norm_periodic | pose geometry | 0 | 0 | 0 |
| dist_back_right_paw_to_front_left_paw_norm_std | kinematics | 0 | 0 | 0 |
| dist_back_right_paw_to_front_left_paw_norm_trend | pose geometry | 0 | 0 | 0 |
| dist_back_right_paw_to_front_left_paw_p10 | pose geometry | 0 | 0 | 0 |
| dist_back_right_paw_to_front_left_paw_p90 | pose geometry | 0 | 0 | 0 |
| dist_back_right_paw_to_front_left_paw_periodicity | pose geometry | 0 | 0 | 0 |
| dist_back_right_paw_to_front_left_paw_std | kinematics | 0 | 0 | 0 |
| dist_back_right_paw_to_front_left_paw_trend | pose geometry | 0 | 0 | 0 |
| dist_back_right_paw_to_front_right_paw_delta | kinematics | 0 | 0 | 0 |
| dist_back_right_paw_to_front_right_paw_energy | kinematics | 0 | 0 | 0 |
| dist_back_right_paw_to_front_right_paw_max | pose geometry | 0 | 0 | 0 |
| dist_back_right_paw_to_front_right_paw_mean | pose geometry | 0 | 0 | 0 |
| dist_back_right_paw_to_front_right_paw_median | pose geometry | 0 | 0 | 0 |
| dist_back_right_paw_to_front_right_paw_norm_delta | kinematics | 0 | 0 | 0 |
| dist_back_right_paw_to_front_right_paw_norm_energy | kinematics | 0 | 0 | 0 |
| dist_back_right_paw_to_front_right_paw_norm_max | pose geometry | 0 | 0 | 0 |

| | | | | |
|---|---|---|---|---|
| dist_back_right_paw_to_front_right_paw_norm_mean | pose geometry | 0 | 0 | 0 |
| dist_back_right_paw_to_front_right_paw_norm_media | pose geometry | 0 | 0 | 0 |
| dist_back_right_paw_to_front_right_paw_norm_p10 | pose geometry | 0 | 0 | 0 |
| dist_back_right_paw_to_front_right_paw_norm_p90 | pose geometry | 0 | 0 | 0 |
| dist_back_right_paw_to_front_right_paw_norm_period | pose geometry | 0 | 0 | 0 |
| dist_back_right_paw_to_front_right_paw_norm_std | kinematics | 0 | 0 | 0 |
| dist_back_right_paw_to_front_right_paw_norm_trend | pose geometry | 0 | 0 | 0 |
| dist_back_right_paw_to_front_right_paw_p10 | pose geometry | 0 | 0 | 0 |
| dist_back_right_paw_to_front_right_paw_p90 | pose geometry | 0 | 0 | 0 |
| dist_back_right_paw_to_front_right_paw_periodicity | pose geometry | 0 | 0 | 0 |
| dist_back_right_paw_to_front_right_paw_std | kinematics | 0 | 0 | 0 |
| dist_back_right_paw_to_front_right_paw_trend | pose geometry | 0 | 0 | 0 |
| dist_back_right_paw_to_left_ear_delta | kinematics | 0 | 0 | 0 |
| dist_back_right_paw_to_left_ear_energy | kinematics | 0 | 0 | 0 |
| dist_back_right_paw_to_left_ear_max | pose geometry | 0 | 0 | 0 |
| dist_back_right_paw_to_left_ear_mean | pose geometry | 0 | 0 | 0 |
| dist_back_right_paw_to_left_ear_median | pose geometry | 0 | 0 | 0 |
| dist_back_right_paw_to_left_ear_norm_delta | kinematics | 0 | 0 | 0 |
| dist_back_right_paw_to_left_ear_norm_energy | kinematics | 0 | 0 | 0 |
| dist_back_right_paw_to_left_ear_norm_max | pose geometry | 0 | 0 | 0 |
| dist_back_right_paw_to_left_ear_norm_mean | pose geometry | 0 | 0 | 0 |
| dist_back_right_paw_to_left_ear_norm_median | pose geometry | 0 | 0 | 0 |
| dist_back_right_paw_to_left_ear_norm_p10 | pose geometry | 0 | 0 | 0 |
| dist_back_right_paw_to_left_ear_norm_p90 | pose geometry | 0 | 0 | 0 |
| dist_back_right_paw_to_left_ear_norm_periodicity | pose geometry | 0 | 0 | 0 |
| dist_back_right_paw_to_left_ear_norm_std | kinematics | 0 | 0 | 0 |
| dist_back_right_paw_to_left_ear_norm_trend | pose geometry | 0 | 0 | 0 |
| dist_back_right_paw_to_left_ear_p10 | pose geometry | 0 | 0 | 0 |
| dist_back_right_paw_to_left_ear_p90 | pose geometry | 0 | 0 | 0 |
| dist_back_right_paw_to_left_ear_periodicity | pose geometry | 0 | 0 | 0 |
| dist_back_right_paw_to_left_ear_std | kinematics | 0 | 0 | 0 |
| dist_back_right_paw_to_left_ear_trend | pose geometry | 0 | 0 | 0 |
| dist_back_right_paw_to_mid_back_delta | kinematics | 0 | 0 | 0 |
| dist_back_right_paw_to_mid_back_energy | kinematics | 0 | 0 | 0 |
| dist_back_right_paw_to_mid_back_max | pose geometry | 0 | 0 | 0 |
| dist_back_right_paw_to_mid_back_mean | pose geometry | 0 | 0 | 0 |
| dist_back_right_paw_to_mid_back_median | pose geometry | 0 | 0 | 0 |
| dist_back_right_paw_to_mid_back_norm_delta | kinematics | 0 | 0 | 0 |
| dist_back_right_paw_to_mid_back_norm_energy | kinematics | 0 | 0 | 0 |
| dist_back_right_paw_to_mid_back_norm_max | pose geometry | 0 | 0 | 0 |
| dist_back_right_paw_to_mid_back_norm_mean | pose geometry | 0 | 0 | 0 |
| dist_back_right_paw_to_mid_back_norm_median | pose geometry | 0 | 0 | 0 |
| dist_back_right_paw_to_mid_back_norm_p10 | pose geometry | 0 | 0 | 0 |
| dist_back_right_paw_to_mid_back_norm_p90 | pose geometry | 0 | 0 | 0 |

| | | | | |
|---|---|---|---|---|
| dist_back_right_paw_to_mid_back_norm_periodicity | pose geometry | 0 | 0 | 0 |
| dist_back_right_paw_to_mid_back_norm_std | kinematics | 0 | 0 | 0 |
| dist_back_right_paw_to_mid_back_norm_trend | pose geometry | 0 | 0 | 0 |
| dist_back_right_paw_to_mid_back_p10 | pose geometry | 0 | 0 | 0 |
| dist_back_right_paw_to_mid_back_p90 | pose geometry | 0 | 0 | 0 |
| dist_back_right_paw_to_mid_back_periodicity | pose geometry | 0 | 0 | 0 |
| dist_back_right_paw_to_mid_back_std | kinematics | 0 | 0 | 0 |
| dist_back_right_paw_to_mid_back_trend | pose geometry | 0 | 0 | 0 |
| dist_back_right_paw_to_nose_delta | kinematics | 0 | 0 | 0 |
| dist_back_right_paw_to_nose_energy | kinematics | 0 | 0 | 0 |
| dist_back_right_paw_to_nose_max | pose geometry | 0 | 0 | 0 |
| dist_back_right_paw_to_nose_mean | pose geometry | 0 | 0 | 0 |
| dist_back_right_paw_to_nose_median | pose geometry | 0 | 0 | 0 |
| dist_back_right_paw_to_nose_norm_delta | kinematics | 0 | 0 | 0 |
| dist_back_right_paw_to_nose_norm_energy | kinematics | 0 | 0 | 0 |
| dist_back_right_paw_to_nose_norm_max | pose geometry | 0 | 0 | 0 |
| dist_back_right_paw_to_nose_norm_mean | pose geometry | 0 | 0 | 0 |
| dist_back_right_paw_to_nose_norm_median | pose geometry | 0 | 0 | 0 |
| dist_back_right_paw_to_nose_norm_p10 | pose geometry | 0 | 0 | 0 |
| dist_back_right_paw_to_nose_norm_p90 | pose geometry | 0 | 0 | 0 |
| dist_back_right_paw_to_nose_norm_periodicity | pose geometry | 0 | 0 | 0 |
| dist_back_right_paw_to_nose_norm_std | kinematics | 0 | 0 | 0 |
| dist_back_right_paw_to_nose_norm_trend | pose geometry | 0 | 0 | 0 |
| dist_back_right_paw_to_nose_p10 | pose geometry | 0 | 0 | 0 |
| dist_back_right_paw_to_nose_p90 | pose geometry | 0 | 0 | 0 |
| dist_back_right_paw_to_nose_periodicity | pose geometry | 0 | 0 | 0 |
| dist_back_right_paw_to_nose_std | kinematics | 0 | 0 | 0 |
| dist_back_right_paw_to_nose_trend | pose geometry | 0 | 0 | 0 |
| dist_back_right_paw_to_right_ear_delta | kinematics | 0 | 0 | 0 |
| dist_back_right_paw_to_right_ear_energy | kinematics | 0 | 0 | 0 |
| dist_back_right_paw_to_right_ear_max | pose geometry | 0 | 0 | 0 |
| dist_back_right_paw_to_right_ear_mean | pose geometry | 0 | 0 | 0 |
| dist_back_right_paw_to_right_ear_median | pose geometry | 0 | 0 | 0 |
| dist_back_right_paw_to_right_ear_norm_delta | kinematics | 0 | 0 | 0 |
| dist_back_right_paw_to_right_ear_norm_energy | kinematics | 0 | 0 | 0 |
| dist_back_right_paw_to_right_ear_norm_max | pose geometry | 0 | 0 | 0 |
| dist_back_right_paw_to_right_ear_norm_mean | pose geometry | 0 | 0 | 0 |
| dist_back_right_paw_to_right_ear_norm_median | pose geometry | 0 | 0 | 0 |
| dist_back_right_paw_to_right_ear_norm_p10 | pose geometry | 0 | 0 | 0 |
| dist_back_right_paw_to_right_ear_norm_p90 | pose geometry | 0 | 0 | 0 |
| dist_back_right_paw_to_right_ear_norm_periodicity | pose geometry | 0 | 0 | 0 |
| dist_back_right_paw_to_right_ear_norm_std | kinematics | 0 | 0 | 0 |
| dist_back_right_paw_to_right_ear_norm_trend | pose geometry | 0 | 0 | 0 |
| dist_back_right_paw_to_right_ear_p10 | pose geometry | 0 | 0 | 0 |

| | | | | |
|---|---|---|---|---|
| dist_back_right_paw_to_right_ear_p90 | pose geometry | 0 | 0 | 0 |
| dist_back_right_paw_to_right_ear_periodicity | pose geometry | 0 | 0 | 0 |
| dist_back_right_paw_to_right_ear_std | kinematics | 0 | 0 | 0 |
| dist_back_right_paw_to_right_ear_trend | pose geometry | 0 | 0 | 0 |
| dist_back_right_paw_to_tail_base_delta | kinematics | 0 | 0 | 0 |
| dist_back_right_paw_to_tail_base_energy | kinematics | 0 | 0 | 0 |
| dist_back_right_paw_to_tail_base_max | pose geometry | 0 | 0 | 0 |
| dist_back_right_paw_to_tail_base_mean | pose geometry | 0 | 0 | 0 |
| dist_back_right_paw_to_tail_base_median | pose geometry | 0 | 0 | 0 |
| dist_back_right_paw_to_tail_base_norm_delta | kinematics | 0 | 0 | 0 |
| dist_back_right_paw_to_tail_base_norm_energy | kinematics | 0 | 0 | 0 |
| dist_back_right_paw_to_tail_base_norm_max | pose geometry | 0 | 0 | 0 |
| dist_back_right_paw_to_tail_base_norm_mean | pose geometry | 0 | 0 | 0 |
| dist_back_right_paw_to_tail_base_norm_median | pose geometry | 0 | 0 | 0 |
| dist_back_right_paw_to_tail_base_norm_p10 | pose geometry | 0 | 0 | 0 |
| dist_back_right_paw_to_tail_base_norm_p90 | pose geometry | 0 | 0 | 0 |
| dist_back_right_paw_to_tail_base_norm_periodicity | pose geometry | 0 | 0 | 0 |
| dist_back_right_paw_to_tail_base_norm_std | kinematics | 0 | 0 | 0 |
| dist_back_right_paw_to_tail_base_norm_trend | pose geometry | 0 | 0 | 0 |
| dist_back_right_paw_to_tail_base_p10 | pose geometry | 0 | 0 | 0 |
| dist_back_right_paw_to_tail_base_p90 | pose geometry | 0 | 0 | 0 |
| dist_back_right_paw_to_tail_base_periodicity | pose geometry | 0 | 0 | 0 |
| dist_back_right_paw_to_tail_base_std | kinematics | 0 | 0 | 0 |
| dist_back_right_paw_to_tail_base_trend | pose geometry | 0 | 0 | 0 |
| dist_body_left_to_body_mid_delta | kinematics | 0 | 1 | 0 |
| dist_body_left_to_body_mid_energy | kinematics | 0 | 1 | 0 |
| dist_body_left_to_body_mid_max | pose geometry | 0 | 1 | 0 |
| dist_body_left_to_body_mid_mean | pose geometry | 0 | 1 | 0 |
| dist_body_left_to_body_mid_median | pose geometry | 0 | 1 | 0 |
| dist_body_left_to_body_mid_norm_delta | kinematics | 0 | 1 | 0 |
| dist_body_left_to_body_mid_norm_energy | kinematics | 0 | 1 | 0 |
| dist_body_left_to_body_mid_norm_max | pose geometry | 0 | 1 | 0 |
| dist_body_left_to_body_mid_norm_mean | pose geometry | 0 | 1 | 0 |
| dist_body_left_to_body_mid_norm_median | pose geometry | 0 | 1 | 0 |
| dist_body_left_to_body_mid_norm_p10 | pose geometry | 0 | 1 | 0 |
| dist_body_left_to_body_mid_norm_p90 | pose geometry | 0 | 1 | 0 |
| dist_body_left_to_body_mid_norm_periodicity | pose geometry | 0 | 1 | 0 |
| dist_body_left_to_body_mid_norm_std | kinematics | 0 | 1 | 0 |
| dist_body_left_to_body_mid_norm_trend | pose geometry | 0 | 1 | 0 |
| dist_body_left_to_body_mid_p10 | pose geometry | 0 | 1 | 0 |
| dist_body_left_to_body_mid_p90 | pose geometry | 0 | 1 | 0 |
| dist_body_left_to_body_mid_periodicity | pose geometry | 0 | 1 | 0 |
| dist_body_left_to_body_mid_std | kinematics | 0 | 1 | 0 |
| dist_body_left_to_body_mid_trend | pose geometry | 0 | 1 | 0 |

| | | | | |
|---|---|---|---|---|
| dist_body_left_to_body_right_delta | kinematics | 0 | 1 | 0 |
| dist_body_left_to_body_right_energy | kinematics | 0 | 1 | 0 |
| dist_body_left_to_body_right_max | pose geometry | 0 | 1 | 0 |
| dist_body_left_to_body_right_mean | pose geometry | 0 | 1 | 0 |
| dist_body_left_to_body_right_median | pose geometry | 0 | 1 | 0 |
| dist_body_left_to_body_right_norm_delta | kinematics | 0 | 1 | 0 |
| dist_body_left_to_body_right_norm_energy | kinematics | 0 | 1 | 0 |
| dist_body_left_to_body_right_norm_max | pose geometry | 0 | 1 | 0 |
| dist_body_left_to_body_right_norm_mean | pose geometry | 0 | 1 | 0 |
| dist_body_left_to_body_right_norm_median | pose geometry | 0 | 1 | 0 |
| dist_body_left_to_body_right_norm_p10 | pose geometry | 0 | 1 | 0 |
| dist_body_left_to_body_right_norm_p90 | pose geometry | 0 | 1 | 0 |
| dist_body_left_to_body_right_norm_periodicity | pose geometry | 0 | 1 | 0 |
| dist_body_left_to_body_right_norm_std | kinematics | 0 | 1 | 0 |
| dist_body_left_to_body_right_norm_trend | pose geometry | 0 | 1 | 0 |
| dist_body_left_to_body_right_p10 | pose geometry | 0 | 1 | 0 |
| dist_body_left_to_body_right_p90 | pose geometry | 0 | 1 | 0 |
| dist_body_left_to_body_right_periodicity | pose geometry | 0 | 1 | 0 |
| dist_body_left_to_body_right_std | kinematics | 0 | 1 | 0 |
| dist_body_left_to_body_right_trend | pose geometry | 0 | 1 | 0 |
| dist_body_left_to_ear_left_delta | kinematics | 0 | 1 | 0 |
| dist_body_left_to_ear_left_energy | kinematics | 0 | 1 | 0 |
| dist_body_left_to_ear_left_max | pose geometry | 0 | 1 | 0 |
| dist_body_left_to_ear_left_mean | pose geometry | 0 | 1 | 0 |
| dist_body_left_to_ear_left_median | pose geometry | 0 | 1 | 0 |
| dist_body_left_to_ear_left_norm_delta | kinematics | 0 | 1 | 0 |
| dist_body_left_to_ear_left_norm_energy | kinematics | 0 | 1 | 0 |
| dist_body_left_to_ear_left_norm_max | pose geometry | 0 | 1 | 0 |
| dist_body_left_to_ear_left_norm_mean | pose geometry | 0 | 1 | 0 |
| dist_body_left_to_ear_left_norm_median | pose geometry | 0 | 1 | 0 |
| dist_body_left_to_ear_left_norm_p10 | pose geometry | 0 | 1 | 0 |
| dist_body_left_to_ear_left_norm_p90 | pose geometry | 0 | 1 | 0 |
| dist_body_left_to_ear_left_norm_periodicity | pose geometry | 0 | 1 | 0 |
| dist_body_left_to_ear_left_norm_std | kinematics | 0 | 1 | 0 |
| dist_body_left_to_ear_left_norm_trend | pose geometry | 0 | 1 | 0 |
| dist_body_left_to_ear_left_p10 | pose geometry | 0 | 1 | 0 |
| dist_body_left_to_ear_left_p90 | pose geometry | 0 | 1 | 0 |
| dist_body_left_to_ear_left_periodicity | pose geometry | 0 | 1 | 0 |
| dist_body_left_to_ear_left_std | kinematics | 0 | 1 | 0 |
| dist_body_left_to_ear_left_trend | pose geometry | 0 | 1 | 0 |
| dist_body_left_to_ear_right_delta | kinematics | 0 | 1 | 0 |
| dist_body_left_to_ear_right_energy | kinematics | 0 | 1 | 0 |
| dist_body_left_to_ear_right_max | pose geometry | 0 | 1 | 0 |
| dist_body_left_to_ear_right_mean | pose geometry | 0 | 1 | 0 |

| | | | | |
|---|---|---|---|---|
| dist_body_left_to_ear_right_median | pose geometry | 0 | 1 | 0 |
| dist_body_left_to_ear_right_norm_delta | kinematics | 0 | 1 | 0 |
| dist_body_left_to_ear_right_norm_energy | kinematics | 0 | 1 | 0 |
| dist_body_left_to_ear_right_norm_max | pose geometry | 0 | 1 | 0 |
| dist_body_left_to_ear_right_norm_mean | pose geometry | 0 | 1 | 0 |
| dist_body_left_to_ear_right_norm_median | pose geometry | 0 | 1 | 0 |
| dist_body_left_to_ear_right_norm_p10 | pose geometry | 0 | 1 | 0 |
| dist_body_left_to_ear_right_norm_p90 | pose geometry | 0 | 1 | 0 |
| dist_body_left_to_ear_right_norm_periodicity | pose geometry | 0 | 1 | 0 |
| dist_body_left_to_ear_right_norm_std | kinematics | 0 | 1 | 0 |
| dist_body_left_to_ear_right_norm_trend | pose geometry | 0 | 1 | 0 |
| dist_body_left_to_ear_right_p10 | pose geometry | 0 | 1 | 0 |
| dist_body_left_to_ear_right_p90 | pose geometry | 0 | 1 | 0 |
| dist_body_left_to_ear_right_periodicity | pose geometry | 0 | 1 | 0 |
| dist_body_left_to_ear_right_std | kinematics | 0 | 1 | 0 |
| dist_body_left_to_ear_right_trend | pose geometry | 0 | 1 | 0 |
| dist_body_left_to_nose_delta | kinematics | 0 | 1 | 0 |
| dist_body_left_to_nose_energy | kinematics | 0 | 1 | 0 |
| dist_body_left_to_nose_max | pose geometry | 0 | 1 | 0 |
| dist_body_left_to_nose_mean | pose geometry | 0 | 1 | 0 |
| dist_body_left_to_nose_median | pose geometry | 0 | 1 | 0 |
| dist_body_left_to_nose_norm_delta | kinematics | 0 | 1 | 0 |
| dist_body_left_to_nose_norm_energy | kinematics | 0 | 1 | 0 |
| dist_body_left_to_nose_norm_max | pose geometry | 0 | 1 | 0 |
| dist_body_left_to_nose_norm_mean | pose geometry | 0 | 1 | 0 |
| dist_body_left_to_nose_norm_median | pose geometry | 0 | 1 | 0 |
| dist_body_left_to_nose_norm_p10 | pose geometry | 0 | 1 | 0 |
| dist_body_left_to_nose_norm_p90 | pose geometry | 0 | 1 | 0 |
| dist_body_left_to_nose_norm_periodicity | pose geometry | 0 | 1 | 0 |
| dist_body_left_to_nose_norm_std | kinematics | 0 | 1 | 0 |
| dist_body_left_to_nose_norm_trend | pose geometry | 0 | 1 | 0 |
| dist_body_left_to_nose_p10 | pose geometry | 0 | 1 | 0 |
| dist_body_left_to_nose_p90 | pose geometry | 0 | 1 | 0 |
| dist_body_left_to_nose_periodicity | pose geometry | 0 | 1 | 0 |
| dist_body_left_to_nose_std | kinematics | 0 | 1 | 0 |
| dist_body_left_to_nose_trend | pose geometry | 0 | 1 | 0 |
| dist_body_left_to_tail_base_delta | kinematics | 0 | 1 | 0 |
| dist_body_left_to_tail_base_energy | kinematics | 0 | 1 | 0 |
| dist_body_left_to_tail_base_max | pose geometry | 0 | 1 | 0 |
| dist_body_left_to_tail_base_mean | pose geometry | 0 | 1 | 0 |
| dist_body_left_to_tail_base_median | pose geometry | 0 | 1 | 0 |
| dist_body_left_to_tail_base_norm_delta | kinematics | 0 | 1 | 0 |
| dist_body_left_to_tail_base_norm_energy | kinematics | 0 | 1 | 0 |
| dist_body_left_to_tail_base_norm_max | pose geometry | 0 | 1 | 0 |

| | | | | |
|---|---|---|---|---|
| dist_body_left_to_tail_base_norm_mean | pose geometry | 0 | 1 | 0 |
| dist_body_left_to_tail_base_norm_median | pose geometry | 0 | 1 | 0 |
| dist_body_left_to_tail_base_norm_p10 | pose geometry | 0 | 1 | 0 |
| dist_body_left_to_tail_base_norm_p90 | pose geometry | 0 | 1 | 0 |
| dist_body_left_to_tail_base_norm_periodicity | pose geometry | 0 | 1 | 0 |
| dist_body_left_to_tail_base_norm_std | kinematics | 0 | 1 | 0 |
| dist_body_left_to_tail_base_norm_trend | pose geometry | 0 | 1 | 0 |
| dist_body_left_to_tail_base_p10 | pose geometry | 0 | 1 | 0 |
| dist_body_left_to_tail_base_p90 | pose geometry | 0 | 1 | 0 |
| dist_body_left_to_tail_base_periodicity | pose geometry | 0 | 1 | 0 |
| dist_body_left_to_tail_base_std | kinematics | 0 | 1 | 0 |
| dist_body_left_to_tail_base_trend | pose geometry | 0 | 1 | 0 |
| dist_body_mid_to_body_right_delta | kinematics | 0 | 1 | 0 |
| dist_body_mid_to_body_right_energy | kinematics | 0 | 1 | 0 |
| dist_body_mid_to_body_right_max | pose geometry | 0 | 1 | 0 |
| dist_body_mid_to_body_right_mean | pose geometry | 0 | 1 | 0 |
| dist_body_mid_to_body_right_median | pose geometry | 0 | 1 | 0 |
| dist_body_mid_to_body_right_norm_delta | kinematics | 0 | 1 | 0 |
| dist_body_mid_to_body_right_norm_energy | kinematics | 0 | 1 | 0 |
| dist_body_mid_to_body_right_norm_max | pose geometry | 0 | 1 | 0 |
| dist_body_mid_to_body_right_norm_mean | pose geometry | 0 | 1 | 0 |
| dist_body_mid_to_body_right_norm_median | pose geometry | 0 | 1 | 0 |
| dist_body_mid_to_body_right_norm_p10 | pose geometry | 0 | 1 | 0 |
| dist_body_mid_to_body_right_norm_p90 | pose geometry | 0 | 1 | 0 |
| dist_body_mid_to_body_right_norm_periodicity | pose geometry | 0 | 1 | 0 |
| dist_body_mid_to_body_right_norm_std | kinematics | 0 | 1 | 0 |
| dist_body_mid_to_body_right_norm_trend | pose geometry | 0 | 1 | 0 |
| dist_body_mid_to_body_right_p10 | pose geometry | 0 | 1 | 0 |
| dist_body_mid_to_body_right_p90 | pose geometry | 0 | 1 | 0 |
| dist_body_mid_to_body_right_periodicity | pose geometry | 0 | 1 | 0 |
| dist_body_mid_to_body_right_std | kinematics | 0 | 1 | 0 |
| dist_body_mid_to_body_right_trend | pose geometry | 0 | 1 | 0 |
| dist_body_mid_to_ear_left_delta | kinematics | 0 | 1 | 0 |
| dist_body_mid_to_ear_left_energy | kinematics | 0 | 1 | 0 |
| dist_body_mid_to_ear_left_max | pose geometry | 0 | 1 | 0 |
| dist_body_mid_to_ear_left_mean | pose geometry | 0 | 1 | 0 |
| dist_body_mid_to_ear_left_median | pose geometry | 0 | 1 | 0 |
| dist_body_mid_to_ear_left_norm_delta | kinematics | 0 | 1 | 0 |
| dist_body_mid_to_ear_left_norm_energy | kinematics | 0 | 1 | 0 |
| dist_body_mid_to_ear_left_norm_max | pose geometry | 0 | 1 | 0 |
| dist_body_mid_to_ear_left_norm_mean | pose geometry | 0 | 1 | 0 |
| dist_body_mid_to_ear_left_norm_median | pose geometry | 0 | 1 | 0 |
| dist_body_mid_to_ear_left_norm_p10 | pose geometry | 0 | 1 | 0 |
| dist_body_mid_to_ear_left_norm_p90 | pose geometry | 0 | 1 | 0 |

| | | | | |
|---|---|---|---|---|
| dist_body_mid_to_ear_left_norm_periodicity | pose geometry | 0 | 1 | 0 |
| dist_body_mid_to_ear_left_norm_std | kinematics | 0 | 1 | 0 |
| dist_body_mid_to_ear_left_norm_trend | pose geometry | 0 | 1 | 0 |
| dist_body_mid_to_ear_left_p10 | pose geometry | 0 | 1 | 0 |
| dist_body_mid_to_ear_left_p90 | pose geometry | 0 | 1 | 0 |
| dist_body_mid_to_ear_left_periodicity | pose geometry | 0 | 1 | 0 |
| dist_body_mid_to_ear_left_std | kinematics | 0 | 1 | 0 |
| dist_body_mid_to_ear_left_trend | pose geometry | 0 | 1 | 0 |
| dist_body_mid_to_ear_right_delta | kinematics | 0 | 1 | 0 |
| dist_body_mid_to_ear_right_energy | kinematics | 0 | 1 | 0 |
| dist_body_mid_to_ear_right_max | pose geometry | 0 | 1 | 0 |
| dist_body_mid_to_ear_right_mean | pose geometry | 0 | 1 | 0 |
| dist_body_mid_to_ear_right_median | pose geometry | 0 | 1 | 0 |
| dist_body_mid_to_ear_right_norm_delta | kinematics | 0 | 1 | 0 |
| dist_body_mid_to_ear_right_norm_energy | kinematics | 0 | 1 | 0 |
| dist_body_mid_to_ear_right_norm_max | pose geometry | 0 | 1 | 0 |
| dist_body_mid_to_ear_right_norm_mean | pose geometry | 0 | 1 | 0 |
| dist_body_mid_to_ear_right_norm_median | pose geometry | 0 | 1 | 0 |
| dist_body_mid_to_ear_right_norm_p10 | pose geometry | 0 | 1 | 0 |
| dist_body_mid_to_ear_right_norm_p90 | pose geometry | 0 | 1 | 0 |
| dist_body_mid_to_ear_right_norm_periodicity | pose geometry | 0 | 1 | 0 |
| dist_body_mid_to_ear_right_norm_std | kinematics | 0 | 1 | 0 |
| dist_body_mid_to_ear_right_norm_trend | pose geometry | 0 | 1 | 0 |
| dist_body_mid_to_ear_right_p10 | pose geometry | 0 | 1 | 0 |
| dist_body_mid_to_ear_right_p90 | pose geometry | 0 | 1 | 0 |
| dist_body_mid_to_ear_right_periodicity | pose geometry | 0 | 1 | 0 |
| dist_body_mid_to_ear_right_std | kinematics | 0 | 1 | 0 |
| dist_body_mid_to_ear_right_trend | pose geometry | 0 | 1 | 0 |
| dist_body_mid_to_nose_delta | kinematics | 0 | 1 | 0 |
| dist_body_mid_to_nose_energy | kinematics | 0 | 1 | 0 |
| dist_body_mid_to_nose_max | pose geometry | 0 | 1 | 0 |
| dist_body_mid_to_nose_mean | pose geometry | 0 | 1 | 0 |
| dist_body_mid_to_nose_median | pose geometry | 0 | 1 | 0 |
| dist_body_mid_to_nose_norm_delta | kinematics | 0 | 1 | 0 |
| dist_body_mid_to_nose_norm_energy | kinematics | 0 | 1 | 0 |
| dist_body_mid_to_nose_norm_max | pose geometry | 0 | 1 | 0 |
| dist_body_mid_to_nose_norm_mean | pose geometry | 0 | 1 | 0 |
| dist_body_mid_to_nose_norm_median | pose geometry | 0 | 1 | 0 |
| dist_body_mid_to_nose_norm_p10 | pose geometry | 0 | 1 | 0 |
| dist_body_mid_to_nose_norm_p90 | pose geometry | 0 | 1 | 0 |
| dist_body_mid_to_nose_norm_periodicity | pose geometry | 0 | 1 | 0 |
| dist_body_mid_to_nose_norm_std | kinematics | 0 | 1 | 0 |
| dist_body_mid_to_nose_norm_trend | pose geometry | 0 | 1 | 0 |
| dist_body_mid_to_nose_p10 | pose geometry | 0 | 1 | 0 |

| | | | | |
|---|---|---|---|---|
| dist_body_mid_to_nose_p90 | pose geometry | 0 | 1 | 0 |
| dist_body_mid_to_nose_periodicity | pose geometry | 0 | 1 | 0 |
| dist_body_mid_to_nose_std | kinematics | 0 | 1 | 0 |
| dist_body_mid_to_nose_trend | pose geometry | 0 | 1 | 0 |
| dist_body_mid_to_tail_base_delta | kinematics | 0 | 1 | 0 |
| dist_body_mid_to_tail_base_energy | kinematics | 0 | 1 | 0 |
| dist_body_mid_to_tail_base_max | pose geometry | 0 | 1 | 0 |
| dist_body_mid_to_tail_base_mean | pose geometry | 0 | 1 | 0 |
| dist_body_mid_to_tail_base_median | pose geometry | 0 | 1 | 0 |
| dist_body_mid_to_tail_base_norm_delta | kinematics | 0 | 1 | 0 |
| dist_body_mid_to_tail_base_norm_energy | kinematics | 0 | 1 | 0 |
| dist_body_mid_to_tail_base_norm_max | pose geometry | 0 | 1 | 0 |
| dist_body_mid_to_tail_base_norm_mean | pose geometry | 0 | 1 | 0 |
| dist_body_mid_to_tail_base_norm_median | pose geometry | 0 | 1 | 0 |
| dist_body_mid_to_tail_base_norm_p10 | pose geometry | 0 | 1 | 0 |
| dist_body_mid_to_tail_base_norm_p90 | pose geometry | 0 | 1 | 0 |
| dist_body_mid_to_tail_base_norm_periodicity | pose geometry | 0 | 1 | 0 |
| dist_body_mid_to_tail_base_norm_std | kinematics | 0 | 1 | 0 |
| dist_body_mid_to_tail_base_norm_trend | pose geometry | 0 | 1 | 0 |
| dist_body_mid_to_tail_base_p10 | pose geometry | 0 | 1 | 0 |
| dist_body_mid_to_tail_base_p90 | pose geometry | 0 | 1 | 0 |
| dist_body_mid_to_tail_base_periodicity | pose geometry | 0 | 1 | 0 |
| dist_body_mid_to_tail_base_std | kinematics | 0 | 1 | 0 |
| dist_body_mid_to_tail_base_trend | pose geometry | 0 | 1 | 0 |
| dist_body_right_to_ear_left_delta | kinematics | 0 | 1 | 0 |
| dist_body_right_to_ear_left_energy | kinematics | 0 | 1 | 0 |
| dist_body_right_to_ear_left_max | pose geometry | 0 | 1 | 0 |
| dist_body_right_to_ear_left_mean | pose geometry | 0 | 1 | 0 |
| dist_body_right_to_ear_left_median | pose geometry | 0 | 1 | 0 |
| dist_body_right_to_ear_left_norm_delta | kinematics | 0 | 1 | 0 |
| dist_body_right_to_ear_left_norm_energy | kinematics | 0 | 1 | 0 |
| dist_body_right_to_ear_left_norm_max | pose geometry | 0 | 1 | 0 |
| dist_body_right_to_ear_left_norm_mean | pose geometry | 0 | 1 | 0 |
| dist_body_right_to_ear_left_norm_median | pose geometry | 0 | 1 | 0 |
| dist_body_right_to_ear_left_norm_p10 | pose geometry | 0 | 1 | 0 |
| dist_body_right_to_ear_left_norm_p90 | pose geometry | 0 | 1 | 0 |
| dist_body_right_to_ear_left_norm_periodicity | pose geometry | 0 | 1 | 0 |
| dist_body_right_to_ear_left_norm_std | kinematics | 0 | 1 | 0 |
| dist_body_right_to_ear_left_norm_trend | pose geometry | 0 | 1 | 0 |
| dist_body_right_to_ear_left_p10 | pose geometry | 0 | 1 | 0 |
| dist_body_right_to_ear_left_p90 | pose geometry | 0 | 1 | 0 |
| dist_body_right_to_ear_left_periodicity | pose geometry | 0 | 1 | 0 |
| dist_body_right_to_ear_left_std | kinematics | 0 | 1 | 0 |
| dist_body_right_to_ear_left_trend | pose geometry | 0 | 1 | 0 |

| | | | | |
|---|---|---|---|---|
| dist_body_right_to_ear_right_delta | kinematics | 0 | 1 | 0 |
| dist_body_right_to_ear_right_energy | kinematics | 0 | 1 | 0 |
| dist_body_right_to_ear_right_max | pose geometry | 0 | 1 | 0 |
| dist_body_right_to_ear_right_mean | pose geometry | 0 | 1 | 0 |
| dist_body_right_to_ear_right_median | pose geometry | 0 | 1 | 0 |
| dist_body_right_to_ear_right_norm_delta | kinematics | 0 | 1 | 0 |
| dist_body_right_to_ear_right_norm_energy | kinematics | 0 | 1 | 0 |
| dist_body_right_to_ear_right_norm_max | pose geometry | 0 | 1 | 0 |
| dist_body_right_to_ear_right_norm_mean | pose geometry | 0 | 1 | 0 |
| dist_body_right_to_ear_right_norm_median | pose geometry | 0 | 1 | 0 |
| dist_body_right_to_ear_right_norm_p10 | pose geometry | 0 | 1 | 0 |
| dist_body_right_to_ear_right_norm_p90 | pose geometry | 0 | 1 | 0 |
| dist_body_right_to_ear_right_norm_periodicity | pose geometry | 0 | 1 | 0 |
| dist_body_right_to_ear_right_norm_std | kinematics | 0 | 1 | 0 |
| dist_body_right_to_ear_right_norm_trend | pose geometry | 0 | 1 | 0 |
| dist_body_right_to_ear_right_p10 | pose geometry | 0 | 1 | 0 |
| dist_body_right_to_ear_right_p90 | pose geometry | 0 | 1 | 0 |
| dist_body_right_to_ear_right_periodicity | pose geometry | 0 | 1 | 0 |
| dist_body_right_to_ear_right_std | kinematics | 0 | 1 | 0 |
| dist_body_right_to_ear_right_trend | pose geometry | 0 | 1 | 0 |
| dist_body_right_to_nose_delta | kinematics | 0 | 1 | 0 |
| dist_body_right_to_nose_energy | kinematics | 0 | 1 | 0 |
| dist_body_right_to_nose_max | pose geometry | 0 | 1 | 0 |
| dist_body_right_to_nose_mean | pose geometry | 0 | 1 | 0 |
| dist_body_right_to_nose_median | pose geometry | 0 | 1 | 0 |
| dist_body_right_to_nose_norm_delta | kinematics | 0 | 1 | 0 |
| dist_body_right_to_nose_norm_energy | kinematics | 0 | 1 | 0 |
| dist_body_right_to_nose_norm_max | pose geometry | 0 | 1 | 0 |
| dist_body_right_to_nose_norm_mean | pose geometry | 0 | 1 | 0 |
| dist_body_right_to_nose_norm_median | pose geometry | 0 | 1 | 0 |
| dist_body_right_to_nose_norm_p10 | pose geometry | 0 | 1 | 0 |
| dist_body_right_to_nose_norm_p90 | pose geometry | 0 | 1 | 0 |
| dist_body_right_to_nose_norm_periodicity | pose geometry | 0 | 1 | 0 |
| dist_body_right_to_nose_norm_std | kinematics | 0 | 1 | 0 |
| dist_body_right_to_nose_norm_trend | pose geometry | 0 | 1 | 0 |
| dist_body_right_to_nose_p10 | pose geometry | 0 | 1 | 0 |
| dist_body_right_to_nose_p90 | pose geometry | 0 | 1 | 0 |
| dist_body_right_to_nose_periodicity | pose geometry | 0 | 1 | 0 |
| dist_body_right_to_nose_std | kinematics | 0 | 1 | 0 |
| dist_body_right_to_nose_trend | pose geometry | 0 | 1 | 0 |
| dist_body_right_to_tail_base_delta | kinematics | 0 | 1 | 0 |
| dist_body_right_to_tail_base_energy | kinematics | 0 | 1 | 0 |
| dist_body_right_to_tail_base_max | pose geometry | 0 | 1 | 0 |
| dist_body_right_to_tail_base_mean | pose geometry | 0 | 1 | 0 |

| | | | | |
|---|---|---|---|---|
| dist_body_right_to_tail_base_median | pose geometry | 0 | 1 | 0 |
| dist_body_right_to_tail_base_norm_delta | kinematics | 0 | 1 | 0 |
| dist_body_right_to_tail_base_norm_energy | kinematics | 0 | 1 | 0 |
| dist_body_right_to_tail_base_norm_max | pose geometry | 0 | 1 | 0 |
| dist_body_right_to_tail_base_norm_mean | pose geometry | 0 | 1 | 0 |
| dist_body_right_to_tail_base_norm_median | pose geometry | 0 | 1 | 0 |
| dist_body_right_to_tail_base_norm_p10 | pose geometry | 0 | 1 | 0 |
| dist_body_right_to_tail_base_norm_p90 | pose geometry | 0 | 1 | 0 |
| dist_body_right_to_tail_base_norm_periodicity | pose geometry | 0 | 1 | 0 |
| dist_body_right_to_tail_base_norm_std | kinematics | 0 | 1 | 0 |
| dist_body_right_to_tail_base_norm_trend | pose geometry | 0 | 1 | 0 |
| dist_body_right_to_tail_base_p10 | pose geometry | 0 | 1 | 0 |
| dist_body_right_to_tail_base_p90 | pose geometry | 0 | 1 | 0 |
| dist_body_right_to_tail_base_periodicity | pose geometry | 0 | 1 | 0 |
| dist_body_right_to_tail_base_std | kinematics | 0 | 1 | 0 |
| dist_body_right_to_tail_base_trend | pose geometry | 0 | 1 | 0 |
| dist_center_body_to_left_body_delta | context (ROI/target) | 1 | 0 | 0 |
| dist_center_body_to_left_body_energy | context (ROI/target) | 1 | 0 | 0 |
| dist_center_body_to_left_body_max | context (ROI/target) | 1 | 0 | 0 |
| dist_center_body_to_left_body_mean | context (ROI/target) | 1 | 0 | 0 |
| dist_center_body_to_left_body_median | context (ROI/target) | 1 | 0 | 0 |
| dist_center_body_to_left_body_norm_delta | context (ROI/target) | 1 | 0 | 0 |
| dist_center_body_to_left_body_norm_energy | context (ROI/target) | 1 | 0 | 0 |
| dist_center_body_to_left_body_norm_max | context (ROI/target) | 1 | 0 | 0 |
| dist_center_body_to_left_body_norm_mean | context (ROI/target) | 1 | 0 | 0 |
| dist_center_body_to_left_body_norm_median | context (ROI/target) | 1 | 0 | 0 |
| dist_center_body_to_left_body_norm_p10 | context (ROI/target) | 1 | 0 | 0 |
| dist_center_body_to_left_body_norm_p90 | context (ROI/target) | 1 | 0 | 0 |
| dist_center_body_to_left_body_norm_periodicity | context (ROI/target) | 1 | 0 | 0 |
| dist_center_body_to_left_body_norm_std | context (ROI/target) | 1 | 0 | 0 |
| dist_center_body_to_left_body_norm_trend | context (ROI/target) | 1 | 0 | 0 |
| dist_center_body_to_left_body_p10 | context (ROI/target) | 1 | 0 | 0 |
| dist_center_body_to_left_body_p90 | context (ROI/target) | 1 | 0 | 0 |
| dist_center_body_to_left_body_periodicity | context (ROI/target) | 1 | 0 | 0 |
| dist_center_body_to_left_body_std | context (ROI/target) | 1 | 0 | 0 |
| dist_center_body_to_left_body_trend | context (ROI/target) | 1 | 0 | 0 |
| dist_center_body_to_left_ear_delta | context (ROI/target) | 1 | 0 | 0 |
| dist_center_body_to_left_ear_energy | context (ROI/target) | 1 | 0 | 0 |
| dist_center_body_to_left_ear_max | context (ROI/target) | 1 | 0 | 0 |
| dist_center_body_to_left_ear_mean | context (ROI/target) | 1 | 0 | 0 |
| dist_center_body_to_left_ear_median | context (ROI/target) | 1 | 0 | 0 |
| dist_center_body_to_left_ear_norm_delta | context (ROI/target) | 1 | 0 | 0 |
| dist_center_body_to_left_ear_norm_energy | context (ROI/target) | 1 | 0 | 0 |
| dist_center_body_to_left_ear_norm_max | context (ROI/target) | 1 | 0 | 0 |

| | | | | |
|---|---|---|---|---|
| dist_center_body_to_left_ear_norm_mean | context (ROI/target) | 1 | 0 | 0 |
| dist_center_body_to_left_ear_norm_median | context (ROI/target) | 1 | 0 | 0 |
| dist_center_body_to_left_ear_norm_p10 | context (ROI/target) | 1 | 0 | 0 |
| dist_center_body_to_left_ear_norm_p90 | context (ROI/target) | 1 | 0 | 0 |
| dist_center_body_to_left_ear_norm_periodicity | context (ROI/target) | 1 | 0 | 0 |
| dist_center_body_to_left_ear_norm_std | context (ROI/target) | 1 | 0 | 0 |
| dist_center_body_to_left_ear_norm_trend | context (ROI/target) | 1 | 0 | 0 |
| dist_center_body_to_left_ear_p10 | context (ROI/target) | 1 | 0 | 0 |
| dist_center_body_to_left_ear_p90 | context (ROI/target) | 1 | 0 | 0 |
| dist_center_body_to_left_ear_periodicity | context (ROI/target) | 1 | 0 | 0 |
| dist_center_body_to_left_ear_std | context (ROI/target) | 1 | 0 | 0 |
| dist_center_body_to_left_ear_trend | context (ROI/target) | 1 | 0 | 0 |
| dist_center_body_to_nose_delta | context (ROI/target) | 1 | 0 | 0 |
| dist_center_body_to_nose_energy | context (ROI/target) | 1 | 0 | 0 |
| dist_center_body_to_nose_max | context (ROI/target) | 1 | 0 | 0 |
| dist_center_body_to_nose_mean | context (ROI/target) | 1 | 0 | 0 |
| dist_center_body_to_nose_median | context (ROI/target) | 1 | 0 | 0 |
| dist_center_body_to_nose_norm_delta | context (ROI/target) | 1 | 0 | 0 |
| dist_center_body_to_nose_norm_energy | context (ROI/target) | 1 | 0 | 0 |
| dist_center_body_to_nose_norm_max | context (ROI/target) | 1 | 0 | 0 |
| dist_center_body_to_nose_norm_mean | context (ROI/target) | 1 | 0 | 0 |
| dist_center_body_to_nose_norm_median | context (ROI/target) | 1 | 0 | 0 |
| dist_center_body_to_nose_norm_p10 | context (ROI/target) | 1 | 0 | 0 |
| dist_center_body_to_nose_norm_p90 | context (ROI/target) | 1 | 0 | 0 |
| dist_center_body_to_nose_norm_periodicity | context (ROI/target) | 1 | 0 | 0 |
| dist_center_body_to_nose_norm_std | context (ROI/target) | 1 | 0 | 0 |
| dist_center_body_to_nose_norm_trend | context (ROI/target) | 1 | 0 | 0 |
| dist_center_body_to_nose_p10 | context (ROI/target) | 1 | 0 | 0 |
| dist_center_body_to_nose_p90 | context (ROI/target) | 1 | 0 | 0 |
| dist_center_body_to_nose_periodicity | context (ROI/target) | 1 | 0 | 0 |
| dist_center_body_to_nose_std | context (ROI/target) | 1 | 0 | 0 |
| dist_center_body_to_nose_trend | context (ROI/target) | 1 | 0 | 0 |
| dist_center_body_to_right_body_delta | context (ROI/target) | 1 | 0 | 0 |
| dist_center_body_to_right_body_energy | context (ROI/target) | 1 | 0 | 0 |
| dist_center_body_to_right_body_max | context (ROI/target) | 1 | 0 | 0 |
| dist_center_body_to_right_body_mean | context (ROI/target) | 1 | 0 | 0 |
| dist_center_body_to_right_body_median | context (ROI/target) | 1 | 0 | 0 |
| dist_center_body_to_right_body_norm_delta | context (ROI/target) | 1 | 0 | 0 |
| dist_center_body_to_right_body_norm_energy | context (ROI/target) | 1 | 0 | 0 |
| dist_center_body_to_right_body_norm_max | context (ROI/target) | 1 | 0 | 0 |
| dist_center_body_to_right_body_norm_mean | context (ROI/target) | 1 | 0 | 0 |
| dist_center_body_to_right_body_norm_median | context (ROI/target) | 1 | 0 | 0 |
| dist_center_body_to_right_body_norm_p10 | context (ROI/target) | 1 | 0 | 0 |
| dist_center_body_to_right_body_norm_p90 | context (ROI/target) | 1 | 0 | 0 |

| | | | | |
|---|---|---|---|---|
| dist_center_body_to_right_body_norm_periodicity | context (ROI/target) | 1 | 0 | 0 |
| dist_center_body_to_right_body_norm_std | context (ROI/target) | 1 | 0 | 0 |
| dist_center_body_to_right_body_norm_trend | context (ROI/target) | 1 | 0 | 0 |
| dist_center_body_to_right_body_p10 | context (ROI/target) | 1 | 0 | 0 |
| dist_center_body_to_right_body_p90 | context (ROI/target) | 1 | 0 | 0 |
| dist_center_body_to_right_body_periodicity | context (ROI/target) | 1 | 0 | 0 |
| dist_center_body_to_right_body_std | context (ROI/target) | 1 | 0 | 0 |
| dist_center_body_to_right_body_trend | context (ROI/target) | 1 | 0 | 0 |
| dist_center_body_to_right_ear_delta | context (ROI/target) | 1 | 0 | 0 |
| dist_center_body_to_right_ear_energy | context (ROI/target) | 1 | 0 | 0 |
| dist_center_body_to_right_ear_max | context (ROI/target) | 1 | 0 | 0 |
| dist_center_body_to_right_ear_mean | context (ROI/target) | 1 | 0 | 0 |
| dist_center_body_to_right_ear_median | context (ROI/target) | 1 | 0 | 0 |
| dist_center_body_to_right_ear_norm_delta | context (ROI/target) | 1 | 0 | 0 |
| dist_center_body_to_right_ear_norm_energy | context (ROI/target) | 1 | 0 | 0 |
| dist_center_body_to_right_ear_norm_max | context (ROI/target) | 1 | 0 | 0 |
| dist_center_body_to_right_ear_norm_mean | context (ROI/target) | 1 | 0 | 0 |
| dist_center_body_to_right_ear_norm_median | context (ROI/target) | 1 | 0 | 0 |
| dist_center_body_to_right_ear_norm_p10 | context (ROI/target) | 1 | 0 | 0 |
| dist_center_body_to_right_ear_norm_p90 | context (ROI/target) | 1 | 0 | 0 |
| dist_center_body_to_right_ear_norm_periodicity | context (ROI/target) | 1 | 0 | 0 |
| dist_center_body_to_right_ear_norm_std | context (ROI/target) | 1 | 0 | 0 |
| dist_center_body_to_right_ear_norm_trend | context (ROI/target) | 1 | 0 | 0 |
| dist_center_body_to_right_ear_p10 | context (ROI/target) | 1 | 0 | 0 |
| dist_center_body_to_right_ear_p90 | context (ROI/target) | 1 | 0 | 0 |
| dist_center_body_to_right_ear_periodicity | context (ROI/target) | 1 | 0 | 0 |
| dist_center_body_to_right_ear_std | context (ROI/target) | 1 | 0 | 0 |
| dist_center_body_to_right_ear_trend | context (ROI/target) | 1 | 0 | 0 |
| dist_center_body_to_tail_base_delta | context (ROI/target) | 1 | 0 | 0 |
| dist_center_body_to_tail_base_energy | context (ROI/target) | 1 | 0 | 0 |
| dist_center_body_to_tail_base_max | context (ROI/target) | 1 | 0 | 0 |
| dist_center_body_to_tail_base_mean | context (ROI/target) | 1 | 0 | 0 |
| dist_center_body_to_tail_base_median | context (ROI/target) | 1 | 0 | 0 |
| dist_center_body_to_tail_base_norm_delta | context (ROI/target) | 1 | 0 | 0 |
| dist_center_body_to_tail_base_norm_energy | context (ROI/target) | 1 | 0 | 0 |
| dist_center_body_to_tail_base_norm_max | context (ROI/target) | 1 | 0 | 0 |
| dist_center_body_to_tail_base_norm_mean | context (ROI/target) | 1 | 0 | 0 |
| dist_center_body_to_tail_base_norm_median | context (ROI/target) | 1 | 0 | 0 |
| dist_center_body_to_tail_base_norm_p10 | context (ROI/target) | 1 | 0 | 0 |
| dist_center_body_to_tail_base_norm_p90 | context (ROI/target) | 1 | 0 | 0 |
| dist_center_body_to_tail_base_norm_periodicity | context (ROI/target) | 1 | 0 | 0 |
| dist_center_body_to_tail_base_norm_std | context (ROI/target) | 1 | 0 | 0 |
| dist_center_body_to_tail_base_norm_trend | context (ROI/target) | 1 | 0 | 0 |
| dist_center_body_to_tail_base_p10 | context (ROI/target) | 1 | 0 | 0 |

| | | | | |
|---|---|---|---|---|
| dist_center_body_to_tail_base_p90 | context (ROI/target) | 1 | 0 | 0 |
| dist_center_body_to_tail_base_periodicity | context (ROI/target) | 1 | 0 | 0 |
| dist_center_body_to_tail_base_std | context (ROI/target) | 1 | 0 | 0 |
| dist_center_body_to_tail_base_trend | context (ROI/target) | 1 | 0 | 0 |
| dist_center_to_ear_left_delta | context (ROI/target) | 0 | 0 | 1 |
| dist_center_to_ear_left_energy | context (ROI/target) | 0 | 0 | 1 |
| dist_center_to_ear_left_max | context (ROI/target) | 0 | 0 | 1 |
| dist_center_to_ear_left_mean | context (ROI/target) | 0 | 0 | 1 |
| dist_center_to_ear_left_median | context (ROI/target) | 0 | 0 | 1 |
| dist_center_to_ear_left_norm_delta | context (ROI/target) | 0 | 0 | 1 |
| dist_center_to_ear_left_norm_energy | context (ROI/target) | 0 | 0 | 1 |
| dist_center_to_ear_left_norm_max | context (ROI/target) | 0 | 0 | 1 |
| dist_center_to_ear_left_norm_mean | context (ROI/target) | 0 | 0 | 1 |
| dist_center_to_ear_left_norm_median | context (ROI/target) | 0 | 0 | 1 |
| dist_center_to_ear_left_norm_p10 | context (ROI/target) | 0 | 0 | 1 |
| dist_center_to_ear_left_norm_p90 | context (ROI/target) | 0 | 0 | 1 |
| dist_center_to_ear_left_norm_periodicity | context (ROI/target) | 0 | 0 | 1 |
| dist_center_to_ear_left_norm_std | context (ROI/target) | 0 | 0 | 1 |
| dist_center_to_ear_left_norm_trend | context (ROI/target) | 0 | 0 | 1 |
| dist_center_to_ear_left_p10 | context (ROI/target) | 0 | 0 | 1 |
| dist_center_to_ear_left_p90 | context (ROI/target) | 0 | 0 | 1 |
| dist_center_to_ear_left_periodicity | context (ROI/target) | 0 | 0 | 1 |
| dist_center_to_ear_left_std | context (ROI/target) | 0 | 0 | 1 |
| dist_center_to_ear_left_trend | context (ROI/target) | 0 | 0 | 1 |
| dist_center_to_ear_right_delta | context (ROI/target) | 0 | 0 | 1 |
| dist_center_to_ear_right_energy | context (ROI/target) | 0 | 0 | 1 |
| dist_center_to_ear_right_max | context (ROI/target) | 0 | 0 | 1 |
| dist_center_to_ear_right_mean | context (ROI/target) | 0 | 0 | 1 |
| dist_center_to_ear_right_median | context (ROI/target) | 0 | 0 | 1 |
| dist_center_to_ear_right_norm_delta | context (ROI/target) | 0 | 0 | 1 |
| dist_center_to_ear_right_norm_energy | context (ROI/target) | 0 | 0 | 1 |
| dist_center_to_ear_right_norm_max | context (ROI/target) | 0 | 0 | 1 |
| dist_center_to_ear_right_norm_mean | context (ROI/target) | 0 | 0 | 1 |
| dist_center_to_ear_right_norm_median | context (ROI/target) | 0 | 0 | 1 |
| dist_center_to_ear_right_norm_p10 | context (ROI/target) | 0 | 0 | 1 |
| dist_center_to_ear_right_norm_p90 | context (ROI/target) | 0 | 0 | 1 |
| dist_center_to_ear_right_norm_periodicity | context (ROI/target) | 0 | 0 | 1 |
| dist_center_to_ear_right_norm_std | context (ROI/target) | 0 | 0 | 1 |
| dist_center_to_ear_right_norm_trend | context (ROI/target) | 0 | 0 | 1 |
| dist_center_to_ear_right_p10 | context (ROI/target) | 0 | 0 | 1 |
| dist_center_to_ear_right_p90 | context (ROI/target) | 0 | 0 | 1 |
| dist_center_to_ear_right_periodicity | context (ROI/target) | 0 | 0 | 1 |
| dist_center_to_ear_right_std | context (ROI/target) | 0 | 0 | 1 |
| dist_center_to_ear_right_trend | context (ROI/target) | 0 | 0 | 1 |

| | | | | |
|---|---|---|---|---|
| dist_center_to_lateral_left_delta | context (ROI/target) | 0 | 0 | 1 |
| dist_center_to_lateral_left_energy | context (ROI/target) | 0 | 0 | 1 |
| dist_center_to_lateral_left_max | context (ROI/target) | 0 | 0 | 1 |
| dist_center_to_lateral_left_mean | context (ROI/target) | 0 | 0 | 1 |
| dist_center_to_lateral_left_median | context (ROI/target) | 0 | 0 | 1 |
| dist_center_to_lateral_left_norm_delta | context (ROI/target) | 0 | 0 | 1 |
| dist_center_to_lateral_left_norm_energy | context (ROI/target) | 0 | 0 | 1 |
| dist_center_to_lateral_left_norm_max | context (ROI/target) | 0 | 0 | 1 |
| dist_center_to_lateral_left_norm_mean | context (ROI/target) | 0 | 0 | 1 |
| dist_center_to_lateral_left_norm_median | context (ROI/target) | 0 | 0 | 1 |
| dist_center_to_lateral_left_norm_p10 | context (ROI/target) | 0 | 0 | 1 |
| dist_center_to_lateral_left_norm_p90 | context (ROI/target) | 0 | 0 | 1 |
| dist_center_to_lateral_left_norm_periodicity | context (ROI/target) | 0 | 0 | 1 |
| dist_center_to_lateral_left_norm_std | context (ROI/target) | 0 | 0 | 1 |
| dist_center_to_lateral_left_norm_trend | context (ROI/target) | 0 | 0 | 1 |
| dist_center_to_lateral_left_p10 | context (ROI/target) | 0 | 0 | 1 |
| dist_center_to_lateral_left_p90 | context (ROI/target) | 0 | 0 | 1 |
| dist_center_to_lateral_left_periodicity | context (ROI/target) | 0 | 0 | 1 |
| dist_center_to_lateral_left_std | context (ROI/target) | 0 | 0 | 1 |
| dist_center_to_lateral_left_trend | context (ROI/target) | 0 | 0 | 1 |
| dist_center_to_lateral_right_delta | context (ROI/target) | 0 | 0 | 1 |
| dist_center_to_lateral_right_energy | context (ROI/target) | 0 | 0 | 1 |
| dist_center_to_lateral_right_max | context (ROI/target) | 0 | 0 | 1 |
| dist_center_to_lateral_right_mean | context (ROI/target) | 0 | 0 | 1 |
| dist_center_to_lateral_right_median | context (ROI/target) | 0 | 0 | 1 |
| dist_center_to_lateral_right_norm_delta | context (ROI/target) | 0 | 0 | 1 |
| dist_center_to_lateral_right_norm_energy | context (ROI/target) | 0 | 0 | 1 |
| dist_center_to_lateral_right_norm_max | context (ROI/target) | 0 | 0 | 1 |
| dist_center_to_lateral_right_norm_mean | context (ROI/target) | 0 | 0 | 1 |
| dist_center_to_lateral_right_norm_median | context (ROI/target) | 0 | 0 | 1 |
| dist_center_to_lateral_right_norm_p10 | context (ROI/target) | 0 | 0 | 1 |
| dist_center_to_lateral_right_norm_p90 | context (ROI/target) | 0 | 0 | 1 |
| dist_center_to_lateral_right_norm_periodicity | context (ROI/target) | 0 | 0 | 1 |
| dist_center_to_lateral_right_norm_std | context (ROI/target) | 0 | 0 | 1 |
| dist_center_to_lateral_right_norm_trend | context (ROI/target) | 0 | 0 | 1 |
| dist_center_to_lateral_right_p10 | context (ROI/target) | 0 | 0 | 1 |
| dist_center_to_lateral_right_p90 | context (ROI/target) | 0 | 0 | 1 |
| dist_center_to_lateral_right_periodicity | context (ROI/target) | 0 | 0 | 1 |
| dist_center_to_lateral_right_std | context (ROI/target) | 0 | 0 | 1 |
| dist_center_to_lateral_right_trend | context (ROI/target) | 0 | 0 | 1 |
| dist_center_to_nose_delta | context (ROI/target) | 0 | 0 | 1 |
| dist_center_to_nose_energy | context (ROI/target) | 0 | 0 | 1 |
| dist_center_to_nose_max | context (ROI/target) | 0 | 0 | 1 |
| dist_center_to_nose_mean | context (ROI/target) | 0 | 0 | 1 |

| | | | | |
|---|---|---|---|---|
| dist_center_to_nose_median | context (ROI/target) | 0 | 0 | 1 |
| dist_center_to_nose_norm_delta | context (ROI/target) | 0 | 0 | 1 |
| dist_center_to_nose_norm_energy | context (ROI/target) | 0 | 0 | 1 |
| dist_center_to_nose_norm_max | context (ROI/target) | 0 | 0 | 1 |
| dist_center_to_nose_norm_mean | context (ROI/target) | 0 | 0 | 1 |
| dist_center_to_nose_norm_median | context (ROI/target) | 0 | 0 | 1 |
| dist_center_to_nose_norm_p10 | context (ROI/target) | 0 | 0 | 1 |
| dist_center_to_nose_norm_p90 | context (ROI/target) | 0 | 0 | 1 |
| dist_center_to_nose_norm_periodicity | context (ROI/target) | 0 | 0 | 1 |
| dist_center_to_nose_norm_std | context (ROI/target) | 0 | 0 | 1 |
| dist_center_to_nose_norm_trend | context (ROI/target) | 0 | 0 | 1 |
| dist_center_to_nose_p10 | context (ROI/target) | 0 | 0 | 1 |
| dist_center_to_nose_p90 | context (ROI/target) | 0 | 0 | 1 |
| dist_center_to_nose_periodicity | context (ROI/target) | 0 | 0 | 1 |
| dist_center_to_nose_std | context (ROI/target) | 0 | 0 | 1 |
| dist_center_to_nose_trend | context (ROI/target) | 0 | 0 | 1 |
| dist_center_to_tail_base_delta | context (ROI/target) | 0 | 0 | 1 |
| dist_center_to_tail_base_energy | context (ROI/target) | 0 | 0 | 1 |
| dist_center_to_tail_base_max | context (ROI/target) | 0 | 0 | 1 |
| dist_center_to_tail_base_mean | context (ROI/target) | 0 | 0 | 1 |
| dist_center_to_tail_base_median | context (ROI/target) | 0 | 0 | 1 |
| dist_center_to_tail_base_norm_delta | context (ROI/target) | 0 | 0 | 1 |
| dist_center_to_tail_base_norm_energy | context (ROI/target) | 0 | 0 | 1 |
| dist_center_to_tail_base_norm_max | context (ROI/target) | 0 | 0 | 1 |
| dist_center_to_tail_base_norm_mean | context (ROI/target) | 0 | 0 | 1 |
| dist_center_to_tail_base_norm_median | context (ROI/target) | 0 | 0 | 1 |
| dist_center_to_tail_base_norm_p10 | context (ROI/target) | 0 | 0 | 1 |
| dist_center_to_tail_base_norm_p90 | context (ROI/target) | 0 | 0 | 1 |
| dist_center_to_tail_base_norm_periodicity | context (ROI/target) | 0 | 0 | 1 |
| dist_center_to_tail_base_norm_std | context (ROI/target) | 0 | 0 | 1 |
| dist_center_to_tail_base_norm_trend | context (ROI/target) | 0 | 0 | 1 |
| dist_center_to_tail_base_p10 | context (ROI/target) | 0 | 0 | 1 |
| dist_center_to_tail_base_p90 | context (ROI/target) | 0 | 0 | 1 |
| dist_center_to_tail_base_periodicity | context (ROI/target) | 0 | 0 | 1 |
| dist_center_to_tail_base_std | context (ROI/target) | 0 | 0 | 1 |
| dist_center_to_tail_base_trend | context (ROI/target) | 0 | 0 | 1 |
| dist_ear_left_to_ear_right_delta | kinematics | 0 | 1 | 1 |
| dist_ear_left_to_ear_right_energy | kinematics | 0 | 1 | 1 |
| dist_ear_left_to_ear_right_max | pose geometry | 0 | 1 | 1 |
| dist_ear_left_to_ear_right_mean | pose geometry | 0 | 1 | 1 |
| dist_ear_left_to_ear_right_median | pose geometry | 0 | 1 | 1 |
| dist_ear_left_to_ear_right_norm_delta | kinematics | 0 | 1 | 1 |
| dist_ear_left_to_ear_right_norm_energy | kinematics | 0 | 1 | 1 |
| dist_ear_left_to_ear_right_norm_max | pose geometry | 0 | 1 | 1 |

| | | | | |
|---|---|---|---|---|
| dist_ear_left_to_ear_right_norm_mean | pose geometry | 0 | 1 | 1 |
| dist_ear_left_to_ear_right_norm_median | pose geometry | 0 | 1 | 1 |
| dist_ear_left_to_ear_right_norm_p10 | pose geometry | 0 | 1 | 1 |
| dist_ear_left_to_ear_right_norm_p90 | pose geometry | 0 | 1 | 1 |
| dist_ear_left_to_ear_right_norm_periodicity | pose geometry | 0 | 1 | 1 |
| dist_ear_left_to_ear_right_norm_std | kinematics | 0 | 1 | 1 |
| dist_ear_left_to_ear_right_norm_trend | pose geometry | 0 | 1 | 1 |
| dist_ear_left_to_ear_right_p10 | pose geometry | 0 | 1 | 1 |
| dist_ear_left_to_ear_right_p90 | pose geometry | 0 | 1 | 1 |
| dist_ear_left_to_ear_right_periodicity | pose geometry | 0 | 1 | 1 |
| dist_ear_left_to_ear_right_std | kinematics | 0 | 1 | 1 |
| dist_ear_left_to_ear_right_trend | pose geometry | 0 | 1 | 1 |
| dist_ear_left_to_lateral_left_delta | kinematics | 0 | 0 | 1 |
| dist_ear_left_to_lateral_left_energy | kinematics | 0 | 0 | 1 |
| dist_ear_left_to_lateral_left_max | pose geometry | 0 | 0 | 1 |
| dist_ear_left_to_lateral_left_mean | pose geometry | 0 | 0 | 1 |
| dist_ear_left_to_lateral_left_median | pose geometry | 0 | 0 | 1 |
| dist_ear_left_to_lateral_left_norm_delta | kinematics | 0 | 0 | 1 |
| dist_ear_left_to_lateral_left_norm_energy | kinematics | 0 | 0 | 1 |
| dist_ear_left_to_lateral_left_norm_max | pose geometry | 0 | 0 | 1 |
| dist_ear_left_to_lateral_left_norm_mean | pose geometry | 0 | 0 | 1 |
| dist_ear_left_to_lateral_left_norm_median | pose geometry | 0 | 0 | 1 |
| dist_ear_left_to_lateral_left_norm_p10 | pose geometry | 0 | 0 | 1 |
| dist_ear_left_to_lateral_left_norm_p90 | pose geometry | 0 | 0 | 1 |
| dist_ear_left_to_lateral_left_norm_periodicity | pose geometry | 0 | 0 | 1 |
| dist_ear_left_to_lateral_left_norm_std | kinematics | 0 | 0 | 1 |
| dist_ear_left_to_lateral_left_norm_trend | pose geometry | 0 | 0 | 1 |
| dist_ear_left_to_lateral_left_p10 | pose geometry | 0 | 0 | 1 |
| dist_ear_left_to_lateral_left_p90 | pose geometry | 0 | 0 | 1 |
| dist_ear_left_to_lateral_left_periodicity | pose geometry | 0 | 0 | 1 |
| dist_ear_left_to_lateral_left_std | kinematics | 0 | 0 | 1 |
| dist_ear_left_to_lateral_left_trend | pose geometry | 0 | 0 | 1 |
| dist_ear_left_to_lateral_right_delta | kinematics | 0 | 0 | 1 |
| dist_ear_left_to_lateral_right_energy | kinematics | 0 | 0 | 1 |
| dist_ear_left_to_lateral_right_max | pose geometry | 0 | 0 | 1 |
| dist_ear_left_to_lateral_right_mean | pose geometry | 0 | 0 | 1 |
| dist_ear_left_to_lateral_right_median | pose geometry | 0 | 0 | 1 |
| dist_ear_left_to_lateral_right_norm_delta | kinematics | 0 | 0 | 1 |
| dist_ear_left_to_lateral_right_norm_energy | kinematics | 0 | 0 | 1 |
| dist_ear_left_to_lateral_right_norm_max | pose geometry | 0 | 0 | 1 |
| dist_ear_left_to_lateral_right_norm_mean | pose geometry | 0 | 0 | 1 |
| dist_ear_left_to_lateral_right_norm_median | pose geometry | 0 | 0 | 1 |
| dist_ear_left_to_lateral_right_norm_p10 | pose geometry | 0 | 0 | 1 |
| dist_ear_left_to_lateral_right_norm_p90 | pose geometry | 0 | 0 | 1 |

| | | | | |
|---|---|---|---|---|
| dist_ear_left_to_lateral_right_norm_periodicity | pose geometry | 0 | 0 | 1 |
| dist_ear_left_to_lateral_right_norm_std | kinematics | 0 | 0 | 1 |
| dist_ear_left_to_lateral_right_norm_trend | pose geometry | 0 | 0 | 1 |
| dist_ear_left_to_lateral_right_p10 | pose geometry | 0 | 0 | 1 |
| dist_ear_left_to_lateral_right_p90 | pose geometry | 0 | 0 | 1 |
| dist_ear_left_to_lateral_right_periodicity | pose geometry | 0 | 0 | 1 |
| dist_ear_left_to_lateral_right_std | kinematics | 0 | 0 | 1 |
| dist_ear_left_to_lateral_right_trend | pose geometry | 0 | 0 | 1 |
| dist_ear_left_to_nose_delta | kinematics | 0 | 1 | 1 |
| dist_ear_left_to_nose_energy | kinematics | 0 | 1 | 1 |
| dist_ear_left_to_nose_max | pose geometry | 0 | 1 | 1 |
| dist_ear_left_to_nose_mean | pose geometry | 0 | 1 | 1 |
| dist_ear_left_to_nose_median | pose geometry | 0 | 1 | 1 |
| dist_ear_left_to_nose_norm_delta | kinematics | 0 | 1 | 1 |
| dist_ear_left_to_nose_norm_energy | kinematics | 0 | 1 | 1 |
| dist_ear_left_to_nose_norm_max | pose geometry | 0 | 1 | 1 |
| dist_ear_left_to_nose_norm_mean | pose geometry | 0 | 1 | 1 |
| dist_ear_left_to_nose_norm_median | pose geometry | 0 | 1 | 1 |
| dist_ear_left_to_nose_norm_p10 | pose geometry | 0 | 1 | 1 |
| dist_ear_left_to_nose_norm_p90 | pose geometry | 0 | 1 | 1 |
| dist_ear_left_to_nose_norm_periodicity | pose geometry | 0 | 1 | 1 |
| dist_ear_left_to_nose_norm_std | kinematics | 0 | 1 | 1 |
| dist_ear_left_to_nose_norm_trend | pose geometry | 0 | 1 | 1 |
| dist_ear_left_to_nose_p10 | pose geometry | 0 | 1 | 1 |
| dist_ear_left_to_nose_p90 | pose geometry | 0 | 1 | 1 |
| dist_ear_left_to_nose_periodicity | pose geometry | 0 | 1 | 1 |
| dist_ear_left_to_nose_std | kinematics | 0 | 1 | 1 |
| dist_ear_left_to_nose_trend | pose geometry | 0 | 1 | 1 |
| dist_ear_left_to_tail_base_delta | kinematics | 0 | 1 | 1 |
| dist_ear_left_to_tail_base_energy | kinematics | 0 | 1 | 1 |
| dist_ear_left_to_tail_base_max | pose geometry | 0 | 1 | 1 |
| dist_ear_left_to_tail_base_mean | pose geometry | 0 | 1 | 1 |
| dist_ear_left_to_tail_base_median | pose geometry | 0 | 1 | 1 |
| dist_ear_left_to_tail_base_norm_delta | kinematics | 0 | 1 | 1 |
| dist_ear_left_to_tail_base_norm_energy | kinematics | 0 | 1 | 1 |
| dist_ear_left_to_tail_base_norm_max | pose geometry | 0 | 1 | 1 |
| dist_ear_left_to_tail_base_norm_mean | pose geometry | 0 | 1 | 1 |
| dist_ear_left_to_tail_base_norm_median | pose geometry | 0 | 1 | 1 |
| dist_ear_left_to_tail_base_norm_p10 | pose geometry | 0 | 1 | 1 |
| dist_ear_left_to_tail_base_norm_p90 | pose geometry | 0 | 1 | 1 |
| dist_ear_left_to_tail_base_norm_periodicity | pose geometry | 0 | 1 | 1 |
| dist_ear_left_to_tail_base_norm_std | kinematics | 0 | 1 | 1 |
| dist_ear_left_to_tail_base_norm_trend | pose geometry | 0 | 1 | 1 |
| dist_ear_left_to_tail_base_p10 | pose geometry | 0 | 1 | 1 |

| | | | | |
|---|---|---|---|---|
| dist_ear_left_to_tail_base_p90 | pose geometry | 0 | 1 | 1 |
| dist_ear_left_to_tail_base_periodicity | pose geometry | 0 | 1 | 1 |
| dist_ear_left_to_tail_base_std | kinematics | 0 | 1 | 1 |
| dist_ear_left_to_tail_base_trend | pose geometry | 0 | 1 | 1 |
| dist_ear_right_to_lateral_left_delta | kinematics | 0 | 0 | 1 |
| dist_ear_right_to_lateral_left_energy | kinematics | 0 | 0 | 1 |
| dist_ear_right_to_lateral_left_max | pose geometry | 0 | 0 | 1 |
| dist_ear_right_to_lateral_left_mean | pose geometry | 0 | 0 | 1 |
| dist_ear_right_to_lateral_left_median | pose geometry | 0 | 0 | 1 |
| dist_ear_right_to_lateral_left_norm_delta | kinematics | 0 | 0 | 1 |
| dist_ear_right_to_lateral_left_norm_energy | kinematics | 0 | 0 | 1 |
| dist_ear_right_to_lateral_left_norm_max | pose geometry | 0 | 0 | 1 |
| dist_ear_right_to_lateral_left_norm_mean | pose geometry | 0 | 0 | 1 |
| dist_ear_right_to_lateral_left_norm_median | pose geometry | 0 | 0 | 1 |
| dist_ear_right_to_lateral_left_norm_p10 | pose geometry | 0 | 0 | 1 |
| dist_ear_right_to_lateral_left_norm_p90 | pose geometry | 0 | 0 | 1 |
| dist_ear_right_to_lateral_left_norm_periodicity | pose geometry | 0 | 0 | 1 |
| dist_ear_right_to_lateral_left_norm_std | kinematics | 0 | 0 | 1 |
| dist_ear_right_to_lateral_left_norm_trend | pose geometry | 0 | 0 | 1 |
| dist_ear_right_to_lateral_left_p10 | pose geometry | 0 | 0 | 1 |
| dist_ear_right_to_lateral_left_p90 | pose geometry | 0 | 0 | 1 |
| dist_ear_right_to_lateral_left_periodicity | pose geometry | 0 | 0 | 1 |
| dist_ear_right_to_lateral_left_std | kinematics | 0 | 0 | 1 |
| dist_ear_right_to_lateral_left_trend | pose geometry | 0 | 0 | 1 |
| dist_ear_right_to_lateral_right_delta | kinematics | 0 | 0 | 1 |
| dist_ear_right_to_lateral_right_energy | kinematics | 0 | 0 | 1 |
| dist_ear_right_to_lateral_right_max | pose geometry | 0 | 0 | 1 |
| dist_ear_right_to_lateral_right_mean | pose geometry | 0 | 0 | 1 |
| dist_ear_right_to_lateral_right_median | pose geometry | 0 | 0 | 1 |
| dist_ear_right_to_lateral_right_norm_delta | kinematics | 0 | 0 | 1 |
| dist_ear_right_to_lateral_right_norm_energy | kinematics | 0 | 0 | 1 |
| dist_ear_right_to_lateral_right_norm_max | pose geometry | 0 | 0 | 1 |
| dist_ear_right_to_lateral_right_norm_mean | pose geometry | 0 | 0 | 1 |
| dist_ear_right_to_lateral_right_norm_median | pose geometry | 0 | 0 | 1 |
| dist_ear_right_to_lateral_right_norm_p10 | pose geometry | 0 | 0 | 1 |
| dist_ear_right_to_lateral_right_norm_p90 | pose geometry | 0 | 0 | 1 |
| dist_ear_right_to_lateral_right_norm_periodicity | pose geometry | 0 | 0 | 1 |
| dist_ear_right_to_lateral_right_norm_std | kinematics | 0 | 0 | 1 |
| dist_ear_right_to_lateral_right_norm_trend | pose geometry | 0 | 0 | 1 |
| dist_ear_right_to_lateral_right_p10 | pose geometry | 0 | 0 | 1 |
| dist_ear_right_to_lateral_right_p90 | pose geometry | 0 | 0 | 1 |
| dist_ear_right_to_lateral_right_periodicity | pose geometry | 0 | 0 | 1 |
| dist_ear_right_to_lateral_right_std | kinematics | 0 | 0 | 1 |
| dist_ear_right_to_lateral_right_trend | pose geometry | 0 | 0 | 1 |

| | | | | |
|---|---|---|---|---|
| dist_ear_right_to_nose_delta | kinematics | 0 | 1 | 1 |
| dist_ear_right_to_nose_energy | kinematics | 0 | 1 | 1 |
| dist_ear_right_to_nose_max | pose geometry | 0 | 1 | 1 |
| dist_ear_right_to_nose_mean | pose geometry | 0 | 1 | 1 |
| dist_ear_right_to_nose_median | pose geometry | 0 | 1 | 1 |
| dist_ear_right_to_nose_norm_delta | kinematics | 0 | 1 | 1 |
| dist_ear_right_to_nose_norm_energy | kinematics | 0 | 1 | 1 |
| dist_ear_right_to_nose_norm_max | pose geometry | 0 | 1 | 1 |
| dist_ear_right_to_nose_norm_mean | pose geometry | 0 | 1 | 1 |
| dist_ear_right_to_nose_norm_median | pose geometry | 0 | 1 | 1 |
| dist_ear_right_to_nose_norm_p10 | pose geometry | 0 | 1 | 1 |
| dist_ear_right_to_nose_norm_p90 | pose geometry | 0 | 1 | 1 |
| dist_ear_right_to_nose_norm_periodicity | pose geometry | 0 | 1 | 1 |
| dist_ear_right_to_nose_norm_std | kinematics | 0 | 1 | 1 |
| dist_ear_right_to_nose_norm_trend | pose geometry | 0 | 1 | 1 |
| dist_ear_right_to_nose_p10 | pose geometry | 0 | 1 | 1 |
| dist_ear_right_to_nose_p90 | pose geometry | 0 | 1 | 1 |
| dist_ear_right_to_nose_periodicity | pose geometry | 0 | 1 | 1 |
| dist_ear_right_to_nose_std | kinematics | 0 | 1 | 1 |
| dist_ear_right_to_nose_trend | pose geometry | 0 | 1 | 1 |
| dist_ear_right_to_tail_base_delta | kinematics | 0 | 1 | 1 |
| dist_ear_right_to_tail_base_energy | kinematics | 0 | 1 | 1 |
| dist_ear_right_to_tail_base_max | pose geometry | 0 | 1 | 1 |
| dist_ear_right_to_tail_base_mean | pose geometry | 0 | 1 | 1 |
| dist_ear_right_to_tail_base_median | pose geometry | 0 | 1 | 1 |
| dist_ear_right_to_tail_base_norm_delta | kinematics | 0 | 1 | 1 |
| dist_ear_right_to_tail_base_norm_energy | kinematics | 0 | 1 | 1 |
| dist_ear_right_to_tail_base_norm_max | pose geometry | 0 | 1 | 1 |
| dist_ear_right_to_tail_base_norm_mean | pose geometry | 0 | 1 | 1 |
| dist_ear_right_to_tail_base_norm_median | pose geometry | 0 | 1 | 1 |
| dist_ear_right_to_tail_base_norm_p10 | pose geometry | 0 | 1 | 1 |
| dist_ear_right_to_tail_base_norm_p90 | pose geometry | 0 | 1 | 1 |
| dist_ear_right_to_tail_base_norm_periodicity | pose geometry | 0 | 1 | 1 |
| dist_ear_right_to_tail_base_norm_std | kinematics | 0 | 1 | 1 |
| dist_ear_right_to_tail_base_norm_trend | pose geometry | 0 | 1 | 1 |
| dist_ear_right_to_tail_base_p10 | pose geometry | 0 | 1 | 1 |
| dist_ear_right_to_tail_base_p90 | pose geometry | 0 | 1 | 1 |
| dist_ear_right_to_tail_base_periodicity | pose geometry | 0 | 1 | 1 |
| dist_ear_right_to_tail_base_std | kinematics | 0 | 1 | 1 |
| dist_ear_right_to_tail_base_trend | pose geometry | 0 | 1 | 1 |
| dist_front_left_paw_to_front_right_paw_delta | kinematics | 0 | 0 | 0 |
| dist_front_left_paw_to_front_right_paw_energy | kinematics | 0 | 0 | 0 |
| dist_front_left_paw_to_front_right_paw_max | pose geometry | 0 | 0 | 0 |
| dist_front_left_paw_to_front_right_paw_mean | pose geometry | 0 | 0 | 0 |

| | | | | |
|---|---|---|---|---|
| dist_front_left_paw_to_front_right_paw_median | pose geometry | 0 | 0 | 0 |
| dist_front_left_paw_to_front_right_paw_norm_delta | kinematics | 0 | 0 | 0 |
| dist_front_left_paw_to_front_right_paw_norm_energy | kinematics | 0 | 0 | 0 |
| dist_front_left_paw_to_front_right_paw_norm_max | pose geometry | 0 | 0 | 0 |
| dist_front_left_paw_to_front_right_paw_norm_mean | pose geometry | 0 | 0 | 0 |
| dist_front_left_paw_to_front_right_paw_norm_median | pose geometry | 0 | 0 | 0 |
| dist_front_left_paw_to_front_right_paw_norm_p10 | pose geometry | 0 | 0 | 0 |
| dist_front_left_paw_to_front_right_paw_norm_p90 | pose geometry | 0 | 0 | 0 |
| dist_front_left_paw_to_front_right_paw_norm_periodic | pose geometry | 0 | 0 | 0 |
| dist_front_left_paw_to_front_right_paw_norm_std | kinematics | 0 | 0 | 0 |
| dist_front_left_paw_to_front_right_paw_norm_trend | pose geometry | 0 | 0 | 0 |
| dist_front_left_paw_to_front_right_paw_p10 | pose geometry | 0 | 0 | 0 |
| dist_front_left_paw_to_front_right_paw_p90 | pose geometry | 0 | 0 | 0 |
| dist_front_left_paw_to_front_right_paw_periodicity | pose geometry | 0 | 0 | 0 |
| dist_front_left_paw_to_front_right_paw_std | kinematics | 0 | 0 | 0 |
| dist_front_left_paw_to_front_right_paw_trend | pose geometry | 0 | 0 | 0 |
| dist_front_left_paw_to_left_ear_delta | kinematics | 0 | 0 | 0 |
| dist_front_left_paw_to_left_ear_energy | kinematics | 0 | 0 | 0 |
| dist_front_left_paw_to_left_ear_max | pose geometry | 0 | 0 | 0 |
| dist_front_left_paw_to_left_ear_mean | pose geometry | 0 | 0 | 0 |
| dist_front_left_paw_to_left_ear_median | pose geometry | 0 | 0 | 0 |
| dist_front_left_paw_to_left_ear_norm_delta | kinematics | 0 | 0 | 0 |
| dist_front_left_paw_to_left_ear_norm_energy | kinematics | 0 | 0 | 0 |
| dist_front_left_paw_to_left_ear_norm_max | pose geometry | 0 | 0 | 0 |
| dist_front_left_paw_to_left_ear_norm_mean | pose geometry | 0 | 0 | 0 |
| dist_front_left_paw_to_left_ear_norm_median | pose geometry | 0 | 0 | 0 |
| dist_front_left_paw_to_left_ear_norm_p10 | pose geometry | 0 | 0 | 0 |
| dist_front_left_paw_to_left_ear_norm_p90 | pose geometry | 0 | 0 | 0 |
| dist_front_left_paw_to_left_ear_norm_periodicity | pose geometry | 0 | 0 | 0 |
| dist_front_left_paw_to_left_ear_norm_std | kinematics | 0 | 0 | 0 |
| dist_front_left_paw_to_left_ear_norm_trend | pose geometry | 0 | 0 | 0 |
| dist_front_left_paw_to_left_ear_p10 | pose geometry | 0 | 0 | 0 |
| dist_front_left_paw_to_left_ear_p90 | pose geometry | 0 | 0 | 0 |
| dist_front_left_paw_to_left_ear_periodicity | pose geometry | 0 | 0 | 0 |
| dist_front_left_paw_to_left_ear_std | kinematics | 0 | 0 | 0 |
| dist_front_left_paw_to_left_ear_trend | pose geometry | 0 | 0 | 0 |
| dist_front_left_paw_to_mid_back_delta | kinematics | 0 | 0 | 0 |
| dist_front_left_paw_to_mid_back_energy | kinematics | 0 | 0 | 0 |
| dist_front_left_paw_to_mid_back_max | pose geometry | 0 | 0 | 0 |
| dist_front_left_paw_to_mid_back_mean | pose geometry | 0 | 0 | 0 |
| dist_front_left_paw_to_mid_back_median | pose geometry | 0 | 0 | 0 |
| dist_front_left_paw_to_mid_back_norm_delta | kinematics | 0 | 0 | 0 |
| dist_front_left_paw_to_mid_back_norm_energy | kinematics | 0 | 0 | 0 |
| dist_front_left_paw_to_mid_back_norm_max | pose geometry | 0 | 0 | 0 |

| | | | | |
|---|---|---|---|---|
| dist_front_left_paw_to_mid_back_norm_mean | pose geometry | 0 | 0 | 0 |
| dist_front_left_paw_to_mid_back_norm_median | pose geometry | 0 | 0 | 0 |
| dist_front_left_paw_to_mid_back_norm_p10 | pose geometry | 0 | 0 | 0 |
| dist_front_left_paw_to_mid_back_norm_p90 | pose geometry | 0 | 0 | 0 |
| dist_front_left_paw_to_mid_back_norm_periodicity | pose geometry | 0 | 0 | 0 |
| dist_front_left_paw_to_mid_back_norm_std | kinematics | 0 | 0 | 0 |
| dist_front_left_paw_to_mid_back_norm_trend | pose geometry | 0 | 0 | 0 |
| dist_front_left_paw_to_mid_back_p10 | pose geometry | 0 | 0 | 0 |
| dist_front_left_paw_to_mid_back_p90 | pose geometry | 0 | 0 | 0 |
| dist_front_left_paw_to_mid_back_periodicity | pose geometry | 0 | 0 | 0 |
| dist_front_left_paw_to_mid_back_std | kinematics | 0 | 0 | 0 |
| dist_front_left_paw_to_mid_back_trend | pose geometry | 0 | 0 | 0 |
| dist_front_left_paw_to_nose_delta | kinematics | 0 | 0 | 0 |
| dist_front_left_paw_to_nose_energy | kinematics | 0 | 0 | 0 |
| dist_front_left_paw_to_nose_max | pose geometry | 0 | 0 | 0 |
| dist_front_left_paw_to_nose_mean | pose geometry | 0 | 0 | 0 |
| dist_front_left_paw_to_nose_median | pose geometry | 0 | 0 | 0 |
| dist_front_left_paw_to_nose_norm_delta | kinematics | 0 | 0 | 0 |
| dist_front_left_paw_to_nose_norm_energy | kinematics | 0 | 0 | 0 |
| dist_front_left_paw_to_nose_norm_max | pose geometry | 0 | 0 | 0 |
| dist_front_left_paw_to_nose_norm_mean | pose geometry | 0 | 0 | 0 |
| dist_front_left_paw_to_nose_norm_median | pose geometry | 0 | 0 | 0 |
| dist_front_left_paw_to_nose_norm_p10 | pose geometry | 0 | 0 | 0 |
| dist_front_left_paw_to_nose_norm_p90 | pose geometry | 0 | 0 | 0 |
| dist_front_left_paw_to_nose_norm_periodicity | pose geometry | 0 | 0 | 0 |
| dist_front_left_paw_to_nose_norm_std | kinematics | 0 | 0 | 0 |
| dist_front_left_paw_to_nose_norm_trend | pose geometry | 0 | 0 | 0 |
| dist_front_left_paw_to_nose_p10 | pose geometry | 0 | 0 | 0 |
| dist_front_left_paw_to_nose_p90 | pose geometry | 0 | 0 | 0 |
| dist_front_left_paw_to_nose_periodicity | pose geometry | 0 | 0 | 0 |
| dist_front_left_paw_to_nose_std | kinematics | 0 | 0 | 0 |
| dist_front_left_paw_to_nose_trend | pose geometry | 0 | 0 | 0 |
| dist_front_left_paw_to_right_ear_delta | kinematics | 0 | 0 | 0 |
| dist_front_left_paw_to_right_ear_energy | kinematics | 0 | 0 | 0 |
| dist_front_left_paw_to_right_ear_max | pose geometry | 0 | 0 | 0 |
| dist_front_left_paw_to_right_ear_mean | pose geometry | 0 | 0 | 0 |
| dist_front_left_paw_to_right_ear_median | pose geometry | 0 | 0 | 0 |
| dist_front_left_paw_to_right_ear_norm_delta | kinematics | 0 | 0 | 0 |
| dist_front_left_paw_to_right_ear_norm_energy | kinematics | 0 | 0 | 0 |
| dist_front_left_paw_to_right_ear_norm_max | pose geometry | 0 | 0 | 0 |
| dist_front_left_paw_to_right_ear_norm_mean | pose geometry | 0 | 0 | 0 |
| dist_front_left_paw_to_right_ear_norm_median | pose geometry | 0 | 0 | 0 |
| dist_front_left_paw_to_right_ear_norm_p10 | pose geometry | 0 | 0 | 0 |
| dist_front_left_paw_to_right_ear_norm_p90 | pose geometry | 0 | 0 | 0 |

| | | | | |
|---|---|---|---|---|
| dist_front_left_paw_to_right_ear_norm_periodicity | pose geometry | 0 | 0 | 0 |
| dist_front_left_paw_to_right_ear_norm_std | kinematics | 0 | 0 | 0 |
| dist_front_left_paw_to_right_ear_norm_trend | pose geometry | 0 | 0 | 0 |
| dist_front_left_paw_to_right_ear_p10 | pose geometry | 0 | 0 | 0 |
| dist_front_left_paw_to_right_ear_p90 | pose geometry | 0 | 0 | 0 |
| dist_front_left_paw_to_right_ear_periodicity | pose geometry | 0 | 0 | 0 |
| dist_front_left_paw_to_right_ear_std | kinematics | 0 | 0 | 0 |
| dist_front_left_paw_to_right_ear_trend | pose geometry | 0 | 0 | 0 |
| dist_front_left_paw_to_tail_base_delta | kinematics | 0 | 0 | 0 |
| dist_front_left_paw_to_tail_base_energy | kinematics | 0 | 0 | 0 |
| dist_front_left_paw_to_tail_base_max | pose geometry | 0 | 0 | 0 |
| dist_front_left_paw_to_tail_base_mean | pose geometry | 0 | 0 | 0 |
| dist_front_left_paw_to_tail_base_median | pose geometry | 0 | 0 | 0 |
| dist_front_left_paw_to_tail_base_norm_delta | kinematics | 0 | 0 | 0 |
| dist_front_left_paw_to_tail_base_norm_energy | kinematics | 0 | 0 | 0 |
| dist_front_left_paw_to_tail_base_norm_max | pose geometry | 0 | 0 | 0 |
| dist_front_left_paw_to_tail_base_norm_mean | pose geometry | 0 | 0 | 0 |
| dist_front_left_paw_to_tail_base_norm_median | pose geometry | 0 | 0 | 0 |
| dist_front_left_paw_to_tail_base_norm_p10 | pose geometry | 0 | 0 | 0 |
| dist_front_left_paw_to_tail_base_norm_p90 | pose geometry | 0 | 0 | 0 |
| dist_front_left_paw_to_tail_base_norm_periodicity | pose geometry | 0 | 0 | 0 |
| dist_front_left_paw_to_tail_base_norm_std | kinematics | 0 | 0 | 0 |
| dist_front_left_paw_to_tail_base_norm_trend | pose geometry | 0 | 0 | 0 |
| dist_front_left_paw_to_tail_base_p10 | pose geometry | 0 | 0 | 0 |
| dist_front_left_paw_to_tail_base_p90 | pose geometry | 0 | 0 | 0 |
| dist_front_left_paw_to_tail_base_periodicity | pose geometry | 0 | 0 | 0 |
| dist_front_left_paw_to_tail_base_std | kinematics | 0 | 0 | 0 |
| dist_front_left_paw_to_tail_base_trend | pose geometry | 0 | 0 | 0 |
| dist_front_right_paw_to_left_ear_delta | kinematics | 0 | 0 | 0 |
| dist_front_right_paw_to_left_ear_energy | kinematics | 0 | 0 | 0 |
| dist_front_right_paw_to_left_ear_max | pose geometry | 0 | 0 | 0 |
| dist_front_right_paw_to_left_ear_mean | pose geometry | 0 | 0 | 0 |
| dist_front_right_paw_to_left_ear_median | pose geometry | 0 | 0 | 0 |
| dist_front_right_paw_to_left_ear_norm_delta | kinematics | 0 | 0 | 0 |
| dist_front_right_paw_to_left_ear_norm_energy | kinematics | 0 | 0 | 0 |
| dist_front_right_paw_to_left_ear_norm_max | pose geometry | 0 | 0 | 0 |
| dist_front_right_paw_to_left_ear_norm_mean | pose geometry | 0 | 0 | 0 |
| dist_front_right_paw_to_left_ear_norm_median | pose geometry | 0 | 0 | 0 |
| dist_front_right_paw_to_left_ear_norm_p10 | pose geometry | 0 | 0 | 0 |
| dist_front_right_paw_to_left_ear_norm_p90 | pose geometry | 0 | 0 | 0 |
| dist_front_right_paw_to_left_ear_norm_periodicity | pose geometry | 0 | 0 | 0 |
| dist_front_right_paw_to_left_ear_norm_std | kinematics | 0 | 0 | 0 |
| dist_front_right_paw_to_left_ear_norm_trend | pose geometry | 0 | 0 | 0 |
| dist_front_right_paw_to_left_ear_p10 | pose geometry | 0 | 0 | 0 |

| | | | | |
|---|---|---|---|---|
| dist_front_right_paw_to_left_ear_p90 | pose geometry | 0 | 0 | 0 |
| dist_front_right_paw_to_left_ear_periodicity | pose geometry | 0 | 0 | 0 |
| dist_front_right_paw_to_left_ear_std | kinematics | 0 | 0 | 0 |
| dist_front_right_paw_to_left_ear_trend | pose geometry | 0 | 0 | 0 |
| dist_front_right_paw_to_mid_back_delta | kinematics | 0 | 0 | 0 |
| dist_front_right_paw_to_mid_back_energy | kinematics | 0 | 0 | 0 |
| dist_front_right_paw_to_mid_back_max | pose geometry | 0 | 0 | 0 |
| dist_front_right_paw_to_mid_back_mean | pose geometry | 0 | 0 | 0 |
| dist_front_right_paw_to_mid_back_median | pose geometry | 0 | 0 | 0 |
| dist_front_right_paw_to_mid_back_norm_delta | kinematics | 0 | 0 | 0 |
| dist_front_right_paw_to_mid_back_norm_energy | kinematics | 0 | 0 | 0 |
| dist_front_right_paw_to_mid_back_norm_max | pose geometry | 0 | 0 | 0 |
| dist_front_right_paw_to_mid_back_norm_mean | pose geometry | 0 | 0 | 0 |
| dist_front_right_paw_to_mid_back_norm_median | pose geometry | 0 | 0 | 0 |
| dist_front_right_paw_to_mid_back_norm_p10 | pose geometry | 0 | 0 | 0 |
| dist_front_right_paw_to_mid_back_norm_p90 | pose geometry | 0 | 0 | 0 |
| dist_front_right_paw_to_mid_back_norm_periodicity | pose geometry | 0 | 0 | 0 |
| dist_front_right_paw_to_mid_back_norm_std | kinematics | 0 | 0 | 0 |
| dist_front_right_paw_to_mid_back_norm_trend | pose geometry | 0 | 0 | 0 |
| dist_front_right_paw_to_mid_back_p10 | pose geometry | 0 | 0 | 0 |
| dist_front_right_paw_to_mid_back_p90 | pose geometry | 0 | 0 | 0 |
| dist_front_right_paw_to_mid_back_periodicity | pose geometry | 0 | 0 | 0 |
| dist_front_right_paw_to_mid_back_std | kinematics | 0 | 0 | 0 |
| dist_front_right_paw_to_mid_back_trend | pose geometry | 0 | 0 | 0 |
| dist_front_right_paw_to_nose_delta | kinematics | 0 | 0 | 0 |
| dist_front_right_paw_to_nose_energy | kinematics | 0 | 0 | 0 |
| dist_front_right_paw_to_nose_max | pose geometry | 0 | 0 | 0 |
| dist_front_right_paw_to_nose_mean | pose geometry | 0 | 0 | 0 |
| dist_front_right_paw_to_nose_median | pose geometry | 0 | 0 | 0 |
| dist_front_right_paw_to_nose_norm_delta | kinematics | 0 | 0 | 0 |
| dist_front_right_paw_to_nose_norm_energy | kinematics | 0 | 0 | 0 |
| dist_front_right_paw_to_nose_norm_max | pose geometry | 0 | 0 | 0 |
| dist_front_right_paw_to_nose_norm_mean | pose geometry | 0 | 0 | 0 |
| dist_front_right_paw_to_nose_norm_median | pose geometry | 0 | 0 | 0 |
| dist_front_right_paw_to_nose_norm_p10 | pose geometry | 0 | 0 | 0 |
| dist_front_right_paw_to_nose_norm_p90 | pose geometry | 0 | 0 | 0 |
| dist_front_right_paw_to_nose_norm_periodicity | pose geometry | 0 | 0 | 0 |
| dist_front_right_paw_to_nose_norm_std | kinematics | 0 | 0 | 0 |
| dist_front_right_paw_to_nose_norm_trend | pose geometry | 0 | 0 | 0 |
| dist_front_right_paw_to_nose_p10 | pose geometry | 0 | 0 | 0 |
| dist_front_right_paw_to_nose_p90 | pose geometry | 0 | 0 | 0 |
| dist_front_right_paw_to_nose_periodicity | pose geometry | 0 | 0 | 0 |
| dist_front_right_paw_to_nose_std | kinematics | 0 | 0 | 0 |
| dist_front_right_paw_to_nose_trend | pose geometry | 0 | 0 | 0 |

| | | | | |
|---|---|---|---|---|
| dist_front_right_paw_to_right_ear_delta | kinematics | 0 | 0 | 0 |
| dist_front_right_paw_to_right_ear_energy | kinematics | 0 | 0 | 0 |
| dist_front_right_paw_to_right_ear_max | pose geometry | 0 | 0 | 0 |
| dist_front_right_paw_to_right_ear_mean | pose geometry | 0 | 0 | 0 |
| dist_front_right_paw_to_right_ear_median | pose geometry | 0 | 0 | 0 |
| dist_front_right_paw_to_right_ear_norm_delta | kinematics | 0 | 0 | 0 |
| dist_front_right_paw_to_right_ear_norm_energy | kinematics | 0 | 0 | 0 |
| dist_front_right_paw_to_right_ear_norm_max | pose geometry | 0 | 0 | 0 |
| dist_front_right_paw_to_right_ear_norm_mean | pose geometry | 0 | 0 | 0 |
| dist_front_right_paw_to_right_ear_norm_median | pose geometry | 0 | 0 | 0 |
| dist_front_right_paw_to_right_ear_norm_p10 | pose geometry | 0 | 0 | 0 |
| dist_front_right_paw_to_right_ear_norm_p90 | pose geometry | 0 | 0 | 0 |
| dist_front_right_paw_to_right_ear_norm_periodicity | pose geometry | 0 | 0 | 0 |
| dist_front_right_paw_to_right_ear_norm_std | kinematics | 0 | 0 | 0 |
| dist_front_right_paw_to_right_ear_norm_trend | pose geometry | 0 | 0 | 0 |
| dist_front_right_paw_to_right_ear_p10 | pose geometry | 0 | 0 | 0 |
| dist_front_right_paw_to_right_ear_p90 | pose geometry | 0 | 0 | 0 |
| dist_front_right_paw_to_right_ear_periodicity | pose geometry | 0 | 0 | 0 |
| dist_front_right_paw_to_right_ear_std | kinematics | 0 | 0 | 0 |
| dist_front_right_paw_to_right_ear_trend | pose geometry | 0 | 0 | 0 |
| dist_front_right_paw_to_tail_base_delta | kinematics | 0 | 0 | 0 |
| dist_front_right_paw_to_tail_base_energy | kinematics | 0 | 0 | 0 |
| dist_front_right_paw_to_tail_base_max | pose geometry | 0 | 0 | 0 |
| dist_front_right_paw_to_tail_base_mean | pose geometry | 0 | 0 | 0 |
| dist_front_right_paw_to_tail_base_median | pose geometry | 0 | 0 | 0 |
| dist_front_right_paw_to_tail_base_norm_delta | kinematics | 0 | 0 | 0 |
| dist_front_right_paw_to_tail_base_norm_energy | kinematics | 0 | 0 | 0 |
| dist_front_right_paw_to_tail_base_norm_max | pose geometry | 0 | 0 | 0 |
| dist_front_right_paw_to_tail_base_norm_mean | pose geometry | 0 | 0 | 0 |
| dist_front_right_paw_to_tail_base_norm_median | pose geometry | 0 | 0 | 0 |
| dist_front_right_paw_to_tail_base_norm_p10 | pose geometry | 0 | 0 | 0 |
| dist_front_right_paw_to_tail_base_norm_p90 | pose geometry | 0 | 0 | 0 |
| dist_front_right_paw_to_tail_base_norm_periodicity | pose geometry | 0 | 0 | 0 |
| dist_front_right_paw_to_tail_base_norm_std | kinematics | 0 | 0 | 0 |
| dist_front_right_paw_to_tail_base_norm_trend | pose geometry | 0 | 0 | 0 |
| dist_front_right_paw_to_tail_base_p10 | pose geometry | 0 | 0 | 0 |
| dist_front_right_paw_to_tail_base_p90 | pose geometry | 0 | 0 | 0 |
| dist_front_right_paw_to_tail_base_periodicity | pose geometry | 0 | 0 | 0 |
| dist_front_right_paw_to_tail_base_std | kinematics | 0 | 0 | 0 |
| dist_front_right_paw_to_tail_base_trend | pose geometry | 0 | 0 | 0 |
| dist_lateral_left_to_lateral_right_delta | kinematics | 0 | 0 | 1 |
| dist_lateral_left_to_lateral_right_energy | kinematics | 0 | 0 | 1 |
| dist_lateral_left_to_lateral_right_max | pose geometry | 0 | 0 | 1 |
| dist_lateral_left_to_lateral_right_mean | pose geometry | 0 | 0 | 1 |

| | | | | |
|---|---|---|---|---|
| dist_lateral_left_to_lateral_right_median | pose geometry | 0 | 0 | 1 |
| dist_lateral_left_to_lateral_right_norm_delta | kinematics | 0 | 0 | 1 |
| dist_lateral_left_to_lateral_right_norm_energy | kinematics | 0 | 0 | 1 |
| dist_lateral_left_to_lateral_right_norm_max | pose geometry | 0 | 0 | 1 |
| dist_lateral_left_to_lateral_right_norm_mean | pose geometry | 0 | 0 | 1 |
| dist_lateral_left_to_lateral_right_norm_median | pose geometry | 0 | 0 | 1 |
| dist_lateral_left_to_lateral_right_norm_p10 | pose geometry | 0 | 0 | 1 |
| dist_lateral_left_to_lateral_right_norm_p90 | pose geometry | 0 | 0 | 1 |
| dist_lateral_left_to_lateral_right_norm_periodicity | pose geometry | 0 | 0 | 1 |
| dist_lateral_left_to_lateral_right_norm_std | kinematics | 0 | 0 | 1 |
| dist_lateral_left_to_lateral_right_norm_trend | pose geometry | 0 | 0 | 1 |
| dist_lateral_left_to_lateral_right_p10 | pose geometry | 0 | 0 | 1 |
| dist_lateral_left_to_lateral_right_p90 | pose geometry | 0 | 0 | 1 |
| dist_lateral_left_to_lateral_right_periodicity | pose geometry | 0 | 0 | 1 |
| dist_lateral_left_to_lateral_right_std | kinematics | 0 | 0 | 1 |
| dist_lateral_left_to_lateral_right_trend | pose geometry | 0 | 0 | 1 |
| dist_lateral_left_to_nose_delta | kinematics | 0 | 0 | 1 |
| dist_lateral_left_to_nose_energy | kinematics | 0 | 0 | 1 |
| dist_lateral_left_to_nose_max | pose geometry | 0 | 0 | 1 |
| dist_lateral_left_to_nose_mean | pose geometry | 0 | 0 | 1 |
| dist_lateral_left_to_nose_median | pose geometry | 0 | 0 | 1 |
| dist_lateral_left_to_nose_norm_delta | kinematics | 0 | 0 | 1 |
| dist_lateral_left_to_nose_norm_energy | kinematics | 0 | 0 | 1 |
| dist_lateral_left_to_nose_norm_max | pose geometry | 0 | 0 | 1 |
| dist_lateral_left_to_nose_norm_mean | pose geometry | 0 | 0 | 1 |
| dist_lateral_left_to_nose_norm_median | pose geometry | 0 | 0 | 1 |
| dist_lateral_left_to_nose_norm_p10 | pose geometry | 0 | 0 | 1 |
| dist_lateral_left_to_nose_norm_p90 | pose geometry | 0 | 0 | 1 |
| dist_lateral_left_to_nose_norm_periodicity | pose geometry | 0 | 0 | 1 |
| dist_lateral_left_to_nose_norm_std | kinematics | 0 | 0 | 1 |
| dist_lateral_left_to_nose_norm_trend | pose geometry | 0 | 0 | 1 |
| dist_lateral_left_to_nose_p10 | pose geometry | 0 | 0 | 1 |
| dist_lateral_left_to_nose_p90 | pose geometry | 0 | 0 | 1 |
| dist_lateral_left_to_nose_periodicity | pose geometry | 0 | 0 | 1 |
| dist_lateral_left_to_nose_std | kinematics | 0 | 0 | 1 |
| dist_lateral_left_to_nose_trend | pose geometry | 0 | 0 | 1 |
| dist_lateral_left_to_tail_base_delta | kinematics | 0 | 0 | 1 |
| dist_lateral_left_to_tail_base_energy | kinematics | 0 | 0 | 1 |
| dist_lateral_left_to_tail_base_max | pose geometry | 0 | 0 | 1 |
| dist_lateral_left_to_tail_base_mean | pose geometry | 0 | 0 | 1 |
| dist_lateral_left_to_tail_base_median | pose geometry | 0 | 0 | 1 |
| dist_lateral_left_to_tail_base_norm_delta | kinematics | 0 | 0 | 1 |
| dist_lateral_left_to_tail_base_norm_energy | kinematics | 0 | 0 | 1 |
| dist_lateral_left_to_tail_base_norm_max | pose geometry | 0 | 0 | 1 |

| | | | | |
|---|---|---|---|---|
| dist_lateral_left_to_tail_base_norm_mean | pose geometry | 0 | 0 | 1 |
| dist_lateral_left_to_tail_base_norm_median | pose geometry | 0 | 0 | 1 |
| dist_lateral_left_to_tail_base_norm_p10 | pose geometry | 0 | 0 | 1 |
| dist_lateral_left_to_tail_base_norm_p90 | pose geometry | 0 | 0 | 1 |
| dist_lateral_left_to_tail_base_norm_periodicity | pose geometry | 0 | 0 | 1 |
| dist_lateral_left_to_tail_base_norm_std | kinematics | 0 | 0 | 1 |
| dist_lateral_left_to_tail_base_norm_trend | pose geometry | 0 | 0 | 1 |
| dist_lateral_left_to_tail_base_p10 | pose geometry | 0 | 0 | 1 |
| dist_lateral_left_to_tail_base_p90 | pose geometry | 0 | 0 | 1 |
| dist_lateral_left_to_tail_base_periodicity | pose geometry | 0 | 0 | 1 |
| dist_lateral_left_to_tail_base_std | kinematics | 0 | 0 | 1 |
| dist_lateral_left_to_tail_base_trend | pose geometry | 0 | 0 | 1 |
| dist_lateral_right_to_nose_delta | kinematics | 0 | 0 | 1 |
| dist_lateral_right_to_nose_energy | kinematics | 0 | 0 | 1 |
| dist_lateral_right_to_nose_max | pose geometry | 0 | 0 | 1 |
| dist_lateral_right_to_nose_mean | pose geometry | 0 | 0 | 1 |
| dist_lateral_right_to_nose_median | pose geometry | 0 | 0 | 1 |
| dist_lateral_right_to_nose_norm_delta | kinematics | 0 | 0 | 1 |
| dist_lateral_right_to_nose_norm_energy | kinematics | 0 | 0 | 1 |
| dist_lateral_right_to_nose_norm_max | pose geometry | 0 | 0 | 1 |
| dist_lateral_right_to_nose_norm_mean | pose geometry | 0 | 0 | 1 |
| dist_lateral_right_to_nose_norm_median | pose geometry | 0 | 0 | 1 |
| dist_lateral_right_to_nose_norm_p10 | pose geometry | 0 | 0 | 1 |
| dist_lateral_right_to_nose_norm_p90 | pose geometry | 0 | 0 | 1 |
| dist_lateral_right_to_nose_norm_periodicity | pose geometry | 0 | 0 | 1 |
| dist_lateral_right_to_nose_norm_std | kinematics | 0 | 0 | 1 |
| dist_lateral_right_to_nose_norm_trend | pose geometry | 0 | 0 | 1 |
| dist_lateral_right_to_nose_p10 | pose geometry | 0 | 0 | 1 |
| dist_lateral_right_to_nose_p90 | pose geometry | 0 | 0 | 1 |
| dist_lateral_right_to_nose_periodicity | pose geometry | 0 | 0 | 1 |
| dist_lateral_right_to_nose_std | kinematics | 0 | 0 | 1 |
| dist_lateral_right_to_nose_trend | pose geometry | 0 | 0 | 1 |
| dist_lateral_right_to_tail_base_delta | kinematics | 0 | 0 | 1 |
| dist_lateral_right_to_tail_base_energy | kinematics | 0 | 0 | 1 |
| dist_lateral_right_to_tail_base_max | pose geometry | 0 | 0 | 1 |
| dist_lateral_right_to_tail_base_mean | pose geometry | 0 | 0 | 1 |
| dist_lateral_right_to_tail_base_median | pose geometry | 0 | 0 | 1 |
| dist_lateral_right_to_tail_base_norm_delta | kinematics | 0 | 0 | 1 |
| dist_lateral_right_to_tail_base_norm_energy | kinematics | 0 | 0 | 1 |
| dist_lateral_right_to_tail_base_norm_max | pose geometry | 0 | 0 | 1 |
| dist_lateral_right_to_tail_base_norm_mean | pose geometry | 0 | 0 | 1 |
| dist_lateral_right_to_tail_base_norm_median | pose geometry | 0 | 0 | 1 |
| dist_lateral_right_to_tail_base_norm_p10 | pose geometry | 0 | 0 | 1 |
| dist_lateral_right_to_tail_base_norm_p90 | pose geometry | 0 | 0 | 1 |

| | | | | |
|---|---|---|---|---|
| dist_lateral_right_to_tail_base_norm_periodicity | pose geometry | 0 | 0 | 1 |
| dist_lateral_right_to_tail_base_norm_std | kinematics | 0 | 0 | 1 |
| dist_lateral_right_to_tail_base_norm_trend | pose geometry | 0 | 0 | 1 |
| dist_lateral_right_to_tail_base_p10 | pose geometry | 0 | 0 | 1 |
| dist_lateral_right_to_tail_base_p90 | pose geometry | 0 | 0 | 1 |
| dist_lateral_right_to_tail_base_periodicity | pose geometry | 0 | 0 | 1 |
| dist_lateral_right_to_tail_base_std | kinematics | 0 | 0 | 1 |
| dist_lateral_right_to_tail_base_trend | pose geometry | 0 | 0 | 1 |
| dist_left_body_to_left_ear_delta | kinematics | 1 | 0 | 0 |
| dist_left_body_to_left_ear_energy | kinematics | 1 | 0 | 0 |
| dist_left_body_to_left_ear_max | pose geometry | 1 | 0 | 0 |
| dist_left_body_to_left_ear_mean | pose geometry | 1 | 0 | 0 |
| dist_left_body_to_left_ear_median | pose geometry | 1 | 0 | 0 |
| dist_left_body_to_left_ear_norm_delta | kinematics | 1 | 0 | 0 |
| dist_left_body_to_left_ear_norm_energy | kinematics | 1 | 0 | 0 |
| dist_left_body_to_left_ear_norm_max | pose geometry | 1 | 0 | 0 |
| dist_left_body_to_left_ear_norm_mean | pose geometry | 1 | 0 | 0 |
| dist_left_body_to_left_ear_norm_median | pose geometry | 1 | 0 | 0 |
| dist_left_body_to_left_ear_norm_p10 | pose geometry | 1 | 0 | 0 |
| dist_left_body_to_left_ear_norm_p90 | pose geometry | 1 | 0 | 0 |
| dist_left_body_to_left_ear_norm_periodicity | pose geometry | 1 | 0 | 0 |
| dist_left_body_to_left_ear_norm_std | kinematics | 1 | 0 | 0 |
| dist_left_body_to_left_ear_norm_trend | pose geometry | 1 | 0 | 0 |
| dist_left_body_to_left_ear_p10 | pose geometry | 1 | 0 | 0 |
| dist_left_body_to_left_ear_p90 | pose geometry | 1 | 0 | 0 |
| dist_left_body_to_left_ear_periodicity | pose geometry | 1 | 0 | 0 |
| dist_left_body_to_left_ear_std | kinematics | 1 | 0 | 0 |
| dist_left_body_to_left_ear_trend | pose geometry | 1 | 0 | 0 |
| dist_left_body_to_nose_delta | kinematics | 1 | 0 | 0 |
| dist_left_body_to_nose_energy | kinematics | 1 | 0 | 0 |
| dist_left_body_to_nose_max | pose geometry | 1 | 0 | 0 |
| dist_left_body_to_nose_mean | pose geometry | 1 | 0 | 0 |
| dist_left_body_to_nose_median | pose geometry | 1 | 0 | 0 |
| dist_left_body_to_nose_norm_delta | kinematics | 1 | 0 | 0 |
| dist_left_body_to_nose_norm_energy | kinematics | 1 | 0 | 0 |
| dist_left_body_to_nose_norm_max | pose geometry | 1 | 0 | 0 |
| dist_left_body_to_nose_norm_mean | pose geometry | 1 | 0 | 0 |
| dist_left_body_to_nose_norm_median | pose geometry | 1 | 0 | 0 |
| dist_left_body_to_nose_norm_p10 | pose geometry | 1 | 0 | 0 |
| dist_left_body_to_nose_norm_p90 | pose geometry | 1 | 0 | 0 |
| dist_left_body_to_nose_norm_periodicity | pose geometry | 1 | 0 | 0 |
| dist_left_body_to_nose_norm_std | kinematics | 1 | 0 | 0 |
| dist_left_body_to_nose_norm_trend | pose geometry | 1 | 0 | 0 |
| dist_left_body_to_nose_p10 | pose geometry | 1 | 0 | 0 |

| | | | | |
|---|---|---|---|---|
| dist_left_body_to_nose_p90 | pose geometry | 1 | 0 | 0 |
| dist_left_body_to_nose_periodicity | pose geometry | 1 | 0 | 0 |
| dist_left_body_to_nose_std | kinematics | 1 | 0 | 0 |
| dist_left_body_to_nose_trend | pose geometry | 1 | 0 | 0 |
| dist_left_body_to_right_body_delta | kinematics | 1 | 0 | 0 |
| dist_left_body_to_right_body_energy | kinematics | 1 | 0 | 0 |
| dist_left_body_to_right_body_max | pose geometry | 1 | 0 | 0 |
| dist_left_body_to_right_body_mean | pose geometry | 1 | 0 | 0 |
| dist_left_body_to_right_body_median | pose geometry | 1 | 0 | 0 |
| dist_left_body_to_right_body_norm_delta | kinematics | 1 | 0 | 0 |
| dist_left_body_to_right_body_norm_energy | kinematics | 1 | 0 | 0 |
| dist_left_body_to_right_body_norm_max | pose geometry | 1 | 0 | 0 |
| dist_left_body_to_right_body_norm_mean | pose geometry | 1 | 0 | 0 |
| dist_left_body_to_right_body_norm_median | pose geometry | 1 | 0 | 0 |
| dist_left_body_to_right_body_norm_p10 | pose geometry | 1 | 0 | 0 |
| dist_left_body_to_right_body_norm_p90 | pose geometry | 1 | 0 | 0 |
| dist_left_body_to_right_body_norm_periodicity | pose geometry | 1 | 0 | 0 |
| dist_left_body_to_right_body_norm_std | kinematics | 1 | 0 | 0 |
| dist_left_body_to_right_body_norm_trend | pose geometry | 1 | 0 | 0 |
| dist_left_body_to_right_body_p10 | pose geometry | 1 | 0 | 0 |
| dist_left_body_to_right_body_p90 | pose geometry | 1 | 0 | 0 |
| dist_left_body_to_right_body_periodicity | pose geometry | 1 | 0 | 0 |
| dist_left_body_to_right_body_std | kinematics | 1 | 0 | 0 |
| dist_left_body_to_right_body_trend | pose geometry | 1 | 0 | 0 |
| dist_left_body_to_right_ear_delta | kinematics | 1 | 0 | 0 |
| dist_left_body_to_right_ear_energy | kinematics | 1 | 0 | 0 |
| dist_left_body_to_right_ear_max | pose geometry | 1 | 0 | 0 |
| dist_left_body_to_right_ear_mean | pose geometry | 1 | 0 | 0 |
| dist_left_body_to_right_ear_median | pose geometry | 1 | 0 | 0 |
| dist_left_body_to_right_ear_norm_delta | kinematics | 1 | 0 | 0 |
| dist_left_body_to_right_ear_norm_energy | kinematics | 1 | 0 | 0 |
| dist_left_body_to_right_ear_norm_max | pose geometry | 1 | 0 | 0 |
| dist_left_body_to_right_ear_norm_mean | pose geometry | 1 | 0 | 0 |
| dist_left_body_to_right_ear_norm_median | pose geometry | 1 | 0 | 0 |
| dist_left_body_to_right_ear_norm_p10 | pose geometry | 1 | 0 | 0 |
| dist_left_body_to_right_ear_norm_p90 | pose geometry | 1 | 0 | 0 |
| dist_left_body_to_right_ear_norm_periodicity | pose geometry | 1 | 0 | 0 |
| dist_left_body_to_right_ear_norm_std | kinematics | 1 | 0 | 0 |
| dist_left_body_to_right_ear_norm_trend | pose geometry | 1 | 0 | 0 |
| dist_left_body_to_right_ear_p10 | pose geometry | 1 | 0 | 0 |
| dist_left_body_to_right_ear_p90 | pose geometry | 1 | 0 | 0 |
| dist_left_body_to_right_ear_periodicity | pose geometry | 1 | 0 | 0 |
| dist_left_body_to_right_ear_std | kinematics | 1 | 0 | 0 |
| dist_left_body_to_right_ear_trend | pose geometry | 1 | 0 | 0 |

| | | | | |
|---|---|---|---|---|
| dist_left_body_to_tail_base_delta | kinematics | 1 | 0 | 0 |
| dist_left_body_to_tail_base_energy | kinematics | 1 | 0 | 0 |
| dist_left_body_to_tail_base_max | pose geometry | 1 | 0 | 0 |
| dist_left_body_to_tail_base_mean | pose geometry | 1 | 0 | 0 |
| dist_left_body_to_tail_base_median | pose geometry | 1 | 0 | 0 |
| dist_left_body_to_tail_base_norm_delta | kinematics | 1 | 0 | 0 |
| dist_left_body_to_tail_base_norm_energy | kinematics | 1 | 0 | 0 |
| dist_left_body_to_tail_base_norm_max | pose geometry | 1 | 0 | 0 |
| dist_left_body_to_tail_base_norm_mean | pose geometry | 1 | 0 | 0 |
| dist_left_body_to_tail_base_norm_median | pose geometry | 1 | 0 | 0 |
| dist_left_body_to_tail_base_norm_p10 | pose geometry | 1 | 0 | 0 |
| dist_left_body_to_tail_base_norm_p90 | pose geometry | 1 | 0 | 0 |
| dist_left_body_to_tail_base_norm_periodicity | pose geometry | 1 | 0 | 0 |
| dist_left_body_to_tail_base_norm_std | kinematics | 1 | 0 | 0 |
| dist_left_body_to_tail_base_norm_trend | pose geometry | 1 | 0 | 0 |
| dist_left_body_to_tail_base_p10 | pose geometry | 1 | 0 | 0 |
| dist_left_body_to_tail_base_p90 | pose geometry | 1 | 0 | 0 |
| dist_left_body_to_tail_base_periodicity | pose geometry | 1 | 0 | 0 |
| dist_left_body_to_tail_base_std | kinematics | 1 | 0 | 0 |
| dist_left_body_to_tail_base_trend | pose geometry | 1 | 0 | 0 |
| dist_left_ear_to_mid_back_delta | kinematics | 0 | 0 | 0 |
| dist_left_ear_to_mid_back_energy | kinematics | 0 | 0 | 0 |
| dist_left_ear_to_mid_back_max | pose geometry | 0 | 0 | 0 |
| dist_left_ear_to_mid_back_mean | pose geometry | 0 | 0 | 0 |
| dist_left_ear_to_mid_back_median | pose geometry | 0 | 0 | 0 |
| dist_left_ear_to_mid_back_norm_delta | kinematics | 0 | 0 | 0 |
| dist_left_ear_to_mid_back_norm_energy | kinematics | 0 | 0 | 0 |
| dist_left_ear_to_mid_back_norm_max | pose geometry | 0 | 0 | 0 |
| dist_left_ear_to_mid_back_norm_mean | pose geometry | 0 | 0 | 0 |
| dist_left_ear_to_mid_back_norm_median | pose geometry | 0 | 0 | 0 |
| dist_left_ear_to_mid_back_norm_p10 | pose geometry | 0 | 0 | 0 |
| dist_left_ear_to_mid_back_norm_p90 | pose geometry | 0 | 0 | 0 |
| dist_left_ear_to_mid_back_norm_periodicity | pose geometry | 0 | 0 | 0 |
| dist_left_ear_to_mid_back_norm_std | kinematics | 0 | 0 | 0 |
| dist_left_ear_to_mid_back_norm_trend | pose geometry | 0 | 0 | 0 |
| dist_left_ear_to_mid_back_p10 | pose geometry | 0 | 0 | 0 |
| dist_left_ear_to_mid_back_p90 | pose geometry | 0 | 0 | 0 |
| dist_left_ear_to_mid_back_periodicity | pose geometry | 0 | 0 | 0 |
| dist_left_ear_to_mid_back_std | kinematics | 0 | 0 | 0 |
| dist_left_ear_to_mid_back_trend | pose geometry | 0 | 0 | 0 |
| dist_left_ear_to_nose_delta | kinematics | 1 | 0 | 0 |
| dist_left_ear_to_nose_energy | kinematics | 1 | 0 | 0 |
| dist_left_ear_to_nose_max | pose geometry | 1 | 0 | 0 |
| dist_left_ear_to_nose_mean | pose geometry | 1 | 0 | 0 |

| | | | | |
|---|---|---|---|---|
| dist_left_ear_to_nose_median | pose geometry | 1 | 0 | 0 |
| dist_left_ear_to_nose_norm_delta | kinematics | 1 | 0 | 0 |
| dist_left_ear_to_nose_norm_energy | kinematics | 1 | 0 | 0 |
| dist_left_ear_to_nose_norm_max | pose geometry | 1 | 0 | 0 |
| dist_left_ear_to_nose_norm_mean | pose geometry | 1 | 0 | 0 |
| dist_left_ear_to_nose_norm_median | pose geometry | 1 | 0 | 0 |
| dist_left_ear_to_nose_norm_p10 | pose geometry | 1 | 0 | 0 |
| dist_left_ear_to_nose_norm_p90 | pose geometry | 1 | 0 | 0 |
| dist_left_ear_to_nose_norm_periodicity | pose geometry | 1 | 0 | 0 |
| dist_left_ear_to_nose_norm_std | kinematics | 1 | 0 | 0 |
| dist_left_ear_to_nose_norm_trend | pose geometry | 1 | 0 | 0 |
| dist_left_ear_to_nose_p10 | pose geometry | 1 | 0 | 0 |
| dist_left_ear_to_nose_p90 | pose geometry | 1 | 0 | 0 |
| dist_left_ear_to_nose_periodicity | pose geometry | 1 | 0 | 0 |
| dist_left_ear_to_nose_std | kinematics | 1 | 0 | 0 |
| dist_left_ear_to_nose_trend | pose geometry | 1 | 0 | 0 |
| dist_left_ear_to_right_body_delta | kinematics | 1 | 0 | 0 |
| dist_left_ear_to_right_body_energy | kinematics | 1 | 0 | 0 |
| dist_left_ear_to_right_body_max | pose geometry | 1 | 0 | 0 |
| dist_left_ear_to_right_body_mean | pose geometry | 1 | 0 | 0 |
| dist_left_ear_to_right_body_median | pose geometry | 1 | 0 | 0 |
| dist_left_ear_to_right_body_norm_delta | kinematics | 1 | 0 | 0 |
| dist_left_ear_to_right_body_norm_energy | kinematics | 1 | 0 | 0 |
| dist_left_ear_to_right_body_norm_max | pose geometry | 1 | 0 | 0 |
| dist_left_ear_to_right_body_norm_mean | pose geometry | 1 | 0 | 0 |
| dist_left_ear_to_right_body_norm_median | pose geometry | 1 | 0 | 0 |
| dist_left_ear_to_right_body_norm_p10 | pose geometry | 1 | 0 | 0 |
| dist_left_ear_to_right_body_norm_p90 | pose geometry | 1 | 0 | 0 |
| dist_left_ear_to_right_body_norm_periodicity | pose geometry | 1 | 0 | 0 |
| dist_left_ear_to_right_body_norm_std | kinematics | 1 | 0 | 0 |
| dist_left_ear_to_right_body_norm_trend | pose geometry | 1 | 0 | 0 |
| dist_left_ear_to_right_body_p10 | pose geometry | 1 | 0 | 0 |
| dist_left_ear_to_right_body_p90 | pose geometry | 1 | 0 | 0 |
| dist_left_ear_to_right_body_periodicity | pose geometry | 1 | 0 | 0 |
| dist_left_ear_to_right_body_std | kinematics | 1 | 0 | 0 |
| dist_left_ear_to_right_body_trend | pose geometry | 1 | 0 | 0 |
| dist_left_ear_to_right_ear_delta | kinematics | 1 | 0 | 0 |
| dist_left_ear_to_right_ear_energy | kinematics | 1 | 0 | 0 |
| dist_left_ear_to_right_ear_max | pose geometry | 1 | 0 | 0 |
| dist_left_ear_to_right_ear_mean | pose geometry | 1 | 0 | 0 |
| dist_left_ear_to_right_ear_median | pose geometry | 1 | 0 | 0 |
| dist_left_ear_to_right_ear_norm_delta | kinematics | 1 | 0 | 0 |
| dist_left_ear_to_right_ear_norm_energy | kinematics | 1 | 0 | 0 |
| dist_left_ear_to_right_ear_norm_max | pose geometry | 1 | 0 | 0 |

| | | | | |
|---|---|---|---|---|
| dist_left_ear_to_right_ear_norm_mean | pose geometry | 1 | 0 | 0 |
| dist_left_ear_to_right_ear_norm_median | pose geometry | 1 | 0 | 0 |
| dist_left_ear_to_right_ear_norm_p10 | pose geometry | 1 | 0 | 0 |
| dist_left_ear_to_right_ear_norm_p90 | pose geometry | 1 | 0 | 0 |
| dist_left_ear_to_right_ear_norm_periodicity | pose geometry | 1 | 0 | 0 |
| dist_left_ear_to_right_ear_norm_std | kinematics | 1 | 0 | 0 |
| dist_left_ear_to_right_ear_norm_trend | pose geometry | 1 | 0 | 0 |
| dist_left_ear_to_right_ear_p10 | pose geometry | 1 | 0 | 0 |
| dist_left_ear_to_right_ear_p90 | pose geometry | 1 | 0 | 0 |
| dist_left_ear_to_right_ear_periodicity | pose geometry | 1 | 0 | 0 |
| dist_left_ear_to_right_ear_std | kinematics | 1 | 0 | 0 |
| dist_left_ear_to_right_ear_trend | pose geometry | 1 | 0 | 0 |
| dist_left_ear_to_tail_base_delta | kinematics | 1 | 0 | 0 |
| dist_left_ear_to_tail_base_energy | kinematics | 1 | 0 | 0 |
| dist_left_ear_to_tail_base_max | pose geometry | 1 | 0 | 0 |
| dist_left_ear_to_tail_base_mean | pose geometry | 1 | 0 | 0 |
| dist_left_ear_to_tail_base_median | pose geometry | 1 | 0 | 0 |
| dist_left_ear_to_tail_base_norm_delta | kinematics | 1 | 0 | 0 |
| dist_left_ear_to_tail_base_norm_energy | kinematics | 1 | 0 | 0 |
| dist_left_ear_to_tail_base_norm_max | pose geometry | 1 | 0 | 0 |
| dist_left_ear_to_tail_base_norm_mean | pose geometry | 1 | 0 | 0 |
| dist_left_ear_to_tail_base_norm_median | pose geometry | 1 | 0 | 0 |
| dist_left_ear_to_tail_base_norm_p10 | pose geometry | 1 | 0 | 0 |
| dist_left_ear_to_tail_base_norm_p90 | pose geometry | 1 | 0 | 0 |
| dist_left_ear_to_tail_base_norm_periodicity | pose geometry | 1 | 0 | 0 |
| dist_left_ear_to_tail_base_norm_std | kinematics | 1 | 0 | 0 |
| dist_left_ear_to_tail_base_norm_trend | pose geometry | 1 | 0 | 0 |
| dist_left_ear_to_tail_base_p10 | pose geometry | 1 | 0 | 0 |
| dist_left_ear_to_tail_base_p90 | pose geometry | 1 | 0 | 0 |
| dist_left_ear_to_tail_base_periodicity | pose geometry | 1 | 0 | 0 |
| dist_left_ear_to_tail_base_std | kinematics | 1 | 0 | 0 |
| dist_left_ear_to_tail_base_trend | pose geometry | 1 | 0 | 0 |
| dist_mid_back_to_nose_delta | kinematics | 0 | 0 | 0 |
| dist_mid_back_to_nose_energy | kinematics | 0 | 0 | 0 |
| dist_mid_back_to_nose_max | pose geometry | 0 | 0 | 0 |
| dist_mid_back_to_nose_mean | pose geometry | 0 | 0 | 0 |
| dist_mid_back_to_nose_median | pose geometry | 0 | 0 | 0 |
| dist_mid_back_to_nose_norm_delta | kinematics | 0 | 0 | 0 |
| dist_mid_back_to_nose_norm_energy | kinematics | 0 | 0 | 0 |
| dist_mid_back_to_nose_norm_max | pose geometry | 0 | 0 | 0 |
| dist_mid_back_to_nose_norm_mean | pose geometry | 0 | 0 | 0 |
| dist_mid_back_to_nose_norm_median | pose geometry | 0 | 0 | 0 |
| dist_mid_back_to_nose_norm_p10 | pose geometry | 0 | 0 | 0 |
| dist_mid_back_to_nose_norm_p90 | pose geometry | 0 | 0 | 0 |

| | | | | |
|---|---|---|---|---|
| dist_mid_back_to_nose_norm_periodicity | pose geometry | 0 | 0 | 0 |
| dist_mid_back_to_nose_norm_std | kinematics | 0 | 0 | 0 |
| dist_mid_back_to_nose_norm_trend | pose geometry | 0 | 0 | 0 |
| dist_mid_back_to_nose_p10 | pose geometry | 0 | 0 | 0 |
| dist_mid_back_to_nose_p90 | pose geometry | 0 | 0 | 0 |
| dist_mid_back_to_nose_periodicity | pose geometry | 0 | 0 | 0 |
| dist_mid_back_to_nose_std | kinematics | 0 | 0 | 0 |
| dist_mid_back_to_nose_trend | pose geometry | 0 | 0 | 0 |
| dist_mid_back_to_right_ear_delta | kinematics | 0 | 0 | 0 |
| dist_mid_back_to_right_ear_energy | kinematics | 0 | 0 | 0 |
| dist_mid_back_to_right_ear_max | pose geometry | 0 | 0 | 0 |
| dist_mid_back_to_right_ear_mean | pose geometry | 0 | 0 | 0 |
| dist_mid_back_to_right_ear_median | pose geometry | 0 | 0 | 0 |
| dist_mid_back_to_right_ear_norm_delta | kinematics | 0 | 0 | 0 |
| dist_mid_back_to_right_ear_norm_energy | kinematics | 0 | 0 | 0 |
| dist_mid_back_to_right_ear_norm_max | pose geometry | 0 | 0 | 0 |
| dist_mid_back_to_right_ear_norm_mean | pose geometry | 0 | 0 | 0 |
| dist_mid_back_to_right_ear_norm_median | pose geometry | 0 | 0 | 0 |
| dist_mid_back_to_right_ear_norm_p10 | pose geometry | 0 | 0 | 0 |
| dist_mid_back_to_right_ear_norm_p90 | pose geometry | 0 | 0 | 0 |
| dist_mid_back_to_right_ear_norm_periodicity | pose geometry | 0 | 0 | 0 |
| dist_mid_back_to_right_ear_norm_std | kinematics | 0 | 0 | 0 |
| dist_mid_back_to_right_ear_norm_trend | pose geometry | 0 | 0 | 0 |
| dist_mid_back_to_right_ear_p10 | pose geometry | 0 | 0 | 0 |
| dist_mid_back_to_right_ear_p90 | pose geometry | 0 | 0 | 0 |
| dist_mid_back_to_right_ear_periodicity | pose geometry | 0 | 0 | 0 |
| dist_mid_back_to_right_ear_std | kinematics | 0 | 0 | 0 |
| dist_mid_back_to_right_ear_trend | pose geometry | 0 | 0 | 0 |
| dist_mid_back_to_tail_base_delta | kinematics | 0 | 0 | 0 |
| dist_mid_back_to_tail_base_energy | kinematics | 0 | 0 | 0 |
| dist_mid_back_to_tail_base_max | pose geometry | 0 | 0 | 0 |
| dist_mid_back_to_tail_base_mean | pose geometry | 0 | 0 | 0 |
| dist_mid_back_to_tail_base_median | pose geometry | 0 | 0 | 0 |
| dist_mid_back_to_tail_base_norm_delta | kinematics | 0 | 0 | 0 |
| dist_mid_back_to_tail_base_norm_energy | kinematics | 0 | 0 | 0 |
| dist_mid_back_to_tail_base_norm_max | pose geometry | 0 | 0 | 0 |
| dist_mid_back_to_tail_base_norm_mean | pose geometry | 0 | 0 | 0 |
| dist_mid_back_to_tail_base_norm_median | pose geometry | 0 | 0 | 0 |
| dist_mid_back_to_tail_base_norm_p10 | pose geometry | 0 | 0 | 0 |
| dist_mid_back_to_tail_base_norm_p90 | pose geometry | 0 | 0 | 0 |
| dist_mid_back_to_tail_base_norm_periodicity | pose geometry | 0 | 0 | 0 |
| dist_mid_back_to_tail_base_norm_std | kinematics | 0 | 0 | 0 |
| dist_mid_back_to_tail_base_norm_trend | pose geometry | 0 | 0 | 0 |
| dist_mid_back_to_tail_base_p10 | pose geometry | 0 | 0 | 0 |

| | | | | |
|---|---|---|---|---|
| dist_mid_back_to_tail_base_p90 | pose geometry | 0 | 0 | 0 |
| dist_mid_back_to_tail_base_periodicity | pose geometry | 0 | 0 | 0 |
| dist_mid_back_to_tail_base_std | kinematics | 0 | 0 | 0 |
| dist_mid_back_to_tail_base_trend | pose geometry | 0 | 0 | 0 |
| dist_nose_to_right_body_delta | kinematics | 1 | 0 | 0 |
| dist_nose_to_right_body_energy | kinematics | 1 | 0 | 0 |
| dist_nose_to_right_body_max | pose geometry | 1 | 0 | 0 |
| dist_nose_to_right_body_mean | pose geometry | 1 | 0 | 0 |
| dist_nose_to_right_body_median | pose geometry | 1 | 0 | 0 |
| dist_nose_to_right_body_norm_delta | kinematics | 1 | 0 | 0 |
| dist_nose_to_right_body_norm_energy | kinematics | 1 | 0 | 0 |
| dist_nose_to_right_body_norm_max | pose geometry | 1 | 0 | 0 |
| dist_nose_to_right_body_norm_mean | pose geometry | 1 | 0 | 0 |
| dist_nose_to_right_body_norm_median | pose geometry | 1 | 0 | 0 |
| dist_nose_to_right_body_norm_p10 | pose geometry | 1 | 0 | 0 |
| dist_nose_to_right_body_norm_p90 | pose geometry | 1 | 0 | 0 |
| dist_nose_to_right_body_norm_periodicity | pose geometry | 1 | 0 | 0 |
| dist_nose_to_right_body_norm_std | kinematics | 1 | 0 | 0 |
| dist_nose_to_right_body_norm_trend | pose geometry | 1 | 0 | 0 |
| dist_nose_to_right_body_p10 | pose geometry | 1 | 0 | 0 |
| dist_nose_to_right_body_p90 | pose geometry | 1 | 0 | 0 |
| dist_nose_to_right_body_periodicity | pose geometry | 1 | 0 | 0 |
| dist_nose_to_right_body_std | kinematics | 1 | 0 | 0 |
| dist_nose_to_right_body_trend | pose geometry | 1 | 0 | 0 |
| dist_nose_to_right_ear_delta | kinematics | 1 | 0 | 0 |
| dist_nose_to_right_ear_energy | kinematics | 1 | 0 | 0 |
| dist_nose_to_right_ear_max | pose geometry | 1 | 0 | 0 |
| dist_nose_to_right_ear_mean | pose geometry | 1 | 0 | 0 |
| dist_nose_to_right_ear_median | pose geometry | 1 | 0 | 0 |
| dist_nose_to_right_ear_norm_delta | kinematics | 1 | 0 | 0 |
| dist_nose_to_right_ear_norm_energy | kinematics | 1 | 0 | 0 |
| dist_nose_to_right_ear_norm_max | pose geometry | 1 | 0 | 0 |
| dist_nose_to_right_ear_norm_mean | pose geometry | 1 | 0 | 0 |
| dist_nose_to_right_ear_norm_median | pose geometry | 1 | 0 | 0 |
| dist_nose_to_right_ear_norm_p10 | pose geometry | 1 | 0 | 0 |
| dist_nose_to_right_ear_norm_p90 | pose geometry | 1 | 0 | 0 |
| dist_nose_to_right_ear_norm_periodicity | pose geometry | 1 | 0 | 0 |
| dist_nose_to_right_ear_norm_std | kinematics | 1 | 0 | 0 |
| dist_nose_to_right_ear_norm_trend | pose geometry | 1 | 0 | 0 |
| dist_nose_to_right_ear_p10 | pose geometry | 1 | 0 | 0 |
| dist_nose_to_right_ear_p90 | pose geometry | 1 | 0 | 0 |
| dist_nose_to_right_ear_periodicity | pose geometry | 1 | 0 | 0 |
| dist_nose_to_right_ear_std | kinematics | 1 | 0 | 0 |
| dist_nose_to_right_ear_trend | pose geometry | 1 | 0 | 0 |

| | | | | |
|---|---|---|---|---|
| dist_nose_to_tail_base_delta | kinematics | 1 | 1 | 1 |
| dist_nose_to_tail_base_energy | kinematics | 1 | 1 | 1 |
| dist_nose_to_tail_base_max | pose geometry | 1 | 1 | 1 |
| dist_nose_to_tail_base_mean | pose geometry | 1 | 1 | 1 |
| dist_nose_to_tail_base_median | pose geometry | 1 | 1 | 1 |
| dist_nose_to_tail_base_p10 | pose geometry | 1 | 1 | 1 |
| dist_nose_to_tail_base_p90 | pose geometry | 1 | 1 | 1 |
| dist_nose_to_tail_base_periodicity | pose geometry | 1 | 1 | 1 |
| dist_nose_to_tail_base_std | kinematics | 1 | 1 | 1 |
| dist_nose_to_tail_base_trend | pose geometry | 1 | 1 | 1 |
| dist_right_body_to_right_ear_delta | kinematics | 1 | 0 | 0 |
| dist_right_body_to_right_ear_energy | kinematics | 1 | 0 | 0 |
| dist_right_body_to_right_ear_max | pose geometry | 1 | 0 | 0 |
| dist_right_body_to_right_ear_mean | pose geometry | 1 | 0 | 0 |
| dist_right_body_to_right_ear_median | pose geometry | 1 | 0 | 0 |
| dist_right_body_to_right_ear_norm_delta | kinematics | 1 | 0 | 0 |
| dist_right_body_to_right_ear_norm_energy | kinematics | 1 | 0 | 0 |
| dist_right_body_to_right_ear_norm_max | pose geometry | 1 | 0 | 0 |
| dist_right_body_to_right_ear_norm_mean | pose geometry | 1 | 0 | 0 |
| dist_right_body_to_right_ear_norm_median | pose geometry | 1 | 0 | 0 |
| dist_right_body_to_right_ear_norm_p10 | pose geometry | 1 | 0 | 0 |
| dist_right_body_to_right_ear_norm_p90 | pose geometry | 1 | 0 | 0 |
| dist_right_body_to_right_ear_norm_periodicity | pose geometry | 1 | 0 | 0 |
| dist_right_body_to_right_ear_norm_std | kinematics | 1 | 0 | 0 |
| dist_right_body_to_right_ear_norm_trend | pose geometry | 1 | 0 | 0 |
| dist_right_body_to_right_ear_p10 | pose geometry | 1 | 0 | 0 |
| dist_right_body_to_right_ear_p90 | pose geometry | 1 | 0 | 0 |
| dist_right_body_to_right_ear_periodicity | pose geometry | 1 | 0 | 0 |
| dist_right_body_to_right_ear_std | kinematics | 1 | 0 | 0 |
| dist_right_body_to_right_ear_trend | pose geometry | 1 | 0 | 0 |
| dist_right_body_to_tail_base_delta | kinematics | 1 | 0 | 0 |
| dist_right_body_to_tail_base_energy | kinematics | 1 | 0 | 0 |
| dist_right_body_to_tail_base_max | pose geometry | 1 | 0 | 0 |
| dist_right_body_to_tail_base_mean | pose geometry | 1 | 0 | 0 |
| dist_right_body_to_tail_base_median | pose geometry | 1 | 0 | 0 |
| dist_right_body_to_tail_base_norm_delta | kinematics | 1 | 0 | 0 |
| dist_right_body_to_tail_base_norm_energy | kinematics | 1 | 0 | 0 |
| dist_right_body_to_tail_base_norm_max | pose geometry | 1 | 0 | 0 |
| dist_right_body_to_tail_base_norm_mean | pose geometry | 1 | 0 | 0 |
| dist_right_body_to_tail_base_norm_median | pose geometry | 1 | 0 | 0 |
| dist_right_body_to_tail_base_norm_p10 | pose geometry | 1 | 0 | 0 |
| dist_right_body_to_tail_base_norm_p90 | pose geometry | 1 | 0 | 0 |
| dist_right_body_to_tail_base_norm_periodicity | pose geometry | 1 | 0 | 0 |
| dist_right_body_to_tail_base_norm_std | kinematics | 1 | 0 | 0 |

| | | | | |
|---|---|---|---|---|
| dist_right_body_to_tail_base_norm_trend | pose geometry | 1 | 0 | 0 |
| dist_right_body_to_tail_base_p10 | pose geometry | 1 | 0 | 0 |
| dist_right_body_to_tail_base_p90 | pose geometry | 1 | 0 | 0 |
| dist_right_body_to_tail_base_periodicity | pose geometry | 1 | 0 | 0 |
| dist_right_body_to_tail_base_std | kinematics | 1 | 0 | 0 |
| dist_right_body_to_tail_base_trend | pose geometry | 1 | 0 | 0 |
| dist_right_ear_to_tail_base_delta | kinematics | 1 | 0 | 0 |
| dist_right_ear_to_tail_base_energy | kinematics | 1 | 0 | 0 |
| dist_right_ear_to_tail_base_max | pose geometry | 1 | 0 | 0 |
| dist_right_ear_to_tail_base_mean | pose geometry | 1 | 0 | 0 |
| dist_right_ear_to_tail_base_median | pose geometry | 1 | 0 | 0 |
| dist_right_ear_to_tail_base_norm_delta | kinematics | 1 | 0 | 0 |
| dist_right_ear_to_tail_base_norm_energy | kinematics | 1 | 0 | 0 |
| dist_right_ear_to_tail_base_norm_max | pose geometry | 1 | 0 | 0 |
| dist_right_ear_to_tail_base_norm_mean | pose geometry | 1 | 0 | 0 |
| dist_right_ear_to_tail_base_norm_median | pose geometry | 1 | 0 | 0 |
| dist_right_ear_to_tail_base_norm_p10 | pose geometry | 1 | 0 | 0 |
| dist_right_ear_to_tail_base_norm_p90 | pose geometry | 1 | 0 | 0 |
| dist_right_ear_to_tail_base_norm_periodicity | pose geometry | 1 | 0 | 0 |
| dist_right_ear_to_tail_base_norm_std | kinematics | 1 | 0 | 0 |
| dist_right_ear_to_tail_base_norm_trend | pose geometry | 1 | 0 | 0 |
| dist_right_ear_to_tail_base_p10 | pose geometry | 1 | 0 | 0 |
| dist_right_ear_to_tail_base_p90 | pose geometry | 1 | 0 | 0 |
| dist_right_ear_to_tail_base_periodicity | pose geometry | 1 | 0 | 0 |
| dist_right_ear_to_tail_base_std | kinematics | 1 | 0 | 0 |
| dist_right_ear_to_tail_base_trend | pose geometry | 1 | 0 | 0 |
| ear_left_acceleration_energy | kinematics | 0 | 1 | 1 |
| ear_left_acceleration_max | kinematics | 0 | 1 | 1 |
| ear_left_acceleration_mean | kinematics | 0 | 1 | 1 |
| ear_left_acceleration_median | kinematics | 0 | 1 | 1 |
| ear_left_acceleration_p10 | kinematics | 0 | 1 | 1 |
| ear_left_acceleration_p90 | kinematics | 0 | 1 | 1 |
| ear_left_acceleration_periodicity | kinematics | 0 | 1 | 1 |
| ear_left_acceleration_std | kinematics | 0 | 1 | 1 |
| ear_left_forward_velocity_energy | kinematics | 0 | 1 | 1 |
| ear_left_forward_velocity_max | kinematics | 0 | 1 | 1 |
| ear_left_forward_velocity_mean | kinematics | 0 | 1 | 1 |
| ear_left_forward_velocity_median | kinematics | 0 | 1 | 1 |
| ear_left_forward_velocity_p10 | kinematics | 0 | 1 | 1 |
| ear_left_forward_velocity_p90 | kinematics | 0 | 1 | 1 |
| ear_left_forward_velocity_periodicity | kinematics | 0 | 1 | 1 |
| ear_left_forward_velocity_std | kinematics | 0 | 1 | 1 |
| ear_left_jerk_energy | kinematics | 0 | 1 | 1 |
| ear_left_jerk_max | kinematics | 0 | 1 | 1 |

| | | | | |
|---|---|---|---|---|
| ear_left_jerk_mean | kinematics | 0 | 1 | 1 |
| ear_left_jerk_median | kinematics | 0 | 1 | 1 |
| ear_left_jerk_p10 | kinematics | 0 | 1 | 1 |
| ear_left_jerk_p90 | kinematics | 0 | 1 | 1 |
| ear_left_jerk_periodicity | kinematics | 0 | 1 | 1 |
| ear_left_jerk_std | kinematics | 0 | 1 | 1 |
| ear_left_lateral_velocity_energy | kinematics | 0 | 1 | 1 |
| ear_left_lateral_velocity_max | kinematics | 0 | 1 | 1 |
| ear_left_lateral_velocity_mean | kinematics | 0 | 1 | 1 |
| ear_left_lateral_velocity_median | kinematics | 0 | 1 | 1 |
| ear_left_lateral_velocity_p10 | kinematics | 0 | 1 | 1 |
| ear_left_lateral_velocity_p90 | kinematics | 0 | 1 | 1 |
| ear_left_lateral_velocity_periodicity | kinematics | 0 | 1 | 1 |
| ear_left_lateral_velocity_std | kinematics | 0 | 1 | 1 |
| ear_left_speed_energy | kinematics | 0 | 1 | 1 |
| ear_left_speed_max | kinematics | 0 | 1 | 1 |
| ear_left_speed_mean | kinematics | 0 | 1 | 1 |
| ear_left_speed_median | kinematics | 0 | 1 | 1 |
| ear_left_speed_p10 | kinematics | 0 | 1 | 1 |
| ear_left_speed_p90 | kinematics | 0 | 1 | 1 |
| ear_left_speed_periodicity | kinematics | 0 | 1 | 1 |
| ear_left_speed_std | kinematics | 0 | 1 | 1 |
| ear_left_velocity_x_energy | kinematics | 0 | 1 | 1 |
| ear_left_velocity_x_max | kinematics | 0 | 1 | 1 |
| ear_left_velocity_x_mean | kinematics | 0 | 1 | 1 |
| ear_left_velocity_x_median | kinematics | 0 | 1 | 1 |
| ear_left_velocity_x_p10 | kinematics | 0 | 1 | 1 |
| ear_left_velocity_x_p90 | kinematics | 0 | 1 | 1 |
| ear_left_velocity_x_periodicity | kinematics | 0 | 1 | 1 |
| ear_left_velocity_x_std | kinematics | 0 | 1 | 1 |
| ear_left_velocity_y_energy | kinematics | 0 | 1 | 1 |
| ear_left_velocity_y_max | kinematics | 0 | 1 | 1 |
| ear_left_velocity_y_mean | kinematics | 0 | 1 | 1 |
| ear_left_velocity_y_median | kinematics | 0 | 1 | 1 |
| ear_left_velocity_y_p10 | kinematics | 0 | 1 | 1 |
| ear_left_velocity_y_p90 | kinematics | 0 | 1 | 1 |
| ear_left_velocity_y_periodicity | kinematics | 0 | 1 | 1 |
| ear_left_velocity_y_std | kinematics | 0 | 1 | 1 |
| ear_right_acceleration_energy | kinematics | 0 | 1 | 1 |
| ear_right_acceleration_max | kinematics | 0 | 1 | 1 |
| ear_right_acceleration_mean | kinematics | 0 | 1 | 1 |
| ear_right_acceleration_median | kinematics | 0 | 1 | 1 |
| ear_right_acceleration_p10 | kinematics | 0 | 1 | 1 |
| ear_right_acceleration_p90 | kinematics | 0 | 1 | 1 |

| | | | | |
|---|---|---|---|---|
| ear_right_acceleration_periodicity | kinematics | 0 | 1 | 1 |
| ear_right_acceleration_std | kinematics | 0 | 1 | 1 |
| ear_right_forward_velocity_energy | kinematics | 0 | 1 | 1 |
| ear_right_forward_velocity_max | kinematics | 0 | 1 | 1 |
| ear_right_forward_velocity_mean | kinematics | 0 | 1 | 1 |
| ear_right_forward_velocity_median | kinematics | 0 | 1 | 1 |
| ear_right_forward_velocity_p10 | kinematics | 0 | 1 | 1 |
| ear_right_forward_velocity_p90 | kinematics | 0 | 1 | 1 |
| ear_right_forward_velocity_periodicity | kinematics | 0 | 1 | 1 |
| ear_right_forward_velocity_std | kinematics | 0 | 1 | 1 |
| ear_right_jerk_energy | kinematics | 0 | 1 | 1 |
| ear_right_jerk_max | kinematics | 0 | 1 | 1 |
| ear_right_jerk_mean | kinematics | 0 | 1 | 1 |
| ear_right_jerk_median | kinematics | 0 | 1 | 1 |
| ear_right_jerk_p10 | kinematics | 0 | 1 | 1 |
| ear_right_jerk_p90 | kinematics | 0 | 1 | 1 |
| ear_right_jerk_periodicity | kinematics | 0 | 1 | 1 |
| ear_right_jerk_std | kinematics | 0 | 1 | 1 |
| ear_right_lateral_velocity_energy | kinematics | 0 | 1 | 1 |
| ear_right_lateral_velocity_max | kinematics | 0 | 1 | 1 |
| ear_right_lateral_velocity_mean | kinematics | 0 | 1 | 1 |
| ear_right_lateral_velocity_median | kinematics | 0 | 1 | 1 |
| ear_right_lateral_velocity_p10 | kinematics | 0 | 1 | 1 |
| ear_right_lateral_velocity_p90 | kinematics | 0 | 1 | 1 |
| ear_right_lateral_velocity_periodicity | kinematics | 0 | 1 | 1 |
| ear_right_lateral_velocity_std | kinematics | 0 | 1 | 1 |
| ear_right_speed_energy | kinematics | 0 | 1 | 1 |
| ear_right_speed_max | kinematics | 0 | 1 | 1 |
| ear_right_speed_mean | kinematics | 0 | 1 | 1 |
| ear_right_speed_median | kinematics | 0 | 1 | 1 |
| ear_right_speed_p10 | kinematics | 0 | 1 | 1 |
| ear_right_speed_p90 | kinematics | 0 | 1 | 1 |
| ear_right_speed_periodicity | kinematics | 0 | 1 | 1 |
| ear_right_speed_std | kinematics | 0 | 1 | 1 |
| ear_right_velocity_x_energy | kinematics | 0 | 1 | 1 |
| ear_right_velocity_x_max | kinematics | 0 | 1 | 1 |
| ear_right_velocity_x_mean | kinematics | 0 | 1 | 1 |
| ear_right_velocity_x_median | kinematics | 0 | 1 | 1 |
| ear_right_velocity_x_p10 | kinematics | 0 | 1 | 1 |
| ear_right_velocity_x_p90 | kinematics | 0 | 1 | 1 |
| ear_right_velocity_x_periodicity | kinematics | 0 | 1 | 1 |
| ear_right_velocity_x_std | kinematics | 0 | 1 | 1 |
| ear_right_velocity_y_energy | kinematics | 0 | 1 | 1 |
| ear_right_velocity_y_max | kinematics | 0 | 1 | 1 |

| | | | | |
|---|---|---|---|---|
| ear_right_velocity_y_mean | kinematics | 0 | 1 | 1 |
| ear_right_velocity_y_median | kinematics | 0 | 1 | 1 |
| ear_right_velocity_y_p10 | kinematics | 0 | 1 | 1 |
| ear_right_velocity_y_p90 | kinematics | 0 | 1 | 1 |
| ear_right_velocity_y_periodicity | kinematics | 0 | 1 | 1 |
| ear_right_velocity_y_std | kinematics | 0 | 1 | 1 |
| ear_spread_energy | kinematics | 1 | 1 | 1 |
| ear_spread_max | pose geometry | 1 | 1 | 1 |
| ear_spread_mean | pose geometry | 1 | 1 | 1 |
| ear_spread_median | pose geometry | 1 | 1 | 1 |
| ear_spread_p10 | pose geometry | 1 | 1 | 1 |
| ear_spread_p90 | pose geometry | 1 | 1 | 1 |
| ear_spread_periodicity | pose geometry | 1 | 1 | 1 |
| ear_spread_std | kinematics | 1 | 1 | 1 |
| flow_directionality_paw_energy | video (motion/flow) | 1 | 1 | 1 |
| flow_directionality_paw_max | video (motion/flow) | 1 | 1 | 1 |
| flow_directionality_paw_mean | video (motion/flow) | 1 | 1 | 1 |
| flow_directionality_paw_median | video (motion/flow) | 1 | 1 | 1 |
| flow_directionality_paw_p10 | video (motion/flow) | 1 | 1 | 1 |
| flow_directionality_paw_p90 | video (motion/flow) | 1 | 1 | 1 |
| flow_directionality_paw_periodicity | video (motion/flow) | 1 | 1 | 1 |
| flow_directionality_paw_std | video (motion/flow) | 1 | 1 | 1 |
| flow_entropy_local_energy | video (motion/flow) | 1 | 1 | 1 |
| flow_entropy_local_max | video (motion/flow) | 1 | 1 | 1 |
| flow_entropy_local_mean | video (motion/flow) | 1 | 1 | 1 |
| flow_entropy_local_median | video (motion/flow) | 1 | 1 | 1 |
| flow_entropy_local_p10 | video (motion/flow) | 1 | 1 | 1 |
| flow_entropy_local_p90 | video (motion/flow) | 1 | 1 | 1 |
| flow_entropy_local_periodicity | video (motion/flow) | 1 | 1 | 1 |
| flow_entropy_local_std | video (motion/flow) | 1 | 1 | 1 |
| flow_mag_near_nose_energy | video (motion/flow) | 1 | 1 | 1 |
| flow_mag_near_nose_max | video (motion/flow) | 1 | 1 | 1 |
| flow_mag_near_nose_mean | video (motion/flow) | 1 | 1 | 1 |
| flow_mag_near_nose_median | video (motion/flow) | 1 | 1 | 1 |
| flow_mag_near_nose_p10 | video (motion/flow) | 1 | 1 | 1 |
| flow_mag_near_nose_p90 | video (motion/flow) | 1 | 1 | 1 |
| flow_mag_near_nose_periodicity | video (motion/flow) | 1 | 1 | 1 |
| flow_mag_near_nose_std | video (motion/flow) | 1 | 1 | 1 |
| flow_mag_near_roi_1_energy | video (motion/flow) | 1 | 1 | 1 |
| flow_mag_near_roi_1_max | video (motion/flow) | 1 | 1 | 1 |
| flow_mag_near_roi_1_mean | video (motion/flow) | 1 | 1 | 1 |
| flow_mag_near_roi_1_median | video (motion/flow) | 1 | 1 | 1 |
| flow_mag_near_roi_1_p10 | video (motion/flow) | 1 | 1 | 1 |
| flow_mag_near_roi_1_p90 | video (motion/flow) | 1 | 1 | 1 |

| | | | | |
|---|---|---|---|---|
| flow_mag_near_roi_1_periodicity | video (motion/flow) | 1 | 1 | 1 |
| flow_mag_near_roi_1_std | video (motion/flow) | 1 | 1 | 1 |
| flow_mag_near_roi_2_energy | video (motion/flow) | 1 | 0 | 0 |
| flow_mag_near_roi_2_max | video (motion/flow) | 1 | 0 | 0 |
| flow_mag_near_roi_2_mean | video (motion/flow) | 1 | 0 | 0 |
| flow_mag_near_roi_2_median | video (motion/flow) | 1 | 0 | 0 |
| flow_mag_near_roi_2_p10 | video (motion/flow) | 1 | 0 | 0 |
| flow_mag_near_roi_2_p90 | video (motion/flow) | 1 | 0 | 0 |
| flow_mag_near_roi_2_periodicity | video (motion/flow) | 1 | 0 | 0 |
| flow_mag_near_roi_2_std | video (motion/flow) | 1 | 0 | 0 |
| flow_mag_near_target_energy | video (motion/flow) | 1 | 1 | 1 |
| flow_mag_near_target_max | video (motion/flow) | 1 | 1 | 1 |
| flow_mag_near_target_mean | video (motion/flow) | 1 | 1 | 1 |
| flow_mag_near_target_median | video (motion/flow) | 1 | 1 | 1 |
| flow_mag_near_target_p10 | video (motion/flow) | 1 | 1 | 1 |
| flow_mag_near_target_p90 | video (motion/flow) | 1 | 1 | 1 |
| flow_mag_near_target_periodicity | video (motion/flow) | 1 | 1 | 1 |
| flow_mag_near_target_std | video (motion/flow) | 1 | 1 | 1 |
| flow_mag_paw_L_energy | video (motion/flow) | 1 | 1 | 1 |
| flow_mag_paw_L_max | video (motion/flow) | 1 | 1 | 1 |
| flow_mag_paw_L_mean | video (motion/flow) | 1 | 1 | 1 |
| flow_mag_paw_L_median | video (motion/flow) | 1 | 1 | 1 |
| flow_mag_paw_L_p10 | video (motion/flow) | 1 | 1 | 1 |
| flow_mag_paw_L_p90 | video (motion/flow) | 1 | 1 | 1 |
| flow_mag_paw_L_periodicity | video (motion/flow) | 1 | 1 | 1 |
| flow_mag_paw_L_std | video (motion/flow) | 1 | 1 | 1 |
| flow_mag_paw_R_energy | video (motion/flow) | 1 | 1 | 1 |
| flow_mag_paw_R_max | video (motion/flow) | 1 | 1 | 1 |
| flow_mag_paw_R_mean | video (motion/flow) | 1 | 1 | 1 |
| flow_mag_paw_R_median | video (motion/flow) | 1 | 1 | 1 |
| flow_mag_paw_R_p10 | video (motion/flow) | 1 | 1 | 1 |
| flow_mag_paw_R_p90 | video (motion/flow) | 1 | 1 | 1 |
| flow_mag_paw_R_periodicity | video (motion/flow) | 1 | 1 | 1 |
| flow_mag_paw_R_std | video (motion/flow) | 1 | 1 | 1 |
| forepaw_autocorr_peak_energy | kinematics | 1 | 1 | 1 |
| forepaw_autocorr_peak_max | pose geometry | 1 | 1 | 1 |
| forepaw_autocorr_peak_mean | pose geometry | 1 | 1 | 1 |
| forepaw_autocorr_peak_median | pose geometry | 1 | 1 | 1 |
| forepaw_autocorr_peak_p10 | pose geometry | 1 | 1 | 1 |
| forepaw_autocorr_peak_p90 | pose geometry | 1 | 1 | 1 |
| forepaw_autocorr_peak_periodicity | pose geometry | 1 | 1 | 1 |
| forepaw_autocorr_peak_std | kinematics | 1 | 1 | 1 |
| forepaw_centroid_roi_1_axial_abs_energy | context (ROI/target) | 1 | 1 | 1 |
| forepaw_centroid_roi_1_axial_abs_max | context (ROI/target) | 1 | 1 | 1 |

| | | | | |
|---|---|---|---|---|
| forepaw_centroid_roi_1_axial_abs_mean | context (ROI/target) | 1 | 1 | 1 |
| forepaw_centroid_roi_1_axial_abs_median | context (ROI/target) | 1 | 1 | 1 |
| forepaw_centroid_roi_1_axial_abs_p10 | context (ROI/target) | 1 | 1 | 1 |
| forepaw_centroid_roi_1_axial_abs_p90 | context (ROI/target) | 1 | 1 | 1 |
| forepaw_centroid_roi_1_axial_abs_periodicity | context (ROI/target) | 1 | 1 | 1 |
| forepaw_centroid_roi_1_axial_abs_std | context (ROI/target) | 1 | 1 | 1 |
| forepaw_centroid_roi_1_axial_energy | context (ROI/target) | 1 | 1 | 1 |
| forepaw_centroid_roi_1_axial_max | context (ROI/target) | 1 | 1 | 1 |
| forepaw_centroid_roi_1_axial_mean | context (ROI/target) | 1 | 1 | 1 |
| forepaw_centroid_roi_1_axial_median | context (ROI/target) | 1 | 1 | 1 |
| forepaw_centroid_roi_1_axial_p10 | context (ROI/target) | 1 | 1 | 1 |
| forepaw_centroid_roi_1_axial_p90 | context (ROI/target) | 1 | 1 | 1 |
| forepaw_centroid_roi_1_axial_periodicity | context (ROI/target) | 1 | 1 | 1 |
| forepaw_centroid_roi_1_axial_std | context (ROI/target) | 1 | 1 | 1 |
| forepaw_centroid_roi_1_lateral_abs_energy | context (ROI/target) | 1 | 1 | 1 |
| forepaw_centroid_roi_1_lateral_abs_max | context (ROI/target) | 1 | 1 | 1 |
| forepaw_centroid_roi_1_lateral_abs_mean | context (ROI/target) | 1 | 1 | 1 |
| forepaw_centroid_roi_1_lateral_abs_median | context (ROI/target) | 1 | 1 | 1 |
| forepaw_centroid_roi_1_lateral_abs_p10 | context (ROI/target) | 1 | 1 | 1 |
| forepaw_centroid_roi_1_lateral_abs_p90 | context (ROI/target) | 1 | 1 | 1 |
| forepaw_centroid_roi_1_lateral_abs_periodicity | context (ROI/target) | 1 | 1 | 1 |
| forepaw_centroid_roi_1_lateral_abs_std | context (ROI/target) | 1 | 1 | 1 |
| forepaw_centroid_roi_1_lateral_energy | context (ROI/target) | 1 | 1 | 1 |
| forepaw_centroid_roi_1_lateral_max | context (ROI/target) | 1 | 1 | 1 |
| forepaw_centroid_roi_1_lateral_mean | context (ROI/target) | 1 | 1 | 1 |
| forepaw_centroid_roi_1_lateral_median | context (ROI/target) | 1 | 1 | 1 |
| forepaw_centroid_roi_1_lateral_p10 | context (ROI/target) | 1 | 1 | 1 |
| forepaw_centroid_roi_1_lateral_p90 | context (ROI/target) | 1 | 1 | 1 |
| forepaw_centroid_roi_1_lateral_periodicity | context (ROI/target) | 1 | 1 | 1 |
| forepaw_centroid_roi_1_lateral_std | context (ROI/target) | 1 | 1 | 1 |
| forepaw_centroid_roi_2_axial_abs_energy | context (ROI/target) | 1 | 0 | 0 |
| forepaw_centroid_roi_2_axial_abs_max | context (ROI/target) | 1 | 0 | 0 |
| forepaw_centroid_roi_2_axial_abs_mean | context (ROI/target) | 1 | 0 | 0 |
| forepaw_centroid_roi_2_axial_abs_median | context (ROI/target) | 1 | 0 | 0 |
| forepaw_centroid_roi_2_axial_abs_p10 | context (ROI/target) | 1 | 0 | 0 |
| forepaw_centroid_roi_2_axial_abs_p90 | context (ROI/target) | 1 | 0 | 0 |
| forepaw_centroid_roi_2_axial_abs_periodicity | context (ROI/target) | 1 | 0 | 0 |
| forepaw_centroid_roi_2_axial_abs_std | context (ROI/target) | 1 | 0 | 0 |
| forepaw_centroid_roi_2_axial_energy | context (ROI/target) | 1 | 0 | 0 |
| forepaw_centroid_roi_2_axial_max | context (ROI/target) | 1 | 0 | 0 |
| forepaw_centroid_roi_2_axial_mean | context (ROI/target) | 1 | 0 | 0 |
| forepaw_centroid_roi_2_axial_median | context (ROI/target) | 1 | 0 | 0 |
| forepaw_centroid_roi_2_axial_p10 | context (ROI/target) | 1 | 0 | 0 |
| forepaw_centroid_roi_2_axial_p90 | context (ROI/target) | 1 | 0 | 0 |

| | | | | |
|---|---|---|---|---|
| forepaw_centroid_roi_2_axial_periodicity | context (ROI/target) | 1 | 0 | 0 |
| forepaw_centroid_roi_2_axial_std | context (ROI/target) | 1 | 0 | 0 |
| forepaw_centroid_roi_2_lateral_abs_energy | context (ROI/target) | 1 | 0 | 0 |
| forepaw_centroid_roi_2_lateral_abs_max | context (ROI/target) | 1 | 0 | 0 |
| forepaw_centroid_roi_2_lateral_abs_mean | context (ROI/target) | 1 | 0 | 0 |
| forepaw_centroid_roi_2_lateral_abs_median | context (ROI/target) | 1 | 0 | 0 |
| forepaw_centroid_roi_2_lateral_abs_p10 | context (ROI/target) | 1 | 0 | 0 |
| forepaw_centroid_roi_2_lateral_abs_p90 | context (ROI/target) | 1 | 0 | 0 |
| forepaw_centroid_roi_2_lateral_abs_periodicity | context (ROI/target) | 1 | 0 | 0 |
| forepaw_centroid_roi_2_lateral_abs_std | context (ROI/target) | 1 | 0 | 0 |
| forepaw_centroid_roi_2_lateral_energy | context (ROI/target) | 1 | 0 | 0 |
| forepaw_centroid_roi_2_lateral_max | context (ROI/target) | 1 | 0 | 0 |
| forepaw_centroid_roi_2_lateral_mean | context (ROI/target) | 1 | 0 | 0 |
| forepaw_centroid_roi_2_lateral_median | context (ROI/target) | 1 | 0 | 0 |
| forepaw_centroid_roi_2_lateral_p10 | context (ROI/target) | 1 | 0 | 0 |
| forepaw_centroid_roi_2_lateral_p90 | context (ROI/target) | 1 | 0 | 0 |
| forepaw_centroid_roi_2_lateral_periodicity | context (ROI/target) | 1 | 0 | 0 |
| forepaw_centroid_roi_2_lateral_std | context (ROI/target) | 1 | 0 | 0 |
| forepaw_centroid_to_roi_1_corner_dist_delta | context (ROI/target) | 1 | 1 | 1 |
| forepaw_centroid_to_roi_1_corner_dist_energy | context (ROI/target) | 1 | 1 | 1 |
| forepaw_centroid_to_roi_1_corner_dist_max | context (ROI/target) | 1 | 1 | 1 |
| forepaw_centroid_to_roi_1_corner_dist_mean | context (ROI/target) | 1 | 1 | 1 |
| forepaw_centroid_to_roi_1_corner_dist_median | context (ROI/target) | 1 | 1 | 1 |
| forepaw_centroid_to_roi_1_corner_dist_p10 | context (ROI/target) | 1 | 1 | 1 |
| forepaw_centroid_to_roi_1_corner_dist_p90 | context (ROI/target) | 1 | 1 | 1 |
| forepaw_centroid_to_roi_1_corner_dist_periodicity | context (ROI/target) | 1 | 1 | 1 |
| forepaw_centroid_to_roi_1_corner_dist_std | context (ROI/target) | 1 | 1 | 1 |
| forepaw_centroid_to_roi_1_corner_dist_trend | context (ROI/target) | 1 | 1 | 1 |
| forepaw_centroid_to_roi_1_dist_delta | context (ROI/target) | 1 | 1 | 1 |
| forepaw_centroid_to_roi_1_dist_energy | context (ROI/target) | 1 | 1 | 1 |
| forepaw_centroid_to_roi_1_dist_max | context (ROI/target) | 1 | 1 | 1 |
| forepaw_centroid_to_roi_1_dist_mean | context (ROI/target) | 1 | 1 | 1 |
| forepaw_centroid_to_roi_1_dist_median | context (ROI/target) | 1 | 1 | 1 |
| forepaw_centroid_to_roi_1_dist_p10 | context (ROI/target) | 1 | 1 | 1 |
| forepaw_centroid_to_roi_1_dist_p90 | context (ROI/target) | 1 | 1 | 1 |
| forepaw_centroid_to_roi_1_dist_periodicity | context (ROI/target) | 1 | 1 | 1 |
| forepaw_centroid_to_roi_1_dist_std | context (ROI/target) | 1 | 1 | 1 |
| forepaw_centroid_to_roi_1_dist_trend | context (ROI/target) | 1 | 1 | 1 |
| forepaw_centroid_to_roi_1_edge_dist_delta | context (ROI/target) | 1 | 1 | 1 |
| forepaw_centroid_to_roi_1_edge_dist_energy | context (ROI/target) | 1 | 1 | 1 |
| forepaw_centroid_to_roi_1_edge_dist_max | context (ROI/target) | 1 | 1 | 1 |
| forepaw_centroid_to_roi_1_edge_dist_mean | context (ROI/target) | 1 | 1 | 1 |
| forepaw_centroid_to_roi_1_edge_dist_median | context (ROI/target) | 1 | 1 | 1 |
| forepaw_centroid_to_roi_1_edge_dist_p10 | context (ROI/target) | 1 | 1 | 1 |

| | | | | |
|---|---|---|---|---|
| forepaw_centroid_to_roi_1_edge_dist_p90 | context (ROI/target) | 1 | 1 | 1 |
| forepaw_centroid_to_roi_1_edge_dist_periodicity | context (ROI/target) | 1 | 1 | 1 |
| forepaw_centroid_to_roi_1_edge_dist_std | context (ROI/target) | 1 | 1 | 1 |
| forepaw_centroid_to_roi_1_edge_dist_trend | context (ROI/target) | 1 | 1 | 1 |
| forepaw_centroid_to_roi_1_signed_dist_delta | context (ROI/target) | 1 | 1 | 1 |
| forepaw_centroid_to_roi_1_signed_dist_energy | context (ROI/target) | 1 | 1 | 1 |
| forepaw_centroid_to_roi_1_signed_dist_max | context (ROI/target) | 1 | 1 | 1 |
| forepaw_centroid_to_roi_1_signed_dist_mean | context (ROI/target) | 1 | 1 | 1 |
| forepaw_centroid_to_roi_1_signed_dist_median | context (ROI/target) | 1 | 1 | 1 |
| forepaw_centroid_to_roi_1_signed_dist_p10 | context (ROI/target) | 1 | 1 | 1 |
| forepaw_centroid_to_roi_1_signed_dist_p90 | context (ROI/target) | 1 | 1 | 1 |
| forepaw_centroid_to_roi_1_signed_dist_periodicity | context (ROI/target) | 1 | 1 | 1 |
| forepaw_centroid_to_roi_1_signed_dist_std | context (ROI/target) | 1 | 1 | 1 |
| forepaw_centroid_to_roi_1_signed_dist_trend | context (ROI/target) | 1 | 1 | 1 |
| forepaw_centroid_to_roi_2_corner_dist_delta | context (ROI/target) | 1 | 0 | 0 |
| forepaw_centroid_to_roi_2_corner_dist_energy | context (ROI/target) | 1 | 0 | 0 |
| forepaw_centroid_to_roi_2_corner_dist_max | context (ROI/target) | 1 | 0 | 0 |
| forepaw_centroid_to_roi_2_corner_dist_mean | context (ROI/target) | 1 | 0 | 0 |
| forepaw_centroid_to_roi_2_corner_dist_median | context (ROI/target) | 1 | 0 | 0 |
| forepaw_centroid_to_roi_2_corner_dist_p10 | context (ROI/target) | 1 | 0 | 0 |
| forepaw_centroid_to_roi_2_corner_dist_p90 | context (ROI/target) | 1 | 0 | 0 |
| forepaw_centroid_to_roi_2_corner_dist_periodicity | context (ROI/target) | 1 | 0 | 0 |
| forepaw_centroid_to_roi_2_corner_dist_std | context (ROI/target) | 1 | 0 | 0 |
| forepaw_centroid_to_roi_2_corner_dist_trend | context (ROI/target) | 1 | 0 | 0 |
| forepaw_centroid_to_roi_2_dist_delta | context (ROI/target) | 1 | 0 | 0 |
| forepaw_centroid_to_roi_2_dist_energy | context (ROI/target) | 1 | 0 | 0 |
| forepaw_centroid_to_roi_2_dist_max | context (ROI/target) | 1 | 0 | 0 |
| forepaw_centroid_to_roi_2_dist_mean | context (ROI/target) | 1 | 0 | 0 |
| forepaw_centroid_to_roi_2_dist_median | context (ROI/target) | 1 | 0 | 0 |
| forepaw_centroid_to_roi_2_dist_p10 | context (ROI/target) | 1 | 0 | 0 |
| forepaw_centroid_to_roi_2_dist_p90 | context (ROI/target) | 1 | 0 | 0 |
| forepaw_centroid_to_roi_2_dist_periodicity | context (ROI/target) | 1 | 0 | 0 |
| forepaw_centroid_to_roi_2_dist_std | context (ROI/target) | 1 | 0 | 0 |
| forepaw_centroid_to_roi_2_dist_trend | context (ROI/target) | 1 | 0 | 0 |
| forepaw_centroid_to_roi_2_edge_dist_delta | context (ROI/target) | 1 | 0 | 0 |
| forepaw_centroid_to_roi_2_edge_dist_energy | context (ROI/target) | 1 | 0 | 0 |
| forepaw_centroid_to_roi_2_edge_dist_max | context (ROI/target) | 1 | 0 | 0 |
| forepaw_centroid_to_roi_2_edge_dist_mean | context (ROI/target) | 1 | 0 | 0 |
| forepaw_centroid_to_roi_2_edge_dist_median | context (ROI/target) | 1 | 0 | 0 |
| forepaw_centroid_to_roi_2_edge_dist_p10 | context (ROI/target) | 1 | 0 | 0 |
| forepaw_centroid_to_roi_2_edge_dist_p90 | context (ROI/target) | 1 | 0 | 0 |
| forepaw_centroid_to_roi_2_edge_dist_periodicity | context (ROI/target) | 1 | 0 | 0 |
| forepaw_centroid_to_roi_2_edge_dist_std | context (ROI/target) | 1 | 0 | 0 |
| forepaw_centroid_to_roi_2_edge_dist_trend | context (ROI/target) | 1 | 0 | 0 |

| | | | | |
|---|---|---|---|---|
| forepaw_centroid_to_roi_2_signed_dist_delta | context (ROI/target) | 1 | 0 | 0 |
| forepaw_centroid_to_roi_2_signed_dist_energy | context (ROI/target) | 1 | 0 | 0 |
| forepaw_centroid_to_roi_2_signed_dist_max | context (ROI/target) | 1 | 0 | 0 |
| forepaw_centroid_to_roi_2_signed_dist_mean | context (ROI/target) | 1 | 0 | 0 |
| forepaw_centroid_to_roi_2_signed_dist_median | context (ROI/target) | 1 | 0 | 0 |
| forepaw_centroid_to_roi_2_signed_dist_p10 | context (ROI/target) | 1 | 0 | 0 |
| forepaw_centroid_to_roi_2_signed_dist_p90 | context (ROI/target) | 1 | 0 | 0 |
| forepaw_centroid_to_roi_2_signed_dist_periodicity | context (ROI/target) | 1 | 0 | 0 |
| forepaw_centroid_to_roi_2_signed_dist_std | context (ROI/target) | 1 | 0 | 0 |
| forepaw_centroid_to_roi_2_signed_dist_trend | context (ROI/target) | 1 | 0 | 0 |
| forepaw_centroid_to_target_dist_delta | context (ROI/target) | 1 | 1 | 1 |
| forepaw_centroid_to_target_dist_energy | context (ROI/target) | 1 | 1 | 1 |
| forepaw_centroid_to_target_dist_max | context (ROI/target) | 1 | 1 | 1 |
| forepaw_centroid_to_target_dist_mean | context (ROI/target) | 1 | 1 | 1 |
| forepaw_centroid_to_target_dist_median | context (ROI/target) | 1 | 1 | 1 |
| forepaw_centroid_to_target_dist_p10 | context (ROI/target) | 1 | 1 | 1 |
| forepaw_centroid_to_target_dist_p90 | context (ROI/target) | 1 | 1 | 1 |
| forepaw_centroid_to_target_dist_periodicity | context (ROI/target) | 1 | 1 | 1 |
| forepaw_centroid_to_target_dist_std | context (ROI/target) | 1 | 1 | 1 |
| forepaw_centroid_to_target_dist_trend | context (ROI/target) | 1 | 1 | 1 |
| forepaw_movement_frequency_energy | kinematics | 1 | 1 | 1 |
| forepaw_movement_frequency_max | kinematics | 1 | 1 | 1 |
| forepaw_movement_frequency_mean | kinematics | 1 | 1 | 1 |
| forepaw_movement_frequency_median | kinematics | 1 | 1 | 1 |
| forepaw_movement_frequency_p10 | kinematics | 1 | 1 | 1 |
| forepaw_movement_frequency_p90 | kinematics | 1 | 1 | 1 |
| forepaw_movement_frequency_periodicity | kinematics | 1 | 1 | 1 |
| forepaw_movement_frequency_std | kinematics | 1 | 1 | 1 |
| forepaw_oscillation_power_energy | kinematics | 1 | 1 | 1 |
| forepaw_oscillation_power_max | pose geometry | 1 | 1 | 1 |
| forepaw_oscillation_power_mean | pose geometry | 1 | 1 | 1 |
| forepaw_oscillation_power_median | pose geometry | 1 | 1 | 1 |
| forepaw_oscillation_power_p10 | pose geometry | 1 | 1 | 1 |
| forepaw_oscillation_power_p90 | pose geometry | 1 | 1 | 1 |
| forepaw_oscillation_power_periodicity | pose geometry | 1 | 1 | 1 |
| forepaw_oscillation_power_std | kinematics | 1 | 1 | 1 |
| forepaw_speed_energy | kinematics | 1 | 1 | 1 |
| forepaw_speed_max | kinematics | 1 | 1 | 1 |
| forepaw_speed_mean | kinematics | 1 | 1 | 1 |
| forepaw_speed_median | kinematics | 1 | 1 | 1 |
| forepaw_speed_p10 | kinematics | 1 | 1 | 1 |
| forepaw_speed_p90 | kinematics | 1 | 1 | 1 |
| forepaw_speed_periodicity | kinematics | 1 | 1 | 1 |
| forepaw_speed_std | kinematics | 1 | 1 | 1 |

| | | | | |
|---|---|---|---|---|
| forepaw_vertical_velocity_energy | kinematics | 1 | 1 | 1 |
| forepaw_vertical_velocity_max | kinematics | 1 | 1 | 1 |
| forepaw_vertical_velocity_mean | kinematics | 1 | 1 | 1 |
| forepaw_vertical_velocity_median | kinematics | 1 | 1 | 1 |
| forepaw_vertical_velocity_p10 | kinematics | 1 | 1 | 1 |
| forepaw_vertical_velocity_p90 | kinematics | 1 | 1 | 1 |
| forepaw_vertical_velocity_periodicity | kinematics | 1 | 1 | 1 |
| forepaw_vertical_velocity_std | kinematics | 1 | 1 | 1 |
| front_left_paw_acceleration_energy | kinematics | 0 | 0 | 0 |
| front_left_paw_acceleration_max | kinematics | 0 | 0 | 0 |
| front_left_paw_acceleration_mean | kinematics | 0 | 0 | 0 |
| front_left_paw_acceleration_median | kinematics | 0 | 0 | 0 |
| front_left_paw_acceleration_p10 | kinematics | 0 | 0 | 0 |
| front_left_paw_acceleration_p90 | kinematics | 0 | 0 | 0 |
| front_left_paw_acceleration_periodicity | kinematics | 0 | 0 | 0 |
| front_left_paw_acceleration_std | kinematics | 0 | 0 | 0 |
| front_left_paw_forward_velocity_energy | kinematics | 0 | 0 | 0 |
| front_left_paw_forward_velocity_max | kinematics | 0 | 0 | 0 |
| front_left_paw_forward_velocity_mean | kinematics | 0 | 0 | 0 |
| front_left_paw_forward_velocity_median | kinematics | 0 | 0 | 0 |
| front_left_paw_forward_velocity_p10 | kinematics | 0 | 0 | 0 |
| front_left_paw_forward_velocity_p90 | kinematics | 0 | 0 | 0 |
| front_left_paw_forward_velocity_periodicity | kinematics | 0 | 0 | 0 |
| front_left_paw_forward_velocity_std | kinematics | 0 | 0 | 0 |
| front_left_paw_jerk_energy | kinematics | 0 | 0 | 0 |
| front_left_paw_jerk_max | kinematics | 0 | 0 | 0 |
| front_left_paw_jerk_mean | kinematics | 0 | 0 | 0 |
| front_left_paw_jerk_median | kinematics | 0 | 0 | 0 |
| front_left_paw_jerk_p10 | kinematics | 0 | 0 | 0 |
| front_left_paw_jerk_p90 | kinematics | 0 | 0 | 0 |
| front_left_paw_jerk_periodicity | kinematics | 0 | 0 | 0 |
| front_left_paw_jerk_std | kinematics | 0 | 0 | 0 |
| front_left_paw_lateral_velocity_energy | kinematics | 0 | 0 | 0 |
| front_left_paw_lateral_velocity_max | kinematics | 0 | 0 | 0 |
| front_left_paw_lateral_velocity_mean | kinematics | 0 | 0 | 0 |
| front_left_paw_lateral_velocity_median | kinematics | 0 | 0 | 0 |
| front_left_paw_lateral_velocity_p10 | kinematics | 0 | 0 | 0 |
| front_left_paw_lateral_velocity_p90 | kinematics | 0 | 0 | 0 |
| front_left_paw_lateral_velocity_periodicity | kinematics | 0 | 0 | 0 |
| front_left_paw_lateral_velocity_std | kinematics | 0 | 0 | 0 |
| front_left_paw_speed_energy | kinematics | 0 | 0 | 0 |
| front_left_paw_speed_max | kinematics | 0 | 0 | 0 |
| front_left_paw_speed_mean | kinematics | 0 | 0 | 0 |
| front_left_paw_speed_median | kinematics | 0 | 0 | 0 |

| | | | | |
|---|---|---|---|---|
| front_left_paw_speed_p10 | kinematics | 0 | 0 | 0 |
| front_left_paw_speed_p90 | kinematics | 0 | 0 | 0 |
| front_left_paw_speed_periodicity | kinematics | 0 | 0 | 0 |
| front_left_paw_speed_std | kinematics | 0 | 0 | 0 |
| front_left_paw_velocity_x_energy | kinematics | 0 | 0 | 0 |
| front_left_paw_velocity_x_max | kinematics | 0 | 0 | 0 |
| front_left_paw_velocity_x_mean | kinematics | 0 | 0 | 0 |
| front_left_paw_velocity_x_median | kinematics | 0 | 0 | 0 |
| front_left_paw_velocity_x_p10 | kinematics | 0 | 0 | 0 |
| front_left_paw_velocity_x_p90 | kinematics | 0 | 0 | 0 |
| front_left_paw_velocity_x_periodicity | kinematics | 0 | 0 | 0 |
| front_left_paw_velocity_x_std | kinematics | 0 | 0 | 0 |
| front_left_paw_velocity_y_energy | kinematics | 0 | 0 | 0 |
| front_left_paw_velocity_y_max | kinematics | 0 | 0 | 0 |
| front_left_paw_velocity_y_mean | kinematics | 0 | 0 | 0 |
| front_left_paw_velocity_y_median | kinematics | 0 | 0 | 0 |
| front_left_paw_velocity_y_p10 | kinematics | 0 | 0 | 0 |
| front_left_paw_velocity_y_p90 | kinematics | 0 | 0 | 0 |
| front_left_paw_velocity_y_periodicity | kinematics | 0 | 0 | 0 |
| front_left_paw_velocity_y_std | kinematics | 0 | 0 | 0 |
| front_right_paw_acceleration_energy | kinematics | 0 | 0 | 0 |
| front_right_paw_acceleration_max | kinematics | 0 | 0 | 0 |
| front_right_paw_acceleration_mean | kinematics | 0 | 0 | 0 |
| front_right_paw_acceleration_median | kinematics | 0 | 0 | 0 |
| front_right_paw_acceleration_p10 | kinematics | 0 | 0 | 0 |
| front_right_paw_acceleration_p90 | kinematics | 0 | 0 | 0 |
| front_right_paw_acceleration_periodicity | kinematics | 0 | 0 | 0 |
| front_right_paw_acceleration_std | kinematics | 0 | 0 | 0 |
| front_right_paw_forward_velocity_energy | kinematics | 0 | 0 | 0 |
| front_right_paw_forward_velocity_max | kinematics | 0 | 0 | 0 |
| front_right_paw_forward_velocity_mean | kinematics | 0 | 0 | 0 |
| front_right_paw_forward_velocity_median | kinematics | 0 | 0 | 0 |
| front_right_paw_forward_velocity_p10 | kinematics | 0 | 0 | 0 |
| front_right_paw_forward_velocity_p90 | kinematics | 0 | 0 | 0 |
| front_right_paw_forward_velocity_periodicity | kinematics | 0 | 0 | 0 |
| front_right_paw_forward_velocity_std | kinematics | 0 | 0 | 0 |
| front_right_paw_jerk_energy | kinematics | 0 | 0 | 0 |
| front_right_paw_jerk_max | kinematics | 0 | 0 | 0 |
| front_right_paw_jerk_mean | kinematics | 0 | 0 | 0 |
| front_right_paw_jerk_median | kinematics | 0 | 0 | 0 |
| front_right_paw_jerk_p10 | kinematics | 0 | 0 | 0 |
| front_right_paw_jerk_p90 | kinematics | 0 | 0 | 0 |
| front_right_paw_jerk_periodicity | kinematics | 0 | 0 | 0 |
| front_right_paw_jerk_std | kinematics | 0 | 0 | 0 |

| | | | | |
|---|---|---|---|---|
| front_right_paw_lateral_velocity_energy | kinematics | 0 | 0 | 0 |
| front_right_paw_lateral_velocity_max | kinematics | 0 | 0 | 0 |
| front_right_paw_lateral_velocity_mean | kinematics | 0 | 0 | 0 |
| front_right_paw_lateral_velocity_median | kinematics | 0 | 0 | 0 |
| front_right_paw_lateral_velocity_p10 | kinematics | 0 | 0 | 0 |
| front_right_paw_lateral_velocity_p90 | kinematics | 0 | 0 | 0 |
| front_right_paw_lateral_velocity_periodicity | kinematics | 0 | 0 | 0 |
| front_right_paw_lateral_velocity_std | kinematics | 0 | 0 | 0 |
| front_right_paw_speed_energy | kinematics | 0 | 0 | 0 |
| front_right_paw_speed_max | kinematics | 0 | 0 | 0 |
| front_right_paw_speed_mean | kinematics | 0 | 0 | 0 |
| front_right_paw_speed_median | kinematics | 0 | 0 | 0 |
| front_right_paw_speed_p10 | kinematics | 0 | 0 | 0 |
| front_right_paw_speed_p90 | kinematics | 0 | 0 | 0 |
| front_right_paw_speed_periodicity | kinematics | 0 | 0 | 0 |
| front_right_paw_speed_std | kinematics | 0 | 0 | 0 |
| front_right_paw_velocity_x_energy | kinematics | 0 | 0 | 0 |
| front_right_paw_velocity_x_max | kinematics | 0 | 0 | 0 |
| front_right_paw_velocity_x_mean | kinematics | 0 | 0 | 0 |
| front_right_paw_velocity_x_median | kinematics | 0 | 0 | 0 |
| front_right_paw_velocity_x_p10 | kinematics | 0 | 0 | 0 |
| front_right_paw_velocity_x_p90 | kinematics | 0 | 0 | 0 |
| front_right_paw_velocity_x_periodicity | kinematics | 0 | 0 | 0 |
| front_right_paw_velocity_x_std | kinematics | 0 | 0 | 0 |
| front_right_paw_velocity_y_energy | kinematics | 0 | 0 | 0 |
| front_right_paw_velocity_y_max | kinematics | 0 | 0 | 0 |
| front_right_paw_velocity_y_mean | kinematics | 0 | 0 | 0 |
| front_right_paw_velocity_y_median | kinematics | 0 | 0 | 0 |
| front_right_paw_velocity_y_p10 | kinematics | 0 | 0 | 0 |
| front_right_paw_velocity_y_p90 | kinematics | 0 | 0 | 0 |
| front_right_paw_velocity_y_periodicity | kinematics | 0 | 0 | 0 |
| front_right_paw_velocity_y_std | kinematics | 0 | 0 | 0 |
| head_angle_to_roi_1_delta | context (ROI/target) | 1 | 1 | 1 |
| head_angle_to_roi_1_energy | context (ROI/target) | 1 | 1 | 1 |
| head_angle_to_roi_1_max | context (ROI/target) | 1 | 1 | 1 |
| head_angle_to_roi_1_mean | context (ROI/target) | 1 | 1 | 1 |
| head_angle_to_roi_1_median | context (ROI/target) | 1 | 1 | 1 |
| head_angle_to_roi_1_p10 | context (ROI/target) | 1 | 1 | 1 |
| head_angle_to_roi_1_p90 | context (ROI/target) | 1 | 1 | 1 |
| head_angle_to_roi_1_periodicity | context (ROI/target) | 1 | 1 | 1 |
| head_angle_to_roi_1_std | context (ROI/target) | 1 | 1 | 1 |
| head_angle_to_roi_1_trend | context (ROI/target) | 1 | 1 | 1 |
| head_angle_to_roi_2_delta | context (ROI/target) | 1 | 0 | 0 |
| head_angle_to_roi_2_energy | context (ROI/target) | 1 | 0 | 0 |

| | | | | |
|---|---|---|---|---|
| head_angle_to_roi_2_max | context (ROI/target) | 1 | 0 | 0 |
| head_angle_to_roi_2_mean | context (ROI/target) | 1 | 0 | 0 |
| head_angle_to_roi_2_median | context (ROI/target) | 1 | 0 | 0 |
| head_angle_to_roi_2_p10 | context (ROI/target) | 1 | 0 | 0 |
| head_angle_to_roi_2_p90 | context (ROI/target) | 1 | 0 | 0 |
| head_angle_to_roi_2_periodicity | context (ROI/target) | 1 | 0 | 0 |
| head_angle_to_roi_2_std | context (ROI/target) | 1 | 0 | 0 |
| head_angle_to_roi_2_trend | context (ROI/target) | 1 | 0 | 0 |
| head_angle_to_target_delta | context (ROI/target) | 1 | 1 | 1 |
| head_angle_to_target_energy | context (ROI/target) | 1 | 1 | 1 |
| head_angle_to_target_max | context (ROI/target) | 1 | 1 | 1 |
| head_angle_to_target_mean | context (ROI/target) | 1 | 1 | 1 |
| head_angle_to_target_median | context (ROI/target) | 1 | 1 | 1 |
| head_angle_to_target_p10 | context (ROI/target) | 1 | 1 | 1 |
| head_angle_to_target_p90 | context (ROI/target) | 1 | 1 | 1 |
| head_angle_to_target_periodicity | context (ROI/target) | 1 | 1 | 1 |
| head_angle_to_target_std | context (ROI/target) | 1 | 1 | 1 |
| head_angle_to_target_trend | context (ROI/target) | 1 | 1 | 1 |
| head_angular_velocity_energy | kinematics | 1 | 1 | 1 |
| head_angular_velocity_max | kinematics | 1 | 1 | 1 |
| head_angular_velocity_mean | kinematics | 1 | 1 | 1 |
| head_angular_velocity_median | kinematics | 1 | 1 | 1 |
| head_angular_velocity_p10 | kinematics | 1 | 1 | 1 |
| head_angular_velocity_p90 | kinematics | 1 | 1 | 1 |
| head_angular_velocity_periodicity | kinematics | 1 | 1 | 1 |
| head_angular_velocity_std | kinematics | 1 | 1 | 1 |
| head_direction_angle_delta | kinematics | 1 | 1 | 1 |
| head_direction_angle_energy | kinematics | 1 | 1 | 1 |
| head_direction_angle_max | pose geometry | 1 | 1 | 1 |
| head_direction_angle_mean | pose geometry | 1 | 1 | 1 |
| head_direction_angle_median | pose geometry | 1 | 1 | 1 |
| head_direction_angle_p10 | pose geometry | 1 | 1 | 1 |
| head_direction_angle_p90 | pose geometry | 1 | 1 | 1 |
| head_direction_angle_periodicity | pose geometry | 1 | 1 | 1 |
| head_direction_angle_std | kinematics | 1 | 1 | 1 |
| head_direction_angle_trend | pose geometry | 1 | 1 | 1 |
| head_forward_speed_energy | kinematics | 1 | 1 | 1 |
| head_forward_speed_max | kinematics | 1 | 1 | 1 |
| head_forward_speed_mean | kinematics | 1 | 1 | 1 |
| head_forward_speed_median | kinematics | 1 | 1 | 1 |
| head_forward_speed_p10 | kinematics | 1 | 1 | 1 |
| head_forward_speed_p90 | kinematics | 1 | 1 | 1 |
| head_forward_speed_periodicity | kinematics | 1 | 1 | 1 |
| head_forward_speed_std | kinematics | 1 | 1 | 1 |

| | | | | |
|---|---|---|---|---|
| head_lateral_speed_energy | kinematics | 1 | 1 | 1 |
| head_lateral_speed_max | kinematics | 1 | 1 | 1 |
| head_lateral_speed_mean | kinematics | 1 | 1 | 1 |
| head_lateral_speed_median | kinematics | 1 | 1 | 1 |
| head_lateral_speed_p10 | kinematics | 1 | 1 | 1 |
| head_lateral_speed_p90 | kinematics | 1 | 1 | 1 |
| head_lateral_speed_periodicity | kinematics | 1 | 1 | 1 |
| head_lateral_speed_std | kinematics | 1 | 1 | 1 |
| head_pitch_delta | kinematics | 1 | 1 | 1 |
| head_pitch_energy | kinematics | 1 | 1 | 1 |
| head_pitch_max | pose geometry | 1 | 1 | 1 |
| head_pitch_mean | pose geometry | 1 | 1 | 1 |
| head_pitch_median | pose geometry | 1 | 1 | 1 |
| head_pitch_p10 | pose geometry | 1 | 1 | 1 |
| head_pitch_p90 | pose geometry | 1 | 1 | 1 |
| head_pitch_periodicity | pose geometry | 1 | 1 | 1 |
| head_pitch_std | kinematics | 1 | 1 | 1 |
| head_pitch_trend | pose geometry | 1 | 1 | 1 |
| in_roi_1_body_centroid_energy | context (ROI/target) | 1 | 1 | 1 |
| in_roi_1_body_centroid_max | context (ROI/target) | 1 | 1 | 1 |
| in_roi_1_body_centroid_mean | context (ROI/target) | 1 | 1 | 1 |
| in_roi_1_body_centroid_median | context (ROI/target) | 1 | 1 | 1 |
| in_roi_1_body_centroid_p10 | context (ROI/target) | 1 | 1 | 1 |
| in_roi_1_body_centroid_p90 | context (ROI/target) | 1 | 1 | 1 |
| in_roi_1_body_centroid_periodicity | context (ROI/target) | 1 | 1 | 1 |
| in_roi_1_body_centroid_std | context (ROI/target) | 1 | 1 | 1 |
| in_roi_1_forepaw_centroid_energy | context (ROI/target) | 1 | 1 | 1 |
| in_roi_1_forepaw_centroid_max | context (ROI/target) | 1 | 1 | 1 |
| in_roi_1_forepaw_centroid_mean | context (ROI/target) | 1 | 1 | 1 |
| in_roi_1_forepaw_centroid_median | context (ROI/target) | 1 | 1 | 1 |
| in_roi_1_forepaw_centroid_p10 | context (ROI/target) | 1 | 1 | 1 |
| in_roi_1_forepaw_centroid_p90 | context (ROI/target) | 1 | 1 | 1 |
| in_roi_1_forepaw_centroid_periodicity | context (ROI/target) | 1 | 1 | 1 |
| in_roi_1_forepaw_centroid_std | context (ROI/target) | 1 | 1 | 1 |
| in_roi_1_nose_energy | context (ROI/target) | 1 | 1 | 1 |
| in_roi_1_nose_max | context (ROI/target) | 1 | 1 | 1 |
| in_roi_1_nose_mean | context (ROI/target) | 1 | 1 | 1 |
| in_roi_1_nose_median | context (ROI/target) | 1 | 1 | 1 |
| in_roi_1_nose_p10 | context (ROI/target) | 1 | 1 | 1 |
| in_roi_1_nose_p90 | context (ROI/target) | 1 | 1 | 1 |
| in_roi_1_nose_periodicity | context (ROI/target) | 1 | 1 | 1 |
| in_roi_1_nose_std | context (ROI/target) | 1 | 1 | 1 |
| in_roi_2_body_centroid_energy | context (ROI/target) | 1 | 0 | 0 |
| in_roi_2_body_centroid_max | context (ROI/target) | 1 | 0 | 0 |

| | | | | |
|---|---|---|---|---|
| in_roi_2_body_centroid_mean | context (ROI/target) | 1 | 0 | 0 |
| in_roi_2_body_centroid_median | context (ROI/target) | 1 | 0 | 0 |
| in_roi_2_body_centroid_p10 | context (ROI/target) | 1 | 0 | 0 |
| in_roi_2_body_centroid_p90 | context (ROI/target) | 1 | 0 | 0 |
| in_roi_2_body_centroid_periodicity | context (ROI/target) | 1 | 0 | 0 |
| in_roi_2_body_centroid_std | context (ROI/target) | 1 | 0 | 0 |
| in_roi_2_forepaw_centroid_energy | context (ROI/target) | 1 | 0 | 0 |
| in_roi_2_forepaw_centroid_max | context (ROI/target) | 1 | 0 | 0 |
| in_roi_2_forepaw_centroid_mean | context (ROI/target) | 1 | 0 | 0 |
| in_roi_2_forepaw_centroid_median | context (ROI/target) | 1 | 0 | 0 |
| in_roi_2_forepaw_centroid_p10 | context (ROI/target) | 1 | 0 | 0 |
| in_roi_2_forepaw_centroid_p90 | context (ROI/target) | 1 | 0 | 0 |
| in_roi_2_forepaw_centroid_periodicity | context (ROI/target) | 1 | 0 | 0 |
| in_roi_2_forepaw_centroid_std | context (ROI/target) | 1 | 0 | 0 |
| in_roi_2_nose_energy | context (ROI/target) | 1 | 0 | 0 |
| in_roi_2_nose_max | context (ROI/target) | 1 | 0 | 0 |
| in_roi_2_nose_mean | context (ROI/target) | 1 | 0 | 0 |
| in_roi_2_nose_median | context (ROI/target) | 1 | 0 | 0 |
| in_roi_2_nose_p10 | context (ROI/target) | 1 | 0 | 0 |
| in_roi_2_nose_p90 | context (ROI/target) | 1 | 0 | 0 |
| in_roi_2_nose_periodicity | context (ROI/target) | 1 | 0 | 0 |
| in_roi_2_nose_std | context (ROI/target) | 1 | 0 | 0 |
| joint_angle_forelimb_spread_delta | kinematics | 0 | 0 | 0 |
| joint_angle_forelimb_spread_energy | kinematics | 0 | 0 | 0 |
| joint_angle_forelimb_spread_max | pose geometry | 0 | 0 | 0 |
| joint_angle_forelimb_spread_mean | pose geometry | 0 | 0 | 0 |
| joint_angle_forelimb_spread_median | pose geometry | 0 | 0 | 0 |
| joint_angle_forelimb_spread_p10 | pose geometry | 0 | 0 | 0 |
| joint_angle_forelimb_spread_p90 | pose geometry | 0 | 0 | 0 |
| joint_angle_forelimb_spread_periodicity | pose geometry | 0 | 0 | 0 |
| joint_angle_forelimb_spread_std | kinematics | 0 | 0 | 0 |
| joint_angle_forelimb_spread_trend | pose geometry | 0 | 0 | 0 |
| joint_angle_head_neck_angle_delta | kinematics | 1 | 1 | 1 |
| joint_angle_head_neck_angle_energy | kinematics | 1 | 1 | 1 |
| joint_angle_head_neck_angle_max | pose geometry | 1 | 1 | 1 |
| joint_angle_head_neck_angle_mean | pose geometry | 1 | 1 | 1 |
| joint_angle_head_neck_angle_median | pose geometry | 1 | 1 | 1 |
| joint_angle_head_neck_angle_p10 | pose geometry | 1 | 1 | 1 |
| joint_angle_head_neck_angle_p90 | pose geometry | 1 | 1 | 1 |
| joint_angle_head_neck_angle_periodicity | pose geometry | 1 | 1 | 1 |
| joint_angle_head_neck_angle_std | kinematics | 1 | 1 | 1 |
| joint_angle_head_neck_angle_trend | pose geometry | 1 | 1 | 1 |
| joint_angle_lateral_torso_delta | kinematics | 1 | 1 | 0 |
| joint_angle_lateral_torso_energy | kinematics | 1 | 1 | 0 |

| | | | | |
|---|---|---|---|---|
| joint_angle_lateral_torso_max | pose geometry | 1 | 1 | 0 |
| joint_angle_lateral_torso_mean | pose geometry | 1 | 1 | 0 |
| joint_angle_lateral_torso_median | pose geometry | 1 | 1 | 0 |
| joint_angle_lateral_torso_p10 | pose geometry | 1 | 1 | 0 |
| joint_angle_lateral_torso_p90 | pose geometry | 1 | 1 | 0 |
| joint_angle_lateral_torso_periodicity | pose geometry | 1 | 1 | 0 |
| joint_angle_lateral_torso_std | kinematics | 1 | 1 | 0 |
| joint_angle_lateral_torso_trend | pose geometry | 1 | 1 | 0 |
| joint_angle_spine_flexion_delta | kinematics | 1 | 1 | 1 |
| joint_angle_spine_flexion_energy | kinematics | 1 | 1 | 1 |
| joint_angle_spine_flexion_max | pose geometry | 1 | 1 | 1 |
| joint_angle_spine_flexion_mean | pose geometry | 1 | 1 | 1 |
| joint_angle_spine_flexion_median | pose geometry | 1 | 1 | 1 |
| joint_angle_spine_flexion_p10 | pose geometry | 1 | 1 | 1 |
| joint_angle_spine_flexion_p90 | pose geometry | 1 | 1 | 1 |
| joint_angle_spine_flexion_periodicity | pose geometry | 1 | 1 | 1 |
| joint_angle_spine_flexion_std | kinematics | 1 | 1 | 1 |
| joint_angle_spine_flexion_trend | pose geometry | 1 | 1 | 1 |
| lateral_left_acceleration_energy | kinematics | 0 | 0 | 1 |
| lateral_left_acceleration_max | kinematics | 0 | 0 | 1 |
| lateral_left_acceleration_mean | kinematics | 0 | 0 | 1 |
| lateral_left_acceleration_median | kinematics | 0 | 0 | 1 |
| lateral_left_acceleration_p10 | kinematics | 0 | 0 | 1 |
| lateral_left_acceleration_p90 | kinematics | 0 | 0 | 1 |
| lateral_left_acceleration_periodicity | kinematics | 0 | 0 | 1 |
| lateral_left_acceleration_std | kinematics | 0 | 0 | 1 |
| lateral_left_forward_velocity_energy | kinematics | 0 | 0 | 1 |
| lateral_left_forward_velocity_max | kinematics | 0 | 0 | 1 |
| lateral_left_forward_velocity_mean | kinematics | 0 | 0 | 1 |
| lateral_left_forward_velocity_median | kinematics | 0 | 0 | 1 |
| lateral_left_forward_velocity_p10 | kinematics | 0 | 0 | 1 |
| lateral_left_forward_velocity_p90 | kinematics | 0 | 0 | 1 |
| lateral_left_forward_velocity_periodicity | kinematics | 0 | 0 | 1 |
| lateral_left_forward_velocity_std | kinematics | 0 | 0 | 1 |
| lateral_left_jerk_energy | kinematics | 0 | 0 | 1 |
| lateral_left_jerk_max | kinematics | 0 | 0 | 1 |
| lateral_left_jerk_mean | kinematics | 0 | 0 | 1 |
| lateral_left_jerk_median | kinematics | 0 | 0 | 1 |
| lateral_left_jerk_p10 | kinematics | 0 | 0 | 1 |
| lateral_left_jerk_p90 | kinematics | 0 | 0 | 1 |
| lateral_left_jerk_periodicity | kinematics | 0 | 0 | 1 |
| lateral_left_jerk_std | kinematics | 0 | 0 | 1 |
| lateral_left_lateral_velocity_energy | kinematics | 0 | 0 | 1 |
| lateral_left_lateral_velocity_max | kinematics | 0 | 0 | 1 |

| | | | | |
|---|---|---|---|---|
| lateral_left_lateral_velocity_mean | kinematics | 0 | 0 | 1 |
| lateral_left_lateral_velocity_median | kinematics | 0 | 0 | 1 |
| lateral_left_lateral_velocity_p10 | kinematics | 0 | 0 | 1 |
| lateral_left_lateral_velocity_p90 | kinematics | 0 | 0 | 1 |
| lateral_left_lateral_velocity_periodicity | kinematics | 0 | 0 | 1 |
| lateral_left_lateral_velocity_std | kinematics | 0 | 0 | 1 |
| lateral_left_speed_energy | kinematics | 0 | 0 | 1 |
| lateral_left_speed_max | kinematics | 0 | 0 | 1 |
| lateral_left_speed_mean | kinematics | 0 | 0 | 1 |
| lateral_left_speed_median | kinematics | 0 | 0 | 1 |
| lateral_left_speed_p10 | kinematics | 0 | 0 | 1 |
| lateral_left_speed_p90 | kinematics | 0 | 0 | 1 |
| lateral_left_speed_periodicity | kinematics | 0 | 0 | 1 |
| lateral_left_speed_std | kinematics | 0 | 0 | 1 |
| lateral_left_velocity_x_energy | kinematics | 0 | 0 | 1 |
| lateral_left_velocity_x_max | kinematics | 0 | 0 | 1 |
| lateral_left_velocity_x_mean | kinematics | 0 | 0 | 1 |
| lateral_left_velocity_x_median | kinematics | 0 | 0 | 1 |
| lateral_left_velocity_x_p10 | kinematics | 0 | 0 | 1 |
| lateral_left_velocity_x_p90 | kinematics | 0 | 0 | 1 |
| lateral_left_velocity_x_periodicity | kinematics | 0 | 0 | 1 |
| lateral_left_velocity_x_std | kinematics | 0 | 0 | 1 |
| lateral_left_velocity_y_energy | kinematics | 0 | 0 | 1 |
| lateral_left_velocity_y_max | kinematics | 0 | 0 | 1 |
| lateral_left_velocity_y_mean | kinematics | 0 | 0 | 1 |
| lateral_left_velocity_y_median | kinematics | 0 | 0 | 1 |
| lateral_left_velocity_y_p10 | kinematics | 0 | 0 | 1 |
| lateral_left_velocity_y_p90 | kinematics | 0 | 0 | 1 |
| lateral_left_velocity_y_periodicity | kinematics | 0 | 0 | 1 |
| lateral_left_velocity_y_std | kinematics | 0 | 0 | 1 |
| lateral_right_acceleration_energy | kinematics | 0 | 0 | 1 |
| lateral_right_acceleration_max | kinematics | 0 | 0 | 1 |
| lateral_right_acceleration_mean | kinematics | 0 | 0 | 1 |
| lateral_right_acceleration_median | kinematics | 0 | 0 | 1 |
| lateral_right_acceleration_p10 | kinematics | 0 | 0 | 1 |
| lateral_right_acceleration_p90 | kinematics | 0 | 0 | 1 |
| lateral_right_acceleration_periodicity | kinematics | 0 | 0 | 1 |
| lateral_right_acceleration_std | kinematics | 0 | 0 | 1 |
| lateral_right_forward_velocity_energy | kinematics | 0 | 0 | 1 |
| lateral_right_forward_velocity_max | kinematics | 0 | 0 | 1 |
| lateral_right_forward_velocity_mean | kinematics | 0 | 0 | 1 |
| lateral_right_forward_velocity_median | kinematics | 0 | 0 | 1 |
| lateral_right_forward_velocity_p10 | kinematics | 0 | 0 | 1 |
| lateral_right_forward_velocity_p90 | kinematics | 0 | 0 | 1 |

| | | | | |
|---|---|---|---|---|
| lateral_right_forward_velocity_periodicity | kinematics | 0 | 0 | 1 |
| lateral_right_forward_velocity_std | kinematics | 0 | 0 | 1 |
| lateral_right_jerk_energy | kinematics | 0 | 0 | 1 |
| lateral_right_jerk_max | kinematics | 0 | 0 | 1 |
| lateral_right_jerk_mean | kinematics | 0 | 0 | 1 |
| lateral_right_jerk_median | kinematics | 0 | 0 | 1 |
| lateral_right_jerk_p10 | kinematics | 0 | 0 | 1 |
| lateral_right_jerk_p90 | kinematics | 0 | 0 | 1 |
| lateral_right_jerk_periodicity | kinematics | 0 | 0 | 1 |
| lateral_right_jerk_std | kinematics | 0 | 0 | 1 |
| lateral_right_lateral_velocity_energy | kinematics | 0 | 0 | 1 |
| lateral_right_lateral_velocity_max | kinematics | 0 | 0 | 1 |
| lateral_right_lateral_velocity_mean | kinematics | 0 | 0 | 1 |
| lateral_right_lateral_velocity_median | kinematics | 0 | 0 | 1 |
| lateral_right_lateral_velocity_p10 | kinematics | 0 | 0 | 1 |
| lateral_right_lateral_velocity_p90 | kinematics | 0 | 0 | 1 |
| lateral_right_lateral_velocity_periodicity | kinematics | 0 | 0 | 1 |
| lateral_right_lateral_velocity_std | kinematics | 0 | 0 | 1 |
| lateral_right_speed_energy | kinematics | 0 | 0 | 1 |
| lateral_right_speed_max | kinematics | 0 | 0 | 1 |
| lateral_right_speed_mean | kinematics | 0 | 0 | 1 |
| lateral_right_speed_median | kinematics | 0 | 0 | 1 |
| lateral_right_speed_p10 | kinematics | 0 | 0 | 1 |
| lateral_right_speed_p90 | kinematics | 0 | 0 | 1 |
| lateral_right_speed_periodicity | kinematics | 0 | 0 | 1 |
| lateral_right_speed_std | kinematics | 0 | 0 | 1 |
| lateral_right_velocity_x_energy | kinematics | 0 | 0 | 1 |
| lateral_right_velocity_x_max | kinematics | 0 | 0 | 1 |
| lateral_right_velocity_x_mean | kinematics | 0 | 0 | 1 |
| lateral_right_velocity_x_median | kinematics | 0 | 0 | 1 |
| lateral_right_velocity_x_p10 | kinematics | 0 | 0 | 1 |
| lateral_right_velocity_x_p90 | kinematics | 0 | 0 | 1 |
| lateral_right_velocity_x_periodicity | kinematics | 0 | 0 | 1 |
| lateral_right_velocity_x_std | kinematics | 0 | 0 | 1 |
| lateral_right_velocity_y_energy | kinematics | 0 | 0 | 1 |
| lateral_right_velocity_y_max | kinematics | 0 | 0 | 1 |
| lateral_right_velocity_y_mean | kinematics | 0 | 0 | 1 |
| lateral_right_velocity_y_median | kinematics | 0 | 0 | 1 |
| lateral_right_velocity_y_p10 | kinematics | 0 | 0 | 1 |
| lateral_right_velocity_y_p90 | kinematics | 0 | 0 | 1 |
| lateral_right_velocity_y_periodicity | kinematics | 0 | 0 | 1 |
| lateral_right_velocity_y_std | kinematics | 0 | 0 | 1 |
| left_body_acceleration_energy | kinematics | 1 | 0 | 0 |
| left_body_acceleration_max | kinematics | 1 | 0 | 0 |

| | | | | |
|---|---|---|---|---|
| left_body_acceleration_mean | kinematics | 1 | 0 | 0 |
| left_body_acceleration_median | kinematics | 1 | 0 | 0 |
| left_body_acceleration_p10 | kinematics | 1 | 0 | 0 |
| left_body_acceleration_p90 | kinematics | 1 | 0 | 0 |
| left_body_acceleration_periodicity | kinematics | 1 | 0 | 0 |
| left_body_acceleration_std | kinematics | 1 | 0 | 0 |
| left_body_forward_velocity_energy | kinematics | 1 | 0 | 0 |
| left_body_forward_velocity_max | kinematics | 1 | 0 | 0 |
| left_body_forward_velocity_mean | kinematics | 1 | 0 | 0 |
| left_body_forward_velocity_median | kinematics | 1 | 0 | 0 |
| left_body_forward_velocity_p10 | kinematics | 1 | 0 | 0 |
| left_body_forward_velocity_p90 | kinematics | 1 | 0 | 0 |
| left_body_forward_velocity_periodicity | kinematics | 1 | 0 | 0 |
| left_body_forward_velocity_std | kinematics | 1 | 0 | 0 |
| left_body_jerk_energy | kinematics | 1 | 0 | 0 |
| left_body_jerk_max | kinematics | 1 | 0 | 0 |
| left_body_jerk_mean | kinematics | 1 | 0 | 0 |
| left_body_jerk_median | kinematics | 1 | 0 | 0 |
| left_body_jerk_p10 | kinematics | 1 | 0 | 0 |
| left_body_jerk_p90 | kinematics | 1 | 0 | 0 |
| left_body_jerk_periodicity | kinematics | 1 | 0 | 0 |
| left_body_jerk_std | kinematics | 1 | 0 | 0 |
| left_body_lateral_velocity_energy | kinematics | 1 | 0 | 0 |
| left_body_lateral_velocity_max | kinematics | 1 | 0 | 0 |
| left_body_lateral_velocity_mean | kinematics | 1 | 0 | 0 |
| left_body_lateral_velocity_median | kinematics | 1 | 0 | 0 |
| left_body_lateral_velocity_p10 | kinematics | 1 | 0 | 0 |
| left_body_lateral_velocity_p90 | kinematics | 1 | 0 | 0 |
| left_body_lateral_velocity_periodicity | kinematics | 1 | 0 | 0 |
| left_body_lateral_velocity_std | kinematics | 1 | 0 | 0 |
| left_body_speed_energy | kinematics | 1 | 0 | 0 |
| left_body_speed_max | kinematics | 1 | 0 | 0 |
| left_body_speed_mean | kinematics | 1 | 0 | 0 |
| left_body_speed_median | kinematics | 1 | 0 | 0 |
| left_body_speed_p10 | kinematics | 1 | 0 | 0 |
| left_body_speed_p90 | kinematics | 1 | 0 | 0 |
| left_body_speed_periodicity | kinematics | 1 | 0 | 0 |
| left_body_speed_std | kinematics | 1 | 0 | 0 |
| left_body_velocity_x_energy | kinematics | 1 | 0 | 0 |
| left_body_velocity_x_max | kinematics | 1 | 0 | 0 |
| left_body_velocity_x_mean | kinematics | 1 | 0 | 0 |
| left_body_velocity_x_median | kinematics | 1 | 0 | 0 |
| left_body_velocity_x_p10 | kinematics | 1 | 0 | 0 |
| left_body_velocity_x_p90 | kinematics | 1 | 0 | 0 |

| | | | | |
|---|---|---|---|---|
| left_body_velocity_x_periodicity | kinematics | 1 | 0 | 0 |
| left_body_velocity_x_std | kinematics | 1 | 0 | 0 |
| left_body_velocity_y_energy | kinematics | 1 | 0 | 0 |
| left_body_velocity_y_max | kinematics | 1 | 0 | 0 |
| left_body_velocity_y_mean | kinematics | 1 | 0 | 0 |
| left_body_velocity_y_median | kinematics | 1 | 0 | 0 |
| left_body_velocity_y_p10 | kinematics | 1 | 0 | 0 |
| left_body_velocity_y_p90 | kinematics | 1 | 0 | 0 |
| left_body_velocity_y_periodicity | kinematics | 1 | 0 | 0 |
| left_body_velocity_y_std | kinematics | 1 | 0 | 0 |
| left_ear_acceleration_energy | kinematics | 1 | 0 | 0 |
| left_ear_acceleration_max | kinematics | 1 | 0 | 0 |
| left_ear_acceleration_mean | kinematics | 1 | 0 | 0 |
| left_ear_acceleration_median | kinematics | 1 | 0 | 0 |
| left_ear_acceleration_p10 | kinematics | 1 | 0 | 0 |
| left_ear_acceleration_p90 | kinematics | 1 | 0 | 0 |
| left_ear_acceleration_periodicity | kinematics | 1 | 0 | 0 |
| left_ear_acceleration_std | kinematics | 1 | 0 | 0 |
| left_ear_forward_velocity_energy | kinematics | 1 | 0 | 0 |
| left_ear_forward_velocity_max | kinematics | 1 | 0 | 0 |
| left_ear_forward_velocity_mean | kinematics | 1 | 0 | 0 |
| left_ear_forward_velocity_median | kinematics | 1 | 0 | 0 |
| left_ear_forward_velocity_p10 | kinematics | 1 | 0 | 0 |
| left_ear_forward_velocity_p90 | kinematics | 1 | 0 | 0 |
| left_ear_forward_velocity_periodicity | kinematics | 1 | 0 | 0 |
| left_ear_forward_velocity_std | kinematics | 1 | 0 | 0 |
| left_ear_jerk_energy | kinematics | 1 | 0 | 0 |
| left_ear_jerk_max | kinematics | 1 | 0 | 0 |
| left_ear_jerk_mean | kinematics | 1 | 0 | 0 |
| left_ear_jerk_median | kinematics | 1 | 0 | 0 |
| left_ear_jerk_p10 | kinematics | 1 | 0 | 0 |
| left_ear_jerk_p90 | kinematics | 1 | 0 | 0 |
| left_ear_jerk_periodicity | kinematics | 1 | 0 | 0 |
| left_ear_jerk_std | kinematics | 1 | 0 | 0 |
| left_ear_lateral_velocity_energy | kinematics | 1 | 0 | 0 |
| left_ear_lateral_velocity_max | kinematics | 1 | 0 | 0 |
| left_ear_lateral_velocity_mean | kinematics | 1 | 0 | 0 |
| left_ear_lateral_velocity_median | kinematics | 1 | 0 | 0 |
| left_ear_lateral_velocity_p10 | kinematics | 1 | 0 | 0 |
| left_ear_lateral_velocity_p90 | kinematics | 1 | 0 | 0 |
| left_ear_lateral_velocity_periodicity | kinematics | 1 | 0 | 0 |
| left_ear_lateral_velocity_std | kinematics | 1 | 0 | 0 |
| left_ear_speed_energy | kinematics | 1 | 0 | 0 |
| left_ear_speed_max | kinematics | 1 | 0 | 0 |

| | | | | |
|---|---|---|---|---|
| left_ear_speed_mean | kinematics | 1 | 0 | 0 |
| left_ear_speed_median | kinematics | 1 | 0 | 0 |
| left_ear_speed_p10 | kinematics | 1 | 0 | 0 |
| left_ear_speed_p90 | kinematics | 1 | 0 | 0 |
| left_ear_speed_periodicity | kinematics | 1 | 0 | 0 |
| left_ear_speed_std | kinematics | 1 | 0 | 0 |
| left_ear_velocity_x_energy | kinematics | 1 | 0 | 0 |
| left_ear_velocity_x_max | kinematics | 1 | 0 | 0 |
| left_ear_velocity_x_mean | kinematics | 1 | 0 | 0 |
| left_ear_velocity_x_median | kinematics | 1 | 0 | 0 |
| left_ear_velocity_x_p10 | kinematics | 1 | 0 | 0 |
| left_ear_velocity_x_p90 | kinematics | 1 | 0 | 0 |
| left_ear_velocity_x_periodicity | kinematics | 1 | 0 | 0 |
| left_ear_velocity_x_std | kinematics | 1 | 0 | 0 |
| left_ear_velocity_y_energy | kinematics | 1 | 0 | 0 |
| left_ear_velocity_y_max | kinematics | 1 | 0 | 0 |
| left_ear_velocity_y_mean | kinematics | 1 | 0 | 0 |
| left_ear_velocity_y_median | kinematics | 1 | 0 | 0 |
| left_ear_velocity_y_p10 | kinematics | 1 | 0 | 0 |
| left_ear_velocity_y_p90 | kinematics | 1 | 0 | 0 |
| left_ear_velocity_y_periodicity | kinematics | 1 | 0 | 0 |
| left_ear_velocity_y_std | kinematics | 1 | 0 | 0 |
| local_surface_change_rate_energy | kinematics | 1 | 1 | 1 |
| local_surface_change_rate_max | pose geometry | 1 | 1 | 1 |
| local_surface_change_rate_mean | pose geometry | 1 | 1 | 1 |
| local_surface_change_rate_median | pose geometry | 1 | 1 | 1 |
| local_surface_change_rate_p10 | pose geometry | 1 | 1 | 1 |
| local_surface_change_rate_p90 | pose geometry | 1 | 1 | 1 |
| local_surface_change_rate_periodicity | pose geometry | 1 | 1 | 1 |
| local_surface_change_rate_std | kinematics | 1 | 1 | 1 |
| local_surface_motion_energy_energy | video (motion/flow) | 1 | 1 | 1 |
| local_surface_motion_energy_max | video (motion/flow) | 1 | 1 | 1 |
| local_surface_motion_energy_mean | video (motion/flow) | 1 | 1 | 1 |
| local_surface_motion_energy_median | video (motion/flow) | 1 | 1 | 1 |
| local_surface_motion_energy_p10 | video (motion/flow) | 1 | 1 | 1 |
| local_surface_motion_energy_p90 | video (motion/flow) | 1 | 1 | 1 |
| local_surface_motion_energy_periodicity | video (motion/flow) | 1 | 1 | 1 |
| local_surface_motion_energy_std | video (motion/flow) | 1 | 1 | 1 |
| local_surface_motion_variance_energy | kinematics | 1 | 1 | 1 |
| local_surface_motion_variance_max | kinematics | 1 | 1 | 1 |
| local_surface_motion_variance_mean | kinematics | 1 | 1 | 1 |
| local_surface_motion_variance_median | kinematics | 1 | 1 | 1 |
| local_surface_motion_variance_p10 | kinematics | 1 | 1 | 1 |
| local_surface_motion_variance_p90 | kinematics | 1 | 1 | 1 |

| | | | | |
|---|---|---|---|---|
| local_surface_motion_variance_periodicity | kinematics | 1 | 1 | 1 |
| local_surface_motion_variance_std | kinematics | 1 | 1 | 1 |
| mid_back_acceleration_energy | kinematics | 0 | 0 | 0 |
| mid_back_acceleration_max | kinematics | 0 | 0 | 0 |
| mid_back_acceleration_mean | kinematics | 0 | 0 | 0 |
| mid_back_acceleration_median | kinematics | 0 | 0 | 0 |
| mid_back_acceleration_p10 | kinematics | 0 | 0 | 0 |
| mid_back_acceleration_p90 | kinematics | 0 | 0 | 0 |
| mid_back_acceleration_periodicity | kinematics | 0 | 0 | 0 |
| mid_back_acceleration_std | kinematics | 0 | 0 | 0 |
| mid_back_forward_velocity_energy | kinematics | 0 | 0 | 0 |
| mid_back_forward_velocity_max | kinematics | 0 | 0 | 0 |
| mid_back_forward_velocity_mean | kinematics | 0 | 0 | 0 |
| mid_back_forward_velocity_median | kinematics | 0 | 0 | 0 |
| mid_back_forward_velocity_p10 | kinematics | 0 | 0 | 0 |
| mid_back_forward_velocity_p90 | kinematics | 0 | 0 | 0 |
| mid_back_forward_velocity_periodicity | kinematics | 0 | 0 | 0 |
| mid_back_forward_velocity_std | kinematics | 0 | 0 | 0 |
| mid_back_jerk_energy | kinematics | 0 | 0 | 0 |
| mid_back_jerk_max | kinematics | 0 | 0 | 0 |
| mid_back_jerk_mean | kinematics | 0 | 0 | 0 |
| mid_back_jerk_median | kinematics | 0 | 0 | 0 |
| mid_back_jerk_p10 | kinematics | 0 | 0 | 0 |
| mid_back_jerk_p90 | kinematics | 0 | 0 | 0 |
| mid_back_jerk_periodicity | kinematics | 0 | 0 | 0 |
| mid_back_jerk_std | kinematics | 0 | 0 | 0 |
| mid_back_lateral_velocity_energy | kinematics | 0 | 0 | 0 |
| mid_back_lateral_velocity_max | kinematics | 0 | 0 | 0 |
| mid_back_lateral_velocity_mean | kinematics | 0 | 0 | 0 |
| mid_back_lateral_velocity_median | kinematics | 0 | 0 | 0 |
| mid_back_lateral_velocity_p10 | kinematics | 0 | 0 | 0 |
| mid_back_lateral_velocity_p90 | kinematics | 0 | 0 | 0 |
| mid_back_lateral_velocity_periodicity | kinematics | 0 | 0 | 0 |
| mid_back_lateral_velocity_std | kinematics | 0 | 0 | 0 |
| mid_back_speed_energy | kinematics | 0 | 0 | 0 |
| mid_back_speed_max | kinematics | 0 | 0 | 0 |
| mid_back_speed_mean | kinematics | 0 | 0 | 0 |
| mid_back_speed_median | kinematics | 0 | 0 | 0 |
| mid_back_speed_p10 | kinematics | 0 | 0 | 0 |
| mid_back_speed_p90 | kinematics | 0 | 0 | 0 |
| mid_back_speed_periodicity | kinematics | 0 | 0 | 0 |
| mid_back_speed_std | kinematics | 0 | 0 | 0 |
| mid_back_velocity_x_energy | kinematics | 0 | 0 | 0 |
| mid_back_velocity_x_max | kinematics | 0 | 0 | 0 |

| | | | | |
|---|---|---|---|---|
| mid_back_velocity_x_mean | kinematics | 0 | 0 | 0 |
| mid_back_velocity_x_median | kinematics | 0 | 0 | 0 |
| mid_back_velocity_x_p10 | kinematics | 0 | 0 | 0 |
| mid_back_velocity_x_p90 | kinematics | 0 | 0 | 0 |
| mid_back_velocity_x_periodicity | kinematics | 0 | 0 | 0 |
| mid_back_velocity_x_std | kinematics | 0 | 0 | 0 |
| mid_back_velocity_y_energy | kinematics | 0 | 0 | 0 |
| mid_back_velocity_y_max | kinematics | 0 | 0 | 0 |
| mid_back_velocity_y_mean | kinematics | 0 | 0 | 0 |
| mid_back_velocity_y_median | kinematics | 0 | 0 | 0 |
| mid_back_velocity_y_p10 | kinematics | 0 | 0 | 0 |
| mid_back_velocity_y_p90 | kinematics | 0 | 0 | 0 |
| mid_back_velocity_y_periodicity | kinematics | 0 | 0 | 0 |
| mid_back_velocity_y_std | kinematics | 0 | 0 | 0 |
| nose_acceleration_energy | kinematics | 1 | 1 | 1 |
| nose_acceleration_max | kinematics | 1 | 1 | 1 |
| nose_acceleration_mean | kinematics | 1 | 1 | 1 |
| nose_acceleration_median | kinematics | 1 | 1 | 1 |
| nose_acceleration_p10 | kinematics | 1 | 1 | 1 |
| nose_acceleration_p90 | kinematics | 1 | 1 | 1 |
| nose_acceleration_periodicity | kinematics | 1 | 1 | 1 |
| nose_acceleration_std | kinematics | 1 | 1 | 1 |
| nose_autocorr_peak_energy | kinematics | 1 | 1 | 1 |
| nose_autocorr_peak_max | pose geometry | 1 | 1 | 1 |
| nose_autocorr_peak_mean | pose geometry | 1 | 1 | 1 |
| nose_autocorr_peak_median | pose geometry | 1 | 1 | 1 |
| nose_autocorr_peak_p10 | pose geometry | 1 | 1 | 1 |
| nose_autocorr_peak_p90 | pose geometry | 1 | 1 | 1 |
| nose_autocorr_peak_periodicity | pose geometry | 1 | 1 | 1 |
| nose_autocorr_peak_std | kinematics | 1 | 1 | 1 |
| nose_forward_velocity_energy | kinematics | 1 | 1 | 1 |
| nose_forward_velocity_max | kinematics | 1 | 1 | 1 |
| nose_forward_velocity_mean | kinematics | 1 | 1 | 1 |
| nose_forward_velocity_median | kinematics | 1 | 1 | 1 |
| nose_forward_velocity_p10 | kinematics | 1 | 1 | 1 |
| nose_forward_velocity_p90 | kinematics | 1 | 1 | 1 |
| nose_forward_velocity_periodicity | kinematics | 1 | 1 | 1 |
| nose_forward_velocity_std | kinematics | 1 | 1 | 1 |
| nose_jerk_energy | kinematics | 1 | 1 | 1 |
| nose_jerk_max | kinematics | 1 | 1 | 1 |
| nose_jerk_mean | kinematics | 1 | 1 | 1 |
| nose_jerk_median | kinematics | 1 | 1 | 1 |
| nose_jerk_p10 | kinematics | 1 | 1 | 1 |
| nose_jerk_p90 | kinematics | 1 | 1 | 1 |

| | | | | |
|---|---|---|---|---|
| nose_jerk_periodicity | kinematics | 1 | 1 | 1 |
| nose_jerk_std | kinematics | 1 | 1 | 1 |
| nose_lateral_velocity_energy | kinematics | 1 | 1 | 1 |
| nose_lateral_velocity_max | kinematics | 1 | 1 | 1 |
| nose_lateral_velocity_mean | kinematics | 1 | 1 | 1 |
| nose_lateral_velocity_median | kinematics | 1 | 1 | 1 |
| nose_lateral_velocity_p10 | kinematics | 1 | 1 | 1 |
| nose_lateral_velocity_p90 | kinematics | 1 | 1 | 1 |
| nose_lateral_velocity_periodicity | kinematics | 1 | 1 | 1 |
| nose_lateral_velocity_std | kinematics | 1 | 1 | 1 |
| nose_local_change_rate_energy | kinematics | 1 | 1 | 1 |
| nose_local_change_rate_max | pose geometry | 1 | 1 | 1 |
| nose_local_change_rate_mean | pose geometry | 1 | 1 | 1 |
| nose_local_change_rate_median | pose geometry | 1 | 1 | 1 |
| nose_local_change_rate_p10 | pose geometry | 1 | 1 | 1 |
| nose_local_change_rate_p90 | pose geometry | 1 | 1 | 1 |
| nose_local_change_rate_periodicity | pose geometry | 1 | 1 | 1 |
| nose_local_change_rate_std | kinematics | 1 | 1 | 1 |
| nose_local_variance_energy | kinematics | 1 | 1 | 1 |
| nose_local_variance_max | kinematics | 1 | 1 | 1 |
| nose_local_variance_mean | kinematics | 1 | 1 | 1 |
| nose_local_variance_median | kinematics | 1 | 1 | 1 |
| nose_local_variance_p10 | kinematics | 1 | 1 | 1 |
| nose_local_variance_p90 | kinematics | 1 | 1 | 1 |
| nose_local_variance_periodicity | kinematics | 1 | 1 | 1 |
| nose_local_variance_std | kinematics | 1 | 1 | 1 |
| nose_movement_frequency_energy | kinematics | 1 | 1 | 1 |
| nose_movement_frequency_max | kinematics | 1 | 1 | 1 |
| nose_movement_frequency_mean | kinematics | 1 | 1 | 1 |
| nose_movement_frequency_median | kinematics | 1 | 1 | 1 |
| nose_movement_frequency_p10 | kinematics | 1 | 1 | 1 |
| nose_movement_frequency_p90 | kinematics | 1 | 1 | 1 |
| nose_movement_frequency_periodicity | kinematics | 1 | 1 | 1 |
| nose_movement_frequency_std | kinematics | 1 | 1 | 1 |
| nose_oscillation_energy_energy | kinematics | 1 | 1 | 1 |
| nose_oscillation_energy_max | kinematics | 1 | 1 | 1 |
| nose_oscillation_energy_mean | kinematics | 1 | 1 | 1 |
| nose_oscillation_energy_median | kinematics | 1 | 1 | 1 |
| nose_oscillation_energy_p10 | kinematics | 1 | 1 | 1 |
| nose_oscillation_energy_p90 | kinematics | 1 | 1 | 1 |
| nose_oscillation_energy_periodicity | kinematics | 1 | 1 | 1 |
| nose_oscillation_energy_std | kinematics | 1 | 1 | 1 |
| nose_oscillation_power_energy | kinematics | 1 | 1 | 1 |
| nose_oscillation_power_max | pose geometry | 1 | 1 | 1 |

| | | | | |
|---|---|---|---|---|
| nose_oscillation_power_mean | pose geometry | 1 | 1 | 1 |
| nose_oscillation_power_median | pose geometry | 1 | 1 | 1 |
| nose_oscillation_power_p10 | pose geometry | 1 | 1 | 1 |
| nose_oscillation_power_p90 | pose geometry | 1 | 1 | 1 |
| nose_oscillation_power_periodicity | pose geometry | 1 | 1 | 1 |
| nose_oscillation_power_std | kinematics | 1 | 1 | 1 |
| nose_roi_1_axial_abs_energy | context (ROI/target) | 1 | 1 | 1 |
| nose_roi_1_axial_abs_max | context (ROI/target) | 1 | 1 | 1 |
| nose_roi_1_axial_abs_mean | context (ROI/target) | 1 | 1 | 1 |
| nose_roi_1_axial_abs_median | context (ROI/target) | 1 | 1 | 1 |
| nose_roi_1_axial_abs_p10 | context (ROI/target) | 1 | 1 | 1 |
| nose_roi_1_axial_abs_p90 | context (ROI/target) | 1 | 1 | 1 |
| nose_roi_1_axial_abs_periodicity | context (ROI/target) | 1 | 1 | 1 |
| nose_roi_1_axial_abs_std | context (ROI/target) | 1 | 1 | 1 |
| nose_roi_1_axial_energy | context (ROI/target) | 1 | 1 | 1 |
| nose_roi_1_axial_max | context (ROI/target) | 1 | 1 | 1 |
| nose_roi_1_axial_mean | context (ROI/target) | 1 | 1 | 1 |
| nose_roi_1_axial_median | context (ROI/target) | 1 | 1 | 1 |
| nose_roi_1_axial_p10 | context (ROI/target) | 1 | 1 | 1 |
| nose_roi_1_axial_p90 | context (ROI/target) | 1 | 1 | 1 |
| nose_roi_1_axial_periodicity | context (ROI/target) | 1 | 1 | 1 |
| nose_roi_1_axial_std | context (ROI/target) | 1 | 1 | 1 |
| nose_roi_1_lateral_abs_energy | context (ROI/target) | 1 | 1 | 1 |
| nose_roi_1_lateral_abs_max | context (ROI/target) | 1 | 1 | 1 |
| nose_roi_1_lateral_abs_mean | context (ROI/target) | 1 | 1 | 1 |
| nose_roi_1_lateral_abs_median | context (ROI/target) | 1 | 1 | 1 |
| nose_roi_1_lateral_abs_p10 | context (ROI/target) | 1 | 1 | 1 |
| nose_roi_1_lateral_abs_p90 | context (ROI/target) | 1 | 1 | 1 |
| nose_roi_1_lateral_abs_periodicity | context (ROI/target) | 1 | 1 | 1 |
| nose_roi_1_lateral_abs_std | context (ROI/target) | 1 | 1 | 1 |
| nose_roi_1_lateral_energy | context (ROI/target) | 1 | 1 | 1 |
| nose_roi_1_lateral_max | context (ROI/target) | 1 | 1 | 1 |
| nose_roi_1_lateral_mean | context (ROI/target) | 1 | 1 | 1 |
| nose_roi_1_lateral_median | context (ROI/target) | 1 | 1 | 1 |
| nose_roi_1_lateral_p10 | context (ROI/target) | 1 | 1 | 1 |
| nose_roi_1_lateral_p90 | context (ROI/target) | 1 | 1 | 1 |
| nose_roi_1_lateral_periodicity | context (ROI/target) | 1 | 1 | 1 |
| nose_roi_1_lateral_std | context (ROI/target) | 1 | 1 | 1 |
| nose_roi_2_axial_abs_energy | context (ROI/target) | 1 | 0 | 0 |
| nose_roi_2_axial_abs_max | context (ROI/target) | 1 | 0 | 0 |
| nose_roi_2_axial_abs_mean | context (ROI/target) | 1 | 0 | 0 |
| nose_roi_2_axial_abs_median | context (ROI/target) | 1 | 0 | 0 |
| nose_roi_2_axial_abs_p10 | context (ROI/target) | 1 | 0 | 0 |
| nose_roi_2_axial_abs_p90 | context (ROI/target) | 1 | 0 | 0 |

| | | | | |
|---|---|---|---|---|
| nose_roi_2_axial_abs_periodicity | context (ROI/target) | 1 | 0 | 0 |
| nose_roi_2_axial_abs_std | context (ROI/target) | 1 | 0 | 0 |
| nose_roi_2_axial_energy | context (ROI/target) | 1 | 0 | 0 |
| nose_roi_2_axial_max | context (ROI/target) | 1 | 0 | 0 |
| nose_roi_2_axial_mean | context (ROI/target) | 1 | 0 | 0 |
| nose_roi_2_axial_median | context (ROI/target) | 1 | 0 | 0 |
| nose_roi_2_axial_p10 | context (ROI/target) | 1 | 0 | 0 |
| nose_roi_2_axial_p90 | context (ROI/target) | 1 | 0 | 0 |
| nose_roi_2_axial_periodicity | context (ROI/target) | 1 | 0 | 0 |
| nose_roi_2_axial_std | context (ROI/target) | 1 | 0 | 0 |
| nose_roi_2_lateral_abs_energy | context (ROI/target) | 1 | 0 | 0 |
| nose_roi_2_lateral_abs_max | context (ROI/target) | 1 | 0 | 0 |
| nose_roi_2_lateral_abs_mean | context (ROI/target) | 1 | 0 | 0 |
| nose_roi_2_lateral_abs_median | context (ROI/target) | 1 | 0 | 0 |
| nose_roi_2_lateral_abs_p10 | context (ROI/target) | 1 | 0 | 0 |
| nose_roi_2_lateral_abs_p90 | context (ROI/target) | 1 | 0 | 0 |
| nose_roi_2_lateral_abs_periodicity | context (ROI/target) | 1 | 0 | 0 |
| nose_roi_2_lateral_abs_std | context (ROI/target) | 1 | 0 | 0 |
| nose_roi_2_lateral_energy | context (ROI/target) | 1 | 0 | 0 |
| nose_roi_2_lateral_max | context (ROI/target) | 1 | 0 | 0 |
| nose_roi_2_lateral_mean | context (ROI/target) | 1 | 0 | 0 |
| nose_roi_2_lateral_median | context (ROI/target) | 1 | 0 | 0 |
| nose_roi_2_lateral_p10 | context (ROI/target) | 1 | 0 | 0 |
| nose_roi_2_lateral_p90 | context (ROI/target) | 1 | 0 | 0 |
| nose_roi_2_lateral_periodicity | context (ROI/target) | 1 | 0 | 0 |
| nose_roi_2_lateral_std | context (ROI/target) | 1 | 0 | 0 |
| nose_speed_energy | kinematics | 1 | 1 | 1 |
| nose_speed_max | kinematics | 1 | 1 | 1 |
| nose_speed_mean | kinematics | 1 | 1 | 1 |
| nose_speed_median | kinematics | 1 | 1 | 1 |
| nose_speed_p10 | kinematics | 1 | 1 | 1 |
| nose_speed_p90 | kinematics | 1 | 1 | 1 |
| nose_speed_periodicity | kinematics | 1 | 1 | 1 |
| nose_speed_std | kinematics | 1 | 1 | 1 |
| nose_surface_change_rate_energy | kinematics | 1 | 1 | 1 |
| nose_surface_change_rate_max | pose geometry | 1 | 1 | 1 |
| nose_surface_change_rate_mean | pose geometry | 1 | 1 | 1 |
| nose_surface_change_rate_median | pose geometry | 1 | 1 | 1 |
| nose_surface_change_rate_p10 | pose geometry | 1 | 1 | 1 |
| nose_surface_change_rate_p90 | pose geometry | 1 | 1 | 1 |
| nose_surface_change_rate_periodicity | pose geometry | 1 | 1 | 1 |
| nose_surface_change_rate_std | kinematics | 1 | 1 | 1 |
| nose_surface_motion_energy_energy | video (motion/flow) | 1 | 1 | 1 |
| nose_surface_motion_energy_max | video (motion/flow) | 1 | 1 | 1 |

| | | | | |
|---|---|---|---|---|
| nose_surface_motion_energy_mean | video (motion/flow) | 1 | 1 | 1 |
| nose_surface_motion_energy_median | video (motion/flow) | 1 | 1 | 1 |
| nose_surface_motion_energy_p10 | video (motion/flow) | 1 | 1 | 1 |
| nose_surface_motion_energy_p90 | video (motion/flow) | 1 | 1 | 1 |
| nose_surface_motion_energy_periodicity | video (motion/flow) | 1 | 1 | 1 |
| nose_surface_motion_energy_std | video (motion/flow) | 1 | 1 | 1 |
| nose_surface_motion_variance_energy | kinematics | 1 | 1 | 1 |
| nose_surface_motion_variance_max | kinematics | 1 | 1 | 1 |
| nose_surface_motion_variance_mean | kinematics | 1 | 1 | 1 |
| nose_surface_motion_variance_median | kinematics | 1 | 1 | 1 |
| nose_surface_motion_variance_p10 | kinematics | 1 | 1 | 1 |
| nose_surface_motion_variance_p90 | kinematics | 1 | 1 | 1 |
| nose_surface_motion_variance_periodicity | kinematics | 1 | 1 | 1 |
| nose_surface_motion_variance_std | kinematics | 1 | 1 | 1 |
| nose_to_roi_1_corner_dist_delta | context (ROI/target) | 1 | 1 | 1 |
| nose_to_roi_1_corner_dist_energy | context (ROI/target) | 1 | 1 | 1 |
| nose_to_roi_1_corner_dist_max | context (ROI/target) | 1 | 1 | 1 |
| nose_to_roi_1_corner_dist_mean | context (ROI/target) | 1 | 1 | 1 |
| nose_to_roi_1_corner_dist_median | context (ROI/target) | 1 | 1 | 1 |
| nose_to_roi_1_corner_dist_p10 | context (ROI/target) | 1 | 1 | 1 |
| nose_to_roi_1_corner_dist_p90 | context (ROI/target) | 1 | 1 | 1 |
| nose_to_roi_1_corner_dist_periodicity | context (ROI/target) | 1 | 1 | 1 |
| nose_to_roi_1_corner_dist_std | context (ROI/target) | 1 | 1 | 1 |
| nose_to_roi_1_corner_dist_trend | context (ROI/target) | 1 | 1 | 1 |
| nose_to_roi_1_dist_delta | context (ROI/target) | 1 | 1 | 1 |
| nose_to_roi_1_dist_energy | context (ROI/target) | 1 | 1 | 1 |
| nose_to_roi_1_dist_max | context (ROI/target) | 1 | 1 | 1 |
| nose_to_roi_1_dist_mean | context (ROI/target) | 1 | 1 | 1 |
| nose_to_roi_1_dist_median | context (ROI/target) | 1 | 1 | 1 |
| nose_to_roi_1_dist_p10 | context (ROI/target) | 1 | 1 | 1 |
| nose_to_roi_1_dist_p90 | context (ROI/target) | 1 | 1 | 1 |
| nose_to_roi_1_dist_periodicity | context (ROI/target) | 1 | 1 | 1 |
| nose_to_roi_1_dist_std | context (ROI/target) | 1 | 1 | 1 |
| nose_to_roi_1_dist_trend | context (ROI/target) | 1 | 1 | 1 |
| nose_to_roi_1_edge_dist_delta | context (ROI/target) | 1 | 1 | 1 |
| nose_to_roi_1_edge_dist_energy | context (ROI/target) | 1 | 1 | 1 |
| nose_to_roi_1_edge_dist_max | context (ROI/target) | 1 | 1 | 1 |
| nose_to_roi_1_edge_dist_mean | context (ROI/target) | 1 | 1 | 1 |
| nose_to_roi_1_edge_dist_median | context (ROI/target) | 1 | 1 | 1 |
| nose_to_roi_1_edge_dist_p10 | context (ROI/target) | 1 | 1 | 1 |
| nose_to_roi_1_edge_dist_p90 | context (ROI/target) | 1 | 1 | 1 |
| nose_to_roi_1_edge_dist_periodicity | context (ROI/target) | 1 | 1 | 1 |
| nose_to_roi_1_edge_dist_std | context (ROI/target) | 1 | 1 | 1 |
| nose_to_roi_1_edge_dist_trend | context (ROI/target) | 1 | 1 | 1 |

| | | | | |
|---|---|---|---|---|
| nose_to_roi_1_signed_dist_delta | context (ROI/target) | 1 | 1 | 1 |
| nose_to_roi_1_signed_dist_energy | context (ROI/target) | 1 | 1 | 1 |
| nose_to_roi_1_signed_dist_max | context (ROI/target) | 1 | 1 | 1 |
| nose_to_roi_1_signed_dist_mean | context (ROI/target) | 1 | 1 | 1 |
| nose_to_roi_1_signed_dist_median | context (ROI/target) | 1 | 1 | 1 |
| nose_to_roi_1_signed_dist_p10 | context (ROI/target) | 1 | 1 | 1 |
| nose_to_roi_1_signed_dist_p90 | context (ROI/target) | 1 | 1 | 1 |
| nose_to_roi_1_signed_dist_periodicity | context (ROI/target) | 1 | 1 | 1 |
| nose_to_roi_1_signed_dist_std | context (ROI/target) | 1 | 1 | 1 |
| nose_to_roi_1_signed_dist_trend | context (ROI/target) | 1 | 1 | 1 |
| nose_to_roi_2_corner_dist_delta | context (ROI/target) | 1 | 0 | 0 |
| nose_to_roi_2_corner_dist_energy | context (ROI/target) | 1 | 0 | 0 |
| nose_to_roi_2_corner_dist_max | context (ROI/target) | 1 | 0 | 0 |
| nose_to_roi_2_corner_dist_mean | context (ROI/target) | 1 | 0 | 0 |
| nose_to_roi_2_corner_dist_median | context (ROI/target) | 1 | 0 | 0 |
| nose_to_roi_2_corner_dist_p10 | context (ROI/target) | 1 | 0 | 0 |
| nose_to_roi_2_corner_dist_p90 | context (ROI/target) | 1 | 0 | 0 |
| nose_to_roi_2_corner_dist_periodicity | context (ROI/target) | 1 | 0 | 0 |
| nose_to_roi_2_corner_dist_std | context (ROI/target) | 1 | 0 | 0 |
| nose_to_roi_2_corner_dist_trend | context (ROI/target) | 1 | 0 | 0 |
| nose_to_roi_2_dist_delta | context (ROI/target) | 1 | 0 | 0 |
| nose_to_roi_2_dist_energy | context (ROI/target) | 1 | 0 | 0 |
| nose_to_roi_2_dist_max | context (ROI/target) | 1 | 0 | 0 |
| nose_to_roi_2_dist_mean | context (ROI/target) | 1 | 0 | 0 |
| nose_to_roi_2_dist_median | context (ROI/target) | 1 | 0 | 0 |
| nose_to_roi_2_dist_p10 | context (ROI/target) | 1 | 0 | 0 |
| nose_to_roi_2_dist_p90 | context (ROI/target) | 1 | 0 | 0 |
| nose_to_roi_2_dist_periodicity | context (ROI/target) | 1 | 0 | 0 |
| nose_to_roi_2_dist_std | context (ROI/target) | 1 | 0 | 0 |
| nose_to_roi_2_dist_trend | context (ROI/target) | 1 | 0 | 0 |
| nose_to_roi_2_edge_dist_delta | context (ROI/target) | 1 | 0 | 0 |
| nose_to_roi_2_edge_dist_energy | context (ROI/target) | 1 | 0 | 0 |
| nose_to_roi_2_edge_dist_max | context (ROI/target) | 1 | 0 | 0 |
| nose_to_roi_2_edge_dist_mean | context (ROI/target) | 1 | 0 | 0 |
| nose_to_roi_2_edge_dist_median | context (ROI/target) | 1 | 0 | 0 |
| nose_to_roi_2_edge_dist_p10 | context (ROI/target) | 1 | 0 | 0 |
| nose_to_roi_2_edge_dist_p90 | context (ROI/target) | 1 | 0 | 0 |
| nose_to_roi_2_edge_dist_periodicity | context (ROI/target) | 1 | 0 | 0 |
| nose_to_roi_2_edge_dist_std | context (ROI/target) | 1 | 0 | 0 |
| nose_to_roi_2_edge_dist_trend | context (ROI/target) | 1 | 0 | 0 |
| nose_to_roi_2_signed_dist_delta | context (ROI/target) | 1 | 0 | 0 |
| nose_to_roi_2_signed_dist_energy | context (ROI/target) | 1 | 0 | 0 |
| nose_to_roi_2_signed_dist_max | context (ROI/target) | 1 | 0 | 0 |
| nose_to_roi_2_signed_dist_mean | context (ROI/target) | 1 | 0 | 0 |

| | | | | |
|---|---|---|---|---|
| nose_to_roi_2_signed_dist_median | context (ROI/target) | 1 | 0 | 0 |
| nose_to_roi_2_signed_dist_p10 | context (ROI/target) | 1 | 0 | 0 |
| nose_to_roi_2_signed_dist_p90 | context (ROI/target) | 1 | 0 | 0 |
| nose_to_roi_2_signed_dist_periodicity | context (ROI/target) | 1 | 0 | 0 |
| nose_to_roi_2_signed_dist_std | context (ROI/target) | 1 | 0 | 0 |
| nose_to_roi_2_signed_dist_trend | context (ROI/target) | 1 | 0 | 0 |
| nose_to_target_dist_delta | context (ROI/target) | 1 | 1 | 1 |
| nose_to_target_dist_energy | context (ROI/target) | 1 | 1 | 1 |
| nose_to_target_dist_max | context (ROI/target) | 1 | 1 | 1 |
| nose_to_target_dist_mean | context (ROI/target) | 1 | 1 | 1 |
| nose_to_target_dist_median | context (ROI/target) | 1 | 1 | 1 |
| nose_to_target_dist_p10 | context (ROI/target) | 1 | 1 | 1 |
| nose_to_target_dist_p90 | context (ROI/target) | 1 | 1 | 1 |
| nose_to_target_dist_periodicity | context (ROI/target) | 1 | 1 | 1 |
| nose_to_target_dist_std | context (ROI/target) | 1 | 1 | 1 |
| nose_to_target_dist_trend | context (ROI/target) | 1 | 1 | 1 |
| nose_velocity_energy | kinematics | 1 | 1 | 1 |
| nose_velocity_max | kinematics | 1 | 1 | 1 |
| nose_velocity_mean | kinematics | 1 | 1 | 1 |
| nose_velocity_median | kinematics | 1 | 1 | 1 |
| nose_velocity_p10 | kinematics | 1 | 1 | 1 |
| nose_velocity_p90 | kinematics | 1 | 1 | 1 |
| nose_velocity_periodicity | kinematics | 1 | 1 | 1 |
| nose_velocity_std | kinematics | 1 | 1 | 1 |
| nose_velocity_x_energy | kinematics | 1 | 1 | 1 |
| nose_velocity_x_max | kinematics | 1 | 1 | 1 |
| nose_velocity_x_mean | kinematics | 1 | 1 | 1 |
| nose_velocity_x_median | kinematics | 1 | 1 | 1 |
| nose_velocity_x_p10 | kinematics | 1 | 1 | 1 |
| nose_velocity_x_p90 | kinematics | 1 | 1 | 1 |
| nose_velocity_x_periodicity | kinematics | 1 | 1 | 1 |
| nose_velocity_x_std | kinematics | 1 | 1 | 1 |
| nose_velocity_y_energy | kinematics | 1 | 1 | 1 |
| nose_velocity_y_max | kinematics | 1 | 1 | 1 |
| nose_velocity_y_mean | kinematics | 1 | 1 | 1 |
| nose_velocity_y_median | kinematics | 1 | 1 | 1 |
| nose_velocity_y_p10 | kinematics | 1 | 1 | 1 |
| nose_velocity_y_p90 | kinematics | 1 | 1 | 1 |
| nose_velocity_y_periodicity | kinematics | 1 | 1 | 1 |
| nose_velocity_y_std | kinematics | 1 | 1 | 1 |
| nose_vertical_velocity_energy | kinematics | 1 | 1 | 1 |
| nose_vertical_velocity_max | kinematics | 1 | 1 | 1 |
| nose_vertical_velocity_mean | kinematics | 1 | 1 | 1 |
| nose_vertical_velocity_median | kinematics | 1 | 1 | 1 |

| | | | | |
|---|---|---|---|---|
| nose_vertical_velocity_p10 | kinematics | 1 | 1 | 1 |
| nose_vertical_velocity_p90 | kinematics | 1 | 1 | 1 |
| nose_vertical_velocity_periodicity | kinematics | 1 | 1 | 1 |
| nose_vertical_velocity_std | kinematics | 1 | 1 | 1 |
| oscillation_energy_energy | kinematics | 1 | 1 | 1 |
| oscillation_energy_max | kinematics | 1 | 1 | 1 |
| oscillation_energy_mean | kinematics | 1 | 1 | 1 |
| oscillation_energy_median | kinematics | 1 | 1 | 1 |
| oscillation_energy_p10 | kinematics | 1 | 1 | 1 |
| oscillation_energy_p90 | kinematics | 1 | 1 | 1 |
| oscillation_energy_periodicity | kinematics | 1 | 1 | 1 |
| oscillation_energy_std | kinematics | 1 | 1 | 1 |
| prediction_prob | pose geometry | 1 | 1 | 1 |
| prediction_variance | kinematics | 1 | 1 | 1 |
| r3d_000 | video (R3D appearance | 1 | 1 | 1 |
| r3d_001 | video (R3D appearance | 1 | 1 | 1 |
| r3d_002 | video (R3D appearance | 1 | 1 | 1 |
| r3d_003 | video (R3D appearance | 1 | 1 | 1 |
| r3d_004 | video (R3D appearance | 1 | 1 | 1 |
| r3d_005 | video (R3D appearance | 1 | 1 | 1 |
| r3d_006 | video (R3D appearance | 1 | 1 | 1 |
| r3d_007 | video (R3D appearance | 1 | 1 | 1 |
| r3d_008 | video (R3D appearance | 1 | 1 | 1 |
| r3d_009 | video (R3D appearance | 1 | 1 | 1 |
| r3d_010 | video (R3D appearance | 1 | 1 | 1 |
| r3d_011 | video (R3D appearance | 1 | 1 | 1 |
| r3d_012 | video (R3D appearance | 1 | 1 | 1 |
| r3d_013 | video (R3D appearance | 1 | 1 | 1 |
| r3d_014 | video (R3D appearance | 1 | 1 | 1 |
| r3d_015 | video (R3D appearance | 1 | 1 | 1 |
| r3d_016 | video (R3D appearance | 1 | 1 | 1 |
| r3d_017 | video (R3D appearance | 1 | 1 | 1 |
| r3d_018 | video (R3D appearance | 1 | 1 | 1 |
| r3d_019 | video (R3D appearance | 1 | 1 | 1 |
| r3d_020 | video (R3D appearance | 1 | 1 | 1 |
| r3d_021 | video (R3D appearance | 1 | 1 | 1 |
| r3d_022 | video (R3D appearance | 1 | 1 | 1 |
| r3d_023 | video (R3D appearance | 1 | 1 | 1 |
| r3d_024 | video (R3D appearance | 1 | 1 | 1 |
| r3d_025 | video (R3D appearance | 1 | 1 | 1 |
| r3d_026 | video (R3D appearance | 1 | 1 | 1 |
| r3d_027 | video (R3D appearance | 1 | 1 | 1 |
| r3d_028 | video (R3D appearance | 1 | 1 | 1 |
| r3d_029 | video (R3D appearance | 1 | 1 | 1 |

| | | | | |
|---|---|---|---|---|
| r3d_030 | video (R3D appearance | 1 | 1 | 1 |
| r3d_031 | video (R3D appearance | 1 | 1 | 1 |
| r3d_032 | video (R3D appearance | 1 | 1 | 1 |
| r3d_033 | video (R3D appearance | 1 | 1 | 1 |
| r3d_034 | video (R3D appearance | 1 | 1 | 1 |
| r3d_035 | video (R3D appearance | 1 | 1 | 1 |
| r3d_036 | video (R3D appearance | 1 | 1 | 1 |
| r3d_037 | video (R3D appearance | 1 | 1 | 1 |
| r3d_038 | video (R3D appearance | 1 | 1 | 1 |
| r3d_039 | video (R3D appearance | 1 | 1 | 1 |
| r3d_040 | video (R3D appearance | 1 | 1 | 1 |
| r3d_041 | video (R3D appearance | 1 | 1 | 1 |
| r3d_042 | video (R3D appearance | 1 | 1 | 1 |
| r3d_043 | video (R3D appearance | 1 | 1 | 1 |
| r3d_044 | video (R3D appearance | 1 | 1 | 1 |
| r3d_045 | video (R3D appearance | 1 | 1 | 1 |
| r3d_046 | video (R3D appearance | 1 | 1 | 1 |
| r3d_047 | video (R3D appearance | 1 | 1 | 1 |
| r3d_048 | video (R3D appearance | 1 | 1 | 1 |
| r3d_049 | video (R3D appearance | 1 | 1 | 1 |
| r3d_050 | video (R3D appearance | 1 | 1 | 1 |
| r3d_051 | video (R3D appearance | 1 | 1 | 1 |
| r3d_052 | video (R3D appearance | 1 | 1 | 1 |
| r3d_053 | video (R3D appearance | 1 | 1 | 1 |
| r3d_054 | video (R3D appearance | 1 | 1 | 1 |
| r3d_055 | video (R3D appearance | 1 | 1 | 1 |
| r3d_056 | video (R3D appearance | 1 | 1 | 1 |
| r3d_057 | video (R3D appearance | 1 | 1 | 1 |
| r3d_058 | video (R3D appearance | 1 | 1 | 1 |
| r3d_059 | video (R3D appearance | 1 | 1 | 1 |
| r3d_060 | video (R3D appearance | 1 | 1 | 1 |
| r3d_061 | video (R3D appearance | 1 | 1 | 1 |
| r3d_062 | video (R3D appearance | 1 | 1 | 1 |
| r3d_063 | video (R3D appearance | 1 | 1 | 1 |
| r3d_064 | video (R3D appearance | 1 | 1 | 1 |
| r3d_065 | video (R3D appearance | 1 | 1 | 1 |
| r3d_066 | video (R3D appearance | 1 | 1 | 1 |
| r3d_067 | video (R3D appearance | 1 | 1 | 1 |
| r3d_068 | video (R3D appearance | 1 | 1 | 1 |
| r3d_069 | video (R3D appearance | 1 | 1 | 1 |
| r3d_070 | video (R3D appearance | 1 | 1 | 1 |
| r3d_071 | video (R3D appearance | 1 | 1 | 1 |
| r3d_072 | video (R3D appearance | 1 | 1 | 1 |
| r3d_073 | video (R3D appearance | 1 | 1 | 1 |

| | | | | |
|---|---|---|---|---|
| r3d_074 | video (R3D appearance | 1 | 1 | 1 |
| r3d_075 | video (R3D appearance | 1 | 1 | 1 |
| r3d_076 | video (R3D appearance | 1 | 1 | 1 |
| r3d_077 | video (R3D appearance | 1 | 1 | 1 |
| r3d_078 | video (R3D appearance | 1 | 1 | 1 |
| r3d_079 | video (R3D appearance | 1 | 1 | 1 |
| r3d_080 | video (R3D appearance | 1 | 1 | 1 |
| r3d_081 | video (R3D appearance | 1 | 1 | 1 |
| r3d_082 | video (R3D appearance | 1 | 1 | 1 |
| r3d_083 | video (R3D appearance | 1 | 1 | 1 |
| r3d_084 | video (R3D appearance | 1 | 1 | 1 |
| r3d_085 | video (R3D appearance | 1 | 1 | 1 |
| r3d_086 | video (R3D appearance | 1 | 1 | 1 |
| r3d_087 | video (R3D appearance | 1 | 1 | 1 |
| r3d_088 | video (R3D appearance | 1 | 1 | 1 |
| r3d_089 | video (R3D appearance | 1 | 1 | 1 |
| r3d_090 | video (R3D appearance | 1 | 1 | 1 |
| r3d_091 | video (R3D appearance | 1 | 1 | 1 |
| r3d_092 | video (R3D appearance | 1 | 1 | 1 |
| r3d_093 | video (R3D appearance | 1 | 1 | 1 |
| r3d_094 | video (R3D appearance | 1 | 1 | 1 |
| r3d_095 | video (R3D appearance | 1 | 1 | 1 |
| r3d_096 | video (R3D appearance | 1 | 1 | 1 |
| r3d_097 | video (R3D appearance | 1 | 1 | 1 |
| r3d_098 | video (R3D appearance | 1 | 1 | 1 |
| r3d_099 | video (R3D appearance | 1 | 1 | 1 |
| r3d_100 | video (R3D appearance | 1 | 1 | 1 |
| r3d_101 | video (R3D appearance | 1 | 1 | 1 |
| r3d_102 | video (R3D appearance | 1 | 1 | 1 |
| r3d_103 | video (R3D appearance | 1 | 1 | 1 |
| r3d_104 | video (R3D appearance | 1 | 1 | 1 |
| r3d_105 | video (R3D appearance | 1 | 1 | 1 |
| r3d_106 | video (R3D appearance | 1 | 1 | 1 |
| r3d_107 | video (R3D appearance | 1 | 1 | 1 |
| r3d_108 | video (R3D appearance | 1 | 1 | 1 |
| r3d_109 | video (R3D appearance | 1 | 1 | 1 |
| r3d_110 | video (R3D appearance | 1 | 1 | 1 |
| r3d_111 | video (R3D appearance | 1 | 1 | 1 |
| r3d_112 | video (R3D appearance | 1 | 1 | 1 |
| r3d_113 | video (R3D appearance | 1 | 1 | 1 |
| r3d_114 | video (R3D appearance | 1 | 1 | 1 |
| r3d_115 | video (R3D appearance | 1 | 1 | 1 |
| r3d_116 | video (R3D appearance | 1 | 1 | 1 |
| r3d_117 | video (R3D appearance | 1 | 1 | 1 |

| | | | | |
|---|---|---|---|---|
| r3d_118 | video (R3D appearance | 1 | 1 | 1 |
| r3d_119 | video (R3D appearance | 1 | 1 | 1 |
| r3d_120 | video (R3D appearance | 1 | 1 | 1 |
| r3d_121 | video (R3D appearance | 1 | 1 | 1 |
| r3d_122 | video (R3D appearance | 1 | 1 | 1 |
| r3d_123 | video (R3D appearance | 1 | 1 | 1 |
| r3d_124 | video (R3D appearance | 1 | 1 | 1 |
| r3d_125 | video (R3D appearance | 1 | 1 | 1 |
| r3d_126 | video (R3D appearance | 1 | 1 | 1 |
| r3d_127 | video (R3D appearance | 1 | 1 | 1 |
| r3d_128 | video (R3D appearance | 1 | 1 | 1 |
| r3d_129 | video (R3D appearance | 1 | 1 | 1 |
| r3d_130 | video (R3D appearance | 1 | 1 | 1 |
| r3d_131 | video (R3D appearance | 1 | 1 | 1 |
| r3d_132 | video (R3D appearance | 1 | 1 | 1 |
| r3d_133 | video (R3D appearance | 1 | 1 | 1 |
| r3d_134 | video (R3D appearance | 1 | 1 | 1 |
| r3d_135 | video (R3D appearance | 1 | 1 | 1 |
| r3d_136 | video (R3D appearance | 1 | 1 | 1 |
| r3d_137 | video (R3D appearance | 1 | 1 | 1 |
| r3d_138 | video (R3D appearance | 1 | 1 | 1 |
| r3d_139 | video (R3D appearance | 1 | 1 | 1 |
| r3d_140 | video (R3D appearance | 1 | 1 | 1 |
| r3d_141 | video (R3D appearance | 1 | 1 | 1 |
| r3d_142 | video (R3D appearance | 1 | 1 | 1 |
| r3d_143 | video (R3D appearance | 1 | 1 | 1 |
| r3d_144 | video (R3D appearance | 1 | 1 | 1 |
| r3d_145 | video (R3D appearance | 1 | 1 | 1 |
| r3d_146 | video (R3D appearance | 1 | 1 | 1 |
| r3d_147 | video (R3D appearance | 1 | 1 | 1 |
| r3d_148 | video (R3D appearance | 1 | 1 | 1 |
| r3d_149 | video (R3D appearance | 1 | 1 | 1 |
| r3d_150 | video (R3D appearance | 1 | 1 | 1 |
| r3d_151 | video (R3D appearance | 1 | 1 | 1 |
| r3d_152 | video (R3D appearance | 1 | 1 | 1 |
| r3d_153 | video (R3D appearance | 1 | 1 | 1 |
| r3d_154 | video (R3D appearance | 1 | 1 | 1 |
| r3d_155 | video (R3D appearance | 1 | 1 | 1 |
| r3d_156 | video (R3D appearance | 1 | 1 | 1 |
| r3d_157 | video (R3D appearance | 1 | 1 | 1 |
| r3d_158 | video (R3D appearance | 1 | 1 | 1 |
| r3d_159 | video (R3D appearance | 1 | 1 | 1 |
| r3d_160 | video (R3D appearance | 1 | 1 | 1 |
| r3d_161 | video (R3D appearance | 1 | 1 | 1 |

| | | | | |
|---|---|---|---|---|
| r3d_162 | video (R3D appearance | 1 | 1 | 1 |
| r3d_163 | video (R3D appearance | 1 | 1 | 1 |
| r3d_164 | video (R3D appearance | 1 | 1 | 1 |
| r3d_165 | video (R3D appearance | 1 | 1 | 1 |
| r3d_166 | video (R3D appearance | 1 | 1 | 1 |
| r3d_167 | video (R3D appearance | 1 | 1 | 1 |
| r3d_168 | video (R3D appearance | 1 | 1 | 1 |
| r3d_169 | video (R3D appearance | 1 | 1 | 1 |
| r3d_170 | video (R3D appearance | 1 | 1 | 1 |
| r3d_171 | video (R3D appearance | 1 | 1 | 1 |
| r3d_172 | video (R3D appearance | 1 | 1 | 1 |
| r3d_173 | video (R3D appearance | 1 | 1 | 1 |
| r3d_174 | video (R3D appearance | 1 | 1 | 1 |
| r3d_175 | video (R3D appearance | 1 | 1 | 1 |
| r3d_176 | video (R3D appearance | 1 | 1 | 1 |
| r3d_177 | video (R3D appearance | 1 | 1 | 1 |
| r3d_178 | video (R3D appearance | 1 | 1 | 1 |
| r3d_179 | video (R3D appearance | 1 | 1 | 1 |
| r3d_180 | video (R3D appearance | 1 | 1 | 1 |
| r3d_181 | video (R3D appearance | 1 | 1 | 1 |
| r3d_182 | video (R3D appearance | 1 | 1 | 1 |
| r3d_183 | video (R3D appearance | 1 | 1 | 1 |
| r3d_184 | video (R3D appearance | 1 | 1 | 1 |
| r3d_185 | video (R3D appearance | 1 | 1 | 1 |
| r3d_186 | video (R3D appearance | 1 | 1 | 1 |
| r3d_187 | video (R3D appearance | 1 | 1 | 1 |
| r3d_188 | video (R3D appearance | 1 | 1 | 1 |
| r3d_189 | video (R3D appearance | 1 | 1 | 1 |
| r3d_190 | video (R3D appearance | 1 | 1 | 1 |
| r3d_191 | video (R3D appearance | 1 | 1 | 1 |
| r3d_192 | video (R3D appearance | 1 | 1 | 1 |
| r3d_193 | video (R3D appearance | 1 | 1 | 1 |
| r3d_194 | video (R3D appearance | 1 | 1 | 1 |
| r3d_195 | video (R3D appearance | 1 | 1 | 1 |
| r3d_196 | video (R3D appearance | 1 | 1 | 1 |
| r3d_197 | video (R3D appearance | 1 | 1 | 1 |
| r3d_198 | video (R3D appearance | 1 | 1 | 1 |
| r3d_199 | video (R3D appearance | 1 | 1 | 1 |
| r3d_200 | video (R3D appearance | 1 | 1 | 1 |
| r3d_201 | video (R3D appearance | 1 | 1 | 1 |
| r3d_202 | video (R3D appearance | 1 | 1 | 1 |
| r3d_203 | video (R3D appearance | 1 | 1 | 1 |
| r3d_204 | video (R3D appearance | 1 | 1 | 1 |
| r3d_205 | video (R3D appearance | 1 | 1 | 1 |

| | | | | |
|---|---|---|---|---|
| r3d_206 | video (R3D appearance | 1 | 1 | 1 |
| r3d_207 | video (R3D appearance | 1 | 1 | 1 |
| r3d_208 | video (R3D appearance | 1 | 1 | 1 |
| r3d_209 | video (R3D appearance | 1 | 1 | 1 |
| r3d_210 | video (R3D appearance | 1 | 1 | 1 |
| r3d_211 | video (R3D appearance | 1 | 1 | 1 |
| r3d_212 | video (R3D appearance | 1 | 1 | 1 |
| r3d_213 | video (R3D appearance | 1 | 1 | 1 |
| r3d_214 | video (R3D appearance | 1 | 1 | 1 |
| r3d_215 | video (R3D appearance | 1 | 1 | 1 |
| r3d_216 | video (R3D appearance | 1 | 1 | 1 |
| r3d_217 | video (R3D appearance | 1 | 1 | 1 |
| r3d_218 | video (R3D appearance | 1 | 1 | 1 |
| r3d_219 | video (R3D appearance | 1 | 1 | 1 |
| r3d_220 | video (R3D appearance | 1 | 1 | 1 |
| r3d_221 | video (R3D appearance | 1 | 1 | 1 |
| r3d_222 | video (R3D appearance | 1 | 1 | 1 |
| r3d_223 | video (R3D appearance | 1 | 1 | 1 |
| r3d_224 | video (R3D appearance | 1 | 1 | 1 |
| r3d_225 | video (R3D appearance | 1 | 1 | 1 |
| r3d_226 | video (R3D appearance | 1 | 1 | 1 |
| r3d_227 | video (R3D appearance | 1 | 1 | 1 |
| r3d_228 | video (R3D appearance | 1 | 1 | 1 |
| r3d_229 | video (R3D appearance | 1 | 1 | 1 |
| r3d_230 | video (R3D appearance | 1 | 1 | 1 |
| r3d_231 | video (R3D appearance | 1 | 1 | 1 |
| r3d_232 | video (R3D appearance | 1 | 1 | 1 |
| r3d_233 | video (R3D appearance | 1 | 1 | 1 |
| r3d_234 | video (R3D appearance | 1 | 1 | 1 |
| r3d_235 | video (R3D appearance | 1 | 1 | 1 |
| r3d_236 | video (R3D appearance | 1 | 1 | 1 |
| r3d_237 | video (R3D appearance | 1 | 1 | 1 |
| r3d_238 | video (R3D appearance | 1 | 1 | 1 |
| r3d_239 | video (R3D appearance | 1 | 1 | 1 |
| r3d_240 | video (R3D appearance | 1 | 1 | 1 |
| r3d_241 | video (R3D appearance | 1 | 1 | 1 |
| r3d_242 | video (R3D appearance | 1 | 1 | 1 |
| r3d_243 | video (R3D appearance | 1 | 1 | 1 |
| r3d_244 | video (R3D appearance | 1 | 1 | 1 |
| r3d_245 | video (R3D appearance | 1 | 1 | 1 |
| r3d_246 | video (R3D appearance | 1 | 1 | 1 |
| r3d_247 | video (R3D appearance | 1 | 1 | 1 |
| r3d_248 | video (R3D appearance | 1 | 1 | 1 |
| r3d_249 | video (R3D appearance | 1 | 1 | 1 |

| | | | | |
|---|---|---|---|---|
| r3d_250 | video (R3D appearance | 1 | 1 | 1 |
| r3d_251 | video (R3D appearance | 1 | 1 | 1 |
| r3d_252 | video (R3D appearance | 1 | 1 | 1 |
| r3d_253 | video (R3D appearance | 1 | 1 | 1 |
| r3d_254 | video (R3D appearance | 1 | 1 | 1 |
| r3d_255 | video (R3D appearance | 1 | 1 | 1 |
| r3d_256 | video (R3D appearance | 1 | 1 | 1 |
| r3d_257 | video (R3D appearance | 1 | 1 | 1 |
| r3d_258 | video (R3D appearance | 1 | 1 | 1 |
| r3d_259 | video (R3D appearance | 1 | 1 | 1 |
| r3d_260 | video (R3D appearance | 1 | 1 | 1 |
| r3d_261 | video (R3D appearance | 1 | 1 | 1 |
| r3d_262 | video (R3D appearance | 1 | 1 | 1 |
| r3d_263 | video (R3D appearance | 1 | 1 | 1 |
| r3d_264 | video (R3D appearance | 1 | 1 | 1 |
| r3d_265 | video (R3D appearance | 1 | 1 | 1 |
| r3d_266 | video (R3D appearance | 1 | 1 | 1 |
| r3d_267 | video (R3D appearance | 1 | 1 | 1 |
| r3d_268 | video (R3D appearance | 1 | 1 | 1 |
| r3d_269 | video (R3D appearance | 1 | 1 | 1 |
| r3d_270 | video (R3D appearance | 1 | 1 | 1 |
| r3d_271 | video (R3D appearance | 1 | 1 | 1 |
| r3d_272 | video (R3D appearance | 1 | 1 | 1 |
| r3d_273 | video (R3D appearance | 1 | 1 | 1 |
| r3d_274 | video (R3D appearance | 1 | 1 | 1 |
| r3d_275 | video (R3D appearance | 1 | 1 | 1 |
| r3d_276 | video (R3D appearance | 1 | 1 | 1 |
| r3d_277 | video (R3D appearance | 1 | 1 | 1 |
| r3d_278 | video (R3D appearance | 1 | 1 | 1 |
| r3d_279 | video (R3D appearance | 1 | 1 | 1 |
| r3d_280 | video (R3D appearance | 1 | 1 | 1 |
| r3d_281 | video (R3D appearance | 1 | 1 | 1 |
| r3d_282 | video (R3D appearance | 1 | 1 | 1 |
| r3d_283 | video (R3D appearance | 1 | 1 | 1 |
| r3d_284 | video (R3D appearance | 1 | 1 | 1 |
| r3d_285 | video (R3D appearance | 1 | 1 | 1 |
| r3d_286 | video (R3D appearance | 1 | 1 | 1 |
| r3d_287 | video (R3D appearance | 1 | 1 | 1 |
| r3d_288 | video (R3D appearance | 1 | 1 | 1 |
| r3d_289 | video (R3D appearance | 1 | 1 | 1 |
| r3d_290 | video (R3D appearance | 1 | 1 | 1 |
| r3d_291 | video (R3D appearance | 1 | 1 | 1 |
| r3d_292 | video (R3D appearance | 1 | 1 | 1 |
| r3d_293 | video (R3D appearance | 1 | 1 | 1 |

| | | | | |
|---|---|---|---|---|
| r3d_294 | video (R3D appearance | 1 | 1 | 1 |
| r3d_295 | video (R3D appearance | 1 | 1 | 1 |
| r3d_296 | video (R3D appearance | 1 | 1 | 1 |
| r3d_297 | video (R3D appearance | 1 | 1 | 1 |
| r3d_298 | video (R3D appearance | 1 | 1 | 1 |
| r3d_299 | video (R3D appearance | 1 | 1 | 1 |
| r3d_300 | video (R3D appearance | 1 | 1 | 1 |
| r3d_301 | video (R3D appearance | 1 | 1 | 1 |
| r3d_302 | video (R3D appearance | 1 | 1 | 1 |
| r3d_303 | video (R3D appearance | 1 | 1 | 1 |
| r3d_304 | video (R3D appearance | 1 | 1 | 1 |
| r3d_305 | video (R3D appearance | 1 | 1 | 1 |
| r3d_306 | video (R3D appearance | 1 | 1 | 1 |
| r3d_307 | video (R3D appearance | 1 | 1 | 1 |
| r3d_308 | video (R3D appearance | 1 | 1 | 1 |
| r3d_309 | video (R3D appearance | 1 | 1 | 1 |
| r3d_310 | video (R3D appearance | 1 | 1 | 1 |
| r3d_311 | video (R3D appearance | 1 | 1 | 1 |
| r3d_312 | video (R3D appearance | 1 | 1 | 1 |
| r3d_313 | video (R3D appearance | 1 | 1 | 1 |
| r3d_314 | video (R3D appearance | 1 | 1 | 1 |
| r3d_315 | video (R3D appearance | 1 | 1 | 1 |
| r3d_316 | video (R3D appearance | 1 | 1 | 1 |
| r3d_317 | video (R3D appearance | 1 | 1 | 1 |
| r3d_318 | video (R3D appearance | 1 | 1 | 1 |
| r3d_319 | video (R3D appearance | 1 | 1 | 1 |
| r3d_320 | video (R3D appearance | 1 | 1 | 1 |
| r3d_321 | video (R3D appearance | 1 | 1 | 1 |
| r3d_322 | video (R3D appearance | 1 | 1 | 1 |
| r3d_323 | video (R3D appearance | 1 | 1 | 1 |
| r3d_324 | video (R3D appearance | 1 | 1 | 1 |
| r3d_325 | video (R3D appearance | 1 | 1 | 1 |
| r3d_326 | video (R3D appearance | 1 | 1 | 1 |
| r3d_327 | video (R3D appearance | 1 | 1 | 1 |
| r3d_328 | video (R3D appearance | 1 | 1 | 1 |
| r3d_329 | video (R3D appearance | 1 | 1 | 1 |
| r3d_330 | video (R3D appearance | 1 | 1 | 1 |
| r3d_331 | video (R3D appearance | 1 | 1 | 1 |
| r3d_332 | video (R3D appearance | 1 | 1 | 1 |
| r3d_333 | video (R3D appearance | 1 | 1 | 1 |
| r3d_334 | video (R3D appearance | 1 | 1 | 1 |
| r3d_335 | video (R3D appearance | 1 | 1 | 1 |
| r3d_336 | video (R3D appearance | 1 | 1 | 1 |
| r3d_337 | video (R3D appearance | 1 | 1 | 1 |

| | | | | |
|---|---|---|---|---|
| r3d_338 | video (R3D appearance | 1 | 1 | 1 |
| r3d_339 | video (R3D appearance | 1 | 1 | 1 |
| r3d_340 | video (R3D appearance | 1 | 1 | 1 |
| r3d_341 | video (R3D appearance | 1 | 1 | 1 |
| r3d_342 | video (R3D appearance | 1 | 1 | 1 |
| r3d_343 | video (R3D appearance | 1 | 1 | 1 |
| r3d_344 | video (R3D appearance | 1 | 1 | 1 |
| r3d_345 | video (R3D appearance | 1 | 1 | 1 |
| r3d_346 | video (R3D appearance | 1 | 1 | 1 |
| r3d_347 | video (R3D appearance | 1 | 1 | 1 |
| r3d_348 | video (R3D appearance | 1 | 1 | 1 |
| r3d_349 | video (R3D appearance | 1 | 1 | 1 |
| r3d_350 | video (R3D appearance | 1 | 1 | 1 |
| r3d_351 | video (R3D appearance | 1 | 1 | 1 |
| r3d_352 | video (R3D appearance | 1 | 1 | 1 |
| r3d_353 | video (R3D appearance | 1 | 1 | 1 |
| r3d_354 | video (R3D appearance | 1 | 1 | 1 |
| r3d_355 | video (R3D appearance | 1 | 1 | 1 |
| r3d_356 | video (R3D appearance | 1 | 1 | 1 |
| r3d_357 | video (R3D appearance | 1 | 1 | 1 |
| r3d_358 | video (R3D appearance | 1 | 1 | 1 |
| r3d_359 | video (R3D appearance | 1 | 1 | 1 |
| r3d_360 | video (R3D appearance | 1 | 1 | 1 |
| r3d_361 | video (R3D appearance | 1 | 1 | 1 |
| r3d_362 | video (R3D appearance | 1 | 1 | 1 |
| r3d_363 | video (R3D appearance | 1 | 1 | 1 |
| r3d_364 | video (R3D appearance | 1 | 1 | 1 |
| r3d_365 | video (R3D appearance | 1 | 1 | 1 |
| r3d_366 | video (R3D appearance | 1 | 1 | 1 |
| r3d_367 | video (R3D appearance | 1 | 1 | 1 |
| r3d_368 | video (R3D appearance | 1 | 1 | 1 |
| r3d_369 | video (R3D appearance | 1 | 1 | 1 |
| r3d_370 | video (R3D appearance | 1 | 1 | 1 |
| r3d_371 | video (R3D appearance | 1 | 1 | 1 |
| r3d_372 | video (R3D appearance | 1 | 1 | 1 |
| r3d_373 | video (R3D appearance | 1 | 1 | 1 |
| r3d_374 | video (R3D appearance | 1 | 1 | 1 |
| r3d_375 | video (R3D appearance | 1 | 1 | 1 |
| r3d_376 | video (R3D appearance | 1 | 1 | 1 |
| r3d_377 | video (R3D appearance | 1 | 1 | 1 |
| r3d_378 | video (R3D appearance | 1 | 1 | 1 |
| r3d_379 | video (R3D appearance | 1 | 1 | 1 |
| r3d_380 | video (R3D appearance | 1 | 1 | 1 |
| r3d_381 | video (R3D appearance | 1 | 1 | 1 |

| | | | | |
|---|---|---|---|---|
| r3d_382 | video (R3D appearance | 1 | 1 | 1 |
| r3d_383 | video (R3D appearance | 1 | 1 | 1 |
| r3d_384 | video (R3D appearance | 1 | 1 | 1 |
| r3d_385 | video (R3D appearance | 1 | 1 | 1 |
| r3d_386 | video (R3D appearance | 1 | 1 | 1 |
| r3d_387 | video (R3D appearance | 1 | 1 | 1 |
| r3d_388 | video (R3D appearance | 1 | 1 | 1 |
| r3d_389 | video (R3D appearance | 1 | 1 | 1 |
| r3d_390 | video (R3D appearance | 1 | 1 | 1 |
| r3d_391 | video (R3D appearance | 1 | 1 | 1 |
| r3d_392 | video (R3D appearance | 1 | 1 | 1 |
| r3d_393 | video (R3D appearance | 1 | 1 | 1 |
| r3d_394 | video (R3D appearance | 1 | 1 | 1 |
| r3d_395 | video (R3D appearance | 1 | 1 | 1 |
| r3d_396 | video (R3D appearance | 1 | 1 | 1 |
| r3d_397 | video (R3D appearance | 1 | 1 | 1 |
| r3d_398 | video (R3D appearance | 1 | 1 | 1 |
| r3d_399 | video (R3D appearance | 1 | 1 | 1 |
| r3d_400 | video (R3D appearance | 1 | 1 | 1 |
| r3d_401 | video (R3D appearance | 1 | 1 | 1 |
| r3d_402 | video (R3D appearance | 1 | 1 | 1 |
| r3d_403 | video (R3D appearance | 1 | 1 | 1 |
| r3d_404 | video (R3D appearance | 1 | 1 | 1 |
| r3d_405 | video (R3D appearance | 1 | 1 | 1 |
| r3d_406 | video (R3D appearance | 1 | 1 | 1 |
| r3d_407 | video (R3D appearance | 1 | 1 | 1 |
| r3d_408 | video (R3D appearance | 1 | 1 | 1 |
| r3d_409 | video (R3D appearance | 1 | 1 | 1 |
| r3d_410 | video (R3D appearance | 1 | 1 | 1 |
| r3d_411 | video (R3D appearance | 1 | 1 | 1 |
| r3d_412 | video (R3D appearance | 1 | 1 | 1 |
| r3d_413 | video (R3D appearance | 1 | 1 | 1 |
| r3d_414 | video (R3D appearance | 1 | 1 | 1 |
| r3d_415 | video (R3D appearance | 1 | 1 | 1 |
| r3d_416 | video (R3D appearance | 1 | 1 | 1 |
| r3d_417 | video (R3D appearance | 1 | 1 | 1 |
| r3d_418 | video (R3D appearance | 1 | 1 | 1 |
| r3d_419 | video (R3D appearance | 1 | 1 | 1 |
| r3d_420 | video (R3D appearance | 1 | 1 | 1 |
| r3d_421 | video (R3D appearance | 1 | 1 | 1 |
| r3d_422 | video (R3D appearance | 1 | 1 | 1 |
| r3d_423 | video (R3D appearance | 1 | 1 | 1 |
| r3d_424 | video (R3D appearance | 1 | 1 | 1 |
| r3d_425 | video (R3D appearance | 1 | 1 | 1 |

| | | | | |
|---|---|---|---|---|
| r3d_426 | video (R3D appearance | 1 | 1 | 1 |
| r3d_427 | video (R3D appearance | 1 | 1 | 1 |
| r3d_428 | video (R3D appearance | 1 | 1 | 1 |
| r3d_429 | video (R3D appearance | 1 | 1 | 1 |
| r3d_430 | video (R3D appearance | 1 | 1 | 1 |
| r3d_431 | video (R3D appearance | 1 | 1 | 1 |
| r3d_432 | video (R3D appearance | 1 | 1 | 1 |
| r3d_433 | video (R3D appearance | 1 | 1 | 1 |
| r3d_434 | video (R3D appearance | 1 | 1 | 1 |
| r3d_435 | video (R3D appearance | 1 | 1 | 1 |
| r3d_436 | video (R3D appearance | 1 | 1 | 1 |
| r3d_437 | video (R3D appearance | 1 | 1 | 1 |
| r3d_438 | video (R3D appearance | 1 | 1 | 1 |
| r3d_439 | video (R3D appearance | 1 | 1 | 1 |
| r3d_440 | video (R3D appearance | 1 | 1 | 1 |
| r3d_441 | video (R3D appearance | 1 | 1 | 1 |
| r3d_442 | video (R3D appearance | 1 | 1 | 1 |
| r3d_443 | video (R3D appearance | 1 | 1 | 1 |
| r3d_444 | video (R3D appearance | 1 | 1 | 1 |
| r3d_445 | video (R3D appearance | 1 | 1 | 1 |
| r3d_446 | video (R3D appearance | 1 | 1 | 1 |
| r3d_447 | video (R3D appearance | 1 | 1 | 1 |
| r3d_448 | video (R3D appearance | 1 | 1 | 1 |
| r3d_449 | video (R3D appearance | 1 | 1 | 1 |
| r3d_450 | video (R3D appearance | 1 | 1 | 1 |
| r3d_451 | video (R3D appearance | 1 | 1 | 1 |
| r3d_452 | video (R3D appearance | 1 | 1 | 1 |
| r3d_453 | video (R3D appearance | 1 | 1 | 1 |
| r3d_454 | video (R3D appearance | 1 | 1 | 1 |
| r3d_455 | video (R3D appearance | 1 | 1 | 1 |
| r3d_456 | video (R3D appearance | 1 | 1 | 1 |
| r3d_457 | video (R3D appearance | 1 | 1 | 1 |
| r3d_458 | video (R3D appearance | 1 | 1 | 1 |
| r3d_459 | video (R3D appearance | 1 | 1 | 1 |
| r3d_460 | video (R3D appearance | 1 | 1 | 1 |
| r3d_461 | video (R3D appearance | 1 | 1 | 1 |
| r3d_462 | video (R3D appearance | 1 | 1 | 1 |
| r3d_463 | video (R3D appearance | 1 | 1 | 1 |
| r3d_464 | video (R3D appearance | 1 | 1 | 1 |
| r3d_465 | video (R3D appearance | 1 | 1 | 1 |
| r3d_466 | video (R3D appearance | 1 | 1 | 1 |
| r3d_467 | video (R3D appearance | 1 | 1 | 1 |
| r3d_468 | video (R3D appearance | 1 | 1 | 1 |
| r3d_469 | video (R3D appearance | 1 | 1 | 1 |

| | | | | |
|---|---|---|---|---|
| r3d_470 | video (R3D appearance | 1 | 1 | 1 |
| r3d_471 | video (R3D appearance | 1 | 1 | 1 |
| r3d_472 | video (R3D appearance | 1 | 1 | 1 |
| r3d_473 | video (R3D appearance | 1 | 1 | 1 |
| r3d_474 | video (R3D appearance | 1 | 1 | 1 |
| r3d_475 | video (R3D appearance | 1 | 1 | 1 |
| r3d_476 | video (R3D appearance | 1 | 1 | 1 |
| r3d_477 | video (R3D appearance | 1 | 1 | 1 |
| r3d_478 | video (R3D appearance | 1 | 1 | 1 |
| r3d_479 | video (R3D appearance | 1 | 1 | 1 |
| r3d_480 | video (R3D appearance | 1 | 1 | 1 |
| r3d_481 | video (R3D appearance | 1 | 1 | 1 |
| r3d_482 | video (R3D appearance | 1 | 1 | 1 |
| r3d_483 | video (R3D appearance | 1 | 1 | 1 |
| r3d_484 | video (R3D appearance | 1 | 1 | 1 |
| r3d_485 | video (R3D appearance | 1 | 1 | 1 |
| r3d_486 | video (R3D appearance | 1 | 1 | 1 |
| r3d_487 | video (R3D appearance | 1 | 1 | 1 |
| r3d_488 | video (R3D appearance | 1 | 1 | 1 |
| r3d_489 | video (R3D appearance | 1 | 1 | 1 |
| r3d_490 | video (R3D appearance | 1 | 1 | 1 |
| r3d_491 | video (R3D appearance | 1 | 1 | 1 |
| r3d_492 | video (R3D appearance | 1 | 1 | 1 |
| r3d_493 | video (R3D appearance | 1 | 1 | 1 |
| r3d_494 | video (R3D appearance | 1 | 1 | 1 |
| r3d_495 | video (R3D appearance | 1 | 1 | 1 |
| r3d_496 | video (R3D appearance | 1 | 1 | 1 |
| r3d_497 | video (R3D appearance | 1 | 1 | 1 |
| r3d_498 | video (R3D appearance | 1 | 1 | 1 |
| r3d_499 | video (R3D appearance | 1 | 1 | 1 |
| r3d_500 | video (R3D appearance | 1 | 1 | 1 |
| r3d_501 | video (R3D appearance | 1 | 1 | 1 |
| r3d_502 | video (R3D appearance | 1 | 1 | 1 |
| r3d_503 | video (R3D appearance | 1 | 1 | 1 |
| r3d_504 | video (R3D appearance | 1 | 1 | 1 |
| r3d_505 | video (R3D appearance | 1 | 1 | 1 |
| r3d_506 | video (R3D appearance | 1 | 1 | 1 |
| r3d_507 | video (R3D appearance | 1 | 1 | 1 |
| r3d_508 | video (R3D appearance | 1 | 1 | 1 |
| r3d_509 | video (R3D appearance | 1 | 1 | 1 |
| r3d_510 | video (R3D appearance | 1 | 1 | 1 |
| r3d_511 | video (R3D appearance | 1 | 1 | 1 |
| right_body_acceleration_energy | kinematics | 1 | 0 | 0 |
| right_body_acceleration_max | kinematics | 1 | 0 | 0 |

| | | | | |
|---|---|---|---|---|
| right_body_acceleration_mean | kinematics | 1 | 0 | 0 |
| right_body_acceleration_median | kinematics | 1 | 0 | 0 |
| right_body_acceleration_p10 | kinematics | 1 | 0 | 0 |
| right_body_acceleration_p90 | kinematics | 1 | 0 | 0 |
| right_body_acceleration_periodicity | kinematics | 1 | 0 | 0 |
| right_body_acceleration_std | kinematics | 1 | 0 | 0 |
| right_body_forward_velocity_energy | kinematics | 1 | 0 | 0 |
| right_body_forward_velocity_max | kinematics | 1 | 0 | 0 |
| right_body_forward_velocity_mean | kinematics | 1 | 0 | 0 |
| right_body_forward_velocity_median | kinematics | 1 | 0 | 0 |
| right_body_forward_velocity_p10 | kinematics | 1 | 0 | 0 |
| right_body_forward_velocity_p90 | kinematics | 1 | 0 | 0 |
| right_body_forward_velocity_periodicity | kinematics | 1 | 0 | 0 |
| right_body_forward_velocity_std | kinematics | 1 | 0 | 0 |
| right_body_jerk_energy | kinematics | 1 | 0 | 0 |
| right_body_jerk_max | kinematics | 1 | 0 | 0 |
| right_body_jerk_mean | kinematics | 1 | 0 | 0 |
| right_body_jerk_median | kinematics | 1 | 0 | 0 |
| right_body_jerk_p10 | kinematics | 1 | 0 | 0 |
| right_body_jerk_p90 | kinematics | 1 | 0 | 0 |
| right_body_jerk_periodicity | kinematics | 1 | 0 | 0 |
| right_body_jerk_std | kinematics | 1 | 0 | 0 |
| right_body_lateral_velocity_energy | kinematics | 1 | 0 | 0 |
| right_body_lateral_velocity_max | kinematics | 1 | 0 | 0 |
| right_body_lateral_velocity_mean | kinematics | 1 | 0 | 0 |
| right_body_lateral_velocity_median | kinematics | 1 | 0 | 0 |
| right_body_lateral_velocity_p10 | kinematics | 1 | 0 | 0 |
| right_body_lateral_velocity_p90 | kinematics | 1 | 0 | 0 |
| right_body_lateral_velocity_periodicity | kinematics | 1 | 0 | 0 |
| right_body_lateral_velocity_std | kinematics | 1 | 0 | 0 |
| right_body_speed_energy | kinematics | 1 | 0 | 0 |
| right_body_speed_max | kinematics | 1 | 0 | 0 |
| right_body_speed_mean | kinematics | 1 | 0 | 0 |
| right_body_speed_median | kinematics | 1 | 0 | 0 |
| right_body_speed_p10 | kinematics | 1 | 0 | 0 |
| right_body_speed_p90 | kinematics | 1 | 0 | 0 |
| right_body_speed_periodicity | kinematics | 1 | 0 | 0 |
| right_body_speed_std | kinematics | 1 | 0 | 0 |
| right_body_velocity_x_energy | kinematics | 1 | 0 | 0 |
| right_body_velocity_x_max | kinematics | 1 | 0 | 0 |
| right_body_velocity_x_mean | kinematics | 1 | 0 | 0 |
| right_body_velocity_x_median | kinematics | 1 | 0 | 0 |
| right_body_velocity_x_p10 | kinematics | 1 | 0 | 0 |
| right_body_velocity_x_p90 | kinematics | 1 | 0 | 0 |

| | | | | |
|---|---|---|---|---|
| right_body_velocity_x_periodicity | kinematics | 1 | 0 | 0 |
| right_body_velocity_x_std | kinematics | 1 | 0 | 0 |
| right_body_velocity_y_energy | kinematics | 1 | 0 | 0 |
| right_body_velocity_y_max | kinematics | 1 | 0 | 0 |
| right_body_velocity_y_mean | kinematics | 1 | 0 | 0 |
| right_body_velocity_y_median | kinematics | 1 | 0 | 0 |
| right_body_velocity_y_p10 | kinematics | 1 | 0 | 0 |
| right_body_velocity_y_p90 | kinematics | 1 | 0 | 0 |
| right_body_velocity_y_periodicity | kinematics | 1 | 0 | 0 |
| right_body_velocity_y_std | kinematics | 1 | 0 | 0 |
| right_ear_acceleration_energy | kinematics | 1 | 0 | 0 |
| right_ear_acceleration_max | kinematics | 1 | 0 | 0 |
| right_ear_acceleration_mean | kinematics | 1 | 0 | 0 |
| right_ear_acceleration_median | kinematics | 1 | 0 | 0 |
| right_ear_acceleration_p10 | kinematics | 1 | 0 | 0 |
| right_ear_acceleration_p90 | kinematics | 1 | 0 | 0 |
| right_ear_acceleration_periodicity | kinematics | 1 | 0 | 0 |
| right_ear_acceleration_std | kinematics | 1 | 0 | 0 |
| right_ear_forward_velocity_energy | kinematics | 1 | 0 | 0 |
| right_ear_forward_velocity_max | kinematics | 1 | 0 | 0 |
| right_ear_forward_velocity_mean | kinematics | 1 | 0 | 0 |
| right_ear_forward_velocity_median | kinematics | 1 | 0 | 0 |
| right_ear_forward_velocity_p10 | kinematics | 1 | 0 | 0 |
| right_ear_forward_velocity_p90 | kinematics | 1 | 0 | 0 |
| right_ear_forward_velocity_periodicity | kinematics | 1 | 0 | 0 |
| right_ear_forward_velocity_std | kinematics | 1 | 0 | 0 |
| right_ear_jerk_energy | kinematics | 1 | 0 | 0 |
| right_ear_jerk_max | kinematics | 1 | 0 | 0 |
| right_ear_jerk_mean | kinematics | 1 | 0 | 0 |
| right_ear_jerk_median | kinematics | 1 | 0 | 0 |
| right_ear_jerk_p10 | kinematics | 1 | 0 | 0 |
| right_ear_jerk_p90 | kinematics | 1 | 0 | 0 |
| right_ear_jerk_periodicity | kinematics | 1 | 0 | 0 |
| right_ear_jerk_std | kinematics | 1 | 0 | 0 |
| right_ear_lateral_velocity_energy | kinematics | 1 | 0 | 0 |
| right_ear_lateral_velocity_max | kinematics | 1 | 0 | 0 |
| right_ear_lateral_velocity_mean | kinematics | 1 | 0 | 0 |
| right_ear_lateral_velocity_median | kinematics | 1 | 0 | 0 |
| right_ear_lateral_velocity_p10 | kinematics | 1 | 0 | 0 |
| right_ear_lateral_velocity_p90 | kinematics | 1 | 0 | 0 |
| right_ear_lateral_velocity_periodicity | kinematics | 1 | 0 | 0 |
| right_ear_lateral_velocity_std | kinematics | 1 | 0 | 0 |
| right_ear_speed_energy | kinematics | 1 | 0 | 0 |
| right_ear_speed_max | kinematics | 1 | 0 | 0 |

| | | | | |
|---|---|---|---|---|
| right_ear_speed_mean | kinematics | 1 | 0 | 0 |
| right_ear_speed_median | kinematics | 1 | 0 | 0 |
| right_ear_speed_p10 | kinematics | 1 | 0 | 0 |
| right_ear_speed_p90 | kinematics | 1 | 0 | 0 |
| right_ear_speed_periodicity | kinematics | 1 | 0 | 0 |
| right_ear_speed_std | kinematics | 1 | 0 | 0 |
| right_ear_velocity_x_energy | kinematics | 1 | 0 | 0 |
| right_ear_velocity_x_max | kinematics | 1 | 0 | 0 |
| right_ear_velocity_x_mean | kinematics | 1 | 0 | 0 |
| right_ear_velocity_x_median | kinematics | 1 | 0 | 0 |
| right_ear_velocity_x_p10 | kinematics | 1 | 0 | 0 |
| right_ear_velocity_x_p90 | kinematics | 1 | 0 | 0 |
| right_ear_velocity_x_periodicity | kinematics | 1 | 0 | 0 |
| right_ear_velocity_x_std | kinematics | 1 | 0 | 0 |
| right_ear_velocity_y_energy | kinematics | 1 | 0 | 0 |
| right_ear_velocity_y_max | kinematics | 1 | 0 | 0 |
| right_ear_velocity_y_mean | kinematics | 1 | 0 | 0 |
| right_ear_velocity_y_median | kinematics | 1 | 0 | 0 |
| right_ear_velocity_y_p10 | kinematics | 1 | 0 | 0 |
| right_ear_velocity_y_p90 | kinematics | 1 | 0 | 0 |
| right_ear_velocity_y_periodicity | kinematics | 1 | 0 | 0 |
| right_ear_velocity_y_std | kinematics | 1 | 0 | 0 |
| roi_1_present_energy | context (ROI/target) | 1 | 1 | 1 |
| roi_1_present_max | context (ROI/target) | 1 | 1 | 1 |
| roi_1_present_mean | context (ROI/target) | 1 | 1 | 1 |
| roi_1_present_median | context (ROI/target) | 1 | 1 | 1 |
| roi_1_present_p10 | context (ROI/target) | 1 | 1 | 1 |
| roi_1_present_p90 | context (ROI/target) | 1 | 1 | 1 |
| roi_1_present_periodicity | context (ROI/target) | 1 | 1 | 1 |
| roi_1_present_std | context (ROI/target) | 1 | 1 | 1 |
| roi_2_present_energy | context (ROI/target) | 1 | 0 | 0 |
| roi_2_present_max | context (ROI/target) | 1 | 0 | 0 |
| roi_2_present_mean | context (ROI/target) | 1 | 0 | 0 |
| roi_2_present_median | context (ROI/target) | 1 | 0 | 0 |
| roi_2_present_p10 | context (ROI/target) | 1 | 0 | 0 |
| roi_2_present_p90 | context (ROI/target) | 1 | 0 | 0 |
| roi_2_present_periodicity | context (ROI/target) | 1 | 0 | 0 |
| roi_2_present_std | context (ROI/target) | 1 | 0 | 0 |
| tail_base_acceleration_energy | kinematics | 1 | 1 | 1 |
| tail_base_acceleration_max | kinematics | 1 | 1 | 1 |
| tail_base_acceleration_mean | kinematics | 1 | 1 | 1 |
| tail_base_acceleration_median | kinematics | 1 | 1 | 1 |
| tail_base_acceleration_p10 | kinematics | 1 | 1 | 1 |
| tail_base_acceleration_p90 | kinematics | 1 | 1 | 1 |

| | | | | |
|---|---|---|---|---|
| tail_base_acceleration_periodicity | kinematics | 1 | 1 | 1 |
| tail_base_acceleration_std | kinematics | 1 | 1 | 1 |
| tail_base_forward_velocity_energy | kinematics | 1 | 1 | 1 |
| tail_base_forward_velocity_max | kinematics | 1 | 1 | 1 |
| tail_base_forward_velocity_mean | kinematics | 1 | 1 | 1 |
| tail_base_forward_velocity_median | kinematics | 1 | 1 | 1 |
| tail_base_forward_velocity_p10 | kinematics | 1 | 1 | 1 |
| tail_base_forward_velocity_p90 | kinematics | 1 | 1 | 1 |
| tail_base_forward_velocity_periodicity | kinematics | 1 | 1 | 1 |
| tail_base_forward_velocity_std | kinematics | 1 | 1 | 1 |
| tail_base_jerk_energy | kinematics | 1 | 1 | 1 |
| tail_base_jerk_max | kinematics | 1 | 1 | 1 |
| tail_base_jerk_mean | kinematics | 1 | 1 | 1 |
| tail_base_jerk_median | kinematics | 1 | 1 | 1 |
| tail_base_jerk_p10 | kinematics | 1 | 1 | 1 |
| tail_base_jerk_p90 | kinematics | 1 | 1 | 1 |
| tail_base_jerk_periodicity | kinematics | 1 | 1 | 1 |
| tail_base_jerk_std | kinematics | 1 | 1 | 1 |
| tail_base_lateral_velocity_energy | kinematics | 1 | 1 | 1 |
| tail_base_lateral_velocity_max | kinematics | 1 | 1 | 1 |
| tail_base_lateral_velocity_mean | kinematics | 1 | 1 | 1 |
| tail_base_lateral_velocity_median | kinematics | 1 | 1 | 1 |
| tail_base_lateral_velocity_p10 | kinematics | 1 | 1 | 1 |
| tail_base_lateral_velocity_p90 | kinematics | 1 | 1 | 1 |
| tail_base_lateral_velocity_periodicity | kinematics | 1 | 1 | 1 |
| tail_base_lateral_velocity_std | kinematics | 1 | 1 | 1 |
| tail_base_speed_energy | kinematics | 1 | 1 | 1 |
| tail_base_speed_max | kinematics | 1 | 1 | 1 |
| tail_base_speed_mean | kinematics | 1 | 1 | 1 |
| tail_base_speed_median | kinematics | 1 | 1 | 1 |
| tail_base_speed_p10 | kinematics | 1 | 1 | 1 |
| tail_base_speed_p90 | kinematics | 1 | 1 | 1 |
| tail_base_speed_periodicity | kinematics | 1 | 1 | 1 |
| tail_base_speed_std | kinematics | 1 | 1 | 1 |
| tail_base_velocity_x_energy | kinematics | 1 | 1 | 1 |
| tail_base_velocity_x_max | kinematics | 1 | 1 | 1 |
| tail_base_velocity_x_mean | kinematics | 1 | 1 | 1 |
| tail_base_velocity_x_median | kinematics | 1 | 1 | 1 |
| tail_base_velocity_x_p10 | kinematics | 1 | 1 | 1 |
| tail_base_velocity_x_p90 | kinematics | 1 | 1 | 1 |
| tail_base_velocity_x_periodicity | kinematics | 1 | 1 | 1 |
| tail_base_velocity_x_std | kinematics | 1 | 1 | 1 |
| tail_base_velocity_y_energy | kinematics | 1 | 1 | 1 |
| tail_base_velocity_y_max | kinematics | 1 | 1 | 1 |

| | | | | |
|---|---|---|---|---|
| tail_base_velocity_y_mean | kinematics | 1 | 1 | 1 |
| tail_base_velocity_y_median | kinematics | 1 | 1 | 1 |
| tail_base_velocity_y_p10 | kinematics | 1 | 1 | 1 |
| tail_base_velocity_y_p90 | kinematics | 1 | 1 | 1 |
| tail_base_velocity_y_periodicity | kinematics | 1 | 1 | 1 |
| tail_base_velocity_y_std | kinematics | 1 | 1 | 1 |
| uncertainty_entropy | pose geometry | 1 | 1 | 1 |
| uncertainty_margin | pose geometry | 1 | 1 | 1 |
| uncertainty_score | pose geometry | 1 | 1 | 1 |

| HC | NOR | NSF | OF | TMT | n_assays |
|---|---|---|---|---|---|
| 0 | 0 | 0 | 0 | 1 | 1 |
| 0 | 0 | 0 | 0 | 1 | 1 |
| 0 | 0 | 0 | 0 | 1 | 1 |
| 0 | 0 | 0 | 0 | 1 | 1 |
| 0 | 0 | 0 | 0 | 1 | 1 |
| 0 | 0 | 0 | 0 | 1 | 1 |
| 0 | 0 | 0 | 0 | 1 | 1 |
| 0 | 0 | 0 | 0 | 1 | 1 |
| 0 | 0 | 0 | 0 | 1 | 1 |
| 0 | 0 | 0 | 0 | 1 | 1 |
| 0 | 0 | 0 | 0 | 1 | 1 |
| 0 | 0 | 0 | 0 | 1 | 1 |
| 0 | 0 | 0 | 0 | 1 | 1 |
| 0 | 0 | 0 | 0 | 1 | 1 |
| 0 | 0 | 0 | 0 | 1 | 1 |
| 0 | 0 | 0 | 0 | 1 | 1 |
| 0 | 0 | 0 | 0 | 1 | 1 |
| 0 | 0 | 0 | 0 | 1 | 1 |
| 0 | 0 | 0 | 0 | 1 | 1 |
| 0 | 0 | 0 | 0 | 1 | 1 |
| 0 | 0 | 0 | 0 | 1 | 1 |
| 0 | 0 | 0 | 0 | 1 | 1 |
| 0 | 0 | 0 | 0 | 1 | 1 |
| 0 | 0 | 0 | 0 | 1 | 1 |
| 0 | 0 | 0 | 0 | 1 | 1 |
| 0 | 0 | 0 | 0 | 1 | 1 |
| 0 | 0 | 0 | 0 | 1 | 1 |
| 0 | 0 | 0 | 0 | 1 | 1 |
| 0 | 0 | 0 | 0 | 1 | 1 |
| 0 | 0 | 0 | 0 | 1 | 1 |
| 0 | 0 | 0 | 0 | 1 | 1 |
| 0 | 0 | 0 | 0 | 1 | 1 |
| 0 | 0 | 0 | 0 | 1 | 1 |
| 0 | 0 | 0 | 0 | 1 | 1 |
| 0 | 0 | 0 | 0 | 1 | 1 |
| 0 | 0 | 0 | 0 | 1 | 1 |
| 0 | 0 | 0 | 0 | 1 | 1 |
| 0 | 0 | 0 | 0 | 1 | 1 |
| 0 | 0 | 0 | 0 | 1 | 1 |
| 0 | 0 | 0 | 0 | 1 | 1 |
| 0 | 0 | 0 | 0 | 1 | 1 |
| 0 | 0 | 0 | 0 | 1 | 1 |
| 0 | 0 | 0 | 0 | 1 | 1 |

| | | | | | |
|---|---|---|---|---|---|
| 0 | 0 | 0 | 0 | 1 | 1 |
| 0 | 0 | 0 | 0 | 1 | 1 |
| 0 | 0 | 0 | 0 | 1 | 1 |
| 0 | 0 | 0 | 0 | 1 | 1 |
| 0 | 0 | 0 | 0 | 1 | 1 |
| 0 | 0 | 0 | 0 | 1 | 1 |
| 0 | 0 | 0 | 0 | 1 | 1 |
| 0 | 0 | 0 | 0 | 1 | 1 |
| 0 | 0 | 0 | 0 | 1 | 1 |
| 0 | 0 | 0 | 0 | 1 | 1 |
| 0 | 0 | 0 | 0 | 1 | 1 |
| 0 | 0 | 0 | 0 | 1 | 1 |
| 0 | 0 | 0 | 0 | 1 | 1 |
| 0 | 0 | 0 | 0 | 1 | 1 |
| 0 | 0 | 0 | 0 | 1 | 1 |
| 0 | 0 | 0 | 0 | 1 | 1 |
| 0 | 0 | 0 | 0 | 1 | 1 |
| 0 | 0 | 0 | 0 | 1 | 1 |
| 0 | 0 | 0 | 0 | 1 | 1 |
| 0 | 0 | 0 | 0 | 1 | 1 |
| 0 | 0 | 0 | 0 | 1 | 1 |
| 0 | 0 | 0 | 0 | 1 | 1 |
| 0 | 0 | 0 | 0 | 1 | 1 |
| 0 | 0 | 0 | 0 | 1 | 1 |
| 0 | 0 | 0 | 0 | 1 | 1 |
| 0 | 0 | 0 | 0 | 1 | 1 |
| 0 | 0 | 0 | 0 | 1 | 1 |
| 0 | 0 | 0 | 0 | 1 | 1 |
| 0 | 0 | 0 | 0 | 1 | 1 |
| 0 | 0 | 0 | 0 | 1 | 1 |
| 0 | 0 | 0 | 0 | 1 | 1 |
| 0 | 0 | 0 | 0 | 1 | 1 |
| 0 | 0 | 0 | 0 | 1 | 1 |
| 0 | 0 | 0 | 0 | 1 | 1 |
| 0 | 0 | 0 | 0 | 1 | 1 |
| 0 | 0 | 0 | 0 | 1 | 1 |
| 0 | 0 | 0 | 0 | 1 | 1 |
| 0 | 0 | 0 | 0 | 1 | 1 |
| 0 | 0 | 0 | 0 | 1 | 1 |
| 0 | 0 | 0 | 0 | 1 | 1 |
| 0 | 0 | 0 | 0 | 1 | 1 |
| 0 | 0 | 0 | 0 | 1 | 1 |
| 0 | 0 | 0 | 0 | 1 | 1 |
| 0 | 0 | 0 | 0 | 1 | 1 |

| | | | | | |
|---|---|---|---|---|---|
| 0 | 0 | 0 | 0 | 1 | 1 |
| 0 | 0 | 0 | 0 | 1 | 1 |
| 0 | 0 | 0 | 0 | 1 | 1 |
| 0 | 0 | 0 | 0 | 1 | 1 |
| 0 | 0 | 0 | 0 | 1 | 1 |
| 0 | 0 | 0 | 0 | 1 | 1 |
| 0 | 0 | 0 | 0 | 1 | 1 |
| 0 | 0 | 0 | 0 | 1 | 1 |
| 0 | 0 | 0 | 0 | 1 | 1 |
| 0 | 0 | 0 | 0 | 1 | 1 |
| 0 | 0 | 0 | 0 | 1 | 1 |
| 0 | 0 | 0 | 0 | 1 | 1 |
| 0 | 0 | 0 | 0 | 1 | 1 |
| 0 | 0 | 0 | 0 | 1 | 1 |
| 0 | 0 | 0 | 0 | 1 | 1 |
| 0 | 0 | 0 | 0 | 1 | 1 |
| 0 | 0 | 0 | 0 | 1 | 1 |
| 0 | 0 | 0 | 0 | 1 | 1 |
| 0 | 0 | 0 | 0 | 1 | 1 |
| 0 | 0 | 0 | 0 | 1 | 1 |
| 0 | 0 | 0 | 0 | 1 | 1 |
| 0 | 0 | 0 | 0 | 1 | 1 |
| 0 | 0 | 0 | 0 | 1 | 1 |
| 0 | 0 | 0 | 0 | 1 | 1 |
| 0 | 0 | 0 | 0 | 1 | 1 |
| 1 | 1 | 1 | 1 | 1 | 8 |
| 1 | 1 | 1 | 1 | 1 | 8 |
| 1 | 1 | 1 | 1 | 1 | 8 |
| 1 | 1 | 1 | 1 | 1 | 8 |
| 1 | 1 | 1 | 1 | 1 | 8 |
| 1 | 1 | 1 | 1 | 1 | 8 |
| 1 | 1 | 1 | 1 | 1 | 8 |
| 1 | 1 | 1 | 1 | 1 | 8 |
| 1 | 1 | 1 | 1 | 1 | 8 |
| 1 | 1 | 1 | 1 | 1 | 8 |
| 0 | 1 | 0 | 0 | 0 | 2 |
| 0 | 1 | 0 | 0 | 0 | 2 |
| 0 | 1 | 0 | 0 | 0 | 2 |
| 0 | 1 | 0 | 0 | 0 | 2 |
| 0 | 1 | 0 | 0 | 0 | 2 |
| 0 | 1 | 0 | 0 | 0 | 2 |
| 0 | 1 | 0 | 0 | 0 | 2 |
| 0 | 1 | 0 | 0 | 0 | 2 |
| 0 | 1 | 0 | 0 | 0 | 2 |

| | | | | | |
|---|---|---|---|---|---|
| 0 | 1 | 0 | 0 | 0 | 2 |
| 1 | 1 | 1 | 1 | 1 | 8 |
| 1 | 1 | 1 | 1 | 1 | 8 |
| 1 | 1 | 1 | 1 | 1 | 8 |
| 1 | 1 | 1 | 1 | 1 | 8 |
| 1 | 1 | 1 | 1 | 1 | 8 |
| 1 | 1 | 1 | 1 | 1 | 8 |
| 1 | 1 | 1 | 1 | 1 | 8 |
| 1 | 1 | 1 | 1 | 1 | 8 |
| 1 | 1 | 1 | 1 | 1 | 8 |
| 1 | 1 | 1 | 1 | 1 | 8 |
| 1 | 1 | 1 | 1 | 1 | 8 |
| 1 | 1 | 1 | 1 | 1 | 8 |
| 1 | 1 | 1 | 1 | 1 | 8 |
| 1 | 1 | 1 | 1 | 1 | 8 |
| 1 | 1 | 1 | 1 | 1 | 8 |
| 1 | 1 | 1 | 1 | 1 | 8 |
| 1 | 1 | 1 | 1 | 1 | 8 |
| 1 | 1 | 1 | 1 | 1 | 8 |
| 1 | 1 | 1 | 1 | 1 | 8 |
| 1 | 1 | 1 | 1 | 1 | 8 |
| 1 | 1 | 1 | 1 | 1 | 8 |
| 1 | 1 | 1 | 1 | 1 | 8 |
| 1 | 1 | 1 | 1 | 1 | 8 |
| 1 | 1 | 1 | 1 | 1 | 8 |
| 1 | 1 | 1 | 1 | 1 | 8 |
| 1 | 1 | 1 | 1 | 1 | 8 |
| 1 | 1 | 1 | 1 | 1 | 8 |
| 1 | 1 | 1 | 1 | 1 | 8 |
| 1 | 1 | 1 | 1 | 1 | 8 |
| 1 | 1 | 1 | 1 | 1 | 8 |
| 1 | 1 | 1 | 1 | 1 | 8 |
| 1 | 1 | 1 | 1 | 1 | 8 |
| 1 | 1 | 1 | 1 | 1 | 8 |
| 1 | 1 | 1 | 1 | 1 | 8 |
| 1 | 1 | 1 | 1 | 1 | 8 |
| 1 | 1 | 1 | 1 | 1 | 8 |
| 1 | 1 | 1 | 1 | 1 | 8 |
| 1 | 1 | 1 | 1 | 1 | 8 |
| 1 | 1 | 1 | 1 | 1 | 8 |
| 1 | 1 | 1 | 1 | 1 | 8 |
| 1 | 1 | 1 | 1 | 1 | 8 |
| 1 | 1 | 1 | 1 | 1 | 8 |
| 0 | 1 | 0 | 0 | 0 | 2 |

| | | | | | |
|---|---|---|---|---|---|
| 0 | 1 | 0 | 0 | 0 | 2 |
| 0 | 1 | 0 | 0 | 0 | 2 |
| 0 | 1 | 0 | 0 | 0 | 2 |
| 0 | 1 | 0 | 0 | 0 | 2 |
| 0 | 1 | 0 | 0 | 0 | 2 |
| 0 | 1 | 0 | 0 | 0 | 2 |
| 0 | 1 | 0 | 0 | 0 | 2 |
| 0 | 1 | 0 | 0 | 0 | 2 |
| 0 | 1 | 0 | 0 | 0 | 2 |
| 0 | 1 | 0 | 0 | 0 | 2 |
| 0 | 1 | 0 | 0 | 0 | 2 |
| 0 | 1 | 0 | 0 | 0 | 2 |
| 0 | 1 | 0 | 0 | 0 | 2 |
| 0 | 1 | 0 | 0 | 0 | 2 |
| 0 | 1 | 0 | 0 | 0 | 2 |
| 0 | 1 | 0 | 0 | 0 | 2 |
| 0 | 1 | 0 | 0 | 0 | 2 |
| 0 | 1 | 0 | 0 | 0 | 2 |
| 0 | 1 | 0 | 0 | 0 | 2 |
| 0 | 1 | 0 | 0 | 0 | 2 |
| 0 | 1 | 0 | 0 | 0 | 2 |
| 0 | 1 | 0 | 0 | 0 | 2 |
| 0 | 1 | 0 | 0 | 0 | 2 |
| 0 | 1 | 0 | 0 | 0 | 2 |
| 0 | 1 | 0 | 0 | 0 | 2 |
| 0 | 1 | 0 | 0 | 0 | 2 |
| 0 | 1 | 0 | 0 | 0 | 2 |
| 0 | 1 | 0 | 0 | 0 | 2 |
| 0 | 1 | 0 | 0 | 0 | 2 |
| 0 | 1 | 0 | 0 | 0 | 2 |
| 0 | 1 | 0 | 0 | 0 | 2 |
| 1 | 1 | 1 | 1 | 1 | 8 |
| 1 | 1 | 1 | 1 | 1 | 8 |
| 1 | 1 | 1 | 1 | 1 | 8 |
| 1 | 1 | 1 | 1 | 1 | 8 |
| 1 | 1 | 1 | 1 | 1 | 8 |
| 1 | 1 | 1 | 1 | 1 | 8 |
| 1 | 1 | 1 | 1 | 1 | 8 |
| 1 | 1 | 1 | 1 | 1 | 8 |
| 1 | 1 | 1 | 1 | 1 | 8 |
| 1 | 1 | 1 | 1 | 1 | 8 |
| 1 | 1 | 1 | 1 | 1 | 8 |
| 1 | 1 | 1 | 1 | 1 | 8 |
| 1 | 1 | 1 | 1 | 1 | 8 |

| | | | | | |
|---|---|---|---|---|---|
| 1 | 1 | 1 | 1 | 1 | 8 |
| 1 | 1 | 1 | 1 | 1 | 8 |
| 1 | 1 | 1 | 1 | 1 | 8 |
| 1 | 1 | 1 | 1 | 1 | 8 |
| 1 | 1 | 1 | 1 | 1 | 8 |
| 1 | 1 | 1 | 1 | 1 | 8 |
| 1 | 1 | 1 | 1 | 1 | 8 |
| 1 | 1 | 1 | 1 | 1 | 8 |
| 1 | 1 | 1 | 1 | 1 | 8 |
| 1 | 1 | 1 | 1 | 1 | 8 |
| 1 | 1 | 1 | 1 | 1 | 8 |
| 1 | 1 | 1 | 1 | 1 | 8 |
| 1 | 1 | 1 | 1 | 1 | 8 |
| 1 | 1 | 1 | 1 | 1 | 8 |
| 1 | 1 | 1 | 1 | 1 | 8 |
| 1 | 1 | 1 | 1 | 1 | 8 |
| 1 | 1 | 1 | 1 | 1 | 8 |
| 1 | 1 | 1 | 1 | 1 | 8 |
| 1 | 1 | 1 | 1 | 1 | 8 |
| 1 | 1 | 1 | 1 | 1 | 8 |
| 1 | 1 | 1 | 1 | 1 | 8 |
| 1 | 1 | 1 | 1 | 1 | 8 |
| 1 | 1 | 1 | 1 | 1 | 8 |
| 1 | 1 | 1 | 1 | 1 | 8 |
| 1 | 1 | 1 | 1 | 1 | 8 |
| 1 | 1 | 1 | 1 | 1 | 8 |
| 1 | 1 | 1 | 1 | 1 | 8 |
| 0 | 1 | 0 | 0 | 0 | 2 |
| 0 | 1 | 0 | 0 | 0 | 2 |
| 0 | 1 | 0 | 0 | 0 | 2 |
| 0 | 1 | 0 | 0 | 0 | 2 |
| 0 | 1 | 0 | 0 | 0 | 2 |
| 0 | 1 | 0 | 0 | 0 | 2 |
| 0 | 1 | 0 | 0 | 0 | 2 |
| 0 | 1 | 0 | 0 | 0 | 2 |
| 0 | 1 | 0 | 0 | 0 | 2 |
| 0 | 1 | 0 | 0 | 0 | 2 |
| 0 | 1 | 0 | 0 | 0 | 2 |
| 0 | 1 | 0 | 0 | 0 | 2 |
| 0 | 1 | 0 | 0 | 0 | 2 |
| 0 | 1 | 0 | 0 | 0 | 2 |
| 0 | 1 | 0 | 0 | 0 | 2 |
| 0 | 1 | 0 | 0 | 0 | 2 |
| 0 | 1 | 0 | 0 | 0 | 2 |

| | | | | | |
|---|---|---|---|---|---|
| 0 | 1 | 0 | 0 | 0 | 2 |
| 0 | 1 | 0 | 0 | 0 | 2 |
| 0 | 1 | 0 | 0 | 0 | 2 |
| 0 | 1 | 0 | 0 | 0 | 2 |
| 0 | 1 | 0 | 0 | 0 | 2 |
| 0 | 1 | 0 | 0 | 0 | 2 |
| 0 | 1 | 0 | 0 | 0 | 2 |
| 0 | 1 | 0 | 0 | 0 | 2 |
| 0 | 1 | 0 | 0 | 0 | 2 |
| 0 | 1 | 0 | 0 | 0 | 2 |
| 0 | 1 | 0 | 0 | 0 | 2 |
| 0 | 1 | 0 | 0 | 0 | 2 |
| 0 | 1 | 0 | 0 | 0 | 2 |
| 0 | 1 | 0 | 0 | 0 | 2 |
| 0 | 1 | 0 | 0 | 0 | 2 |
| 0 | 1 | 0 | 0 | 0 | 2 |
| 0 | 1 | 0 | 0 | 0 | 2 |
| 0 | 1 | 0 | 0 | 0 | 2 |
| 0 | 1 | 0 | 0 | 0 | 2 |
| 0 | 1 | 0 | 0 | 0 | 2 |
| 0 | 1 | 0 | 0 | 0 | 2 |
| 0 | 1 | 0 | 0 | 0 | 2 |
| 0 | 1 | 0 | 0 | 0 | 2 |
| 1 | 1 | 1 | 1 | 1 | 8 |
| 1 | 1 | 1 | 1 | 1 | 8 |
| 1 | 1 | 1 | 1 | 1 | 8 |
| 1 | 1 | 1 | 1 | 1 | 8 |
| 1 | 1 | 1 | 1 | 1 | 8 |
| 1 | 1 | 1 | 1 | 1 | 8 |
| 1 | 1 | 1 | 1 | 1 | 8 |
| 1 | 1 | 1 | 1 | 1 | 8 |
| 1 | 1 | 1 | 1 | 1 | 8 |
| 1 | 1 | 1 | 1 | 1 | 8 |
| 0 | 0 | 0 | 0 | 0 | 1 |
| 0 | 0 | 0 | 0 | 0 | 1 |
| 0 | 0 | 0 | 0 | 0 | 1 |
| 0 | 0 | 0 | 0 | 0 | 1 |
| 0 | 0 | 0 | 0 | 0 | 1 |
| 0 | 0 | 0 | 0 | 0 | 1 |
| 0 | 0 | 0 | 0 | 0 | 1 |
| 0 | 0 | 0 | 0 | 0 | 1 |
| 0 | 0 | 0 | 0 | 0 | 1 |
| 0 | 0 | 0 | 0 | 0 | 1 |
| 0 | 0 | 0 | 0 | 0 | 1 |

| | | | | | |
|---|---|---|---|---|---|
| 0 | 0 | 0 | 0 | 0 | 1 |
| 0 | 0 | 0 | 0 | 0 | 1 |
| 0 | 0 | 0 | 0 | 0 | 1 |
| 0 | 0 | 0 | 0 | 0 | 1 |
| 0 | 0 | 0 | 0 | 0 | 1 |
| 0 | 0 | 0 | 0 | 0 | 1 |
| 0 | 0 | 0 | 0 | 0 | 1 |
| 0 | 0 | 0 | 0 | 0 | 1 |
| 0 | 0 | 0 | 0 | 0 | 1 |
| 0 | 0 | 0 | 0 | 0 | 1 |
| 0 | 0 | 0 | 0 | 0 | 1 |
| 0 | 0 | 0 | 0 | 0 | 1 |
| 0 | 0 | 0 | 0 | 0 | 1 |
| 0 | 0 | 0 | 0 | 0 | 1 |
| 0 | 0 | 0 | 0 | 0 | 1 |
| 0 | 0 | 0 | 0 | 0 | 1 |
| 0 | 0 | 0 | 0 | 0 | 1 |
| 0 | 0 | 0 | 0 | 0 | 1 |
| 0 | 0 | 0 | 0 | 0 | 1 |
| 0 | 0 | 0 | 0 | 0 | 1 |
| 0 | 0 | 0 | 0 | 0 | 1 |
| 0 | 0 | 0 | 0 | 0 | 1 |
| 0 | 0 | 0 | 0 | 0 | 1 |
| 0 | 0 | 0 | 0 | 0 | 1 |
| 0 | 0 | 0 | 0 | 0 | 1 |
| 0 | 0 | 0 | 0 | 0 | 1 |
| 0 | 0 | 0 | 0 | 0 | 1 |
| 0 | 0 | 0 | 0 | 0 | 1 |
| 0 | 0 | 0 | 0 | 0 | 1 |
| 0 | 0 | 0 | 0 | 0 | 1 |
| 0 | 0 | 0 | 0 | 0 | 1 |
| 0 | 0 | 0 | 0 | 0 | 1 |
| 0 | 0 | 0 | 0 | 0 | 1 |
| 0 | 0 | 0 | 0 | 0 | 1 |
| 0 | 0 | 0 | 0 | 0 | 1 |
| 0 | 0 | 0 | 0 | 0 | 1 |
| 0 | 0 | 0 | 0 | 0 | 1 |
| 0 | 0 | 0 | 0 | 0 | 1 |
| 0 | 0 | 0 | 0 | 0 | 1 |
| 0 | 0 | 0 | 0 | 0 | 1 |
| 0 | 0 | 0 | 0 | 0 | 1 |
| 0 | 0 | 0 | 0 | 0 | 1 |
| 0 | 0 | 0 | 0 | 0 | 1 |
| 0 | 0 | 0 | 0 | 0 | 1 |

| | | | | | |
|---|---|---|---|---|---|
| 0 | 0 | 0 | 0 | 0 | 1 |
| 1 | 1 | 1 | 1 | 1 | 8 |
| 1 | 1 | 1 | 1 | 1 | 8 |
| 1 | 1 | 1 | 1 | 1 | 8 |
| 1 | 1 | 1 | 1 | 1 | 8 |
| 1 | 1 | 1 | 1 | 1 | 8 |
| 1 | 1 | 1 | 1 | 1 | 8 |
| 1 | 1 | 1 | 1 | 1 | 8 |
| 1 | 1 | 1 | 1 | 1 | 8 |
| 0 | 0 | 0 | 0 | 0 | 1 |
| 0 | 0 | 0 | 0 | 0 | 1 |
| 0 | 0 | 0 | 0 | 0 | 1 |
| 0 | 0 | 0 | 0 | 0 | 1 |
| 0 | 0 | 0 | 0 | 0 | 1 |
| 0 | 0 | 0 | 0 | 0 | 1 |
| 0 | 0 | 0 | 0 | 0 | 1 |
| 0 | 0 | 0 | 0 | 0 | 1 |
| 0 | 0 | 0 | 0 | 0 | 1 |
| 0 | 0 | 0 | 0 | 0 | 1 |
| 0 | 0 | 0 | 0 | 0 | 1 |
| 0 | 0 | 0 | 0 | 0 | 1 |
| 0 | 0 | 0 | 0 | 0 | 1 |
| 0 | 0 | 0 | 0 | 0 | 1 |
| 0 | 0 | 0 | 0 | 0 | 1 |
| 0 | 0 | 0 | 0 | 0 | 1 |
| 0 | 0 | 0 | 0 | 0 | 1 |
| 0 | 0 | 0 | 0 | 0 | 1 |
| 0 | 0 | 0 | 0 | 0 | 1 |
| 0 | 0 | 0 | 0 | 0 | 1 |
| 0 | 0 | 0 | 0 | 0 | 1 |
| 0 | 0 | 0 | 0 | 0 | 1 |
| 0 | 0 | 0 | 0 | 0 | 1 |
| 0 | 0 | 0 | 0 | 0 | 1 |
| 0 | 0 | 0 | 0 | 0 | 1 |
| 0 | 0 | 0 | 0 | 0 | 1 |
| 0 | 0 | 0 | 0 | 0 | 1 |
| 0 | 0 | 0 | 0 | 0 | 1 |
| 0 | 0 | 0 | 0 | 0 | 1 |
| 0 | 0 | 0 | 0 | 0 | 1 |
| 0 | 0 | 0 | 0 | 0 | 1 |
| 0 | 0 | 0 | 0 | 0 | 1 |
| 0 | 0 | 0 | 0 | 0 | 1 |
| 0 | 0 | 0 | 0 | 0 | 1 |
| 0 | 0 | 0 | 0 | 0 | 1 |

| | | | | | |
|---|---|---|---|---|---|
| 0 | 0 | 0 | 0 | 0 | 1 |
| 0 | 0 | 0 | 0 | 0 | 1 |
| 0 | 0 | 0 | 0 | 0 | 1 |
| 0 | 0 | 0 | 0 | 0 | 1 |
| 0 | 0 | 0 | 0 | 0 | 1 |
| 0 | 0 | 0 | 0 | 0 | 1 |
| 0 | 0 | 0 | 0 | 0 | 1 |
| 0 | 0 | 0 | 0 | 0 | 1 |
| 0 | 0 | 0 | 0 | 0 | 1 |
| 0 | 0 | 0 | 0 | 0 | 1 |
| 0 | 0 | 0 | 0 | 0 | 1 |
| 0 | 0 | 0 | 0 | 0 | 1 |
| 0 | 0 | 0 | 0 | 0 | 1 |
| 0 | 0 | 0 | 0 | 0 | 1 |
| 0 | 0 | 0 | 0 | 0 | 1 |
| 0 | 0 | 0 | 0 | 0 | 1 |
| 0 | 0 | 0 | 0 | 0 | 1 |
| 0 | 0 | 0 | 0 | 0 | 1 |
| 0 | 0 | 0 | 0 | 0 | 1 |
| 0 | 0 | 0 | 0 | 0 | 1 |
| 0 | 0 | 0 | 0 | 0 | 1 |
| 1 | 1 | 1 | 1 | 1 | 8 |
| 1 | 1 | 1 | 1 | 1 | 8 |
| 1 | 1 | 1 | 1 | 1 | 8 |
| 1 | 1 | 1 | 1 | 1 | 8 |
| 1 | 1 | 1 | 1 | 1 | 8 |
| 1 | 1 | 1 | 1 | 1 | 8 |
| 1 | 1 | 1 | 1 | 1 | 8 |
| 1 | 1 | 1 | 1 | 1 | 8 |
| 1 | 1 | 1 | 1 | 1 | 8 |
| 1 | 1 | 1 | 1 | 1 | 8 |
| 0 | 0 | 0 | 0 | 0 | 1 |
| 0 | 0 | 0 | 0 | 0 | 1 |
| 0 | 0 | 0 | 0 | 0 | 1 |
| 0 | 0 | 0 | 0 | 0 | 1 |
| 0 | 0 | 0 | 0 | 0 | 1 |
| 0 | 0 | 0 | 0 | 0 | 1 |
| 0 | 0 | 0 | 0 | 0 | 1 |
| 0 | 0 | 0 | 0 | 0 | 1 |
| 0 | 0 | 0 | 0 | 0 | 1 |
| 0 | 0 | 0 | 0 | 0 | 1 |
| 0 | 0 | 0 | 0 | 0 | 1 |
| 0 | 0 | 0 | 0 | 0 | 1 |
| 0 | 0 | 0 | 0 | 0 | 1 |

| | | | | | |
|---|---|---|---|---|---|
| 0 | 0 | 0 | 0 | 0 | 1 |
| 0 | 0 | 0 | 0 | 0 | 1 |
| 0 | 0 | 0 | 0 | 0 | 1 |
| 0 | 0 | 0 | 0 | 0 | 1 |
| 0 | 0 | 0 | 0 | 0 | 1 |
| 0 | 0 | 0 | 0 | 0 | 1 |
| 0 | 0 | 0 | 0 | 0 | 1 |
| 0 | 0 | 0 | 0 | 0 | 1 |
| 0 | 0 | 0 | 0 | 0 | 1 |
| 0 | 0 | 0 | 0 | 0 | 1 |
| 0 | 0 | 0 | 0 | 0 | 1 |
| 0 | 0 | 0 | 0 | 0 | 1 |
| 0 | 0 | 0 | 0 | 0 | 1 |
| 0 | 0 | 0 | 0 | 0 | 1 |
| 0 | 0 | 0 | 0 | 0 | 1 |
| 0 | 0 | 0 | 0 | 0 | 1 |
| 0 | 0 | 0 | 0 | 0 | 1 |
| 0 | 0 | 0 | 0 | 0 | 1 |
| 0 | 0 | 0 | 0 | 0 | 1 |
| 0 | 0 | 0 | 0 | 0 | 1 |
| 0 | 0 | 0 | 0 | 0 | 1 |
| 0 | 0 | 0 | 0 | 0 | 1 |
| 0 | 0 | 0 | 0 | 0 | 1 |
| 0 | 0 | 0 | 0 | 0 | 1 |
| 0 | 0 | 0 | 0 | 0 | 1 |
| 0 | 0 | 0 | 0 | 0 | 1 |
| 0 | 0 | 0 | 0 | 0 | 1 |
| 0 | 0 | 0 | 0 | 0 | 1 |
| 0 | 0 | 0 | 0 | 0 | 1 |
| 0 | 0 | 0 | 0 | 0 | 1 |
| 0 | 0 | 0 | 0 | 0 | 1 |
| 0 | 0 | 0 | 0 | 0 | 1 |
| 0 | 0 | 0 | 0 | 0 | 1 |
| 0 | 0 | 0 | 0 | 0 | 1 |
| 0 | 0 | 0 | 0 | 0 | 1 |
| 0 | 0 | 0 | 0 | 0 | 1 |
| 0 | 0 | 0 | 0 | 0 | 1 |
| 0 | 0 | 0 | 0 | 0 | 1 |
| 0 | 0 | 0 | 0 | 0 | 1 |
| 0 | 0 | 0 | 0 | 0 | 1 |
| 0 | 0 | 0 | 0 | 0 | 1 |
| 0 | 0 | 0 | 0 | 0 | 1 |
| 0 | 0 | 0 | 0 | 0 | 1 |
| 1 | 0 | 0 | 0 | 0 | 2 |

| | | | | | |
|---|---|---|---|---|---|
| 1 | 0 | 0 | 0 | 0 | 2 |
| 1 | 0 | 0 | 0 | 0 | 2 |
| 1 | 0 | 0 | 0 | 0 | 2 |
| 1 | 0 | 0 | 0 | 0 | 2 |
| 1 | 0 | 0 | 0 | 0 | 2 |
| 1 | 0 | 0 | 0 | 0 | 2 |
| 1 | 0 | 0 | 0 | 0 | 2 |
| 0 | 1 | 1 | 1 | 0 | 4 |
| 0 | 1 | 1 | 1 | 0 | 4 |
| 0 | 1 | 1 | 1 | 0 | 4 |
| 0 | 1 | 1 | 1 | 0 | 4 |
| 0 | 1 | 1 | 1 | 0 | 4 |
| 0 | 1 | 1 | 1 | 0 | 4 |
| 0 | 1 | 1 | 1 | 0 | 4 |
| 0 | 1 | 1 | 1 | 0 | 4 |
| 0 | 1 | 1 | 1 | 0 | 4 |
| 0 | 1 | 1 | 1 | 0 | 4 |
| 0 | 1 | 1 | 1 | 0 | 4 |
| 0 | 1 | 1 | 1 | 0 | 4 |
| 0 | 1 | 1 | 1 | 0 | 4 |
| 0 | 1 | 1 | 1 | 0 | 4 |
| 0 | 1 | 1 | 1 | 0 | 4 |
| 0 | 1 | 1 | 1 | 0 | 4 |
| 0 | 1 | 1 | 1 | 0 | 4 |
| 0 | 1 | 1 | 1 | 0 | 4 |
| 0 | 1 | 1 | 1 | 0 | 4 |
| 0 | 1 | 1 | 1 | 0 | 4 |
| 0 | 1 | 1 | 1 | 0 | 4 |
| 0 | 1 | 1 | 1 | 0 | 4 |
| 0 | 1 | 1 | 1 | 0 | 4 |
| 0 | 1 | 1 | 1 | 0 | 4 |
| 0 | 1 | 1 | 1 | 0 | 4 |
| 0 | 1 | 1 | 1 | 0 | 4 |
| 0 | 1 | 1 | 1 | 0 | 4 |
| 0 | 1 | 1 | 1 | 0 | 4 |
| 0 | 1 | 1 | 1 | 0 | 4 |
| 0 | 1 | 1 | 1 | 0 | 4 |
| 0 | 1 | 1 | 1 | 0 | 4 |
| 0 | 1 | 1 | 1 | 0 | 4 |
| 0 | 1 | 1 | 1 | 0 | 4 |
| 0 | 1 | 1 | 1 | 0 | 4 |
| 0 | 1 | 1 | 1 | 0 | 4 |
| 0 | 1 | 1 | 1 | 0 | 4 |
| 0 | 1 | 1 | 1 | 0 | 4 |

| | | | | | |
|---|---|---|---|---|---|
| 0 | 1 | 1 | 1 | 0 | 4 |
| 0 | 1 | 1 | 1 | 0 | 4 |
| 0 | 1 | 1 | 1 | 0 | 4 |
| 0 | 1 | 1 | 1 | 0 | 4 |
| 0 | 1 | 1 | 1 | 0 | 4 |
| 0 | 1 | 1 | 1 | 0 | 4 |
| 0 | 1 | 1 | 1 | 0 | 4 |
| 0 | 1 | 1 | 1 | 0 | 4 |
| 0 | 1 | 1 | 1 | 0 | 4 |
| 0 | 1 | 1 | 1 | 0 | 4 |
| 0 | 1 | 1 | 1 | 0 | 4 |
| 0 | 1 | 1 | 1 | 0 | 4 |
| 0 | 1 | 1 | 1 | 0 | 4 |
| 0 | 1 | 1 | 1 | 0 | 4 |
| 0 | 1 | 1 | 1 | 0 | 4 |
| 0 | 1 | 1 | 1 | 0 | 4 |
| 0 | 1 | 1 | 1 | 0 | 4 |
| 0 | 1 | 1 | 1 | 0 | 4 |
| 0 | 1 | 1 | 1 | 0 | 4 |
| 1 | 0 | 0 | 0 | 0 | 2 |
| 1 | 0 | 0 | 0 | 0 | 2 |
| 1 | 0 | 0 | 0 | 0 | 2 |
| 1 | 0 | 0 | 0 | 0 | 2 |
| 1 | 0 | 0 | 0 | 0 | 2 |
| 1 | 0 | 0 | 0 | 0 | 2 |
| 1 | 0 | 0 | 0 | 0 | 2 |
| 1 | 0 | 0 | 0 | 0 | 2 |
| 1 | 0 | 0 | 0 | 0 | 2 |
| 1 | 0 | 0 | 0 | 0 | 2 |
| 1 | 0 | 0 | 0 | 0 | 2 |
| 1 | 0 | 0 | 0 | 0 | 2 |
| 1 | 0 | 0 | 0 | 0 | 2 |
| 1 | 0 | 0 | 0 | 0 | 2 |
| 1 | 0 | 0 | 0 | 0 | 2 |
| 1 | 0 | 0 | 0 | 0 | 2 |
| 1 | 0 | 0 | 0 | 0 | 2 |
| 1 | 0 | 0 | 0 | 0 | 2 |
| 1 | 0 | 0 | 0 | 0 | 2 |
| 1 | 0 | 0 | 0 | 0 | 2 |
| 1 | 0 | 0 | 0 | 0 | 2 |
| 1 | 0 | 0 | 0 | 0 | 2 |
| 1 | 0 | 0 | 0 | 0 | 2 |
| 1 | 0 | 0 | 0 | 0 | 2 |
| 1 | 0 | 0 | 0 | 0 | 2 |

| | | | | | |
|---|---|---|---|---|---|
| 1 | 0 | 0 | 0 | 0 | 2 |
| 1 | 0 | 0 | 0 | 0 | 2 |
| 1 | 0 | 0 | 0 | 0 | 2 |
| 1 | 0 | 0 | 0 | 0 | 2 |
| 1 | 0 | 0 | 0 | 0 | 2 |
| 1 | 0 | 0 | 0 | 0 | 2 |
| 1 | 0 | 0 | 0 | 0 | 2 |
| 1 | 0 | 0 | 0 | 0 | 2 |
| 1 | 0 | 0 | 0 | 0 | 2 |
| 1 | 0 | 0 | 0 | 0 | 2 |
| 1 | 0 | 0 | 0 | 0 | 2 |
| 1 | 0 | 0 | 0 | 0 | 2 |
| 1 | 0 | 0 | 0 | 0 | 2 |
| 1 | 0 | 0 | 0 | 0 | 2 |
| 1 | 0 | 0 | 0 | 0 | 2 |
| 1 | 0 | 0 | 0 | 0 | 2 |
| 1 | 0 | 0 | 0 | 0 | 2 |
| 1 | 0 | 0 | 0 | 0 | 2 |
| 1 | 0 | 0 | 0 | 0 | 2 |
| 1 | 0 | 0 | 0 | 0 | 2 |
| 1 | 0 | 0 | 0 | 0 | 2 |
| 1 | 0 | 0 | 0 | 0 | 2 |
| 1 | 0 | 0 | 0 | 0 | 2 |
| 1 | 1 | 1 | 1 | 1 | 8 |
| 1 | 1 | 1 | 1 | 1 | 8 |
| 1 | 1 | 1 | 1 | 1 | 8 |
| 1 | 1 | 1 | 1 | 1 | 8 |
| 1 | 1 | 1 | 1 | 1 | 8 |
| 1 | 1 | 1 | 1 | 1 | 8 |
| 1 | 1 | 1 | 1 | 1 | 8 |
| 1 | 1 | 1 | 1 | 1 | 8 |
| 1 | 1 | 1 | 1 | 1 | 8 |
| 0 | 0 | 0 | 0 | 1 | 1 |
| 0 | 0 | 0 | 0 | 1 | 1 |
| 0 | 0 | 0 | 0 | 1 | 1 |
| 0 | 0 | 0 | 0 | 1 | 1 |
| 0 | 0 | 0 | 0 | 1 | 1 |
| 0 | 0 | 0 | 0 | 1 | 1 |
| 0 | 0 | 0 | 0 | 1 | 1 |
| 0 | 0 | 0 | 0 | 1 | 1 |
| 0 | 0 | 0 | 0 | 1 | 1 |
| 0 | 0 | 0 | 0 | 1 | 1 |
| 0 | 0 | 0 | 0 | 1 | 1 |
| 0 | 0 | 0 | 0 | 1 | 1 |

| | | | | | |
|---|---|---|---|---|---|
| 0 | 0 | 0 | 0 | 1 | 1 |
| 0 | 0 | 0 | 0 | 1 | 1 |
| 0 | 0 | 0 | 0 | 1 | 1 |
| 0 | 0 | 0 | 0 | 1 | 1 |
| 0 | 0 | 0 | 0 | 1 | 1 |
| 0 | 0 | 0 | 0 | 1 | 1 |
| 0 | 0 | 0 | 0 | 1 | 1 |
| 0 | 0 | 0 | 0 | 1 | 1 |
| 0 | 0 | 0 | 0 | 1 | 1 |
| 0 | 0 | 0 | 0 | 1 | 1 |
| 0 | 0 | 0 | 0 | 1 | 1 |
| 0 | 0 | 0 | 0 | 1 | 1 |
| 0 | 0 | 0 | 0 | 1 | 1 |
| 0 | 0 | 0 | 0 | 1 | 1 |
| 0 | 0 | 0 | 0 | 1 | 1 |
| 0 | 0 | 0 | 0 | 1 | 1 |
| 0 | 0 | 0 | 0 | 1 | 1 |
| 0 | 0 | 0 | 0 | 1 | 1 |
| 0 | 0 | 0 | 0 | 1 | 1 |
| 0 | 0 | 0 | 0 | 1 | 1 |
| 0 | 0 | 0 | 0 | 1 | 1 |
| 0 | 0 | 0 | 0 | 1 | 1 |
| 0 | 0 | 0 | 0 | 1 | 1 |
| 0 | 0 | 0 | 0 | 1 | 1 |
| 0 | 0 | 0 | 0 | 1 | 1 |
| 0 | 0 | 0 | 0 | 1 | 1 |
| 0 | 0 | 0 | 0 | 1 | 1 |
| 0 | 0 | 0 | 0 | 1 | 1 |
| 0 | 0 | 0 | 0 | 1 | 1 |
| 0 | 0 | 0 | 0 | 1 | 1 |
| 0 | 0 | 0 | 0 | 1 | 1 |
| 0 | 0 | 0 | 0 | 1 | 1 |
| 0 | 0 | 0 | 0 | 1 | 1 |
| 0 | 0 | 0 | 0 | 1 | 1 |
| 0 | 0 | 0 | 0 | 1 | 1 |
| 0 | 0 | 0 | 0 | 1 | 1 |
| 0 | 0 | 0 | 0 | 1 | 1 |
| 0 | 0 | 0 | 0 | 1 | 1 |
| 0 | 0 | 0 | 0 | 1 | 1 |
| 0 | 0 | 0 | 0 | 1 | 1 |
| 0 | 0 | 0 | 0 | 1 | 1 |
| 0 | 0 | 0 | 0 | 1 | 1 |
| 0 | 0 | 0 | 0 | 1 | 1 |
| 0 | 0 | 0 | 0 | 1 | 1 |

| | | | | | |
|---|---|---|---|---|---|
| 0 | 0 | 0 | 0 | 1 | 1 |
| 0 | 0 | 0 | 0 | 1 | 1 |
| 0 | 0 | 0 | 0 | 1 | 1 |
| 0 | 0 | 0 | 0 | 1 | 1 |
| 0 | 0 | 0 | 0 | 1 | 1 |
| 0 | 0 | 0 | 0 | 1 | 1 |
| 0 | 0 | 0 | 0 | 1 | 1 |
| 0 | 0 | 0 | 0 | 1 | 1 |
| 0 | 0 | 0 | 0 | 1 | 1 |
| 0 | 0 | 0 | 0 | 1 | 1 |
| 0 | 0 | 0 | 0 | 1 | 1 |
| 0 | 0 | 0 | 0 | 1 | 1 |
| 0 | 0 | 0 | 0 | 1 | 1 |
| 0 | 0 | 0 | 0 | 1 | 1 |
| 0 | 0 | 0 | 0 | 1 | 1 |
| 0 | 0 | 0 | 0 | 1 | 1 |
| 0 | 0 | 0 | 0 | 1 | 1 |
| 0 | 0 | 0 | 0 | 1 | 1 |
| 0 | 0 | 0 | 0 | 1 | 1 |
| 0 | 0 | 0 | 0 | 1 | 1 |
| 0 | 0 | 0 | 0 | 1 | 1 |
| 0 | 0 | 0 | 0 | 1 | 1 |
| 0 | 0 | 0 | 0 | 1 | 1 |
| 0 | 0 | 0 | 0 | 1 | 1 |
| 0 | 0 | 0 | 0 | 1 | 1 |
| 0 | 0 | 0 | 0 | 1 | 1 |
| 0 | 0 | 0 | 0 | 1 | 1 |
| 0 | 0 | 0 | 0 | 1 | 1 |
| 0 | 0 | 0 | 0 | 1 | 1 |
| 0 | 0 | 0 | 0 | 1 | 1 |
| 0 | 0 | 0 | 0 | 1 | 1 |
| 0 | 0 | 0 | 0 | 1 | 1 |
| 0 | 0 | 0 | 0 | 1 | 1 |
| 0 | 0 | 0 | 0 | 1 | 1 |
| 0 | 0 | 0 | 0 | 1 | 1 |
| 0 | 0 | 0 | 0 | 1 | 1 |
| 0 | 0 | 0 | 0 | 1 | 1 |
| 0 | 0 | 0 | 0 | 1 | 1 |
| 0 | 0 | 0 | 0 | 1 | 1 |
| 0 | 0 | 0 | 0 | 1 | 1 |
| 0 | 0 | 0 | 0 | 1 | 1 |
| 0 | 0 | 0 | 0 | 1 | 1 |
| 0 | 0 | 0 | 0 | 1 | 1 |
| 0 | 0 | 0 | 0 | 1 | 1 |

| | | | | | |
|---|---|---|---|---|---|
| 0 | 0 | 0 | 0 | 1 | 1 |
| 0 | 0 | 0 | 0 | 1 | 1 |
| 0 | 0 | 0 | 0 | 1 | 1 |
| 0 | 0 | 0 | 0 | 1 | 1 |
| 0 | 0 | 0 | 0 | 1 | 1 |
| 0 | 0 | 0 | 0 | 1 | 1 |
| 0 | 0 | 0 | 0 | 1 | 1 |
| 0 | 0 | 0 | 0 | 1 | 1 |
| 0 | 0 | 0 | 0 | 1 | 1 |
| 0 | 0 | 0 | 0 | 1 | 1 |
| 0 | 0 | 0 | 0 | 1 | 1 |
| 0 | 0 | 0 | 0 | 1 | 1 |
| 0 | 0 | 0 | 0 | 1 | 1 |
| 0 | 0 | 0 | 0 | 1 | 1 |
| 0 | 0 | 0 | 0 | 1 | 1 |
| 0 | 0 | 0 | 0 | 1 | 1 |
| 0 | 0 | 0 | 0 | 1 | 1 |
| 0 | 0 | 0 | 0 | 1 | 1 |
| 0 | 0 | 0 | 0 | 1 | 1 |
| 0 | 0 | 0 | 0 | 1 | 1 |
| 0 | 0 | 0 | 0 | 1 | 1 |
| 0 | 0 | 0 | 0 | 1 | 1 |
| 0 | 0 | 0 | 0 | 1 | 1 |
| 0 | 0 | 0 | 0 | 1 | 1 |
| 0 | 0 | 0 | 0 | 1 | 1 |
| 0 | 0 | 0 | 0 | 1 | 1 |
| 0 | 0 | 0 | 0 | 1 | 1 |
| 0 | 0 | 0 | 0 | 1 | 1 |
| 0 | 0 | 0 | 0 | 1 | 1 |
| 0 | 0 | 0 | 0 | 1 | 1 |
| 0 | 0 | 0 | 0 | 1 | 1 |
| 0 | 0 | 0 | 0 | 1 | 1 |
| 0 | 0 | 0 | 0 | 1 | 1 |
| 0 | 0 | 0 | 0 | 1 | 1 |
| 0 | 0 | 0 | 0 | 1 | 1 |
| 0 | 0 | 0 | 0 | 1 | 1 |
| 0 | 0 | 0 | 0 | 1 | 1 |
| 0 | 0 | 0 | 0 | 1 | 1 |
| 0 | 0 | 0 | 0 | 1 | 1 |
| 0 | 0 | 0 | 0 | 1 | 1 |
| 0 | 0 | 0 | 0 | 1 | 1 |
| 0 | 0 | 0 | 0 | 1 | 1 |
| 0 | 0 | 0 | 0 | 1 | 1 |
| 0 | 0 | 0 | 0 | 1 | 1 |

| | | | | | |
|---|---|---|---|---|---|
| 0 | 0 | 0 | 0 | 1 | 1 |
| 0 | 0 | 0 | 0 | 1 | 1 |
| 0 | 0 | 0 | 0 | 1 | 1 |
| 0 | 0 | 0 | 0 | 1 | 1 |
| 0 | 0 | 0 | 0 | 1 | 1 |
| 0 | 0 | 0 | 0 | 1 | 1 |
| 0 | 0 | 0 | 0 | 1 | 1 |
| 0 | 0 | 0 | 0 | 1 | 1 |
| 0 | 0 | 0 | 0 | 1 | 1 |
| 0 | 0 | 0 | 0 | 1 | 1 |
| 0 | 0 | 0 | 0 | 1 | 1 |
| 0 | 0 | 0 | 0 | 1 | 1 |
| 0 | 0 | 0 | 0 | 1 | 1 |
| 0 | 0 | 0 | 0 | 1 | 1 |
| 0 | 0 | 0 | 0 | 1 | 1 |
| 0 | 0 | 0 | 0 | 1 | 1 |
| 0 | 0 | 0 | 0 | 1 | 1 |
| 0 | 0 | 0 | 0 | 1 | 1 |
| 0 | 0 | 0 | 0 | 1 | 1 |
| 0 | 0 | 0 | 0 | 1 | 1 |
| 0 | 0 | 0 | 0 | 1 | 1 |
| 0 | 0 | 0 | 0 | 1 | 1 |
| 0 | 0 | 0 | 0 | 1 | 1 |
| 0 | 0 | 0 | 0 | 1 | 1 |
| 0 | 0 | 0 | 0 | 1 | 1 |
| 0 | 0 | 0 | 0 | 1 | 1 |
| 0 | 0 | 0 | 0 | 1 | 1 |
| 0 | 0 | 0 | 0 | 1 | 1 |
| 0 | 0 | 0 | 0 | 1 | 1 |
| 0 | 0 | 0 | 0 | 1 | 1 |
| 0 | 0 | 0 | 0 | 1 | 1 |
| 0 | 0 | 0 | 0 | 1 | 1 |
| 0 | 0 | 0 | 0 | 1 | 1 |
| 0 | 0 | 0 | 0 | 1 | 1 |
| 0 | 0 | 0 | 0 | 1 | 1 |
| 0 | 0 | 0 | 0 | 1 | 1 |
| 0 | 0 | 0 | 0 | 1 | 1 |
| 0 | 0 | 0 | 0 | 1 | 1 |
| 0 | 0 | 0 | 0 | 1 | 1 |
| 0 | 0 | 0 | 0 | 1 | 1 |
| 0 | 0 | 0 | 0 | 1 | 1 |
| 0 | 0 | 0 | 0 | 1 | 1 |
| 0 | 0 | 0 | 0 | 1 | 1 |
| 0 | 0 | 0 | 0 | 1 | 1 |

| | | | | | |
|---|---|---|---|---|---|
| 0 | 0 | 0 | 0 | 1 | 1 |
| 0 | 0 | 0 | 0 | 1 | 1 |
| 0 | 0 | 0 | 0 | 1 | 1 |
| 0 | 0 | 0 | 0 | 1 | 1 |
| 0 | 0 | 0 | 0 | 1 | 1 |
| 0 | 0 | 0 | 0 | 1 | 1 |
| 0 | 0 | 0 | 0 | 1 | 1 |
| 0 | 0 | 0 | 0 | 1 | 1 |
| 0 | 0 | 0 | 0 | 1 | 1 |
| 0 | 0 | 0 | 0 | 1 | 1 |
| 0 | 0 | 0 | 0 | 1 | 1 |
| 0 | 0 | 0 | 0 | 1 | 1 |
| 0 | 0 | 0 | 0 | 1 | 1 |
| 0 | 0 | 0 | 0 | 1 | 1 |
| 0 | 0 | 0 | 0 | 1 | 1 |
| 0 | 0 | 0 | 0 | 1 | 1 |
| 0 | 0 | 0 | 0 | 1 | 1 |
| 0 | 0 | 0 | 0 | 1 | 1 |
| 0 | 0 | 0 | 0 | 1 | 1 |
| 0 | 0 | 0 | 0 | 1 | 1 |
| 0 | 0 | 0 | 0 | 1 | 1 |
| 0 | 0 | 0 | 0 | 1 | 1 |
| 0 | 0 | 0 | 0 | 1 | 1 |
| 0 | 0 | 0 | 0 | 1 | 1 |
| 0 | 0 | 0 | 0 | 1 | 1 |
| 0 | 0 | 0 | 0 | 1 | 1 |
| 0 | 0 | 0 | 0 | 1 | 1 |
| 0 | 0 | 0 | 0 | 1 | 1 |
| 0 | 0 | 0 | 0 | 1 | 1 |
| 0 | 0 | 0 | 0 | 1 | 1 |
| 0 | 0 | 0 | 0 | 1 | 1 |
| 0 | 0 | 0 | 0 | 1 | 1 |
| 0 | 0 | 0 | 0 | 1 | 1 |
| 0 | 0 | 0 | 0 | 1 | 1 |
| 0 | 0 | 0 | 0 | 1 | 1 |
| 0 | 0 | 0 | 0 | 1 | 1 |
| 0 | 0 | 0 | 0 | 1 | 1 |
| 0 | 0 | 0 | 0 | 1 | 1 |
| 0 | 0 | 0 | 0 | 1 | 1 |
| 0 | 0 | 0 | 0 | 1 | 1 |
| 0 | 0 | 0 | 0 | 1 | 1 |
| 0 | 0 | 0 | 0 | 1 | 1 |
| 0 | 0 | 0 | 0 | 1 | 1 |
| 0 | 0 | 0 | 0 | 1 | 1 |

| | | | | | |
|---|---|---|---|---|---|
| 0 | 0 | 0 | 0 | 1 | 1 |
| 0 | 0 | 0 | 0 | 1 | 1 |
| 0 | 0 | 0 | 0 | 1 | 1 |
| 0 | 0 | 0 | 0 | 1 | 1 |
| 0 | 0 | 0 | 0 | 1 | 1 |
| 0 | 0 | 0 | 0 | 1 | 1 |
| 0 | 0 | 0 | 0 | 1 | 1 |
| 0 | 0 | 0 | 0 | 1 | 1 |
| 0 | 0 | 0 | 0 | 1 | 1 |
| 0 | 0 | 0 | 0 | 1 | 1 |
| 0 | 0 | 0 | 0 | 1 | 1 |
| 0 | 0 | 0 | 0 | 1 | 1 |
| 0 | 0 | 0 | 0 | 1 | 1 |
| 0 | 0 | 0 | 0 | 1 | 1 |
| 0 | 0 | 0 | 0 | 1 | 1 |
| 0 | 0 | 0 | 0 | 1 | 1 |
| 0 | 0 | 0 | 0 | 1 | 1 |
| 0 | 0 | 0 | 0 | 1 | 1 |
| 0 | 0 | 0 | 0 | 1 | 1 |
| 0 | 0 | 0 | 0 | 1 | 1 |
| 0 | 0 | 0 | 0 | 1 | 1 |
| 0 | 0 | 0 | 0 | 1 | 1 |
| 0 | 0 | 0 | 0 | 1 | 1 |
| 0 | 0 | 0 | 0 | 1 | 1 |
| 0 | 0 | 0 | 0 | 1 | 1 |
| 0 | 0 | 0 | 0 | 1 | 1 |
| 0 | 0 | 0 | 0 | 1 | 1 |
| 0 | 0 | 0 | 0 | 1 | 1 |
| 0 | 0 | 0 | 0 | 1 | 1 |
| 0 | 0 | 0 | 0 | 1 | 1 |
| 0 | 0 | 0 | 0 | 1 | 1 |
| 0 | 0 | 0 | 0 | 1 | 1 |
| 0 | 0 | 0 | 0 | 1 | 1 |
| 0 | 0 | 0 | 0 | 1 | 1 |
| 0 | 0 | 0 | 0 | 1 | 1 |
| 0 | 0 | 0 | 0 | 1 | 1 |
| 0 | 0 | 0 | 0 | 1 | 1 |
| 0 | 0 | 0 | 0 | 1 | 1 |
| 0 | 0 | 0 | 0 | 1 | 1 |
| 0 | 0 | 0 | 0 | 1 | 1 |
| 0 | 0 | 0 | 0 | 1 | 1 |
| 0 | 0 | 0 | 0 | 1 | 1 |
| 0 | 0 | 0 | 0 | 1 | 1 |
| 0 | 0 | 0 | 0 | 1 | 1 |

| | | | | | |
|---|---|---|---|---|---|
| 0 | 0 | 0 | 0 | 1 | 1 |
| 0 | 0 | 0 | 0 | 1 | 1 |
| 0 | 0 | 0 | 0 | 1 | 1 |
| 0 | 0 | 0 | 0 | 1 | 1 |
| 0 | 0 | 0 | 0 | 1 | 1 |
| 0 | 0 | 0 | 0 | 1 | 1 |
| 0 | 0 | 0 | 0 | 1 | 1 |
| 0 | 0 | 0 | 0 | 1 | 1 |
| 0 | 0 | 0 | 0 | 1 | 1 |
| 0 | 0 | 0 | 0 | 1 | 1 |
| 0 | 0 | 0 | 0 | 1 | 1 |
| 0 | 0 | 0 | 0 | 1 | 1 |
| 0 | 0 | 0 | 0 | 1 | 1 |
| 0 | 0 | 0 | 0 | 1 | 1 |
| 0 | 0 | 0 | 0 | 1 | 1 |
| 0 | 0 | 0 | 0 | 1 | 1 |
| 0 | 0 | 0 | 0 | 1 | 1 |
| 0 | 0 | 0 | 0 | 1 | 1 |
| 0 | 0 | 0 | 0 | 1 | 1 |
| 0 | 0 | 0 | 0 | 1 | 1 |
| 0 | 0 | 0 | 0 | 1 | 1 |
| 0 | 0 | 0 | 0 | 1 | 1 |
| 0 | 0 | 0 | 0 | 1 | 1 |
| 0 | 0 | 0 | 0 | 1 | 1 |
| 0 | 0 | 0 | 0 | 0 | 1 |
| 0 | 0 | 0 | 0 | 0 | 1 |
| 0 | 0 | 0 | 0 | 0 | 1 |
| 0 | 0 | 0 | 0 | 0 | 1 |
| 0 | 0 | 0 | 0 | 0 | 1 |
| 0 | 0 | 0 | 0 | 0 | 1 |
| 0 | 0 | 0 | 0 | 0 | 1 |
| 0 | 0 | 0 | 0 | 0 | 1 |
| 0 | 0 | 0 | 0 | 0 | 1 |
| 0 | 0 | 0 | 0 | 0 | 1 |
| 0 | 0 | 0 | 0 | 0 | 1 |
| 0 | 0 | 0 | 0 | 0 | 1 |
| 0 | 0 | 0 | 0 | 0 | 1 |
| 0 | 0 | 0 | 0 | 0 | 1 |
| 0 | 0 | 0 | 0 | 0 | 1 |
| 0 | 0 | 0 | 0 | 0 | 1 |
| 0 | 0 | 0 | 0 | 0 | 1 |
| 0 | 0 | 0 | 0 | 0 | 1 |
| 0 | 0 | 0 | 0 | 0 | 1 |
| 0 | 0 | 0 | 0 | 0 | 1 |

| | | | | | |
|---|---|---|---|---|---|
| 0 | 0 | 0 | 0 | 0 | 1 |
| 0 | 0 | 0 | 0 | 0 | 1 |
| 0 | 0 | 0 | 0 | 0 | 1 |
| 0 | 0 | 0 | 0 | 0 | 1 |
| 0 | 0 | 0 | 0 | 0 | 1 |
| 0 | 0 | 0 | 0 | 0 | 1 |
| 0 | 0 | 0 | 0 | 0 | 1 |
| 0 | 0 | 0 | 0 | 0 | 1 |
| 0 | 0 | 0 | 0 | 0 | 1 |
| 0 | 0 | 0 | 0 | 0 | 1 |
| 0 | 0 | 0 | 0 | 0 | 1 |
| 0 | 0 | 0 | 0 | 0 | 1 |
| 0 | 0 | 0 | 0 | 0 | 1 |
| 0 | 0 | 0 | 0 | 0 | 1 |
| 0 | 0 | 0 | 0 | 0 | 1 |
| 0 | 0 | 0 | 0 | 0 | 1 |
| 0 | 0 | 0 | 0 | 0 | 1 |
| 0 | 0 | 0 | 0 | 0 | 1 |
| 0 | 0 | 0 | 0 | 0 | 1 |
| 0 | 0 | 0 | 0 | 0 | 1 |
| 0 | 0 | 0 | 0 | 0 | 1 |
| 0 | 0 | 0 | 0 | 0 | 1 |
| 0 | 0 | 0 | 0 | 0 | 1 |
| 0 | 0 | 0 | 0 | 0 | 1 |
| 0 | 0 | 0 | 0 | 0 | 1 |
| 0 | 0 | 0 | 0 | 0 | 1 |
| 0 | 0 | 0 | 0 | 0 | 1 |
| 0 | 0 | 0 | 0 | 0 | 1 |
| 0 | 0 | 0 | 0 | 0 | 1 |
| 0 | 0 | 0 | 0 | 0 | 1 |
| 0 | 0 | 0 | 0 | 0 | 1 |
| 0 | 0 | 0 | 0 | 0 | 1 |
| 0 | 0 | 0 | 0 | 0 | 1 |
| 0 | 0 | 0 | 0 | 0 | 1 |
| 0 | 0 | 0 | 0 | 0 | 1 |
| 0 | 0 | 0 | 0 | 0 | 1 |
| 0 | 0 | 0 | 0 | 0 | 1 |
| 0 | 0 | 0 | 0 | 0 | 1 |
| 0 | 0 | 0 | 0 | 0 | 1 |
| 0 | 0 | 0 | 0 | 0 | 1 |
| 0 | 0 | 0 | 0 | 0 | 1 |
| 0 | 0 | 0 | 0 | 0 | 1 |
| 0 | 0 | 0 | 0 | 0 | 1 |
| 0 | 0 | 0 | 0 | 0 | 1 |

| | | | | | |
|---|---|---|---|---|---|
| 0 | 0 | 0 | 0 | 0 | 1 |
| 0 | 0 | 0 | 0 | 0 | 1 |
| 0 | 0 | 0 | 0 | 0 | 1 |
| 0 | 0 | 0 | 0 | 0 | 1 |
| 0 | 0 | 0 | 0 | 0 | 1 |
| 0 | 0 | 0 | 0 | 0 | 1 |
| 0 | 0 | 0 | 0 | 0 | 1 |
| 0 | 0 | 0 | 0 | 0 | 1 |
| 0 | 0 | 0 | 0 | 0 | 1 |
| 0 | 0 | 0 | 0 | 0 | 1 |
| 0 | 0 | 0 | 0 | 0 | 1 |
| 0 | 0 | 0 | 0 | 0 | 1 |
| 0 | 0 | 0 | 0 | 0 | 1 |
| 0 | 0 | 0 | 0 | 0 | 1 |
| 0 | 0 | 0 | 0 | 0 | 1 |
| 0 | 0 | 0 | 0 | 0 | 1 |
| 0 | 0 | 0 | 0 | 0 | 1 |
| 0 | 0 | 0 | 0 | 0 | 1 |
| 0 | 0 | 0 | 0 | 0 | 1 |
| 0 | 0 | 0 | 0 | 0 | 1 |
| 0 | 0 | 0 | 0 | 0 | 1 |
| 0 | 0 | 0 | 0 | 0 | 1 |
| 0 | 0 | 0 | 0 | 0 | 1 |
| 0 | 0 | 0 | 0 | 0 | 1 |
| 0 | 0 | 0 | 0 | 0 | 1 |
| 0 | 0 | 0 | 0 | 0 | 1 |
| 0 | 0 | 0 | 0 | 0 | 1 |
| 0 | 0 | 0 | 0 | 0 | 1 |
| 0 | 0 | 0 | 0 | 0 | 1 |
| 0 | 0 | 0 | 0 | 0 | 1 |
| 0 | 0 | 0 | 0 | 0 | 1 |
| 0 | 0 | 0 | 0 | 0 | 1 |
| 0 | 0 | 0 | 0 | 0 | 1 |
| 0 | 0 | 0 | 0 | 0 | 1 |
| 0 | 0 | 0 | 0 | 0 | 1 |
| 0 | 0 | 0 | 0 | 0 | 1 |
| 0 | 0 | 0 | 0 | 0 | 1 |
| 0 | 0 | 0 | 0 | 0 | 1 |
| 0 | 0 | 0 | 0 | 0 | 1 |
| 0 | 0 | 0 | 0 | 0 | 1 |
| 0 | 0 | 0 | 0 | 0 | 1 |
| 0 | 0 | 0 | 0 | 0 | 1 |
| 0 | 0 | 0 | 0 | 0 | 1 |
| 0 | 0 | 0 | 0 | 0 | 1 |

| | | | | | |
|---|---|---|---|---|---|
| 0 | 0 | 0 | 0 | 0 | 1 |
| 0 | 0 | 0 | 0 | 0 | 1 |
| 0 | 0 | 0 | 0 | 0 | 1 |
| 0 | 0 | 0 | 0 | 0 | 1 |
| 0 | 0 | 0 | 0 | 0 | 1 |
| 0 | 0 | 0 | 0 | 0 | 1 |
| 0 | 0 | 0 | 0 | 0 | 1 |
| 0 | 0 | 0 | 0 | 0 | 1 |
| 0 | 0 | 0 | 0 | 0 | 1 |
| 0 | 0 | 0 | 0 | 0 | 1 |
| 0 | 0 | 0 | 0 | 0 | 1 |
| 0 | 0 | 0 | 0 | 0 | 1 |
| 0 | 0 | 0 | 0 | 0 | 1 |
| 0 | 0 | 0 | 0 | 0 | 1 |
| 0 | 0 | 0 | 0 | 0 | 1 |
| 0 | 0 | 0 | 0 | 0 | 1 |
| 0 | 0 | 0 | 0 | 0 | 1 |
| 0 | 0 | 0 | 0 | 0 | 1 |
| 0 | 0 | 0 | 0 | 0 | 1 |
| 0 | 0 | 0 | 0 | 0 | 1 |
| 0 | 0 | 0 | 0 | 0 | 1 |
| 0 | 0 | 0 | 0 | 0 | 1 |
| 0 | 0 | 0 | 0 | 0 | 1 |
| 0 | 0 | 0 | 0 | 0 | 1 |
| 0 | 0 | 0 | 0 | 0 | 1 |
| 0 | 0 | 0 | 0 | 0 | 1 |
| 0 | 0 | 0 | 0 | 0 | 1 |
| 0 | 0 | 0 | 0 | 0 | 1 |
| 0 | 0 | 0 | 0 | 0 | 1 |
| 0 | 0 | 0 | 0 | 0 | 1 |
| 0 | 0 | 0 | 0 | 0 | 1 |
| 0 | 0 | 0 | 0 | 0 | 1 |
| 0 | 0 | 0 | 0 | 0 | 1 |
| 0 | 0 | 0 | 0 | 0 | 1 |
| 0 | 0 | 0 | 0 | 0 | 1 |
| 0 | 0 | 0 | 0 | 0 | 1 |
| 0 | 0 | 0 | 0 | 0 | 1 |
| 0 | 0 | 0 | 0 | 0 | 1 |
| 0 | 0 | 0 | 0 | 0 | 1 |
| 0 | 0 | 0 | 0 | 0 | 1 |
| 0 | 0 | 0 | 0 | 0 | 1 |
| 0 | 0 | 0 | 0 | 0 | 1 |
| 0 | 0 | 0 | 0 | 0 | 1 |
| 0 | 0 | 0 | 0 | 0 | 1 |

| | | | | | |
|---|---|---|---|---|---|
| 0 | 0 | 0 | 0 | 0 | 1 |
| 0 | 0 | 0 | 0 | 0 | 1 |
| 0 | 0 | 0 | 0 | 0 | 1 |
| 0 | 0 | 0 | 0 | 0 | 1 |
| 0 | 0 | 0 | 0 | 0 | 1 |
| 0 | 0 | 0 | 0 | 0 | 1 |
| 0 | 0 | 0 | 0 | 0 | 1 |
| 0 | 0 | 0 | 0 | 0 | 1 |
| 0 | 0 | 0 | 0 | 0 | 1 |
| 0 | 0 | 0 | 0 | 0 | 1 |
| 0 | 0 | 0 | 0 | 0 | 1 |
| 0 | 0 | 0 | 0 | 0 | 1 |
| 0 | 0 | 0 | 0 | 0 | 1 |
| 0 | 0 | 0 | 0 | 0 | 1 |
| 0 | 0 | 0 | 0 | 0 | 1 |
| 0 | 0 | 0 | 0 | 0 | 1 |
| 0 | 0 | 0 | 0 | 0 | 1 |
| 0 | 0 | 0 | 0 | 0 | 1 |
| 0 | 0 | 0 | 0 | 0 | 1 |
| 0 | 0 | 0 | 0 | 0 | 1 |
| 0 | 0 | 0 | 0 | 0 | 1 |
| 0 | 0 | 0 | 0 | 0 | 1 |
| 0 | 0 | 0 | 0 | 0 | 1 |
| 0 | 0 | 0 | 0 | 0 | 1 |
| 0 | 0 | 0 | 0 | 0 | 1 |
| 0 | 0 | 0 | 0 | 0 | 1 |
| 0 | 0 | 0 | 0 | 0 | 1 |
| 0 | 0 | 0 | 0 | 0 | 1 |
| 0 | 0 | 0 | 0 | 0 | 1 |
| 0 | 0 | 0 | 0 | 0 | 1 |
| 0 | 0 | 0 | 0 | 0 | 1 |
| 0 | 0 | 0 | 0 | 0 | 1 |
| 0 | 0 | 0 | 0 | 0 | 1 |
| 0 | 0 | 0 | 0 | 0 | 1 |
| 0 | 0 | 0 | 0 | 0 | 1 |
| 0 | 0 | 0 | 0 | 0 | 1 |
| 0 | 0 | 0 | 0 | 0 | 1 |
| 0 | 0 | 0 | 0 | 0 | 1 |
| 0 | 0 | 0 | 0 | 0 | 1 |
| 0 | 0 | 0 | 0 | 0 | 1 |
| 0 | 0 | 0 | 0 | 0 | 1 |
| 0 | 0 | 0 | 0 | 0 | 1 |
| 0 | 0 | 0 | 0 | 0 | 1 |
| 0 | 0 | 0 | 0 | 0 | 1 |

| | | | | | |
|---|---|---|---|---|---|
| 0 | 0 | 0 | 0 | 0 | 1 |
| 0 | 0 | 0 | 0 | 0 | 1 |
| 0 | 0 | 0 | 0 | 0 | 1 |
| 0 | 0 | 0 | 0 | 0 | 1 |
| 0 | 0 | 0 | 0 | 0 | 1 |
| 0 | 0 | 0 | 0 | 0 | 1 |
| 0 | 0 | 0 | 0 | 0 | 1 |
| 0 | 0 | 0 | 0 | 0 | 1 |
| 0 | 0 | 0 | 0 | 0 | 1 |
| 0 | 0 | 0 | 0 | 0 | 1 |
| 0 | 0 | 0 | 0 | 0 | 1 |
| 0 | 0 | 0 | 0 | 0 | 1 |
| 0 | 0 | 0 | 0 | 0 | 1 |
| 0 | 0 | 0 | 0 | 0 | 1 |
| 0 | 0 | 0 | 0 | 0 | 1 |
| 0 | 0 | 0 | 0 | 0 | 1 |
| 0 | 0 | 0 | 0 | 0 | 1 |
| 0 | 0 | 0 | 0 | 0 | 1 |
| 0 | 0 | 0 | 0 | 0 | 1 |
| 0 | 0 | 0 | 0 | 0 | 1 |
| 0 | 0 | 0 | 0 | 0 | 1 |
| 0 | 0 | 0 | 0 | 0 | 1 |
| 0 | 0 | 0 | 0 | 0 | 1 |
| 0 | 0 | 0 | 0 | 0 | 1 |
| 0 | 0 | 0 | 0 | 0 | 1 |
| 0 | 0 | 0 | 0 | 0 | 1 |
| 0 | 0 | 0 | 0 | 0 | 1 |
| 0 | 0 | 0 | 0 | 0 | 1 |
| 0 | 0 | 0 | 0 | 0 | 1 |
| 0 | 0 | 0 | 0 | 0 | 1 |
| 0 | 0 | 0 | 0 | 0 | 1 |
| 0 | 0 | 0 | 0 | 0 | 1 |
| 0 | 0 | 0 | 0 | 0 | 1 |
| 0 | 0 | 0 | 0 | 0 | 1 |
| 0 | 0 | 0 | 0 | 0 | 1 |
| 0 | 0 | 0 | 0 | 0 | 1 |
| 0 | 0 | 0 | 0 | 0 | 1 |
| 0 | 0 | 0 | 0 | 0 | 1 |
| 0 | 0 | 0 | 0 | 0 | 1 |
| 0 | 0 | 0 | 0 | 0 | 1 |
| 0 | 0 | 0 | 0 | 0 | 1 |
| 0 | 0 | 0 | 0 | 0 | 1 |
| 0 | 0 | 0 | 0 | 0 | 1 |
| 0 | 0 | 0 | 0 | 0 | 1 |

| | | | | | |
|---|---|---|---|---|---|
| 0 | 0 | 0 | 0 | 0 | 1 |
| 0 | 0 | 0 | 0 | 0 | 1 |
| 0 | 0 | 0 | 0 | 0 | 1 |
| 0 | 0 | 0 | 0 | 0 | 1 |
| 0 | 0 | 0 | 0 | 0 | 1 |
| 0 | 0 | 0 | 0 | 0 | 1 |
| 0 | 0 | 0 | 0 | 0 | 1 |
| 0 | 0 | 0 | 0 | 0 | 1 |
| 0 | 0 | 0 | 0 | 0 | 1 |
| 0 | 0 | 0 | 0 | 0 | 1 |
| 0 | 0 | 0 | 0 | 0 | 1 |
| 0 | 0 | 0 | 0 | 0 | 1 |
| 0 | 0 | 0 | 0 | 0 | 1 |
| 0 | 0 | 0 | 0 | 0 | 1 |
| 0 | 0 | 0 | 0 | 0 | 1 |
| 0 | 0 | 0 | 0 | 0 | 1 |
| 0 | 0 | 0 | 0 | 0 | 1 |
| 0 | 0 | 0 | 0 | 0 | 1 |
| 0 | 0 | 0 | 0 | 0 | 1 |
| 0 | 0 | 0 | 0 | 0 | 1 |
| 0 | 0 | 0 | 0 | 0 | 1 |
| 0 | 0 | 0 | 0 | 0 | 1 |
| 0 | 0 | 0 | 0 | 0 | 1 |
| 0 | 0 | 0 | 0 | 0 | 1 |
| 0 | 0 | 0 | 0 | 0 | 1 |
| 0 | 0 | 0 | 0 | 0 | 1 |
| 0 | 0 | 0 | 0 | 0 | 1 |
| 0 | 0 | 0 | 0 | 0 | 1 |
| 0 | 0 | 0 | 0 | 0 | 1 |
| 0 | 0 | 0 | 0 | 0 | 1 |
| 0 | 0 | 0 | 0 | 0 | 1 |
| 0 | 0 | 0 | 0 | 0 | 1 |
| 0 | 0 | 0 | 0 | 0 | 1 |
| 0 | 0 | 0 | 0 | 0 | 1 |
| 0 | 0 | 0 | 0 | 0 | 1 |
| 0 | 0 | 0 | 0 | 0 | 1 |
| 0 | 0 | 0 | 0 | 0 | 1 |
| 0 | 0 | 0 | 0 | 0 | 1 |
| 0 | 0 | 0 | 0 | 0 | 1 |
| 0 | 0 | 0 | 0 | 0 | 1 |
| 0 | 0 | 0 | 0 | 0 | 1 |
| 0 | 0 | 0 | 0 | 0 | 1 |
| 0 | 0 | 0 | 0 | 0 | 1 |
| 0 | 0 | 0 | 0 | 0 | 1 |

| | | | | | |
|---|---|---|---|---|---|
| 0 | 0 | 0 | 0 | 0 | 1 |
| 0 | 0 | 0 | 0 | 0 | 1 |
| 0 | 0 | 0 | 0 | 0 | 1 |
| 0 | 0 | 0 | 0 | 0 | 1 |
| 0 | 0 | 0 | 0 | 0 | 1 |
| 0 | 0 | 0 | 0 | 0 | 1 |
| 0 | 0 | 0 | 0 | 0 | 1 |
| 0 | 0 | 0 | 0 | 0 | 1 |
| 0 | 0 | 0 | 0 | 0 | 1 |
| 0 | 0 | 0 | 0 | 0 | 1 |
| 0 | 0 | 0 | 0 | 0 | 1 |
| 0 | 0 | 0 | 0 | 0 | 1 |
| 0 | 0 | 0 | 0 | 0 | 1 |
| 0 | 0 | 0 | 0 | 0 | 1 |
| 0 | 0 | 0 | 0 | 0 | 1 |
| 0 | 0 | 0 | 0 | 0 | 1 |
| 0 | 1 | 1 | 1 | 0 | 4 |
| 0 | 1 | 1 | 1 | 0 | 4 |
| 0 | 1 | 1 | 1 | 0 | 4 |
| 0 | 1 | 1 | 1 | 0 | 4 |
| 0 | 1 | 1 | 1 | 0 | 4 |
| 0 | 1 | 1 | 1 | 0 | 4 |
| 0 | 1 | 1 | 1 | 0 | 4 |
| 0 | 1 | 1 | 1 | 0 | 4 |
| 0 | 1 | 1 | 1 | 0 | 4 |
| 0 | 1 | 1 | 1 | 0 | 4 |
| 0 | 1 | 1 | 1 | 0 | 4 |
| 0 | 1 | 1 | 1 | 0 | 4 |
| 0 | 1 | 1 | 1 | 0 | 4 |
| 0 | 1 | 1 | 1 | 0 | 4 |
| 0 | 1 | 1 | 1 | 0 | 4 |
| 0 | 1 | 1 | 1 | 0 | 4 |
| 0 | 1 | 1 | 1 | 0 | 4 |
| 0 | 1 | 1 | 1 | 0 | 4 |
| 0 | 1 | 1 | 1 | 0 | 4 |
| 0 | 1 | 1 | 1 | 0 | 4 |
| 0 | 1 | 1 | 1 | 0 | 4 |
| 0 | 1 | 1 | 1 | 0 | 4 |
| 0 | 1 | 1 | 1 | 0 | 4 |
| 0 | 1 | 1 | 1 | 0 | 4 |
| 0 | 1 | 1 | 1 | 0 | 4 |
| 0 | 1 | 1 | 1 | 0 | 4 |
| 0 | 1 | 1 | 1 | 0 | 4 |
| 0 | 1 | 1 | 1 | 0 | 4 |

| | | | | | |
|---|---|---|---|---|---|
| 0 | 1 | 1 | 1 | 0 | 4 |
| 0 | 1 | 1 | 1 | 0 | 4 |
| 0 | 1 | 1 | 1 | 0 | 4 |
| 0 | 1 | 1 | 1 | 0 | 4 |
| 0 | 1 | 1 | 1 | 0 | 4 |
| 0 | 1 | 1 | 1 | 0 | 4 |
| 0 | 1 | 1 | 1 | 0 | 4 |
| 0 | 1 | 1 | 1 | 0 | 4 |
| 0 | 1 | 1 | 1 | 0 | 4 |
| 0 | 1 | 1 | 1 | 0 | 4 |
| 0 | 1 | 1 | 1 | 0 | 4 |
| 0 | 1 | 1 | 1 | 0 | 4 |
| 0 | 1 | 1 | 1 | 0 | 4 |
| 0 | 1 | 1 | 1 | 0 | 4 |
| 0 | 1 | 1 | 1 | 0 | 4 |
| 0 | 1 | 1 | 1 | 0 | 4 |
| 0 | 1 | 1 | 1 | 0 | 4 |
| 0 | 1 | 1 | 1 | 0 | 4 |
| 0 | 1 | 1 | 1 | 0 | 4 |
| 0 | 1 | 1 | 1 | 0 | 4 |
| 0 | 1 | 1 | 1 | 0 | 4 |
| 0 | 1 | 1 | 1 | 0 | 4 |
| 0 | 1 | 1 | 1 | 0 | 4 |
| 0 | 1 | 1 | 1 | 0 | 4 |
| 0 | 1 | 1 | 1 | 0 | 4 |
| 0 | 1 | 1 | 1 | 0 | 4 |
| 0 | 1 | 1 | 1 | 0 | 4 |
| 0 | 1 | 1 | 1 | 0 | 4 |
| 0 | 1 | 1 | 1 | 0 | 4 |
| 0 | 1 | 1 | 1 | 0 | 4 |
| 0 | 1 | 1 | 1 | 0 | 4 |
| 0 | 1 | 1 | 1 | 0 | 4 |
| 0 | 1 | 1 | 1 | 0 | 4 |
| 0 | 1 | 1 | 1 | 0 | 4 |
| 0 | 1 | 1 | 1 | 0 | 4 |
| 0 | 1 | 1 | 1 | 0 | 4 |
| 0 | 1 | 1 | 1 | 0 | 4 |
| 0 | 1 | 1 | 1 | 0 | 4 |
| 0 | 1 | 1 | 1 | 0 | 4 |
| 0 | 1 | 1 | 1 | 0 | 4 |
| 0 | 1 | 1 | 1 | 0 | 4 |
| 0 | 1 | 1 | 1 | 0 | 4 |
| 0 | 1 | 1 | 1 | 0 | 4 |
| 0 | 1 | 1 | 1 | 0 | 4 |

| | | | | | |
|---|---|---|---|---|---|
| 0 | 1 | 1 | 1 | 0 | 4 |
| 0 | 1 | 1 | 1 | 0 | 4 |
| 0 | 1 | 1 | 1 | 0 | 4 |
| 0 | 1 | 1 | 1 | 0 | 4 |
| 0 | 1 | 1 | 1 | 0 | 4 |
| 0 | 1 | 1 | 1 | 0 | 4 |
| 0 | 1 | 1 | 1 | 0 | 4 |
| 0 | 1 | 1 | 1 | 0 | 4 |
| 0 | 1 | 1 | 1 | 0 | 4 |
| 0 | 1 | 1 | 1 | 0 | 4 |
| 0 | 1 | 1 | 1 | 0 | 4 |
| 0 | 1 | 1 | 1 | 0 | 4 |
| 0 | 1 | 1 | 1 | 0 | 4 |
| 0 | 1 | 1 | 1 | 0 | 4 |
| 0 | 1 | 1 | 1 | 0 | 4 |
| 0 | 1 | 1 | 1 | 0 | 4 |
| 0 | 1 | 1 | 1 | 0 | 4 |
| 0 | 1 | 1 | 1 | 0 | 4 |
| 0 | 1 | 1 | 1 | 0 | 4 |
| 0 | 1 | 1 | 1 | 0 | 4 |
| 0 | 1 | 1 | 1 | 0 | 4 |
| 0 | 1 | 1 | 1 | 0 | 4 |
| 0 | 1 | 1 | 1 | 0 | 4 |
| 0 | 1 | 1 | 1 | 0 | 4 |
| 0 | 1 | 1 | 1 | 0 | 4 |
| 0 | 1 | 1 | 1 | 0 | 4 |
| 0 | 1 | 1 | 1 | 0 | 4 |
| 0 | 1 | 1 | 1 | 0 | 4 |
| 0 | 1 | 1 | 1 | 0 | 4 |
| 0 | 1 | 1 | 1 | 0 | 4 |
| 0 | 1 | 1 | 1 | 0 | 4 |
| 0 | 1 | 1 | 1 | 0 | 4 |
| 0 | 1 | 1 | 1 | 0 | 4 |
| 0 | 1 | 1 | 1 | 0 | 4 |
| 0 | 1 | 1 | 1 | 0 | 4 |
| 0 | 1 | 1 | 1 | 0 | 4 |
| 0 | 1 | 1 | 1 | 0 | 4 |
| 0 | 1 | 1 | 1 | 0 | 4 |
| 0 | 1 | 1 | 1 | 0 | 4 |
| 0 | 1 | 1 | 1 | 0 | 4 |
| 0 | 1 | 1 | 1 | 0 | 4 |
| 0 | 1 | 1 | 1 | 0 | 4 |
| 0 | 1 | 1 | 1 | 0 | 4 |
| 0 | 1 | 1 | 1 | 0 | 4 |

| | | | | | |
|---|---|---|---|---|---|
| 0 | 1 | 1 | 1 | 0 | 4 |
| 0 | 1 | 1 | 1 | 0 | 4 |
| 0 | 1 | 1 | 1 | 0 | 4 |
| 0 | 1 | 1 | 1 | 0 | 4 |
| 1 | 0 | 0 | 0 | 0 | 2 |
| 1 | 0 | 0 | 0 | 0 | 2 |
| 1 | 0 | 0 | 0 | 0 | 2 |
| 1 | 0 | 0 | 0 | 0 | 2 |
| 1 | 0 | 0 | 0 | 0 | 2 |
| 1 | 0 | 0 | 0 | 0 | 2 |
| 1 | 0 | 0 | 0 | 0 | 2 |
| 1 | 0 | 0 | 0 | 0 | 2 |
| 1 | 0 | 0 | 0 | 0 | 2 |
| 1 | 0 | 0 | 0 | 0 | 2 |
| 1 | 0 | 0 | 0 | 0 | 2 |
| 1 | 0 | 0 | 0 | 0 | 2 |
| 1 | 0 | 0 | 0 | 0 | 2 |
| 1 | 0 | 0 | 0 | 0 | 2 |
| 1 | 0 | 0 | 0 | 0 | 2 |
| 1 | 0 | 0 | 0 | 0 | 2 |
| 1 | 0 | 0 | 0 | 0 | 2 |
| 1 | 0 | 0 | 0 | 0 | 2 |
| 1 | 0 | 0 | 0 | 0 | 2 |
| 1 | 0 | 0 | 0 | 0 | 2 |
| 1 | 0 | 0 | 0 | 0 | 2 |
| 1 | 0 | 0 | 0 | 0 | 2 |
| 1 | 0 | 0 | 0 | 0 | 2 |
| 1 | 0 | 0 | 0 | 0 | 2 |
| 1 | 0 | 0 | 0 | 0 | 2 |
| 1 | 0 | 0 | 0 | 0 | 2 |
| 1 | 0 | 0 | 0 | 0 | 2 |
| 1 | 0 | 0 | 0 | 0 | 2 |
| 1 | 0 | 0 | 0 | 0 | 2 |
| 1 | 0 | 0 | 0 | 0 | 2 |
| 1 | 0 | 0 | 0 | 0 | 2 |
| 1 | 0 | 0 | 0 | 0 | 2 |
| 1 | 0 | 0 | 0 | 0 | 2 |
| 1 | 0 | 0 | 0 | 0 | 2 |
| 1 | 0 | 0 | 0 | 0 | 2 |
| 1 | 0 | 0 | 0 | 0 | 2 |
| 1 | 0 | 0 | 0 | 0 | 2 |
| 1 | 0 | 0 | 0 | 0 | 2 |
| 1 | 0 | 0 | 0 | 0 | 2 |
| 1 | 0 | 0 | 0 | 0 | 2 |

| | | | | | |
|---|---|---|---|---|---|
| 1 | 0 | 0 | 0 | 0 | 2 |
| 1 | 0 | 0 | 0 | 0 | 2 |
| 1 | 0 | 0 | 0 | 0 | 2 |
| 1 | 0 | 0 | 0 | 0 | 2 |
| 1 | 0 | 0 | 0 | 0 | 2 |
| 1 | 0 | 0 | 0 | 0 | 2 |
| 1 | 0 | 0 | 0 | 0 | 2 |
| 1 | 0 | 0 | 0 | 0 | 2 |
| 1 | 0 | 0 | 0 | 0 | 2 |
| 1 | 0 | 0 | 0 | 0 | 2 |
| 1 | 0 | 0 | 0 | 0 | 2 |
| 1 | 0 | 0 | 0 | 0 | 2 |
| 1 | 0 | 0 | 0 | 0 | 2 |
| 1 | 0 | 0 | 0 | 0 | 2 |
| 1 | 0 | 0 | 0 | 0 | 2 |
| 1 | 0 | 0 | 0 | 0 | 2 |
| 1 | 0 | 0 | 0 | 0 | 2 |
| 1 | 0 | 0 | 0 | 0 | 2 |
| 1 | 0 | 0 | 0 | 0 | 2 |
| 1 | 0 | 0 | 0 | 0 | 2 |
| 1 | 0 | 0 | 0 | 0 | 2 |
| 1 | 0 | 0 | 0 | 0 | 2 |
| 1 | 0 | 0 | 0 | 0 | 2 |
| 1 | 0 | 0 | 0 | 0 | 2 |
| 1 | 0 | 0 | 0 | 0 | 2 |
| 1 | 0 | 0 | 0 | 0 | 2 |
| 1 | 0 | 0 | 0 | 0 | 2 |
| 1 | 0 | 0 | 0 | 0 | 2 |
| 1 | 0 | 0 | 0 | 0 | 2 |
| 1 | 0 | 0 | 0 | 0 | 2 |
| 1 | 0 | 0 | 0 | 0 | 2 |
| 1 | 0 | 0 | 0 | 0 | 2 |
| 1 | 0 | 0 | 0 | 0 | 2 |
| 1 | 0 | 0 | 0 | 0 | 2 |
| 1 | 0 | 0 | 0 | 0 | 2 |
| 1 | 0 | 0 | 0 | 0 | 2 |
| 1 | 0 | 0 | 0 | 0 | 2 |
| 1 | 0 | 0 | 0 | 0 | 2 |
| 1 | 0 | 0 | 0 | 0 | 2 |
| 1 | 0 | 0 | 0 | 0 | 2 |
| 1 | 0 | 0 | 0 | 0 | 2 |
| 1 | 0 | 0 | 0 | 0 | 2 |
| 1 | 0 | 0 | 0 | 0 | 2 |
| 1 | 0 | 0 | 0 | 0 | 2 |

| | | | | | |
|---|---|---|---|---|---|
| 1 | 0 | 0 | 0 | 0 | 2 |
| 1 | 0 | 0 | 0 | 0 | 2 |
| 1 | 0 | 0 | 0 | 0 | 2 |
| 1 | 0 | 0 | 0 | 0 | 2 |
| 1 | 0 | 0 | 0 | 0 | 2 |
| 1 | 0 | 0 | 0 | 0 | 2 |
| 1 | 0 | 0 | 0 | 0 | 2 |
| 1 | 0 | 0 | 0 | 0 | 2 |
| 1 | 0 | 0 | 0 | 0 | 2 |
| 1 | 0 | 0 | 0 | 0 | 2 |
| 1 | 0 | 0 | 0 | 0 | 2 |
| 1 | 0 | 0 | 0 | 0 | 2 |
| 1 | 0 | 0 | 0 | 0 | 2 |
| 1 | 0 | 0 | 0 | 0 | 2 |
| 1 | 0 | 0 | 0 | 0 | 2 |
| 1 | 0 | 0 | 0 | 0 | 2 |
| 1 | 0 | 0 | 0 | 0 | 2 |
| 1 | 0 | 0 | 0 | 0 | 2 |
| 1 | 0 | 0 | 0 | 0 | 2 |
| 1 | 0 | 0 | 0 | 0 | 2 |
| 1 | 0 | 0 | 0 | 0 | 2 |
| 1 | 0 | 0 | 0 | 0 | 2 |
| 1 | 0 | 0 | 0 | 0 | 2 |
| 1 | 0 | 0 | 0 | 0 | 2 |
| 1 | 0 | 0 | 0 | 0 | 2 |
| 1 | 0 | 0 | 0 | 0 | 2 |
| 1 | 0 | 0 | 0 | 0 | 2 |
| 1 | 0 | 0 | 0 | 0 | 2 |
| 1 | 0 | 0 | 0 | 0 | 2 |
| 1 | 0 | 0 | 0 | 0 | 2 |
| 1 | 0 | 0 | 0 | 0 | 2 |
| 1 | 0 | 0 | 0 | 0 | 2 |
| 1 | 0 | 0 | 0 | 0 | 2 |
| 1 | 0 | 0 | 0 | 0 | 2 |
| 1 | 0 | 0 | 0 | 0 | 2 |
| 1 | 0 | 0 | 0 | 0 | 2 |
| 1 | 0 | 0 | 0 | 0 | 3 |
| 1 | 0 | 0 | 0 | 0 | 3 |
| 1 | 0 | 0 | 0 | 0 | 3 |
| 1 | 0 | 0 | 0 | 0 | 3 |
| 1 | 0 | 0 | 0 | 0 | 3 |
| 1 | 0 | 0 | 0 | 0 | 3 |
| 1 | 0 | 0 | 0 | 0 | 3 |
| 1 | 0 | 0 | 0 | 0 | 3 |

| | | | | | |
|---|---|---|---|---|---|
| 1 | 0 | 0 | 0 | 0 | 3 |
| 1 | 0 | 0 | 0 | 0 | 3 |
| 1 | 0 | 0 | 0 | 0 | 3 |
| 1 | 0 | 0 | 0 | 0 | 3 |
| 1 | 0 | 0 | 0 | 0 | 3 |
| 1 | 0 | 0 | 0 | 0 | 3 |
| 1 | 0 | 0 | 0 | 0 | 3 |
| 1 | 0 | 0 | 0 | 0 | 3 |
| 1 | 0 | 0 | 0 | 0 | 3 |
| 1 | 0 | 0 | 0 | 0 | 3 |
| 1 | 0 | 0 | 0 | 0 | 3 |
| 1 | 0 | 0 | 0 | 0 | 3 |
| 1 | 0 | 0 | 0 | 0 | 2 |
| 1 | 0 | 0 | 0 | 0 | 2 |
| 1 | 0 | 0 | 0 | 0 | 2 |
| 1 | 0 | 0 | 0 | 0 | 2 |
| 1 | 0 | 0 | 0 | 0 | 2 |
| 1 | 0 | 0 | 0 | 0 | 2 |
| 1 | 0 | 0 | 0 | 0 | 2 |
| 1 | 0 | 0 | 0 | 0 | 2 |
| 1 | 0 | 0 | 0 | 0 | 2 |
| 1 | 0 | 0 | 0 | 0 | 2 |
| 1 | 0 | 0 | 0 | 0 | 2 |
| 1 | 0 | 0 | 0 | 0 | 2 |
| 1 | 0 | 0 | 0 | 0 | 2 |
| 1 | 0 | 0 | 0 | 0 | 2 |
| 1 | 0 | 0 | 0 | 0 | 2 |
| 1 | 0 | 0 | 0 | 0 | 2 |
| 1 | 0 | 0 | 0 | 0 | 2 |
| 1 | 0 | 0 | 0 | 0 | 2 |
| 1 | 0 | 0 | 0 | 0 | 2 |
| 1 | 0 | 0 | 0 | 0 | 2 |
| 1 | 0 | 0 | 0 | 0 | 2 |
| 1 | 0 | 0 | 0 | 0 | 2 |
| 1 | 0 | 0 | 0 | 0 | 2 |
| 1 | 0 | 0 | 0 | 0 | 2 |
| 1 | 0 | 0 | 0 | 0 | 2 |
| 1 | 0 | 0 | 0 | 0 | 2 |
| 1 | 0 | 0 | 0 | 0 | 2 |
| 1 | 0 | 0 | 0 | 0 | 2 |
| 1 | 0 | 0 | 0 | 0 | 2 |
| 1 | 0 | 0 | 0 | 0 | 2 |
| 1 | 0 | 0 | 0 | 0 | 2 |
| 1 | 0 | 0 | 0 | 0 | 2 |

| | | | | | |
|---|---|---|---|---|---|
| 1 | 0 | 0 | 0 | 0 | 2 |
| 1 | 0 | 0 | 0 | 0 | 2 |
| 1 | 0 | 0 | 0 | 0 | 2 |
| 1 | 0 | 0 | 0 | 0 | 2 |
| 1 | 0 | 0 | 0 | 0 | 2 |
| 1 | 0 | 0 | 0 | 0 | 2 |
| 1 | 0 | 0 | 0 | 0 | 2 |
| 1 | 0 | 0 | 0 | 0 | 2 |
| 1 | 0 | 0 | 0 | 0 | 3 |
| 1 | 0 | 0 | 0 | 0 | 3 |
| 1 | 0 | 0 | 0 | 0 | 3 |
| 1 | 0 | 0 | 0 | 0 | 3 |
| 1 | 0 | 0 | 0 | 0 | 3 |
| 1 | 0 | 0 | 0 | 0 | 3 |
| 1 | 0 | 0 | 0 | 0 | 3 |
| 1 | 0 | 0 | 0 | 0 | 3 |
| 1 | 0 | 0 | 0 | 0 | 3 |
| 1 | 0 | 0 | 0 | 0 | 3 |
| 1 | 0 | 0 | 0 | 0 | 3 |
| 1 | 0 | 0 | 0 | 0 | 3 |
| 1 | 0 | 0 | 0 | 0 | 3 |
| 1 | 0 | 0 | 0 | 0 | 3 |
| 1 | 0 | 0 | 0 | 0 | 3 |
| 1 | 0 | 0 | 0 | 0 | 3 |
| 1 | 0 | 0 | 0 | 0 | 3 |
| 1 | 0 | 0 | 0 | 0 | 3 |
| 1 | 0 | 0 | 0 | 0 | 3 |
| 1 | 0 | 0 | 0 | 0 | 3 |
| 1 | 0 | 0 | 0 | 0 | 3 |
| 1 | 0 | 0 | 0 | 0 | 3 |
| 1 | 0 | 0 | 0 | 0 | 3 |
| 1 | 0 | 0 | 0 | 0 | 3 |
| 1 | 0 | 0 | 0 | 0 | 3 |
| 1 | 0 | 0 | 0 | 0 | 3 |
| 1 | 0 | 0 | 0 | 0 | 3 |
| 1 | 0 | 0 | 0 | 0 | 3 |
| 1 | 0 | 0 | 0 | 0 | 3 |
| 1 | 0 | 0 | 0 | 0 | 3 |
| 1 | 0 | 0 | 0 | 0 | 3 |
| 1 | 0 | 0 | 0 | 0 | 3 |
| 1 | 0 | 0 | 0 | 0 | 3 |
| 1 | 0 | 0 | 0 | 0 | 3 |
| 1 | 0 | 0 | 0 | 0 | 3 |
| 1 | 0 | 0 | 0 | 0 | 3 |

| | | | | | |
|---|---|---|---|---|---|
| 1 | 0 | 0 | 0 | 0 | 3 |
| 1 | 0 | 0 | 0 | 0 | 3 |
| 1 | 0 | 0 | 0 | 0 | 3 |
| 1 | 0 | 0 | 0 | 0 | 3 |
| 1 | 0 | 0 | 0 | 0 | 2 |
| 1 | 0 | 0 | 0 | 0 | 2 |
| 1 | 0 | 0 | 0 | 0 | 2 |
| 1 | 0 | 0 | 0 | 0 | 2 |
| 1 | 0 | 0 | 0 | 0 | 2 |
| 1 | 0 | 0 | 0 | 0 | 2 |
| 1 | 0 | 0 | 0 | 0 | 2 |
| 1 | 0 | 0 | 0 | 0 | 2 |
| 1 | 0 | 0 | 0 | 0 | 2 |
| 1 | 0 | 0 | 0 | 0 | 2 |
| 1 | 0 | 0 | 0 | 0 | 2 |
| 1 | 0 | 0 | 0 | 0 | 2 |
| 1 | 0 | 0 | 0 | 0 | 2 |
| 1 | 0 | 0 | 0 | 0 | 2 |
| 1 | 0 | 0 | 0 | 0 | 2 |
| 1 | 0 | 0 | 0 | 0 | 2 |
| 1 | 0 | 0 | 0 | 0 | 2 |
| 1 | 0 | 0 | 0 | 0 | 2 |
| 1 | 0 | 0 | 0 | 0 | 2 |
| 1 | 0 | 0 | 0 | 0 | 2 |
| 1 | 0 | 0 | 0 | 0 | 2 |
| 1 | 0 | 0 | 0 | 0 | 2 |
| 1 | 0 | 0 | 0 | 0 | 2 |
| 1 | 0 | 0 | 0 | 0 | 2 |
| 1 | 0 | 0 | 0 | 0 | 2 |
| 1 | 0 | 0 | 0 | 0 | 2 |
| 1 | 0 | 0 | 0 | 0 | 2 |
| 1 | 0 | 0 | 0 | 0 | 2 |
| 1 | 0 | 0 | 0 | 0 | 2 |
| 1 | 0 | 0 | 0 | 0 | 2 |
| 1 | 0 | 0 | 0 | 0 | 2 |
| 1 | 0 | 0 | 0 | 0 | 2 |
| 1 | 0 | 0 | 0 | 0 | 2 |
| 1 | 0 | 0 | 0 | 0 | 2 |
| 1 | 0 | 0 | 0 | 0 | 2 |
| 1 | 0 | 0 | 0 | 0 | 2 |
| 1 | 0 | 0 | 0 | 0 | 2 |
| 1 | 0 | 0 | 0 | 0 | 2 |
| 1 | 0 | 0 | 0 | 0 | 2 |
| 1 | 0 | 0 | 0 | 0 | 2 |

| | | | | | |
|---|---|---|---|---|---|
| 1 | 0 | 0 | 0 | 0 | 3 |
| 1 | 0 | 0 | 0 | 0 | 3 |
| 1 | 0 | 0 | 0 | 0 | 3 |
| 1 | 0 | 0 | 0 | 0 | 3 |
| 1 | 0 | 0 | 0 | 0 | 3 |
| 1 | 0 | 0 | 0 | 0 | 3 |
| 1 | 0 | 0 | 0 | 0 | 3 |
| 1 | 0 | 0 | 0 | 0 | 3 |
| 1 | 0 | 0 | 0 | 0 | 3 |
| 1 | 0 | 0 | 0 | 0 | 3 |
| 1 | 0 | 0 | 0 | 0 | 3 |
| 1 | 0 | 0 | 0 | 0 | 3 |
| 1 | 0 | 0 | 0 | 0 | 3 |
| 1 | 0 | 0 | 0 | 0 | 3 |
| 1 | 0 | 0 | 0 | 0 | 3 |
| 1 | 0 | 0 | 0 | 0 | 3 |
| 1 | 0 | 0 | 0 | 0 | 3 |
| 1 | 0 | 0 | 0 | 0 | 3 |
| 1 | 0 | 0 | 0 | 0 | 3 |
| 1 | 0 | 0 | 0 | 0 | 3 |
| 1 | 0 | 0 | 0 | 0 | 3 |
| 1 | 0 | 0 | 0 | 0 | 3 |
| 1 | 0 | 0 | 0 | 0 | 3 |
| 1 | 0 | 0 | 0 | 0 | 3 |
| 1 | 0 | 0 | 0 | 0 | 3 |
| 1 | 0 | 0 | 0 | 0 | 3 |
| 1 | 0 | 0 | 0 | 0 | 3 |
| 1 | 0 | 0 | 0 | 0 | 3 |
| 1 | 0 | 0 | 0 | 0 | 3 |
| 1 | 0 | 0 | 0 | 0 | 3 |
| 1 | 0 | 0 | 0 | 0 | 3 |
| 1 | 0 | 0 | 0 | 0 | 3 |
| 1 | 0 | 0 | 0 | 0 | 3 |
| 1 | 0 | 0 | 0 | 0 | 3 |
| 1 | 0 | 0 | 0 | 0 | 3 |
| 1 | 0 | 0 | 0 | 0 | 3 |
| 1 | 0 | 0 | 0 | 0 | 3 |
| 1 | 0 | 0 | 0 | 0 | 3 |
| 1 | 0 | 0 | 0 | 0 | 3 |
| 1 | 0 | 0 | 0 | 0 | 3 |
| 0 | 0 | 0 | 0 | 1 | 1 |
| 0 | 0 | 0 | 0 | 1 | 1 |
| 0 | 0 | 0 | 0 | 1 | 1 |
| 0 | 0 | 0 | 0 | 1 | 1 |

| | | | | | |
|---|---|---|---|---|---|
| 0 | 0 | 0 | 0 | 1 | 1 |
| 0 | 0 | 0 | 0 | 1 | 1 |
| 0 | 0 | 0 | 0 | 1 | 1 |
| 0 | 0 | 0 | 0 | 1 | 1 |
| 0 | 0 | 0 | 0 | 1 | 1 |
| 0 | 0 | 0 | 0 | 1 | 1 |
| 0 | 0 | 0 | 0 | 1 | 1 |
| 0 | 0 | 0 | 0 | 1 | 1 |
| 0 | 0 | 0 | 0 | 1 | 1 |
| 0 | 0 | 0 | 0 | 1 | 1 |
| 0 | 0 | 0 | 0 | 1 | 1 |
| 0 | 0 | 0 | 0 | 1 | 1 |
| 0 | 0 | 0 | 0 | 1 | 1 |
| 0 | 0 | 0 | 0 | 1 | 1 |
| 0 | 0 | 0 | 0 | 1 | 1 |
| 0 | 0 | 0 | 0 | 1 | 1 |
| 0 | 0 | 0 | 0 | 1 | 1 |
| 0 | 0 | 0 | 0 | 1 | 1 |
| 0 | 0 | 0 | 0 | 1 | 1 |
| 0 | 0 | 0 | 0 | 1 | 1 |
| 0 | 0 | 0 | 0 | 1 | 1 |
| 0 | 0 | 0 | 0 | 1 | 1 |
| 0 | 0 | 0 | 0 | 1 | 1 |
| 0 | 0 | 0 | 0 | 1 | 1 |
| 0 | 0 | 0 | 0 | 1 | 1 |
| 0 | 0 | 0 | 0 | 1 | 1 |
| 0 | 0 | 0 | 0 | 1 | 1 |
| 0 | 0 | 0 | 0 | 1 | 1 |
| 0 | 0 | 0 | 0 | 1 | 1 |
| 0 | 0 | 0 | 0 | 1 | 1 |
| 0 | 0 | 0 | 0 | 1 | 1 |
| 0 | 0 | 0 | 0 | 1 | 1 |
| 0 | 0 | 0 | 0 | 1 | 1 |
| 0 | 0 | 0 | 0 | 1 | 1 |
| 0 | 0 | 0 | 0 | 1 | 1 |
| 0 | 0 | 0 | 0 | 1 | 1 |
| 0 | 0 | 0 | 0 | 1 | 1 |
| 0 | 0 | 0 | 0 | 1 | 1 |
| 0 | 0 | 0 | 0 | 1 | 1 |
| 0 | 0 | 0 | 0 | 1 | 1 |
| 0 | 0 | 0 | 0 | 1 | 1 |
| 0 | 0 | 0 | 0 | 1 | 1 |
| 0 | 0 | 0 | 0 | 1 | 1 |
| 0 | 0 | 0 | 0 | 1 | 1 |

| | | | | | |
|---|---|---|---|---|---|
| 0 | 0 | 0 | 0 | 1 | 1 |
| 0 | 0 | 0 | 0 | 1 | 1 |
| 0 | 0 | 0 | 0 | 1 | 1 |
| 0 | 0 | 0 | 0 | 1 | 1 |
| 0 | 0 | 0 | 0 | 1 | 1 |
| 0 | 0 | 0 | 0 | 1 | 1 |
| 0 | 0 | 0 | 0 | 1 | 1 |
| 0 | 0 | 0 | 0 | 1 | 1 |
| 0 | 0 | 0 | 0 | 1 | 1 |
| 0 | 0 | 0 | 0 | 1 | 1 |
| 0 | 0 | 0 | 0 | 1 | 1 |
| 0 | 0 | 0 | 0 | 1 | 1 |
| 0 | 0 | 0 | 0 | 1 | 1 |
| 0 | 0 | 0 | 0 | 1 | 1 |
| 0 | 0 | 0 | 0 | 1 | 1 |
| 0 | 0 | 0 | 0 | 1 | 1 |
| 0 | 0 | 0 | 0 | 1 | 1 |
| 0 | 0 | 0 | 0 | 1 | 1 |
| 0 | 0 | 0 | 0 | 1 | 1 |
| 0 | 0 | 0 | 0 | 1 | 1 |
| 0 | 0 | 0 | 0 | 1 | 1 |
| 0 | 0 | 0 | 0 | 1 | 1 |
| 0 | 0 | 0 | 0 | 1 | 1 |
| 0 | 0 | 0 | 0 | 1 | 1 |
| 0 | 0 | 0 | 0 | 1 | 1 |
| 0 | 0 | 0 | 0 | 1 | 1 |
| 0 | 0 | 0 | 0 | 1 | 1 |
| 0 | 0 | 0 | 0 | 1 | 1 |
| 0 | 0 | 0 | 0 | 1 | 1 |
| 0 | 0 | 0 | 0 | 1 | 1 |
| 0 | 0 | 0 | 0 | 1 | 1 |
| 0 | 0 | 0 | 0 | 1 | 1 |
| 0 | 0 | 0 | 0 | 1 | 1 |
| 0 | 0 | 0 | 0 | 1 | 1 |
| 0 | 0 | 0 | 0 | 1 | 1 |
| 0 | 0 | 0 | 0 | 1 | 1 |
| 0 | 0 | 0 | 0 | 1 | 1 |
| 0 | 0 | 0 | 0 | 1 | 1 |
| 0 | 0 | 0 | 0 | 1 | 1 |
| 0 | 0 | 0 | 0 | 1 | 1 |
| 0 | 0 | 0 | 0 | 1 | 1 |
| 0 | 0 | 0 | 0 | 1 | 1 |
| 0 | 0 | 0 | 0 | 1 | 1 |
| 0 | 0 | 0 | 0 | 1 | 1 |

| | | | | | |
|---|---|---|---|---|---|
| 0 | 0 | 0 | 0 | 1 | 1 |
| 0 | 0 | 0 | 0 | 1 | 1 |
| 0 | 0 | 0 | 0 | 1 | 1 |
| 0 | 0 | 0 | 0 | 1 | 1 |
| 0 | 0 | 0 | 0 | 1 | 1 |
| 0 | 0 | 0 | 0 | 1 | 1 |
| 0 | 0 | 0 | 0 | 1 | 1 |
| 0 | 0 | 0 | 0 | 1 | 1 |
| 0 | 0 | 0 | 0 | 1 | 1 |
| 0 | 0 | 0 | 0 | 1 | 1 |
| 0 | 0 | 0 | 0 | 1 | 1 |
| 0 | 0 | 0 | 0 | 1 | 1 |
| 0 | 0 | 0 | 0 | 1 | 1 |
| 0 | 0 | 0 | 0 | 1 | 1 |
| 0 | 0 | 0 | 0 | 1 | 1 |
| 0 | 0 | 0 | 0 | 1 | 1 |
| 0 | 0 | 0 | 0 | 1 | 1 |
| 0 | 0 | 0 | 0 | 1 | 1 |
| 0 | 0 | 0 | 0 | 1 | 1 |
| 0 | 0 | 0 | 0 | 1 | 1 |
| 0 | 0 | 0 | 0 | 1 | 1 |
| 0 | 0 | 0 | 0 | 1 | 1 |
| 0 | 0 | 0 | 0 | 1 | 1 |
| 0 | 0 | 0 | 0 | 1 | 1 |
| 0 | 0 | 0 | 0 | 1 | 1 |
| 0 | 0 | 0 | 0 | 1 | 1 |
| 0 | 0 | 0 | 0 | 1 | 1 |
| 0 | 0 | 0 | 0 | 1 | 1 |
| 0 | 0 | 0 | 0 | 1 | 1 |
| 0 | 0 | 0 | 0 | 1 | 1 |
| 0 | 0 | 0 | 0 | 1 | 1 |
| 0 | 0 | 0 | 0 | 1 | 1 |
| 0 | 0 | 0 | 0 | 1 | 1 |
| 0 | 0 | 0 | 0 | 1 | 1 |
| 0 | 0 | 0 | 0 | 1 | 1 |
| 0 | 0 | 0 | 0 | 1 | 1 |
| 0 | 0 | 0 | 0 | 1 | 1 |
| 0 | 0 | 0 | 0 | 1 | 1 |
| 0 | 0 | 0 | 0 | 1 | 1 |
| 0 | 0 | 0 | 0 | 1 | 1 |
| 0 | 0 | 0 | 0 | 1 | 1 |
| 0 | 0 | 0 | 0 | 1 | 1 |
| 0 | 0 | 0 | 0 | 1 | 1 |
| 0 | 0 | 0 | 0 | 1 | 1 |

| | | | | | |
|---|---|---|---|---|---|
| 0 | 0 | 0 | 0 | 1 | 1 |
| 0 | 0 | 0 | 0 | 1 | 1 |
| 0 | 0 | 0 | 0 | 1 | 1 |
| 0 | 0 | 0 | 0 | 1 | 1 |
| 0 | 0 | 0 | 0 | 1 | 1 |
| 0 | 0 | 0 | 0 | 1 | 1 |
| 0 | 0 | 0 | 0 | 1 | 1 |
| 0 | 0 | 0 | 0 | 1 | 1 |
| 0 | 0 | 0 | 0 | 1 | 1 |
| 0 | 0 | 0 | 0 | 1 | 1 |
| 0 | 0 | 0 | 0 | 1 | 1 |
| 0 | 0 | 0 | 0 | 1 | 1 |
| 0 | 0 | 0 | 0 | 1 | 1 |
| 0 | 0 | 0 | 0 | 1 | 1 |
| 0 | 0 | 0 | 0 | 1 | 1 |
| 0 | 0 | 0 | 0 | 1 | 1 |
| 0 | 0 | 0 | 0 | 1 | 1 |
| 0 | 0 | 0 | 0 | 1 | 1 |
| 0 | 0 | 0 | 0 | 1 | 1 |
| 0 | 0 | 0 | 0 | 1 | 1 |
| 0 | 0 | 0 | 0 | 1 | 1 |
| 0 | 0 | 0 | 0 | 1 | 1 |
| 0 | 0 | 0 | 0 | 1 | 1 |
| 0 | 0 | 0 | 0 | 1 | 1 |
| 0 | 0 | 0 | 0 | 1 | 1 |
| 0 | 0 | 0 | 0 | 1 | 1 |
| 0 | 0 | 0 | 0 | 1 | 1 |
| 0 | 0 | 0 | 0 | 1 | 1 |
| 0 | 0 | 0 | 0 | 1 | 1 |
| 0 | 0 | 0 | 0 | 1 | 1 |
| 0 | 0 | 0 | 0 | 1 | 1 |
| 0 | 0 | 0 | 0 | 1 | 1 |
| 0 | 0 | 0 | 0 | 1 | 1 |
| 0 | 0 | 0 | 0 | 1 | 1 |
| 0 | 0 | 0 | 0 | 1 | 1 |
| 0 | 0 | 0 | 0 | 1 | 1 |
| 0 | 0 | 0 | 0 | 1 | 1 |
| 0 | 0 | 0 | 0 | 1 | 1 |
| 0 | 0 | 0 | 0 | 1 | 1 |
| 0 | 0 | 0 | 0 | 1 | 1 |
| 0 | 0 | 0 | 0 | 1 | 1 |
| 0 | 0 | 0 | 0 | 1 | 1 |
| 0 | 0 | 0 | 0 | 1 | 1 |
| 0 | 0 | 0 | 0 | 1 | 1 |

| | | | | | |
|---|---|---|---|---|---|
| 0 | 0 | 0 | 0 | 1 | 1 |
| 0 | 0 | 0 | 0 | 1 | 1 |
| 0 | 0 | 0 | 0 | 1 | 1 |
| 0 | 0 | 0 | 0 | 1 | 1 |
| 0 | 0 | 0 | 0 | 1 | 1 |
| 0 | 0 | 0 | 0 | 1 | 1 |
| 0 | 0 | 0 | 0 | 1 | 1 |
| 0 | 0 | 0 | 0 | 1 | 1 |
| 0 | 0 | 0 | 0 | 1 | 1 |
| 0 | 0 | 0 | 0 | 1 | 1 |
| 0 | 0 | 0 | 0 | 1 | 1 |
| 0 | 0 | 0 | 0 | 1 | 1 |
| 0 | 0 | 0 | 0 | 1 | 1 |
| 0 | 0 | 0 | 0 | 1 | 1 |
| 0 | 0 | 0 | 0 | 1 | 1 |
| 0 | 0 | 0 | 0 | 1 | 1 |
| 0 | 0 | 0 | 0 | 1 | 1 |
| 0 | 0 | 0 | 0 | 1 | 1 |
| 0 | 0 | 0 | 0 | 1 | 1 |
| 0 | 0 | 0 | 0 | 1 | 1 |
| 0 | 0 | 0 | 0 | 1 | 1 |
| 0 | 0 | 0 | 0 | 1 | 1 |
| 0 | 0 | 0 | 0 | 1 | 1 |
| 0 | 0 | 0 | 0 | 1 | 1 |
| 0 | 0 | 0 | 0 | 1 | 1 |
| 0 | 0 | 0 | 0 | 1 | 1 |
| 0 | 0 | 0 | 0 | 1 | 1 |
| 0 | 0 | 0 | 0 | 1 | 1 |
| 0 | 0 | 0 | 0 | 1 | 1 |
| 0 | 0 | 0 | 0 | 1 | 1 |
| 0 | 0 | 0 | 0 | 1 | 1 |
| 0 | 0 | 0 | 0 | 1 | 1 |
| 0 | 0 | 0 | 0 | 1 | 1 |
| 0 | 0 | 0 | 0 | 1 | 1 |
| 0 | 0 | 0 | 0 | 1 | 1 |
| 0 | 0 | 0 | 0 | 1 | 1 |
| 0 | 0 | 0 | 0 | 1 | 1 |
| 0 | 0 | 0 | 0 | 1 | 1 |
| 0 | 0 | 0 | 0 | 1 | 1 |
| 0 | 0 | 0 | 0 | 1 | 1 |
| 1 | 0 | 0 | 0 | 0 | 2 |
| 1 | 0 | 0 | 0 | 0 | 2 |
| 1 | 0 | 0 | 0 | 0 | 2 |
| 1 | 0 | 0 | 0 | 0 | 2 |

| | | | | | |
|---|---|---|---|---|---|
| 1 | 0 | 0 | 0 | 0 | 2 |
| 1 | 0 | 0 | 0 | 0 | 2 |
| 1 | 0 | 0 | 0 | 0 | 2 |
| 1 | 0 | 0 | 0 | 0 | 2 |
| 1 | 0 | 0 | 0 | 0 | 2 |
| 1 | 0 | 0 | 0 | 0 | 2 |
| 1 | 0 | 0 | 0 | 0 | 2 |
| 1 | 0 | 0 | 0 | 0 | 2 |
| 1 | 0 | 0 | 0 | 0 | 2 |
| 1 | 0 | 0 | 0 | 0 | 2 |
| 1 | 0 | 0 | 0 | 0 | 2 |
| 1 | 0 | 0 | 0 | 0 | 2 |
| 1 | 0 | 0 | 0 | 0 | 2 |
| 1 | 0 | 0 | 0 | 0 | 2 |
| 1 | 0 | 0 | 0 | 0 | 2 |
| 1 | 0 | 0 | 0 | 0 | 2 |
| 1 | 0 | 0 | 0 | 0 | 2 |
| 1 | 0 | 0 | 0 | 0 | 2 |
| 1 | 0 | 0 | 0 | 0 | 2 |
| 1 | 0 | 0 | 0 | 0 | 2 |
| 1 | 0 | 0 | 0 | 0 | 2 |
| 1 | 0 | 0 | 0 | 0 | 2 |
| 1 | 0 | 0 | 0 | 0 | 2 |
| 1 | 0 | 0 | 0 | 0 | 2 |
| 1 | 0 | 0 | 0 | 0 | 2 |
| 1 | 0 | 0 | 0 | 0 | 2 |
| 1 | 0 | 0 | 0 | 0 | 2 |
| 1 | 0 | 0 | 0 | 0 | 2 |
| 1 | 0 | 0 | 0 | 0 | 2 |
| 1 | 0 | 0 | 0 | 0 | 2 |
| 1 | 0 | 0 | 0 | 0 | 2 |
| 1 | 0 | 0 | 0 | 0 | 2 |
| 1 | 0 | 0 | 0 | 0 | 2 |
| 1 | 0 | 0 | 0 | 0 | 2 |
| 1 | 0 | 0 | 0 | 0 | 2 |
| 1 | 0 | 0 | 0 | 0 | 2 |
| 1 | 0 | 0 | 0 | 0 | 2 |
| 1 | 0 | 0 | 0 | 0 | 2 |
| 1 | 0 | 0 | 0 | 0 | 2 |
| 1 | 0 | 0 | 0 | 0 | 2 |
| 1 | 0 | 0 | 0 | 0 | 2 |
| 1 | 0 | 0 | 0 | 0 | 2 |
| 1 | 0 | 0 | 0 | 0 | 2 |
| 1 | 0 | 0 | 0 | 0 | 2 |

| | | | | | |
|---|---|---|---|---|---|
| 1 | 0 | 0 | 0 | 0 | 2 |
| 1 | 0 | 0 | 0 | 0 | 2 |
| 1 | 0 | 0 | 0 | 0 | 2 |
| 1 | 0 | 0 | 0 | 0 | 2 |
| 1 | 0 | 0 | 0 | 0 | 2 |
| 1 | 0 | 0 | 0 | 0 | 2 |
| 1 | 0 | 0 | 0 | 0 | 2 |
| 1 | 0 | 0 | 0 | 0 | 2 |
| 1 | 0 | 0 | 0 | 0 | 2 |
| 1 | 0 | 0 | 0 | 0 | 2 |
| 1 | 0 | 0 | 0 | 0 | 2 |
| 1 | 0 | 0 | 0 | 0 | 2 |
| 1 | 0 | 0 | 0 | 0 | 2 |
| 1 | 0 | 0 | 0 | 0 | 2 |
| 1 | 0 | 0 | 0 | 0 | 2 |
| 1 | 0 | 0 | 0 | 0 | 2 |
| 1 | 0 | 0 | 0 | 0 | 2 |
| 1 | 0 | 0 | 0 | 0 | 2 |
| 1 | 0 | 0 | 0 | 0 | 2 |
| 1 | 0 | 0 | 0 | 0 | 2 |
| 1 | 0 | 0 | 0 | 0 | 2 |
| 1 | 0 | 0 | 0 | 0 | 2 |
| 1 | 0 | 0 | 0 | 0 | 2 |
| 1 | 0 | 0 | 0 | 0 | 2 |
| 1 | 0 | 0 | 0 | 0 | 2 |
| 1 | 0 | 0 | 0 | 0 | 2 |
| 1 | 0 | 0 | 0 | 0 | 2 |
| 1 | 0 | 0 | 0 | 0 | 2 |
| 1 | 0 | 0 | 0 | 0 | 2 |
| 1 | 0 | 0 | 0 | 0 | 2 |
| 1 | 0 | 0 | 0 | 0 | 2 |
| 1 | 0 | 0 | 0 | 0 | 2 |
| 1 | 0 | 0 | 0 | 0 | 2 |
| 1 | 0 | 0 | 0 | 0 | 2 |
| 1 | 0 | 0 | 0 | 0 | 2 |
| 1 | 0 | 0 | 0 | 0 | 2 |
| 1 | 0 | 0 | 0 | 0 | 2 |
| 1 | 0 | 0 | 0 | 0 | 2 |
| 1 | 0 | 0 | 0 | 0 | 2 |
| 1 | 0 | 0 | 0 | 0 | 2 |
| 1 | 0 | 0 | 0 | 0 | 2 |
| 1 | 0 | 0 | 0 | 0 | 2 |
| 1 | 0 | 0 | 0 | 0 | 2 |
| 1 | 0 | 0 | 0 | 0 | 2 |

| | | | | | |
|---|---|---|---|---|---|
| 1 | 0 | 0 | 0 | 0 | 2 |
| 1 | 0 | 0 | 0 | 0 | 2 |
| 1 | 0 | 0 | 0 | 0 | 2 |
| 1 | 0 | 0 | 0 | 0 | 2 |
| 1 | 0 | 0 | 0 | 0 | 2 |
| 1 | 0 | 0 | 0 | 0 | 2 |
| 1 | 0 | 0 | 0 | 0 | 2 |
| 1 | 0 | 0 | 0 | 0 | 2 |
| 0 | 1 | 1 | 1 | 0 | 4 |
| 0 | 1 | 1 | 1 | 0 | 4 |
| 0 | 1 | 1 | 1 | 0 | 4 |
| 0 | 1 | 1 | 1 | 0 | 4 |
| 0 | 1 | 1 | 1 | 0 | 4 |
| 0 | 1 | 1 | 1 | 0 | 4 |
| 0 | 1 | 1 | 1 | 0 | 4 |
| 0 | 1 | 1 | 1 | 0 | 4 |
| 0 | 1 | 1 | 1 | 0 | 4 |
| 0 | 1 | 1 | 1 | 0 | 4 |
| 0 | 1 | 1 | 1 | 0 | 4 |
| 0 | 1 | 1 | 1 | 0 | 4 |
| 0 | 1 | 1 | 1 | 0 | 4 |
| 0 | 1 | 1 | 1 | 0 | 4 |
| 0 | 1 | 1 | 1 | 0 | 4 |
| 0 | 1 | 1 | 1 | 0 | 4 |
| 0 | 1 | 1 | 1 | 0 | 4 |
| 0 | 1 | 1 | 1 | 0 | 4 |
| 0 | 1 | 1 | 1 | 0 | 4 |
| 0 | 1 | 1 | 1 | 0 | 4 |
| 0 | 1 | 1 | 1 | 0 | 4 |
| 0 | 1 | 1 | 1 | 0 | 4 |
| 0 | 1 | 1 | 1 | 0 | 4 |
| 0 | 1 | 1 | 1 | 0 | 4 |
| 0 | 1 | 1 | 1 | 0 | 4 |
| 0 | 1 | 1 | 1 | 0 | 4 |
| 0 | 1 | 1 | 1 | 0 | 4 |
| 0 | 1 | 1 | 1 | 0 | 4 |
| 0 | 1 | 1 | 1 | 0 | 4 |
| 0 | 1 | 1 | 1 | 0 | 4 |
| 0 | 1 | 1 | 1 | 0 | 4 |
| 0 | 1 | 1 | 1 | 0 | 4 |
| 0 | 1 | 1 | 1 | 0 | 4 |
| 0 | 1 | 1 | 1 | 0 | 4 |
| 0 | 1 | 1 | 1 | 0 | 4 |
| 0 | 1 | 1 | 1 | 0 | 4 |

| | | | | | |
|---|---|---|---|---|---|
| 0 | 1 | 1 | 1 | 0 | 4 |
| 0 | 1 | 1 | 1 | 0 | 4 |
| 0 | 1 | 1 | 1 | 0 | 4 |
| 0 | 1 | 1 | 1 | 0 | 4 |
| 0 | 1 | 1 | 1 | 0 | 4 |
| 0 | 1 | 1 | 1 | 0 | 4 |
| 0 | 1 | 1 | 1 | 0 | 4 |
| 0 | 1 | 1 | 1 | 0 | 4 |
| 0 | 1 | 1 | 1 | 0 | 4 |
| 0 | 1 | 1 | 1 | 0 | 4 |
| 0 | 1 | 1 | 1 | 0 | 4 |
| 0 | 1 | 1 | 1 | 0 | 4 |
| 0 | 1 | 1 | 1 | 0 | 4 |
| 0 | 1 | 1 | 1 | 0 | 4 |
| 0 | 1 | 1 | 1 | 0 | 4 |
| 0 | 1 | 1 | 1 | 0 | 4 |
| 0 | 1 | 1 | 1 | 0 | 4 |
| 0 | 1 | 1 | 1 | 0 | 4 |
| 0 | 1 | 1 | 1 | 0 | 4 |
| 0 | 1 | 1 | 1 | 0 | 4 |
| 0 | 1 | 1 | 1 | 0 | 4 |
| 0 | 1 | 1 | 1 | 0 | 4 |
| 0 | 1 | 1 | 1 | 0 | 4 |
| 0 | 1 | 1 | 1 | 0 | 4 |
| 0 | 1 | 1 | 1 | 0 | 4 |
| 0 | 1 | 1 | 1 | 0 | 4 |
| 0 | 1 | 1 | 1 | 0 | 4 |
| 0 | 1 | 1 | 1 | 0 | 4 |
| 0 | 1 | 1 | 1 | 0 | 4 |
| 0 | 1 | 1 | 1 | 0 | 4 |
| 0 | 1 | 1 | 1 | 0 | 4 |
| 0 | 1 | 1 | 1 | 0 | 4 |
| 0 | 1 | 1 | 1 | 0 | 4 |
| 0 | 1 | 1 | 1 | 0 | 4 |
| 0 | 1 | 1 | 1 | 0 | 4 |
| 0 | 1 | 1 | 1 | 0 | 4 |
| 0 | 1 | 1 | 1 | 0 | 4 |
| 0 | 1 | 1 | 1 | 0 | 4 |
| 0 | 1 | 1 | 1 | 0 | 4 |
| 0 | 1 | 1 | 1 | 0 | 4 |
| 0 | 1 | 1 | 1 | 0 | 4 |
| 0 | 1 | 1 | 1 | 0 | 4 |
| 0 | 1 | 1 | 1 | 0 | 4 |
| 0 | 1 | 1 | 1 | 0 | 4 |

| | | | | | |
|---|---|---|---|---|---|
| 0 | 1 | 1 | 1 | 0 | 4 |
| 0 | 1 | 1 | 1 | 0 | 4 |
| 0 | 1 | 1 | 1 | 0 | 4 |
| 0 | 1 | 1 | 1 | 0 | 4 |
| 0 | 1 | 1 | 1 | 0 | 4 |
| 0 | 1 | 1 | 1 | 0 | 4 |
| 0 | 1 | 1 | 1 | 0 | 4 |
| 0 | 1 | 1 | 1 | 0 | 4 |
| 0 | 1 | 1 | 1 | 0 | 4 |
| 0 | 1 | 1 | 1 | 0 | 4 |
| 0 | 1 | 1 | 1 | 0 | 4 |
| 0 | 1 | 1 | 1 | 0 | 4 |
| 0 | 1 | 1 | 1 | 0 | 4 |
| 0 | 1 | 1 | 1 | 0 | 4 |
| 0 | 1 | 1 | 1 | 0 | 4 |
| 0 | 1 | 1 | 1 | 0 | 4 |
| 0 | 1 | 1 | 1 | 0 | 4 |
| 0 | 1 | 1 | 1 | 0 | 4 |
| 0 | 1 | 1 | 1 | 0 | 4 |
| 0 | 1 | 1 | 1 | 0 | 4 |
| 0 | 0 | 0 | 0 | 1 | 1 |
| 0 | 0 | 0 | 0 | 1 | 1 |
| 0 | 0 | 0 | 0 | 1 | 1 |
| 0 | 0 | 0 | 0 | 1 | 1 |
| 0 | 0 | 0 | 0 | 1 | 1 |
| 0 | 0 | 0 | 0 | 1 | 1 |
| 0 | 0 | 0 | 0 | 1 | 1 |
| 0 | 0 | 0 | 0 | 1 | 1 |
| 0 | 0 | 0 | 0 | 1 | 1 |
| 0 | 0 | 0 | 0 | 1 | 1 |
| 0 | 0 | 0 | 0 | 1 | 1 |
| 0 | 0 | 0 | 0 | 1 | 1 |
| 0 | 0 | 0 | 0 | 1 | 1 |
| 0 | 0 | 0 | 0 | 1 | 1 |
| 0 | 0 | 0 | 0 | 1 | 1 |
| 0 | 0 | 0 | 0 | 1 | 1 |
| 0 | 0 | 0 | 0 | 1 | 1 |
| 0 | 0 | 0 | 0 | 1 | 1 |
| 0 | 0 | 0 | 0 | 1 | 1 |
| 0 | 0 | 0 | 0 | 1 | 1 |
| 0 | 1 | 1 | 1 | 1 | 5 |
| 0 | 1 | 1 | 1 | 1 | 5 |
| 0 | 1 | 1 | 1 | 1 | 5 |
| 0 | 1 | 1 | 1 | 1 | 5 |

| | | | | | |
|---|---|---|---|---|---|
| 0 | 1 | 1 | 1 | 1 | 5 |
| 0 | 1 | 1 | 1 | 1 | 5 |
| 0 | 1 | 1 | 1 | 1 | 5 |
| 0 | 1 | 1 | 1 | 1 | 5 |
| 0 | 1 | 1 | 1 | 1 | 5 |
| 0 | 1 | 1 | 1 | 1 | 5 |
| 0 | 1 | 1 | 1 | 1 | 5 |
| 0 | 1 | 1 | 1 | 1 | 5 |
| 0 | 1 | 1 | 1 | 1 | 5 |
| 0 | 1 | 1 | 1 | 1 | 5 |
| 0 | 1 | 1 | 1 | 1 | 5 |
| 0 | 1 | 1 | 1 | 1 | 5 |
| 0 | 1 | 1 | 1 | 1 | 5 |
| 0 | 1 | 1 | 1 | 1 | 5 |
| 0 | 1 | 1 | 1 | 1 | 5 |
| 0 | 1 | 1 | 1 | 1 | 5 |
| 0 | 1 | 1 | 1 | 0 | 4 |
| 0 | 1 | 1 | 1 | 0 | 4 |
| 0 | 1 | 1 | 1 | 0 | 4 |
| 0 | 1 | 1 | 1 | 0 | 4 |
| 0 | 1 | 1 | 1 | 0 | 4 |
| 0 | 1 | 1 | 1 | 0 | 4 |
| 0 | 1 | 1 | 1 | 0 | 4 |
| 0 | 1 | 1 | 1 | 0 | 4 |
| 0 | 1 | 1 | 1 | 0 | 4 |
| 0 | 1 | 1 | 1 | 0 | 4 |
| 0 | 1 | 1 | 1 | 0 | 4 |
| 0 | 1 | 1 | 1 | 0 | 4 |
| 0 | 1 | 1 | 1 | 0 | 4 |
| 0 | 1 | 1 | 1 | 0 | 4 |
| 0 | 1 | 1 | 1 | 0 | 4 |
| 0 | 1 | 1 | 1 | 0 | 4 |
| 0 | 1 | 1 | 1 | 0 | 4 |
| 0 | 1 | 1 | 1 | 0 | 4 |
| 0 | 1 | 1 | 1 | 0 | 4 |
| 0 | 1 | 1 | 1 | 0 | 4 |
| 0 | 1 | 1 | 1 | 1 | 5 |
| 0 | 1 | 1 | 1 | 1 | 5 |
| 0 | 1 | 1 | 1 | 1 | 5 |
| 0 | 1 | 1 | 1 | 1 | 5 |
| 0 | 1 | 1 | 1 | 1 | 5 |
| 0 | 1 | 1 | 1 | 1 | 5 |
| 0 | 1 | 1 | 1 | 1 | 5 |
| 0 | 1 | 1 | 1 | 1 | 5 |

| | | | | | |
|---|---|---|---|---|---|
| 0 | 1 | 1 | 1 | 1 | 5 |
| 0 | 1 | 1 | 1 | 1 | 5 |
| 0 | 1 | 1 | 1 | 1 | 5 |
| 0 | 1 | 1 | 1 | 1 | 5 |
| 0 | 1 | 1 | 1 | 1 | 5 |
| 0 | 1 | 1 | 1 | 1 | 5 |
| 0 | 1 | 1 | 1 | 1 | 5 |
| 0 | 1 | 1 | 1 | 1 | 5 |
| 0 | 1 | 1 | 1 | 1 | 5 |
| 0 | 1 | 1 | 1 | 1 | 5 |
| 0 | 1 | 1 | 1 | 1 | 5 |
| 0 | 1 | 1 | 1 | 1 | 5 |
| 0 | 1 | 1 | 1 | 1 | 5 |
| 0 | 1 | 1 | 1 | 1 | 5 |
| 0 | 1 | 1 | 1 | 1 | 5 |
| 0 | 1 | 1 | 1 | 1 | 5 |
| 0 | 1 | 1 | 1 | 1 | 5 |
| 0 | 1 | 1 | 1 | 1 | 5 |
| 0 | 1 | 1 | 1 | 1 | 5 |
| 0 | 1 | 1 | 1 | 1 | 5 |
| 0 | 1 | 1 | 1 | 1 | 5 |
| 0 | 1 | 1 | 1 | 1 | 5 |
| 0 | 1 | 1 | 1 | 1 | 5 |
| 0 | 1 | 1 | 1 | 1 | 5 |
| 0 | 1 | 1 | 1 | 1 | 5 |
| 0 | 1 | 1 | 1 | 1 | 5 |
| 0 | 1 | 1 | 1 | 1 | 5 |
| 0 | 1 | 1 | 1 | 1 | 5 |
| 0 | 1 | 1 | 1 | 1 | 5 |
| 0 | 1 | 1 | 1 | 1 | 5 |
| 0 | 1 | 1 | 1 | 1 | 5 |
| 0 | 1 | 1 | 1 | 1 | 5 |
| 0 | 0 | 0 | 0 | 1 | 1 |
| 0 | 0 | 0 | 0 | 1 | 1 |
| 0 | 0 | 0 | 0 | 1 | 1 |
| 0 | 0 | 0 | 0 | 1 | 1 |
| 0 | 0 | 0 | 0 | 1 | 1 |
| 0 | 0 | 0 | 0 | 1 | 1 |
| 0 | 0 | 0 | 0 | 1 | 1 |
| 0 | 0 | 0 | 0 | 1 | 1 |
| 0 | 0 | 0 | 0 | 1 | 1 |
| 0 | 0 | 0 | 0 | 1 | 1 |
| 0 | 0 | 0 | 0 | 1 | 1 |
| 0 | 0 | 0 | 0 | 1 | 1 |

| | | | | | |
|---|---|---|---|---|---|
| 0 | 0 | 0 | 0 | 1 | 1 |
| 0 | 0 | 0 | 0 | 1 | 1 |
| 0 | 0 | 0 | 0 | 1 | 1 |
| 0 | 0 | 0 | 0 | 1 | 1 |
| 0 | 0 | 0 | 0 | 1 | 1 |
| 0 | 0 | 0 | 0 | 1 | 1 |
| 0 | 0 | 0 | 0 | 1 | 1 |
| 0 | 0 | 0 | 0 | 1 | 1 |
| 0 | 0 | 0 | 0 | 1 | 1 |
| 0 | 0 | 0 | 0 | 1 | 1 |
| 0 | 0 | 0 | 0 | 1 | 1 |
| 0 | 0 | 0 | 0 | 1 | 1 |
| 0 | 0 | 0 | 0 | 1 | 1 |
| 0 | 0 | 0 | 0 | 1 | 1 |
| 0 | 0 | 0 | 0 | 1 | 1 |
| 0 | 0 | 0 | 0 | 1 | 1 |
| 0 | 0 | 0 | 0 | 1 | 1 |
| 0 | 0 | 0 | 0 | 1 | 1 |
| 0 | 0 | 0 | 0 | 1 | 1 |
| 0 | 0 | 0 | 0 | 1 | 1 |
| 0 | 0 | 0 | 0 | 1 | 1 |
| 0 | 0 | 0 | 0 | 1 | 1 |
| 0 | 0 | 0 | 0 | 1 | 1 |
| 0 | 0 | 0 | 0 | 1 | 1 |
| 0 | 0 | 0 | 0 | 1 | 1 |
| 0 | 0 | 0 | 0 | 1 | 1 |
| 0 | 0 | 0 | 0 | 1 | 1 |
| 0 | 0 | 0 | 0 | 1 | 1 |
| 0 | 0 | 0 | 0 | 1 | 1 |
| 0 | 0 | 0 | 0 | 1 | 1 |
| 0 | 0 | 0 | 0 | 1 | 1 |
| 0 | 0 | 0 | 0 | 1 | 1 |
| 0 | 0 | 0 | 0 | 1 | 1 |
| 0 | 0 | 0 | 0 | 1 | 1 |
| 0 | 0 | 0 | 0 | 1 | 1 |
| 0 | 0 | 0 | 0 | 1 | 1 |
| 0 | 0 | 0 | 0 | 1 | 1 |
| 0 | 0 | 0 | 0 | 1 | 1 |
| 0 | 0 | 0 | 0 | 1 | 1 |
| 0 | 0 | 0 | 0 | 1 | 1 |
| 0 | 0 | 0 | 0 | 1 | 1 |
| 0 | 0 | 0 | 0 | 1 | 1 |
| 0 | 0 | 0 | 0 | 1 | 1 |
| 0 | 0 | 0 | 0 | 1 | 1 |

| | | | | | |
|---|---|---|---|---|---|
| 0 | 0 | 0 | 0 | 1 | 1 |
| 0 | 0 | 0 | 0 | 1 | 1 |
| 0 | 0 | 0 | 0 | 1 | 1 |
| 0 | 0 | 0 | 0 | 1 | 1 |
| 0 | 1 | 1 | 1 | 0 | 4 |
| 0 | 1 | 1 | 1 | 0 | 4 |
| 0 | 1 | 1 | 1 | 0 | 4 |
| 0 | 1 | 1 | 1 | 0 | 4 |
| 0 | 1 | 1 | 1 | 0 | 4 |
| 0 | 1 | 1 | 1 | 0 | 4 |
| 0 | 1 | 1 | 1 | 0 | 4 |
| 0 | 1 | 1 | 1 | 0 | 4 |
| 0 | 1 | 1 | 1 | 0 | 4 |
| 0 | 1 | 1 | 1 | 0 | 4 |
| 0 | 1 | 1 | 1 | 0 | 4 |
| 0 | 1 | 1 | 1 | 0 | 4 |
| 0 | 1 | 1 | 1 | 0 | 4 |
| 0 | 1 | 1 | 1 | 0 | 4 |
| 0 | 1 | 1 | 1 | 0 | 4 |
| 0 | 1 | 1 | 1 | 0 | 4 |
| 0 | 1 | 1 | 1 | 0 | 4 |
| 0 | 1 | 1 | 1 | 0 | 4 |
| 0 | 1 | 1 | 1 | 0 | 4 |
| 0 | 1 | 1 | 1 | 0 | 4 |
| 0 | 1 | 1 | 1 | 1 | 5 |
| 0 | 1 | 1 | 1 | 1 | 5 |
| 0 | 1 | 1 | 1 | 1 | 5 |
| 0 | 1 | 1 | 1 | 1 | 5 |
| 0 | 1 | 1 | 1 | 1 | 5 |
| 0 | 1 | 1 | 1 | 1 | 5 |
| 0 | 1 | 1 | 1 | 1 | 5 |
| 0 | 1 | 1 | 1 | 1 | 5 |
| 0 | 1 | 1 | 1 | 1 | 5 |
| 0 | 1 | 1 | 1 | 1 | 5 |
| 0 | 1 | 1 | 1 | 1 | 5 |
| 0 | 1 | 1 | 1 | 1 | 5 |
| 0 | 1 | 1 | 1 | 1 | 5 |
| 0 | 1 | 1 | 1 | 1 | 5 |
| 0 | 1 | 1 | 1 | 1 | 5 |
| 0 | 1 | 1 | 1 | 1 | 5 |
| 0 | 1 | 1 | 1 | 1 | 5 |
| 0 | 1 | 1 | 1 | 1 | 5 |
| 0 | 1 | 1 | 1 | 1 | 5 |
| 0 | 1 | 1 | 1 | 1 | 5 |

| | | | | | |
|---|---|---|---|---|---|
| 1 | 1 | 1 | 1 | 1 | 8 |
| 1 | 1 | 1 | 1 | 1 | 8 |
| 1 | 1 | 1 | 1 | 1 | 8 |
| 1 | 1 | 1 | 1 | 1 | 8 |
| 1 | 1 | 1 | 1 | 1 | 8 |
| 1 | 1 | 1 | 1 | 1 | 8 |
| 1 | 1 | 1 | 1 | 1 | 8 |
| 1 | 1 | 1 | 1 | 1 | 8 |
| 1 | 1 | 1 | 1 | 1 | 8 |
| 1 | 1 | 1 | 1 | 1 | 8 |
| 0 | 1 | 1 | 1 | 0 | 4 |
| 0 | 1 | 1 | 1 | 0 | 4 |
| 0 | 1 | 1 | 1 | 0 | 4 |
| 0 | 1 | 1 | 1 | 0 | 4 |
| 0 | 1 | 1 | 1 | 0 | 4 |
| 0 | 1 | 1 | 1 | 0 | 4 |
| 0 | 1 | 1 | 1 | 0 | 4 |
| 0 | 1 | 1 | 1 | 0 | 4 |
| 0 | 1 | 1 | 1 | 0 | 4 |
| 0 | 1 | 1 | 1 | 0 | 4 |
| 0 | 1 | 1 | 1 | 0 | 4 |
| 0 | 1 | 1 | 1 | 0 | 4 |
| 0 | 1 | 1 | 1 | 0 | 4 |
| 0 | 1 | 1 | 1 | 0 | 4 |
| 0 | 1 | 1 | 1 | 0 | 4 |
| 0 | 1 | 1 | 1 | 0 | 4 |
| 0 | 1 | 1 | 1 | 0 | 4 |
| 0 | 1 | 1 | 1 | 0 | 4 |
| 0 | 1 | 1 | 1 | 0 | 4 |
| 0 | 1 | 1 | 1 | 0 | 4 |
| 0 | 1 | 1 | 1 | 0 | 4 |
| 0 | 1 | 1 | 1 | 0 | 4 |
| 0 | 1 | 1 | 1 | 0 | 4 |
| 0 | 1 | 1 | 1 | 0 | 4 |
| 0 | 1 | 1 | 1 | 0 | 4 |
| 0 | 1 | 1 | 1 | 0 | 4 |
| 0 | 1 | 1 | 1 | 0 | 4 |
| 0 | 1 | 1 | 1 | 0 | 4 |
| 0 | 1 | 1 | 1 | 0 | 4 |
| 0 | 1 | 1 | 1 | 0 | 4 |
| 0 | 1 | 1 | 1 | 0 | 4 |
| 0 | 1 | 1 | 1 | 0 | 4 |
| 0 | 1 | 1 | 1 | 0 | 4 |
| 0 | 1 | 1 | 1 | 0 | 4 |

| | | | | | |
|---|---|---|---|---|---|
| 0 | 1 | 1 | 1 | 0 | 4 |
| 0 | 1 | 1 | 1 | 0 | 4 |
| 0 | 1 | 1 | 1 | 0 | 4 |
| 0 | 1 | 1 | 1 | 0 | 4 |
| 0 | 1 | 1 | 1 | 0 | 4 |
| 0 | 1 | 1 | 1 | 0 | 4 |
| 0 | 1 | 1 | 1 | 1 | 5 |
| 0 | 1 | 1 | 1 | 1 | 5 |
| 0 | 1 | 1 | 1 | 1 | 5 |
| 0 | 1 | 1 | 1 | 1 | 5 |
| 0 | 1 | 1 | 1 | 1 | 5 |
| 0 | 1 | 1 | 1 | 1 | 5 |
| 0 | 1 | 1 | 1 | 1 | 5 |
| 0 | 1 | 1 | 1 | 1 | 5 |
| 0 | 1 | 1 | 1 | 1 | 5 |
| 0 | 1 | 1 | 1 | 1 | 5 |
| 0 | 1 | 1 | 1 | 1 | 5 |
| 0 | 1 | 1 | 1 | 1 | 5 |
| 0 | 1 | 1 | 1 | 1 | 5 |
| 0 | 1 | 1 | 1 | 1 | 5 |
| 0 | 1 | 1 | 1 | 1 | 5 |
| 0 | 1 | 1 | 1 | 1 | 5 |
| 0 | 1 | 1 | 1 | 1 | 5 |
| 0 | 1 | 1 | 1 | 1 | 5 |
| 0 | 1 | 1 | 1 | 1 | 5 |
| 0 | 1 | 1 | 1 | 1 | 5 |
| 1 | 0 | 0 | 0 | 0 | 3 |
| 1 | 0 | 0 | 0 | 0 | 3 |
| 1 | 0 | 0 | 0 | 0 | 3 |
| 1 | 0 | 0 | 0 | 0 | 3 |
| 1 | 0 | 0 | 0 | 0 | 3 |
| 1 | 0 | 0 | 0 | 0 | 3 |
| 1 | 0 | 0 | 0 | 0 | 3 |
| 1 | 0 | 0 | 0 | 0 | 3 |
| 1 | 0 | 0 | 0 | 0 | 3 |
| 1 | 0 | 0 | 0 | 0 | 3 |
| 1 | 0 | 0 | 0 | 0 | 3 |
| 1 | 0 | 0 | 0 | 0 | 3 |
| 1 | 0 | 0 | 0 | 0 | 3 |
| 1 | 0 | 0 | 0 | 0 | 3 |
| 1 | 0 | 0 | 0 | 0 | 3 |
| 1 | 0 | 0 | 0 | 0 | 3 |
| 1 | 0 | 0 | 0 | 0 | 3 |
| 1 | 0 | 0 | 0 | 0 | 3 |

| | | | | | |
|---|---|---|---|---|---|
| 1 | 0 | 0 | 0 | 0 | 3 |
| 1 | 0 | 0 | 0 | 0 | 3 |
| 1 | 0 | 0 | 0 | 0 | 3 |
| 1 | 0 | 0 | 0 | 0 | 3 |
| 1 | 0 | 0 | 0 | 0 | 3 |
| 1 | 0 | 0 | 0 | 0 | 3 |
| 1 | 0 | 0 | 0 | 0 | 3 |
| 1 | 0 | 0 | 0 | 0 | 3 |
| 1 | 0 | 0 | 0 | 0 | 3 |
| 1 | 0 | 0 | 0 | 0 | 3 |
| 1 | 0 | 0 | 0 | 0 | 3 |
| 1 | 0 | 0 | 0 | 0 | 3 |
| 1 | 0 | 0 | 0 | 0 | 3 |
| 1 | 0 | 0 | 0 | 0 | 3 |
| 1 | 0 | 0 | 0 | 0 | 3 |
| 1 | 0 | 0 | 0 | 0 | 3 |
| 1 | 0 | 0 | 0 | 0 | 3 |
| 1 | 0 | 0 | 0 | 0 | 3 |
| 1 | 0 | 0 | 0 | 0 | 3 |
| 1 | 0 | 0 | 0 | 0 | 3 |
| 1 | 0 | 0 | 0 | 0 | 3 |
| 1 | 0 | 0 | 0 | 0 | 3 |
| 1 | 0 | 0 | 0 | 0 | 3 |
| 1 | 0 | 0 | 0 | 0 | 3 |
| 1 | 0 | 0 | 0 | 0 | 3 |
| 1 | 0 | 0 | 0 | 0 | 3 |
| 1 | 0 | 0 | 0 | 0 | 3 |
| 1 | 0 | 0 | 0 | 0 | 3 |
| 1 | 0 | 0 | 0 | 0 | 3 |
| 1 | 0 | 0 | 0 | 0 | 3 |
| 1 | 0 | 0 | 0 | 0 | 3 |
| 1 | 0 | 0 | 0 | 0 | 3 |
| 1 | 0 | 0 | 0 | 0 | 3 |
| 1 | 0 | 0 | 0 | 0 | 3 |
| 1 | 0 | 0 | 0 | 0 | 3 |
| 1 | 0 | 0 | 0 | 0 | 3 |
| 1 | 0 | 0 | 0 | 0 | 3 |
| 1 | 0 | 0 | 0 | 0 | 3 |
| 1 | 0 | 0 | 0 | 0 | 3 |
| 1 | 0 | 0 | 0 | 0 | 3 |
| 1 | 0 | 0 | 0 | 0 | 3 |
| 1 | 0 | 0 | 0 | 0 | 3 |
| 1 | 0 | 0 | 0 | 0 | 3 |
| 1 | 0 | 0 | 0 | 0 | 3 |

| | | | | | |
|---|---|---|---|---|---|
| 1 | 0 | 0 | 0 | 0 | 3 |
| 1 | 0 | 0 | 0 | 0 | 3 |
| 1 | 0 | 0 | 0 | 0 | 3 |
| 1 | 0 | 0 | 0 | 0 | 3 |
| 1 | 0 | 0 | 0 | 0 | 3 |
| 1 | 0 | 0 | 0 | 0 | 3 |
| 1 | 0 | 0 | 0 | 0 | 3 |
| 1 | 0 | 0 | 0 | 0 | 3 |
| 1 | 0 | 0 | 0 | 0 | 3 |
| 1 | 0 | 0 | 0 | 0 | 3 |
| 1 | 0 | 0 | 0 | 0 | 3 |
| 1 | 0 | 0 | 0 | 0 | 3 |
| 1 | 0 | 0 | 0 | 0 | 3 |
| 1 | 0 | 0 | 0 | 0 | 3 |
| 1 | 0 | 0 | 0 | 0 | 3 |
| 1 | 0 | 0 | 0 | 0 | 3 |
| 1 | 0 | 0 | 0 | 0 | 3 |
| 1 | 0 | 0 | 0 | 0 | 3 |
| 1 | 0 | 0 | 0 | 0 | 3 |
| 1 | 0 | 0 | 0 | 0 | 3 |
| 1 | 0 | 0 | 0 | 0 | 3 |
| 1 | 0 | 0 | 0 | 0 | 3 |
| 1 | 0 | 0 | 0 | 0 | 3 |
| 1 | 0 | 0 | 0 | 0 | 3 |
| 1 | 0 | 0 | 0 | 0 | 3 |
| 1 | 0 | 0 | 0 | 0 | 3 |
| 1 | 0 | 0 | 0 | 0 | 3 |
| 1 | 0 | 0 | 0 | 0 | 3 |
| 1 | 0 | 0 | 0 | 0 | 3 |
| 1 | 0 | 0 | 0 | 0 | 3 |
| 1 | 0 | 0 | 0 | 0 | 3 |
| 1 | 0 | 0 | 0 | 0 | 3 |
| 1 | 0 | 0 | 0 | 0 | 3 |
| 1 | 0 | 0 | 0 | 0 | 3 |
| 1 | 0 | 0 | 0 | 0 | 3 |
| 1 | 0 | 0 | 0 | 0 | 3 |
| 1 | 0 | 0 | 0 | 0 | 3 |
| 1 | 0 | 0 | 0 | 0 | 3 |
| 1 | 0 | 0 | 0 | 0 | 3 |
| 1 | 0 | 0 | 0 | 0 | 3 |
| 1 | 0 | 0 | 0 | 0 | 3 |
| 1 | 0 | 0 | 0 | 0 | 3 |
| 1 | 0 | 0 | 0 | 0 | 3 |
| 1 | 0 | 0 | 0 | 0 | 3 |

| | | | | | |
|---|---|---|---|---|---|
| 1 | 0 | 0 | 0 | 0 | 3 |
| 1 | 0 | 0 | 0 | 0 | 3 |
| 1 | 0 | 0 | 0 | 0 | 3 |
| 1 | 0 | 0 | 0 | 0 | 3 |
| 1 | 0 | 0 | 0 | 0 | 3 |
| 1 | 0 | 0 | 0 | 0 | 3 |
| 1 | 1 | 1 | 1 | 1 | 8 |
| 1 | 1 | 1 | 1 | 1 | 8 |
| 1 | 1 | 1 | 1 | 1 | 8 |
| 1 | 1 | 1 | 1 | 1 | 8 |
| 1 | 1 | 1 | 1 | 1 | 8 |
| 1 | 1 | 1 | 1 | 1 | 8 |
| 1 | 1 | 1 | 1 | 1 | 8 |
| 1 | 1 | 1 | 1 | 1 | 8 |
| 1 | 1 | 1 | 1 | 1 | 8 |
| 1 | 1 | 1 | 1 | 1 | 8 |
| 1 | 1 | 1 | 1 | 1 | 8 |
| 1 | 1 | 1 | 1 | 1 | 8 |
| 1 | 1 | 1 | 1 | 1 | 8 |
| 1 | 1 | 1 | 1 | 1 | 8 |
| 1 | 1 | 1 | 1 | 1 | 8 |
| 1 | 1 | 1 | 1 | 1 | 8 |
| 1 | 1 | 1 | 1 | 1 | 8 |
| 1 | 1 | 1 | 1 | 1 | 8 |
| 1 | 1 | 1 | 1 | 1 | 8 |
| 1 | 1 | 1 | 1 | 1 | 8 |
| 1 | 1 | 1 | 1 | 1 | 8 |
| 1 | 1 | 1 | 1 | 1 | 8 |
| 1 | 1 | 1 | 1 | 1 | 8 |
| 1 | 1 | 1 | 1 | 1 | 8 |
| 1 | 1 | 1 | 1 | 1 | 8 |
| 1 | 1 | 1 | 1 | 1 | 8 |
| 1 | 1 | 1 | 1 | 1 | 8 |
| 1 | 1 | 1 | 1 | 1 | 8 |
| 1 | 1 | 1 | 1 | 1 | 8 |
| 1 | 1 | 1 | 1 | 1 | 8 |
| 1 | 1 | 1 | 1 | 1 | 8 |
| 1 | 1 | 1 | 1 | 1 | 8 |
| 1 | 1 | 1 | 1 | 1 | 8 |
| 1 | 1 | 1 | 1 | 1 | 8 |
| 1 | 1 | 1 | 1 | 1 | 8 |
| 1 | 1 | 1 | 1 | 1 | 8 |
| 1 | 1 | 1 | 1 | 1 | 8 |
| 1 | 1 | 1 | 1 | 1 | 8 |

| | | | | | |
|---|---|---|---|---|---|
| 1 | 1 | 1 | 1 | 1 | 8 |
| 1 | 1 | 1 | 1 | 1 | 8 |
| 0 | 1 | 0 | 0 | 0 | 2 |
| 0 | 1 | 0 | 0 | 0 | 2 |
| 0 | 1 | 0 | 0 | 0 | 2 |
| 0 | 1 | 0 | 0 | 0 | 2 |
| 0 | 1 | 0 | 0 | 0 | 2 |
| 0 | 1 | 0 | 0 | 0 | 2 |
| 0 | 1 | 0 | 0 | 0 | 2 |
| 0 | 1 | 0 | 0 | 0 | 2 |
| 1 | 1 | 1 | 1 | 1 | 8 |
| 1 | 1 | 1 | 1 | 1 | 8 |
| 1 | 1 | 1 | 1 | 1 | 8 |
| 1 | 1 | 1 | 1 | 1 | 8 |
| 1 | 1 | 1 | 1 | 1 | 8 |
| 1 | 1 | 1 | 1 | 1 | 8 |
| 1 | 1 | 1 | 1 | 1 | 8 |
| 1 | 1 | 1 | 1 | 1 | 8 |
| 1 | 1 | 1 | 1 | 1 | 8 |
| 1 | 1 | 1 | 1 | 1 | 8 |
| 1 | 1 | 1 | 1 | 1 | 8 |
| 1 | 1 | 1 | 1 | 1 | 8 |
| 1 | 1 | 1 | 1 | 1 | 8 |
| 1 | 1 | 1 | 1 | 1 | 8 |
| 1 | 1 | 1 | 1 | 1 | 8 |
| 1 | 1 | 1 | 1 | 1 | 8 |
| 1 | 1 | 1 | 1 | 1 | 8 |
| 1 | 1 | 1 | 1 | 1 | 8 |
| 1 | 1 | 1 | 1 | 1 | 8 |
| 1 | 1 | 1 | 1 | 1 | 8 |
| 1 | 1 | 1 | 1 | 1 | 8 |
| 1 | 1 | 1 | 1 | 1 | 8 |
| 1 | 1 | 1 | 1 | 1 | 8 |
| 1 | 1 | 1 | 1 | 1 | 8 |
| 1 | 1 | 1 | 1 | 1 | 8 |
| 1 | 1 | 1 | 1 | 1 | 8 |
| 1 | 1 | 1 | 1 | 1 | 8 |
| 1 | 1 | 1 | 1 | 1 | 8 |
| 1 | 1 | 1 | 1 | 1 | 8 |
| 1 | 1 | 1 | 1 | 1 | 8 |
| 1 | 1 | 1 | 1 | 1 | 8 |
| 1 | 1 | 1 | 1 | 1 | 8 |
| 1 | 1 | 1 | 1 | 1 | 8 |
| 1 | 1 | 1 | 1 | 1 | 8 |

| | | | | | |
|---|---|---|---|---|---|
| 1 | 1 | 1 | 1 | 1 | 8 |
| 1 | 1 | 1 | 1 | 1 | 8 |
| 1 | 1 | 1 | 1 | 1 | 8 |
| 1 | 1 | 1 | 1 | 1 | 8 |
| 1 | 1 | 1 | 1 | 1 | 8 |
| 1 | 1 | 1 | 1 | 1 | 8 |
| 1 | 1 | 1 | 1 | 1 | 8 |
| 1 | 1 | 1 | 1 | 1 | 8 |
| 1 | 1 | 1 | 1 | 1 | 8 |
| 1 | 1 | 1 | 1 | 1 | 8 |
| 1 | 1 | 1 | 1 | 1 | 8 |
| 1 | 1 | 1 | 1 | 1 | 8 |
| 1 | 1 | 1 | 1 | 1 | 8 |
| 1 | 1 | 1 | 1 | 1 | 8 |
| 1 | 1 | 1 | 1 | 1 | 8 |
| 1 | 1 | 1 | 1 | 1 | 8 |
| 1 | 1 | 1 | 1 | 1 | 8 |
| 1 | 1 | 1 | 1 | 1 | 8 |
| 1 | 1 | 1 | 1 | 1 | 8 |
| 1 | 1 | 1 | 1 | 1 | 8 |
| 1 | 1 | 1 | 1 | 1 | 8 |
| 1 | 1 | 1 | 1 | 1 | 8 |
| 1 | 1 | 1 | 1 | 1 | 8 |
| 1 | 1 | 1 | 1 | 1 | 8 |
| 1 | 1 | 1 | 1 | 1 | 8 |
| 1 | 1 | 1 | 1 | 1 | 8 |
| 1 | 1 | 1 | 1 | 1 | 8 |
| 1 | 1 | 1 | 1 | 1 | 8 |
| 1 | 1 | 1 | 1 | 1 | 8 |
| 1 | 1 | 1 | 1 | 1 | 8 |
| 0 | 1 | 0 | 0 | 0 | 2 |
| 0 | 1 | 0 | 0 | 0 | 2 |
| 0 | 1 | 0 | 0 | 0 | 2 |
| 0 | 1 | 0 | 0 | 0 | 2 |
| 0 | 1 | 0 | 0 | 0 | 2 |
| 0 | 1 | 0 | 0 | 0 | 2 |
| 0 | 1 | 0 | 0 | 0 | 2 |
| 0 | 1 | 0 | 0 | 0 | 2 |
| 0 | 1 | 0 | 0 | 0 | 2 |
| 0 | 1 | 0 | 0 | 0 | 2 |
| 0 | 1 | 0 | 0 | 0 | 2 |
| 0 | 1 | 0 | 0 | 0 | 2 |
| 0 | 1 | 0 | 0 | 0 | 2 |
| 0 | 1 | 0 | 0 | 0 | 2 |

| | | | | | |
|---|---|---|---|---|---|
| 0 | 1 | 0 | 0 | 0 | 2 |
| 0 | 1 | 0 | 0 | 0 | 2 |
| 0 | 1 | 0 | 0 | 0 | 2 |
| 0 | 1 | 0 | 0 | 0 | 2 |
| 0 | 1 | 0 | 0 | 0 | 2 |
| 0 | 1 | 0 | 0 | 0 | 2 |
| 0 | 1 | 0 | 0 | 0 | 2 |
| 0 | 1 | 0 | 0 | 0 | 2 |
| 0 | 1 | 0 | 0 | 0 | 2 |
| 0 | 1 | 0 | 0 | 0 | 2 |
| 0 | 1 | 0 | 0 | 0 | 2 |
| 0 | 1 | 0 | 0 | 0 | 2 |
| 0 | 1 | 0 | 0 | 0 | 2 |
| 0 | 1 | 0 | 0 | 0 | 2 |
| 0 | 1 | 0 | 0 | 0 | 2 |
| 0 | 1 | 0 | 0 | 0 | 2 |
| 0 | 1 | 0 | 0 | 0 | 2 |
| 0 | 1 | 0 | 0 | 0 | 2 |
| 1 | 1 | 1 | 1 | 1 | 8 |
| 1 | 1 | 1 | 1 | 1 | 8 |
| 1 | 1 | 1 | 1 | 1 | 8 |
| 1 | 1 | 1 | 1 | 1 | 8 |
| 1 | 1 | 1 | 1 | 1 | 8 |
| 1 | 1 | 1 | 1 | 1 | 8 |
| 1 | 1 | 1 | 1 | 1 | 8 |
| 1 | 1 | 1 | 1 | 1 | 8 |
| 1 | 1 | 1 | 1 | 1 | 8 |
| 1 | 1 | 1 | 1 | 1 | 8 |
| 1 | 1 | 1 | 1 | 1 | 8 |
| 1 | 1 | 1 | 1 | 1 | 8 |
| 1 | 1 | 1 | 1 | 1 | 8 |
| 1 | 1 | 1 | 1 | 1 | 8 |
| 1 | 1 | 1 | 1 | 1 | 8 |
| 1 | 1 | 1 | 1 | 1 | 8 |
| 1 | 1 | 1 | 1 | 1 | 8 |
| 1 | 1 | 1 | 1 | 1 | 8 |
| 1 | 1 | 1 | 1 | 1 | 8 |
| 1 | 1 | 1 | 1 | 1 | 8 |
| 1 | 1 | 1 | 1 | 1 | 8 |
| 1 | 1 | 1 | 1 | 1 | 8 |
| 1 | 1 | 1 | 1 | 1 | 8 |
| 1 | 1 | 1 | 1 | 1 | 8 |
| 1 | 1 | 1 | 1 | 1 | 8 |
| 1 | 1 | 1 | 1 | 1 | 8 |

| | | | | | |
|---|---|---|---|---|---|
| 1 | 1 | 1 | 1 | 1 | 8 |
| 1 | 1 | 1 | 1 | 1 | 8 |
| 1 | 1 | 1 | 1 | 1 | 8 |
| 1 | 1 | 1 | 1 | 1 | 8 |
| 1 | 1 | 1 | 1 | 1 | 8 |
| 1 | 1 | 1 | 1 | 1 | 8 |
| 1 | 1 | 1 | 1 | 1 | 8 |
| 1 | 1 | 1 | 1 | 1 | 8 |
| 1 | 1 | 1 | 1 | 1 | 8 |
| 1 | 1 | 1 | 1 | 1 | 8 |
| 1 | 1 | 1 | 1 | 1 | 8 |
| 1 | 1 | 1 | 1 | 1 | 8 |
| 1 | 1 | 1 | 1 | 1 | 8 |
| 1 | 1 | 1 | 1 | 1 | 8 |
| 0 | 1 | 0 | 0 | 0 | 2 |
| 0 | 1 | 0 | 0 | 0 | 2 |
| 0 | 1 | 0 | 0 | 0 | 2 |
| 0 | 1 | 0 | 0 | 0 | 2 |
| 0 | 1 | 0 | 0 | 0 | 2 |
| 0 | 1 | 0 | 0 | 0 | 2 |
| 0 | 1 | 0 | 0 | 0 | 2 |
| 0 | 1 | 0 | 0 | 0 | 2 |
| 0 | 1 | 0 | 0 | 0 | 2 |
| 0 | 1 | 0 | 0 | 0 | 2 |
| 0 | 1 | 0 | 0 | 0 | 2 |
| 0 | 1 | 0 | 0 | 0 | 2 |
| 0 | 1 | 0 | 0 | 0 | 2 |
| 0 | 1 | 0 | 0 | 0 | 2 |
| 0 | 1 | 0 | 0 | 0 | 2 |
| 0 | 1 | 0 | 0 | 0 | 2 |
| 0 | 1 | 0 | 0 | 0 | 2 |
| 0 | 1 | 0 | 0 | 0 | 2 |
| 0 | 1 | 0 | 0 | 0 | 2 |
| 0 | 1 | 0 | 0 | 0 | 2 |
| 0 | 1 | 0 | 0 | 0 | 2 |
| 0 | 1 | 0 | 0 | 0 | 2 |
| 0 | 1 | 0 | 0 | 0 | 2 |
| 0 | 1 | 0 | 0 | 0 | 2 |
| 0 | 1 | 0 | 0 | 0 | 2 |
| 0 | 1 | 0 | 0 | 0 | 2 |
| 0 | 1 | 0 | 0 | 0 | 2 |
| 0 | 1 | 0 | 0 | 0 | 2 |
| 0 | 1 | 0 | 0 | 0 | 2 |
| 0 | 1 | 0 | 0 | 0 | 2 |

| | | | | | |
|---|---|---|---|---|---|
| 0 | 1 | 0 | 0 | 0 | 2 |
| 0 | 1 | 0 | 0 | 0 | 2 |
| 0 | 1 | 0 | 0 | 0 | 2 |
| 0 | 1 | 0 | 0 | 0 | 2 |
| 0 | 1 | 0 | 0 | 0 | 2 |
| 0 | 1 | 0 | 0 | 0 | 2 |
| 0 | 1 | 0 | 0 | 0 | 2 |
| 0 | 1 | 0 | 0 | 0 | 2 |
| 0 | 1 | 0 | 0 | 0 | 2 |
| 0 | 1 | 0 | 0 | 0 | 2 |
| 1 | 1 | 1 | 1 | 1 | 8 |
| 1 | 1 | 1 | 1 | 1 | 8 |
| 1 | 1 | 1 | 1 | 1 | 8 |
| 1 | 1 | 1 | 1 | 1 | 8 |
| 1 | 1 | 1 | 1 | 1 | 8 |
| 1 | 1 | 1 | 1 | 1 | 8 |
| 1 | 1 | 1 | 1 | 1 | 8 |
| 1 | 1 | 1 | 1 | 1 | 8 |
| 1 | 1 | 1 | 1 | 1 | 8 |
| 1 | 1 | 1 | 1 | 1 | 8 |
| 1 | 1 | 1 | 1 | 1 | 8 |
| 1 | 1 | 1 | 1 | 1 | 8 |
| 1 | 1 | 1 | 1 | 1 | 8 |
| 1 | 1 | 1 | 1 | 1 | 8 |
| 1 | 1 | 1 | 1 | 1 | 8 |
| 1 | 1 | 1 | 1 | 1 | 8 |
| 1 | 1 | 1 | 1 | 1 | 8 |
| 1 | 1 | 1 | 1 | 1 | 8 |
| 1 | 1 | 1 | 1 | 1 | 8 |
| 1 | 1 | 1 | 1 | 1 | 8 |
| 1 | 1 | 1 | 1 | 1 | 8 |
| 1 | 1 | 1 | 1 | 1 | 8 |
| 1 | 1 | 1 | 1 | 1 | 8 |
| 1 | 1 | 1 | 1 | 1 | 8 |
| 1 | 1 | 1 | 1 | 1 | 8 |
| 1 | 1 | 1 | 1 | 1 | 8 |
| 1 | 1 | 1 | 1 | 1 | 8 |
| 1 | 1 | 1 | 1 | 1 | 8 |
| 1 | 1 | 1 | 1 | 1 | 8 |
| 1 | 1 | 1 | 1 | 1 | 8 |
| 1 | 1 | 1 | 1 | 1 | 8 |
| 1 | 1 | 1 | 1 | 1 | 8 |
| 1 | 1 | 1 | 1 | 1 | 8 |
| 1 | 1 | 1 | 1 | 1 | 8 |

| | | | | | |
|---|---|---|---|---|---|
| 1 | 1 | 1 | 1 | 1 | 8 |
| 1 | 1 | 1 | 1 | 1 | 8 |
| 1 | 1 | 1 | 1 | 1 | 8 |
| 1 | 1 | 1 | 1 | 1 | 8 |
| 1 | 1 | 1 | 1 | 1 | 8 |
| 1 | 1 | 1 | 1 | 1 | 8 |
| 1 | 1 | 1 | 1 | 1 | 8 |
| 1 | 1 | 1 | 1 | 1 | 8 |
| 0 | 0 | 0 | 0 | 1 | 1 |
| 0 | 0 | 0 | 0 | 1 | 1 |
| 0 | 0 | 0 | 0 | 1 | 1 |
| 0 | 0 | 0 | 0 | 1 | 1 |
| 0 | 0 | 0 | 0 | 1 | 1 |
| 0 | 0 | 0 | 0 | 1 | 1 |
| 0 | 0 | 0 | 0 | 1 | 1 |
| 0 | 0 | 0 | 0 | 1 | 1 |
| 0 | 0 | 0 | 0 | 1 | 1 |
| 0 | 0 | 0 | 0 | 1 | 1 |
| 0 | 0 | 0 | 0 | 1 | 1 |
| 0 | 0 | 0 | 0 | 1 | 1 |
| 0 | 0 | 0 | 0 | 1 | 1 |
| 0 | 0 | 0 | 0 | 1 | 1 |
| 0 | 0 | 0 | 0 | 1 | 1 |
| 0 | 0 | 0 | 0 | 1 | 1 |
| 0 | 0 | 0 | 0 | 1 | 1 |
| 0 | 0 | 0 | 0 | 1 | 1 |
| 0 | 0 | 0 | 0 | 1 | 1 |
| 0 | 0 | 0 | 0 | 1 | 1 |
| 0 | 0 | 0 | 0 | 1 | 1 |
| 0 | 0 | 0 | 0 | 1 | 1 |
| 0 | 0 | 0 | 0 | 1 | 1 |
| 0 | 0 | 0 | 0 | 1 | 1 |
| 0 | 0 | 0 | 0 | 1 | 1 |
| 0 | 0 | 0 | 0 | 1 | 1 |
| 0 | 0 | 0 | 0 | 1 | 1 |
| 0 | 0 | 0 | 0 | 1 | 1 |
| 0 | 0 | 0 | 0 | 1 | 1 |
| 0 | 0 | 0 | 0 | 1 | 1 |
| 0 | 0 | 0 | 0 | 1 | 1 |
| 0 | 0 | 0 | 0 | 1 | 1 |
| 0 | 0 | 0 | 0 | 1 | 1 |
| 0 | 0 | 0 | 0 | 1 | 1 |
| 0 | 0 | 0 | 0 | 1 | 1 |
| 0 | 0 | 0 | 0 | 1 | 1 |

| | | | | | |
|---|---|---|---|---|---|
| 0 | 0 | 0 | 0 | 1 | 1 |
| 0 | 0 | 0 | 0 | 1 | 1 |
| 0 | 0 | 0 | 0 | 1 | 1 |
| 0 | 0 | 0 | 0 | 1 | 1 |
| 0 | 0 | 0 | 0 | 1 | 1 |
| 0 | 0 | 0 | 0 | 1 | 1 |
| 0 | 0 | 0 | 0 | 1 | 1 |
| 0 | 0 | 0 | 0 | 1 | 1 |
| 0 | 0 | 0 | 0 | 1 | 1 |
| 0 | 0 | 0 | 0 | 1 | 1 |
| 0 | 0 | 0 | 0 | 1 | 1 |
| 0 | 0 | 0 | 0 | 1 | 1 |
| 0 | 0 | 0 | 0 | 1 | 1 |
| 0 | 0 | 0 | 0 | 1 | 1 |
| 0 | 0 | 0 | 0 | 1 | 1 |
| 0 | 0 | 0 | 0 | 1 | 1 |
| 0 | 0 | 0 | 0 | 1 | 1 |
| 0 | 0 | 0 | 0 | 1 | 1 |
| 0 | 0 | 0 | 0 | 1 | 1 |
| 0 | 0 | 0 | 0 | 1 | 1 |
| 0 | 0 | 0 | 0 | 1 | 1 |
| 0 | 0 | 0 | 0 | 1 | 1 |
| 0 | 0 | 0 | 0 | 1 | 1 |
| 0 | 0 | 0 | 0 | 1 | 1 |
| 0 | 0 | 0 | 0 | 1 | 1 |
| 0 | 0 | 0 | 0 | 1 | 1 |
| 0 | 0 | 0 | 0 | 1 | 1 |
| 0 | 0 | 0 | 0 | 1 | 1 |
| 0 | 0 | 0 | 0 | 1 | 1 |
| 0 | 0 | 0 | 0 | 1 | 1 |
| 0 | 0 | 0 | 0 | 1 | 1 |
| 0 | 0 | 0 | 0 | 1 | 1 |
| 0 | 0 | 0 | 0 | 1 | 1 |
| 0 | 0 | 0 | 0 | 1 | 1 |
| 0 | 0 | 0 | 0 | 1 | 1 |
| 0 | 0 | 0 | 0 | 1 | 1 |
| 0 | 0 | 0 | 0 | 1 | 1 |
| 0 | 0 | 0 | 0 | 1 | 1 |
| 0 | 0 | 0 | 0 | 1 | 1 |
| 0 | 0 | 0 | 0 | 1 | 1 |
| 0 | 0 | 0 | 0 | 1 | 1 |
| 0 | 0 | 0 | 0 | 1 | 1 |
| 0 | 0 | 0 | 0 | 1 | 1 |
| 0 | 0 | 0 | 0 | 1 | 1 |

| | | | | | |
|---|---|---|---|---|---|
| 0 | 0 | 0 | 0 | 1 | 1 |
| 0 | 0 | 0 | 0 | 1 | 1 |
| 0 | 0 | 0 | 0 | 1 | 1 |
| 0 | 0 | 0 | 0 | 1 | 1 |
| 0 | 0 | 0 | 0 | 1 | 1 |
| 0 | 0 | 0 | 0 | 1 | 1 |
| 0 | 0 | 0 | 0 | 1 | 1 |
| 0 | 0 | 0 | 0 | 1 | 1 |
| 0 | 0 | 0 | 0 | 1 | 1 |
| 0 | 0 | 0 | 0 | 1 | 1 |
| 0 | 0 | 0 | 0 | 1 | 1 |
| 0 | 0 | 0 | 0 | 1 | 1 |
| 0 | 0 | 0 | 0 | 1 | 1 |
| 0 | 0 | 0 | 0 | 1 | 1 |
| 0 | 0 | 0 | 0 | 1 | 1 |
| 0 | 0 | 0 | 0 | 1 | 1 |
| 0 | 0 | 0 | 0 | 1 | 1 |
| 0 | 0 | 0 | 0 | 1 | 1 |
| 0 | 0 | 0 | 0 | 1 | 1 |
| 0 | 0 | 0 | 0 | 1 | 1 |
| 0 | 0 | 0 | 0 | 1 | 1 |
| 0 | 0 | 0 | 0 | 1 | 1 |
| 0 | 0 | 0 | 0 | 1 | 1 |
| 0 | 0 | 0 | 0 | 1 | 1 |
| 0 | 0 | 0 | 0 | 1 | 1 |
| 0 | 0 | 0 | 0 | 1 | 1 |
| 0 | 0 | 0 | 0 | 1 | 1 |
| 0 | 0 | 0 | 0 | 1 | 1 |
| 0 | 0 | 0 | 0 | 1 | 1 |
| 0 | 0 | 0 | 0 | 1 | 1 |
| 0 | 0 | 0 | 0 | 1 | 1 |
| 0 | 0 | 0 | 0 | 1 | 1 |
| 1 | 1 | 1 | 1 | 1 | 8 |
| 1 | 1 | 1 | 1 | 1 | 8 |
| 1 | 1 | 1 | 1 | 1 | 8 |
| 1 | 1 | 1 | 1 | 1 | 8 |
| 1 | 1 | 1 | 1 | 1 | 8 |
| 1 | 1 | 1 | 1 | 1 | 8 |
| 1 | 1 | 1 | 1 | 1 | 8 |
| 1 | 1 | 1 | 1 | 1 | 8 |
| 1 | 1 | 1 | 1 | 1 | 8 |
| 1 | 1 | 1 | 1 | 1 | 8 |
| 0 | 1 | 0 | 0 | 0 | 2 |
| 0 | 1 | 0 | 0 | 0 | 2 |

| | | | | | |
|---|---|---|---|---|---|
| 0 | 1 | 0 | 0 | 0 | 2 |
| 0 | 1 | 0 | 0 | 0 | 2 |
| 0 | 1 | 0 | 0 | 0 | 2 |
| 0 | 1 | 0 | 0 | 0 | 2 |
| 0 | 1 | 0 | 0 | 0 | 2 |
| 0 | 1 | 0 | 0 | 0 | 2 |
| 0 | 1 | 0 | 0 | 0 | 2 |
| 0 | 1 | 0 | 0 | 0 | 2 |
| 1 | 1 | 1 | 1 | 1 | 8 |
| 1 | 1 | 1 | 1 | 1 | 8 |
| 1 | 1 | 1 | 1 | 1 | 8 |
| 1 | 1 | 1 | 1 | 1 | 8 |
| 1 | 1 | 1 | 1 | 1 | 8 |
| 1 | 1 | 1 | 1 | 1 | 8 |
| 1 | 1 | 1 | 1 | 1 | 8 |
| 1 | 1 | 1 | 1 | 1 | 8 |
| 1 | 1 | 1 | 1 | 1 | 8 |
| 1 | 1 | 1 | 1 | 1 | 8 |
| 1 | 1 | 1 | 1 | 1 | 8 |
| 1 | 1 | 1 | 1 | 1 | 8 |
| 1 | 1 | 1 | 1 | 1 | 8 |
| 1 | 1 | 1 | 1 | 1 | 8 |
| 1 | 1 | 1 | 1 | 1 | 8 |
| 1 | 1 | 1 | 1 | 1 | 8 |
| 1 | 1 | 1 | 1 | 1 | 8 |
| 1 | 1 | 1 | 1 | 1 | 8 |
| 1 | 1 | 1 | 1 | 1 | 8 |
| 1 | 1 | 1 | 1 | 1 | 8 |
| 1 | 1 | 1 | 1 | 1 | 8 |
| 1 | 1 | 1 | 1 | 1 | 8 |
| 1 | 1 | 1 | 1 | 1 | 8 |
| 1 | 1 | 1 | 1 | 1 | 8 |
| 1 | 1 | 1 | 1 | 1 | 8 |
| 1 | 1 | 1 | 1 | 1 | 8 |
| 1 | 1 | 1 | 1 | 1 | 8 |
| 1 | 1 | 1 | 1 | 1 | 8 |
| 1 | 1 | 1 | 1 | 1 | 8 |
| 1 | 1 | 1 | 1 | 1 | 8 |
| 1 | 1 | 1 | 1 | 1 | 8 |
| 1 | 1 | 1 | 1 | 1 | 8 |
| 1 | 1 | 1 | 1 | 1 | 8 |
| 1 | 1 | 1 | 1 | 1 | 8 |
| 1 | 1 | 1 | 1 | 1 | 8 |
| 1 | 1 | 1 | 1 | 1 | 8 |

| | | | | | |
|---|---|---|---|---|---|
| 1 | 1 | 1 | 1 | 1 | 8 |
| 1 | 1 | 1 | 1 | 1 | 8 |
| 1 | 1 | 1 | 1 | 1 | 8 |
| 1 | 1 | 1 | 1 | 1 | 8 |
| 1 | 1 | 1 | 1 | 1 | 8 |
| 1 | 1 | 1 | 1 | 1 | 8 |
| 1 | 1 | 1 | 1 | 1 | 8 |
| 1 | 1 | 1 | 1 | 1 | 8 |
| 1 | 1 | 1 | 1 | 1 | 8 |
| 1 | 1 | 1 | 1 | 1 | 8 |
| 1 | 1 | 1 | 1 | 1 | 8 |
| 1 | 1 | 1 | 1 | 1 | 8 |
| 1 | 1 | 1 | 1 | 1 | 8 |
| 1 | 1 | 1 | 1 | 1 | 8 |
| 1 | 1 | 1 | 1 | 1 | 8 |
| 1 | 1 | 1 | 1 | 1 | 8 |
| 1 | 1 | 1 | 1 | 1 | 8 |
| 1 | 1 | 1 | 1 | 1 | 8 |
| 1 | 1 | 1 | 1 | 1 | 8 |
| 1 | 1 | 1 | 1 | 1 | 8 |
| 1 | 1 | 1 | 1 | 1 | 8 |
| 1 | 1 | 1 | 1 | 1 | 8 |
| 1 | 1 | 1 | 1 | 1 | 8 |
| 1 | 1 | 1 | 1 | 1 | 8 |
| 1 | 1 | 1 | 1 | 1 | 8 |
| 1 | 1 | 1 | 1 | 1 | 8 |
| 1 | 1 | 1 | 1 | 1 | 8 |
| 1 | 1 | 1 | 1 | 1 | 8 |
| 1 | 1 | 1 | 1 | 1 | 8 |
| 1 | 1 | 1 | 1 | 1 | 8 |
| 1 | 1 | 1 | 1 | 1 | 8 |
| 1 | 1 | 1 | 1 | 1 | 8 |
| 1 | 1 | 1 | 1 | 1 | 8 |
| 1 | 1 | 1 | 1 | 1 | 8 |
| 1 | 1 | 1 | 1 | 1 | 8 |
| 1 | 1 | 1 | 1 | 1 | 8 |
| 1 | 1 | 1 | 1 | 1 | 8 |
| 1 | 1 | 1 | 1 | 1 | 8 |
| 1 | 1 | 1 | 1 | 1 | 8 |
| 1 | 1 | 1 | 1 | 1 | 8 |
| 1 | 1 | 1 | 1 | 1 | 8 |
| 1 | 1 | 1 | 1 | 1 | 8 |
| 0 | 1 | 0 | 0 | 0 | 2 |
| 0 | 1 | 0 | 0 | 0 | 2 |

| | | | | | |
|---|---|---|---|---|---|
| 0 | 1 | 0 | 0 | 0 | 2 |
| 0 | 1 | 0 | 0 | 0 | 2 |
| 0 | 1 | 0 | 0 | 0 | 2 |
| 0 | 1 | 0 | 0 | 0 | 2 |
| 0 | 1 | 0 | 0 | 0 | 2 |
| 0 | 1 | 0 | 0 | 0 | 2 |
| 0 | 1 | 0 | 0 | 0 | 2 |
| 0 | 1 | 0 | 0 | 0 | 2 |
| 0 | 1 | 0 | 0 | 0 | 2 |
| 0 | 1 | 0 | 0 | 0 | 2 |
| 0 | 1 | 0 | 0 | 0 | 2 |
| 0 | 1 | 0 | 0 | 0 | 2 |
| 0 | 1 | 0 | 0 | 0 | 2 |
| 0 | 1 | 0 | 0 | 0 | 2 |
| 0 | 1 | 0 | 0 | 0 | 2 |
| 0 | 1 | 0 | 0 | 0 | 2 |
| 0 | 1 | 0 | 0 | 0 | 2 |
| 0 | 1 | 0 | 0 | 0 | 2 |
| 0 | 1 | 0 | 0 | 0 | 2 |
| 0 | 1 | 0 | 0 | 0 | 2 |
| 0 | 1 | 0 | 0 | 0 | 2 |
| 0 | 1 | 0 | 0 | 0 | 2 |
| 0 | 0 | 0 | 0 | 1 | 1 |
| 0 | 0 | 0 | 0 | 1 | 1 |
| 0 | 0 | 0 | 0 | 1 | 1 |
| 0 | 0 | 0 | 0 | 1 | 1 |
| 0 | 0 | 0 | 0 | 1 | 1 |
| 0 | 0 | 0 | 0 | 1 | 1 |
| 0 | 0 | 0 | 0 | 1 | 1 |
| 0 | 0 | 0 | 0 | 1 | 1 |
| 0 | 0 | 0 | 0 | 1 | 1 |
| 0 | 0 | 0 | 0 | 1 | 1 |
| 1 | 1 | 1 | 1 | 1 | 8 |
| 1 | 1 | 1 | 1 | 1 | 8 |
| 1 | 1 | 1 | 1 | 1 | 8 |
| 1 | 1 | 1 | 1 | 1 | 8 |
| 1 | 1 | 1 | 1 | 1 | 8 |
| 1 | 1 | 1 | 1 | 1 | 8 |
| 1 | 1 | 1 | 1 | 1 | 8 |
| 1 | 1 | 1 | 1 | 1 | 8 |
| 1 | 1 | 1 | 1 | 1 | 8 |
| 1 | 1 | 1 | 1 | 1 | 8 |
| 0 | 1 | 1 | 1 | 0 | 5 |
| 0 | 1 | 1 | 1 | 0 | 5 |

| | | | | | |
|---|---|---|---|---|---|
| 0 | 1 | 1 | 1 | 0 | 5 |
| 0 | 1 | 1 | 1 | 0 | 5 |
| 0 | 1 | 1 | 1 | 0 | 5 |
| 0 | 1 | 1 | 1 | 0 | 5 |
| 0 | 1 | 1 | 1 | 0 | 5 |
| 0 | 1 | 1 | 1 | 0 | 5 |
| 0 | 1 | 1 | 1 | 0 | 5 |
| 0 | 1 | 1 | 1 | 0 | 5 |
| 1 | 1 | 1 | 1 | 1 | 8 |
| 1 | 1 | 1 | 1 | 1 | 8 |
| 1 | 1 | 1 | 1 | 1 | 8 |
| 1 | 1 | 1 | 1 | 1 | 8 |
| 1 | 1 | 1 | 1 | 1 | 8 |
| 1 | 1 | 1 | 1 | 1 | 8 |
| 1 | 1 | 1 | 1 | 1 | 8 |
| 1 | 1 | 1 | 1 | 1 | 8 |
| 1 | 1 | 1 | 1 | 1 | 8 |
| 1 | 1 | 1 | 1 | 1 | 8 |
| 1 | 0 | 0 | 0 | 0 | 2 |
| 1 | 0 | 0 | 0 | 0 | 2 |
| 1 | 0 | 0 | 0 | 0 | 2 |
| 1 | 0 | 0 | 0 | 0 | 2 |
| 1 | 0 | 0 | 0 | 0 | 2 |
| 1 | 0 | 0 | 0 | 0 | 2 |
| 1 | 0 | 0 | 0 | 0 | 2 |
| 1 | 0 | 0 | 0 | 0 | 2 |
| 1 | 0 | 0 | 0 | 0 | 2 |
| 1 | 0 | 0 | 0 | 0 | 2 |
| 1 | 0 | 0 | 0 | 0 | 2 |
| 1 | 0 | 0 | 0 | 0 | 2 |
| 1 | 0 | 0 | 0 | 0 | 2 |
| 1 | 0 | 0 | 0 | 0 | 2 |
| 1 | 0 | 0 | 0 | 0 | 2 |
| 1 | 0 | 0 | 0 | 0 | 2 |
| 1 | 0 | 0 | 0 | 0 | 2 |
| 1 | 0 | 0 | 0 | 0 | 2 |
| 1 | 0 | 0 | 0 | 0 | 2 |
| 1 | 0 | 0 | 0 | 0 | 2 |
| 1 | 0 | 0 | 0 | 0 | 2 |
| 1 | 0 | 0 | 0 | 0 | 2 |
| 1 | 0 | 0 | 0 | 0 | 2 |
| 1 | 0 | 0 | 0 | 0 | 2 |
| 1 | 0 | 0 | 0 | 0 | 2 |
| 1 | 0 | 0 | 0 | 0 | 2 |

| | | | | | |
|---|---|---|---|---|---|
| 1 | 0 | 0 | 0 | 0 | 2 |
| 1 | 0 | 0 | 0 | 0 | 2 |
| 1 | 0 | 0 | 0 | 0 | 2 |
| 1 | 0 | 0 | 0 | 0 | 2 |
| 1 | 0 | 0 | 0 | 0 | 2 |
| 1 | 0 | 0 | 0 | 0 | 2 |
| 1 | 0 | 0 | 0 | 0 | 2 |
| 1 | 0 | 0 | 0 | 0 | 2 |
| 1 | 0 | 0 | 0 | 0 | 2 |
| 1 | 0 | 0 | 0 | 0 | 2 |
| 1 | 0 | 0 | 0 | 0 | 2 |
| 1 | 0 | 0 | 0 | 0 | 2 |
| 1 | 0 | 0 | 0 | 0 | 2 |
| 1 | 0 | 0 | 0 | 0 | 2 |
| 1 | 0 | 0 | 0 | 0 | 2 |
| 1 | 0 | 0 | 0 | 0 | 2 |
| 1 | 0 | 0 | 0 | 0 | 2 |
| 1 | 0 | 0 | 0 | 0 | 2 |
| 1 | 0 | 0 | 0 | 0 | 2 |
| 1 | 0 | 0 | 0 | 0 | 2 |
| 1 | 0 | 0 | 0 | 0 | 2 |
| 1 | 0 | 0 | 0 | 0 | 2 |
| 1 | 0 | 0 | 0 | 0 | 2 |
| 1 | 0 | 0 | 0 | 0 | 2 |
| 1 | 0 | 0 | 0 | 0 | 2 |
| 1 | 0 | 0 | 0 | 0 | 2 |
| 1 | 0 | 0 | 0 | 0 | 2 |
| 1 | 0 | 0 | 0 | 0 | 2 |
| 1 | 0 | 0 | 0 | 0 | 2 |
| 1 | 0 | 0 | 0 | 0 | 2 |
| 1 | 0 | 0 | 0 | 0 | 2 |
| 1 | 0 | 0 | 0 | 0 | 2 |
| 1 | 0 | 0 | 0 | 0 | 2 |
| 1 | 0 | 0 | 0 | 0 | 2 |
| 1 | 0 | 0 | 0 | 0 | 2 |
| 1 | 0 | 0 | 0 | 0 | 2 |
| 1 | 0 | 0 | 0 | 0 | 2 |
| 1 | 0 | 0 | 0 | 0 | 2 |
| 1 | 0 | 0 | 0 | 0 | 2 |
| 1 | 0 | 0 | 0 | 0 | 2 |
| 1 | 0 | 0 | 0 | 0 | 2 |
| 1 | 0 | 0 | 0 | 0 | 2 |
| 1 | 0 | 0 | 0 | 0 | 2 |
| 1 | 0 | 0 | 0 | 0 | 2 |

| | | | | | |
|---|---|---|---|---|---|
| 1 | 0 | 0 | 0 | 0 | 2 |
| 1 | 0 | 0 | 0 | 0 | 2 |
| 1 | 0 | 0 | 0 | 0 | 2 |
| 1 | 0 | 0 | 0 | 0 | 2 |
| 1 | 0 | 0 | 0 | 0 | 2 |
| 1 | 0 | 0 | 0 | 0 | 2 |
| 1 | 0 | 0 | 0 | 0 | 2 |
| 1 | 0 | 0 | 0 | 0 | 2 |
| 1 | 0 | 0 | 0 | 0 | 2 |
| 1 | 0 | 0 | 0 | 0 | 2 |
| 1 | 0 | 0 | 0 | 0 | 2 |
| 1 | 0 | 0 | 0 | 0 | 2 |
| 1 | 0 | 0 | 0 | 0 | 2 |
| 1 | 0 | 0 | 0 | 0 | 2 |
| 1 | 0 | 0 | 0 | 0 | 2 |
| 1 | 0 | 0 | 0 | 0 | 2 |
| 1 | 0 | 0 | 0 | 0 | 2 |
| 1 | 0 | 0 | 0 | 0 | 2 |
| 1 | 0 | 0 | 0 | 0 | 2 |
| 1 | 0 | 0 | 0 | 0 | 2 |
| 1 | 0 | 0 | 0 | 0 | 2 |
| 1 | 0 | 0 | 0 | 0 | 2 |
| 1 | 0 | 0 | 0 | 0 | 2 |
| 1 | 0 | 0 | 0 | 0 | 2 |
| 1 | 0 | 0 | 0 | 0 | 2 |
| 1 | 0 | 0 | 0 | 0 | 2 |
| 1 | 0 | 0 | 0 | 0 | 2 |
| 1 | 0 | 0 | 0 | 0 | 2 |
| 1 | 0 | 0 | 0 | 0 | 2 |
| 1 | 0 | 0 | 0 | 0 | 2 |
| 1 | 0 | 0 | 0 | 0 | 2 |
| 1 | 0 | 0 | 0 | 0 | 2 |
| 1 | 0 | 0 | 0 | 0 | 2 |
| 1 | 0 | 0 | 0 | 0 | 2 |
| 1 | 0 | 0 | 0 | 0 | 2 |
| 1 | 0 | 0 | 0 | 0 | 2 |
| 1 | 0 | 0 | 0 | 0 | 2 |
| 1 | 0 | 0 | 0 | 0 | 2 |
| 1 | 0 | 0 | 0 | 0 | 2 |
| 1 | 0 | 0 | 0 | 0 | 2 |
| 1 | 0 | 0 | 0 | 0 | 2 |
| 1 | 0 | 0 | 0 | 0 | 2 |
| 0 | 1 | 1 | 1 | 0 | 4 |
| 0 | 1 | 1 | 1 | 0 | 4 |

| | | | | | |
|---|---|---|---|---|---|
| 0 | 1 | 1 | 1 | 0 | 4 |
| 0 | 1 | 1 | 1 | 0 | 4 |
| 0 | 1 | 1 | 1 | 0 | 4 |
| 0 | 1 | 1 | 1 | 0 | 4 |
| 0 | 1 | 1 | 1 | 0 | 4 |
| 0 | 1 | 1 | 1 | 0 | 4 |
| 0 | 1 | 1 | 1 | 0 | 4 |
| 0 | 1 | 1 | 1 | 0 | 4 |
| 0 | 1 | 1 | 1 | 0 | 4 |
| 0 | 1 | 1 | 1 | 0 | 4 |
| 0 | 1 | 1 | 1 | 0 | 4 |
| 0 | 1 | 1 | 1 | 0 | 4 |
| 0 | 1 | 1 | 1 | 0 | 4 |
| 0 | 1 | 1 | 1 | 0 | 4 |
| 0 | 1 | 1 | 1 | 0 | 4 |
| 0 | 1 | 1 | 1 | 0 | 4 |
| 0 | 1 | 1 | 1 | 0 | 4 |
| 0 | 1 | 1 | 1 | 0 | 4 |
| 0 | 1 | 1 | 1 | 0 | 4 |
| 0 | 1 | 1 | 1 | 0 | 4 |
| 0 | 1 | 1 | 1 | 0 | 4 |
| 0 | 1 | 1 | 1 | 0 | 4 |
| 0 | 1 | 1 | 1 | 0 | 4 |
| 0 | 1 | 1 | 1 | 0 | 4 |
| 0 | 1 | 1 | 1 | 0 | 4 |
| 0 | 1 | 1 | 1 | 0 | 4 |
| 0 | 1 | 1 | 1 | 0 | 4 |
| 0 | 1 | 1 | 1 | 0 | 4 |
| 0 | 1 | 1 | 1 | 0 | 4 |
| 0 | 1 | 1 | 1 | 0 | 4 |
| 0 | 1 | 1 | 1 | 0 | 4 |
| 0 | 1 | 1 | 1 | 0 | 4 |
| 0 | 1 | 1 | 1 | 0 | 4 |
| 0 | 1 | 1 | 1 | 0 | 4 |
| 0 | 1 | 1 | 1 | 0 | 4 |
| 0 | 1 | 1 | 1 | 0 | 4 |
| 0 | 1 | 1 | 1 | 0 | 4 |
| 0 | 1 | 1 | 1 | 0 | 4 |
| 0 | 1 | 1 | 1 | 0 | 4 |
| 0 | 1 | 1 | 1 | 0 | 4 |
| 0 | 1 | 1 | 1 | 0 | 4 |
| 0 | 1 | 1 | 1 | 0 | 4 |
| 0 | 1 | 1 | 1 | 0 | 4 |
| 0 | 1 | 1 | 1 | 0 | 4 |

| | | | | | |
|---|---|---|---|---|---|
| 0 | 1 | 1 | 1 | 0 | 4 |
| 0 | 1 | 1 | 1 | 0 | 4 |
| 0 | 1 | 1 | 1 | 0 | 4 |
| 0 | 1 | 1 | 1 | 0 | 4 |
| 0 | 1 | 1 | 1 | 0 | 4 |
| 0 | 1 | 1 | 1 | 0 | 4 |
| 0 | 1 | 1 | 1 | 0 | 4 |
| 0 | 1 | 1 | 1 | 0 | 4 |
| 0 | 1 | 1 | 1 | 0 | 4 |
| 0 | 1 | 1 | 1 | 0 | 4 |
| 0 | 1 | 1 | 1 | 1 | 5 |
| 0 | 1 | 1 | 1 | 1 | 5 |
| 0 | 1 | 1 | 1 | 1 | 5 |
| 0 | 1 | 1 | 1 | 1 | 5 |
| 0 | 1 | 1 | 1 | 1 | 5 |
| 0 | 1 | 1 | 1 | 1 | 5 |
| 0 | 1 | 1 | 1 | 1 | 5 |
| 0 | 1 | 1 | 1 | 1 | 5 |
| 0 | 1 | 1 | 1 | 1 | 5 |
| 0 | 1 | 1 | 1 | 1 | 5 |
| 0 | 1 | 1 | 1 | 1 | 5 |
| 0 | 1 | 1 | 1 | 1 | 5 |
| 0 | 1 | 1 | 1 | 1 | 5 |
| 0 | 1 | 1 | 1 | 1 | 5 |
| 0 | 1 | 1 | 1 | 1 | 5 |
| 0 | 1 | 1 | 1 | 1 | 5 |
| 0 | 1 | 1 | 1 | 1 | 5 |
| 0 | 1 | 1 | 1 | 1 | 5 |
| 0 | 1 | 1 | 1 | 1 | 5 |
| 0 | 1 | 1 | 1 | 1 | 5 |
| 0 | 1 | 1 | 1 | 1 | 5 |
| 0 | 1 | 1 | 1 | 1 | 5 |
| 0 | 1 | 1 | 1 | 1 | 5 |
| 0 | 1 | 1 | 1 | 1 | 5 |
| 0 | 1 | 1 | 1 | 1 | 5 |
| 0 | 1 | 1 | 1 | 1 | 5 |
| 0 | 1 | 1 | 1 | 1 | 5 |
| 0 | 1 | 1 | 1 | 1 | 5 |
| 0 | 1 | 1 | 1 | 1 | 5 |
| 0 | 1 | 1 | 1 | 1 | 5 |
| 0 | 1 | 1 | 1 | 1 | 5 |
| 0 | 1 | 1 | 1 | 1 | 5 |
| 0 | 1 | 1 | 1 | 1 | 5 |
| 0 | 1 | 1 | 1 | 1 | 5 |

| | | | | | |
|---|---|---|---|---|---|
| 0 | 1 | 1 | 1 | 1 | 5 |
| 0 | 1 | 1 | 1 | 1 | 5 |
| 0 | 1 | 1 | 1 | 1 | 5 |
| 0 | 1 | 1 | 1 | 1 | 5 |
| 0 | 1 | 1 | 1 | 1 | 5 |
| 0 | 1 | 1 | 1 | 1 | 5 |
| 0 | 1 | 1 | 1 | 1 | 5 |
| 0 | 1 | 1 | 1 | 1 | 5 |
| 0 | 1 | 1 | 1 | 1 | 5 |
| 0 | 1 | 1 | 1 | 1 | 5 |
| 0 | 1 | 1 | 1 | 1 | 5 |
| 0 | 1 | 1 | 1 | 1 | 5 |
| 0 | 1 | 1 | 1 | 1 | 5 |
| 0 | 1 | 1 | 1 | 1 | 5 |
| 0 | 1 | 1 | 1 | 1 | 5 |
| 0 | 1 | 1 | 1 | 1 | 5 |
| 0 | 1 | 1 | 1 | 1 | 5 |
| 0 | 1 | 1 | 1 | 1 | 5 |
| 0 | 1 | 1 | 1 | 1 | 5 |
| 0 | 1 | 1 | 1 | 1 | 5 |
| 0 | 1 | 1 | 1 | 1 | 5 |
| 0 | 1 | 1 | 1 | 1 | 5 |
| 1 | 1 | 1 | 1 | 1 | 8 |
| 1 | 1 | 1 | 1 | 1 | 8 |
| 1 | 1 | 1 | 1 | 1 | 8 |
| 1 | 1 | 1 | 1 | 1 | 8 |
| 1 | 1 | 1 | 1 | 1 | 8 |
| 1 | 1 | 1 | 1 | 1 | 8 |
| 1 | 1 | 1 | 1 | 1 | 8 |
| 1 | 1 | 1 | 1 | 1 | 8 |
| 1 | 1 | 1 | 1 | 1 | 8 |
| 1 | 1 | 1 | 1 | 1 | 8 |
| 1 | 1 | 1 | 1 | 1 | 8 |
| 1 | 1 | 1 | 1 | 1 | 8 |
| 1 | 1 | 1 | 1 | 1 | 8 |
| 1 | 1 | 1 | 1 | 1 | 8 |
| 1 | 1 | 1 | 1 | 1 | 8 |
| 1 | 1 | 1 | 1 | 1 | 8 |
| 1 | 1 | 1 | 1 | 1 | 8 |
| 1 | 1 | 1 | 1 | 1 | 8 |
| 1 | 1 | 1 | 1 | 1 | 8 |
| 1 | 1 | 1 | 1 | 1 | 8 |
| 1 | 1 | 1 | 1 | 1 | 8 |
| 1 | 1 | 1 | 1 | 1 | 8 |

| | | | | | |
|---|---|---|---|---|---|
| 1 | 1 | 1 | 1 | 1 | 8 |
| 1 | 1 | 1 | 1 | 1 | 8 |
| 0 | 0 | 0 | 0 | 1 | 1 |
| 0 | 0 | 0 | 0 | 1 | 1 |
| 0 | 0 | 0 | 0 | 1 | 1 |
| 0 | 0 | 0 | 0 | 1 | 1 |
| 0 | 0 | 0 | 0 | 1 | 1 |
| 0 | 0 | 0 | 0 | 1 | 1 |
| 0 | 0 | 0 | 0 | 1 | 1 |
| 0 | 0 | 0 | 0 | 1 | 1 |
| 0 | 0 | 0 | 0 | 1 | 1 |
| 0 | 0 | 0 | 0 | 1 | 1 |
| 0 | 0 | 0 | 0 | 1 | 1 |
| 0 | 0 | 0 | 0 | 1 | 1 |
| 0 | 0 | 0 | 0 | 1 | 1 |
| 0 | 0 | 0 | 0 | 1 | 1 |
| 0 | 0 | 0 | 0 | 1 | 1 |
| 0 | 0 | 0 | 0 | 1 | 1 |
| 0 | 0 | 0 | 0 | 1 | 1 |
| 0 | 0 | 0 | 0 | 1 | 1 |
| 0 | 0 | 0 | 0 | 1 | 1 |
| 0 | 0 | 0 | 0 | 1 | 1 |
| 0 | 0 | 0 | 0 | 1 | 1 |
| 0 | 0 | 0 | 0 | 1 | 1 |
| 0 | 0 | 0 | 0 | 1 | 1 |
| 0 | 0 | 0 | 0 | 1 | 1 |
| 0 | 0 | 0 | 0 | 1 | 1 |
| 0 | 0 | 0 | 0 | 1 | 1 |
| 0 | 0 | 0 | 0 | 1 | 1 |
| 0 | 0 | 0 | 0 | 1 | 1 |
| 0 | 0 | 0 | 0 | 1 | 1 |
| 0 | 0 | 0 | 0 | 1 | 1 |
| 0 | 0 | 0 | 0 | 1 | 1 |
| 0 | 0 | 0 | 0 | 1 | 1 |
| 0 | 0 | 0 | 0 | 1 | 1 |
| 0 | 0 | 0 | 0 | 1 | 1 |
| 0 | 0 | 0 | 0 | 1 | 1 |
| 0 | 0 | 0 | 0 | 1 | 1 |
| 0 | 0 | 0 | 0 | 1 | 1 |
| 0 | 0 | 0 | 0 | 1 | 1 |
| 0 | 0 | 0 | 0 | 1 | 1 |
| 0 | 0 | 0 | 0 | 1 | 1 |
| 0 | 0 | 0 | 0 | 1 | 1 |
| 0 | 0 | 0 | 0 | 1 | 1 |

| | | | | | |
|---|---|---|---|---|---|
| 0 | 0 | 0 | 0 | 1 | 1 |
| 0 | 0 | 0 | 0 | 1 | 1 |
| 0 | 0 | 0 | 0 | 1 | 1 |
| 0 | 0 | 0 | 0 | 1 | 1 |
| 0 | 0 | 0 | 0 | 1 | 1 |
| 0 | 0 | 0 | 0 | 1 | 1 |
| 0 | 0 | 0 | 0 | 1 | 1 |
| 0 | 0 | 0 | 0 | 1 | 1 |
| 0 | 0 | 0 | 0 | 1 | 1 |
| 0 | 0 | 0 | 0 | 1 | 1 |
| 0 | 0 | 0 | 0 | 1 | 1 |
| 0 | 0 | 0 | 0 | 1 | 1 |
| 0 | 0 | 0 | 0 | 1 | 1 |
| 0 | 0 | 0 | 0 | 1 | 1 |
| 1 | 1 | 1 | 1 | 1 | 8 |
| 1 | 1 | 1 | 1 | 1 | 8 |
| 1 | 1 | 1 | 1 | 1 | 8 |
| 1 | 1 | 1 | 1 | 1 | 8 |
| 1 | 1 | 1 | 1 | 1 | 8 |
| 1 | 1 | 1 | 1 | 1 | 8 |
| 1 | 1 | 1 | 1 | 1 | 8 |
| 1 | 1 | 1 | 1 | 1 | 8 |
| 1 | 1 | 1 | 1 | 1 | 8 |
| 1 | 1 | 1 | 1 | 1 | 8 |
| 1 | 1 | 1 | 1 | 1 | 8 |
| 1 | 1 | 1 | 1 | 1 | 8 |
| 1 | 1 | 1 | 1 | 1 | 8 |
| 1 | 1 | 1 | 1 | 1 | 8 |
| 1 | 1 | 1 | 1 | 1 | 8 |
| 1 | 1 | 1 | 1 | 1 | 8 |
| 1 | 1 | 1 | 1 | 1 | 8 |
| 1 | 1 | 1 | 1 | 1 | 8 |
| 1 | 1 | 1 | 1 | 1 | 8 |
| 1 | 1 | 1 | 1 | 1 | 8 |
| 1 | 1 | 1 | 1 | 1 | 8 |
| 1 | 1 | 1 | 1 | 1 | 8 |
| 1 | 1 | 1 | 1 | 1 | 8 |
| 1 | 1 | 1 | 1 | 1 | 8 |
| 1 | 1 | 1 | 1 | 1 | 8 |
| 1 | 1 | 1 | 1 | 1 | 8 |
| 1 | 1 | 1 | 1 | 1 | 8 |
| 1 | 1 | 1 | 1 | 1 | 8 |
| 1 | 1 | 1 | 1 | 1 | 8 |
| 1 | 1 | 1 | 1 | 1 | 8 |

| | | | | | |
|---|---|---|---|---|---|
| 1 | 1 | 1 | 1 | 1 | 8 |
| 1 | 1 | 1 | 1 | 1 | 8 |
| 1 | 1 | 1 | 1 | 1 | 8 |
| 1 | 1 | 1 | 1 | 1 | 8 |
| 1 | 1 | 1 | 1 | 1 | 8 |
| 1 | 1 | 1 | 1 | 1 | 8 |
| 1 | 1 | 1 | 1 | 1 | 8 |
| 1 | 1 | 1 | 1 | 1 | 8 |
| 1 | 1 | 1 | 1 | 1 | 8 |
| 1 | 1 | 1 | 1 | 1 | 8 |
| 1 | 1 | 1 | 1 | 1 | 8 |
| 1 | 1 | 1 | 1 | 1 | 8 |
| 1 | 1 | 1 | 1 | 1 | 8 |
| 1 | 1 | 1 | 1 | 1 | 8 |
| 1 | 1 | 1 | 1 | 1 | 8 |
| 1 | 1 | 1 | 1 | 1 | 8 |
| 1 | 1 | 1 | 1 | 1 | 8 |
| 1 | 1 | 1 | 1 | 1 | 8 |
| 1 | 1 | 1 | 1 | 1 | 8 |
| 1 | 1 | 1 | 1 | 1 | 8 |
| 1 | 1 | 1 | 1 | 1 | 8 |
| 1 | 1 | 1 | 1 | 1 | 8 |
| 1 | 1 | 1 | 1 | 1 | 8 |
| 1 | 1 | 1 | 1 | 1 | 8 |
| 1 | 1 | 1 | 1 | 1 | 8 |
| 1 | 1 | 1 | 1 | 1 | 8 |
| 1 | 1 | 1 | 1 | 1 | 8 |
| 1 | 1 | 1 | 1 | 1 | 8 |
| 1 | 1 | 1 | 1 | 1 | 8 |
| 1 | 1 | 1 | 1 | 1 | 8 |
| 1 | 1 | 1 | 1 | 1 | 8 |
| 1 | 1 | 1 | 1 | 1 | 8 |
| 1 | 1 | 1 | 1 | 1 | 8 |
| 1 | 1 | 1 | 1 | 1 | 8 |
| 1 | 1 | 1 | 1 | 1 | 8 |
| 1 | 1 | 1 | 1 | 1 | 8 |
| 1 | 1 | 1 | 1 | 1 | 8 |
| 1 | 1 | 1 | 1 | 1 | 8 |
| 1 | 1 | 1 | 1 | 1 | 8 |
| 1 | 1 | 1 | 1 | 1 | 8 |
| 1 | 1 | 1 | 1 | 1 | 8 |
| 1 | 1 | 1 | 1 | 1 | 8 |
| 1 | 1 | 1 | 1 | 1 | 8 |
| 1 | 1 | 1 | 1 | 1 | 8 |

| | | | | | |
|---|---|---|---|---|---|
| 1 | 1 | 1 | 1 | 1 | 8 |
| 1 | 1 | 1 | 1 | 1 | 8 |
| 1 | 1 | 1 | 1 | 1 | 8 |
| 1 | 1 | 1 | 1 | 1 | 8 |
| 1 | 1 | 1 | 1 | 1 | 8 |
| 1 | 1 | 1 | 1 | 1 | 8 |
| 1 | 1 | 1 | 1 | 1 | 8 |
| 1 | 1 | 1 | 1 | 1 | 8 |
| 1 | 1 | 1 | 1 | 1 | 8 |
| 1 | 1 | 1 | 1 | 1 | 8 |
| 1 | 1 | 1 | 1 | 1 | 8 |
| 1 | 1 | 1 | 1 | 1 | 8 |
| 1 | 1 | 1 | 1 | 1 | 8 |
| 1 | 1 | 1 | 1 | 1 | 8 |
| 1 | 1 | 1 | 1 | 1 | 8 |
| 1 | 1 | 1 | 1 | 1 | 8 |
| 1 | 1 | 1 | 1 | 1 | 8 |
| 1 | 1 | 1 | 1 | 1 | 8 |
| 1 | 1 | 1 | 1 | 1 | 8 |
| 1 | 1 | 1 | 1 | 1 | 8 |
| 1 | 1 | 1 | 1 | 1 | 8 |
| 1 | 1 | 1 | 1 | 1 | 8 |
| 1 | 1 | 1 | 1 | 1 | 8 |
| 1 | 1 | 1 | 1 | 1 | 8 |
| 1 | 1 | 1 | 1 | 1 | 8 |
| 1 | 1 | 1 | 1 | 1 | 8 |
| 1 | 1 | 1 | 1 | 1 | 8 |
| 1 | 1 | 1 | 1 | 1 | 8 |
| 1 | 1 | 1 | 1 | 1 | 8 |
| 1 | 1 | 1 | 1 | 1 | 8 |
| 1 | 1 | 1 | 1 | 1 | 8 |
| 1 | 1 | 1 | 1 | 1 | 8 |
| 1 | 1 | 1 | 1 | 1 | 8 |
| 1 | 1 | 1 | 1 | 1 | 8 |
| 1 | 1 | 1 | 1 | 1 | 8 |
| 1 | 1 | 1 | 1 | 1 | 8 |
| 1 | 1 | 1 | 1 | 1 | 8 |
| 1 | 1 | 1 | 1 | 1 | 8 |
| 0 | 1 | 0 | 0 | 0 | 2 |
| 0 | 1 | 0 | 0 | 0 | 2 |
| 0 | 1 | 0 | 0 | 0 | 2 |
| 0 | 1 | 0 | 0 | 0 | 2 |
| 0 | 1 | 0 | 0 | 0 | 2 |
| 0 | 1 | 0 | 0 | 0 | 2 |

| | | | | | |
|---|---|---|---|---|---|
| 0 | 1 | 0 | 0 | 0 | 2 |
| 0 | 1 | 0 | 0 | 0 | 2 |
| 0 | 1 | 0 | 0 | 0 | 2 |
| 0 | 1 | 0 | 0 | 0 | 2 |
| 0 | 1 | 0 | 0 | 0 | 2 |
| 0 | 1 | 0 | 0 | 0 | 2 |
| 0 | 1 | 0 | 0 | 0 | 2 |
| 0 | 1 | 0 | 0 | 0 | 2 |
| 0 | 1 | 0 | 0 | 0 | 2 |
| 0 | 1 | 0 | 0 | 0 | 2 |
| 0 | 1 | 0 | 0 | 0 | 2 |
| 0 | 1 | 0 | 0 | 0 | 2 |
| 0 | 1 | 0 | 0 | 0 | 2 |
| 0 | 1 | 0 | 0 | 0 | 2 |
| 0 | 1 | 0 | 0 | 0 | 2 |
| 0 | 1 | 0 | 0 | 0 | 2 |
| 0 | 1 | 0 | 0 | 0 | 2 |
| 0 | 1 | 0 | 0 | 0 | 2 |
| 0 | 1 | 0 | 0 | 0 | 2 |
| 0 | 1 | 0 | 0 | 0 | 2 |
| 0 | 1 | 0 | 0 | 0 | 2 |
| 0 | 1 | 0 | 0 | 0 | 2 |
| 0 | 1 | 0 | 0 | 0 | 2 |
| 0 | 1 | 0 | 0 | 0 | 2 |
| 0 | 1 | 0 | 0 | 0 | 2 |
| 0 | 1 | 0 | 0 | 0 | 2 |
| 1 | 1 | 1 | 1 | 1 | 8 |
| 1 | 1 | 1 | 1 | 1 | 8 |
| 1 | 1 | 1 | 1 | 1 | 8 |
| 1 | 1 | 1 | 1 | 1 | 8 |
| 1 | 1 | 1 | 1 | 1 | 8 |
| 1 | 1 | 1 | 1 | 1 | 8 |
| 1 | 1 | 1 | 1 | 1 | 8 |
| 1 | 1 | 1 | 1 | 1 | 8 |
| 1 | 1 | 1 | 1 | 1 | 8 |
| 1 | 1 | 1 | 1 | 1 | 8 |
| 1 | 1 | 1 | 1 | 1 | 8 |
| 1 | 1 | 1 | 1 | 1 | 8 |
| 1 | 1 | 1 | 1 | 1 | 8 |
| 1 | 1 | 1 | 1 | 1 | 8 |
| 1 | 1 | 1 | 1 | 1 | 8 |
| 1 | 1 | 1 | 1 | 1 | 8 |
| 1 | 1 | 1 | 1 | 1 | 8 |
| 1 | 1 | 1 | 1 | 1 | 8 |

| | | | | | |
|---|---|---|---|---|---|
| 1 | 1 | 1 | 1 | 1 | 8 |
| 1 | 1 | 1 | 1 | 1 | 8 |
| 1 | 1 | 1 | 1 | 1 | 8 |
| 1 | 1 | 1 | 1 | 1 | 8 |
| 1 | 1 | 1 | 1 | 1 | 8 |
| 1 | 1 | 1 | 1 | 1 | 8 |
| 1 | 1 | 1 | 1 | 1 | 8 |
| 1 | 1 | 1 | 1 | 1 | 8 |
| 1 | 1 | 1 | 1 | 1 | 8 |
| 1 | 1 | 1 | 1 | 1 | 8 |
| 1 | 1 | 1 | 1 | 1 | 8 |
| 1 | 1 | 1 | 1 | 1 | 8 |
| 1 | 1 | 1 | 1 | 1 | 8 |
| 1 | 1 | 1 | 1 | 1 | 8 |
| 1 | 1 | 1 | 1 | 1 | 8 |
| 1 | 1 | 1 | 1 | 1 | 8 |
| 1 | 1 | 1 | 1 | 1 | 8 |
| 1 | 1 | 1 | 1 | 1 | 8 |
| 1 | 1 | 1 | 1 | 1 | 8 |
| 1 | 1 | 1 | 1 | 1 | 8 |
| 1 | 1 | 1 | 1 | 1 | 8 |
| 1 | 1 | 1 | 1 | 1 | 8 |
| 1 | 1 | 1 | 1 | 1 | 8 |
| 1 | 1 | 1 | 1 | 1 | 8 |
| 1 | 1 | 1 | 1 | 1 | 8 |
| 1 | 1 | 1 | 1 | 1 | 8 |
| 1 | 1 | 1 | 1 | 1 | 8 |
| 1 | 1 | 1 | 1 | 1 | 8 |
| 1 | 1 | 1 | 1 | 1 | 8 |
| 1 | 1 | 1 | 1 | 1 | 8 |
| 1 | 1 | 1 | 1 | 1 | 8 |
| 1 | 1 | 1 | 1 | 1 | 8 |
| 1 | 1 | 1 | 1 | 1 | 8 |
| 1 | 1 | 1 | 1 | 1 | 8 |
| 1 | 1 | 1 | 1 | 1 | 8 |
| 1 | 1 | 1 | 1 | 1 | 8 |
| 1 | 1 | 1 | 1 | 1 | 8 |
| 1 | 1 | 1 | 1 | 1 | 8 |
| 1 | 1 | 1 | 1 | 1 | 8 |
| 1 | 1 | 1 | 1 | 1 | 8 |
| 1 | 1 | 1 | 1 | 1 | 8 |
| 1 | 1 | 1 | 1 | 1 | 8 |
| 1 | 1 | 1 | 1 | 1 | 8 |
| 1 | 1 | 1 | 1 | 1 | 8 |

| | | | | | |
|---|---|---|---|---|---|
| 1 | 1 | 1 | 1 | 1 | 8 |
| 1 | 1 | 1 | 1 | 1 | 8 |
| 1 | 1 | 1 | 1 | 1 | 8 |
| 1 | 1 | 1 | 1 | 1 | 8 |
| 1 | 1 | 1 | 1 | 1 | 8 |
| 1 | 1 | 1 | 1 | 1 | 8 |
| 1 | 1 | 1 | 1 | 1 | 8 |
| 1 | 1 | 1 | 1 | 1 | 8 |
| 1 | 1 | 1 | 1 | 1 | 8 |
| 1 | 1 | 1 | 1 | 1 | 8 |
| 0 | 1 | 0 | 0 | 0 | 2 |
| 0 | 1 | 0 | 0 | 0 | 2 |
| 0 | 1 | 0 | 0 | 0 | 2 |
| 0 | 1 | 0 | 0 | 0 | 2 |
| 0 | 1 | 0 | 0 | 0 | 2 |
| 0 | 1 | 0 | 0 | 0 | 2 |
| 0 | 1 | 0 | 0 | 0 | 2 |
| 0 | 1 | 0 | 0 | 0 | 2 |
| 0 | 1 | 0 | 0 | 0 | 2 |
| 0 | 1 | 0 | 0 | 0 | 2 |
| 0 | 1 | 0 | 0 | 0 | 2 |
| 0 | 1 | 0 | 0 | 0 | 2 |
| 0 | 1 | 0 | 0 | 0 | 2 |
| 0 | 1 | 0 | 0 | 0 | 2 |
| 0 | 1 | 0 | 0 | 0 | 2 |
| 0 | 1 | 0 | 0 | 0 | 2 |
| 0 | 1 | 0 | 0 | 0 | 2 |
| 0 | 1 | 0 | 0 | 0 | 2 |
| 0 | 1 | 0 | 0 | 0 | 2 |
| 0 | 1 | 0 | 0 | 0 | 2 |
| 0 | 1 | 0 | 0 | 0 | 2 |
| 0 | 1 | 0 | 0 | 0 | 2 |
| 0 | 1 | 0 | 0 | 0 | 2 |
| 0 | 1 | 0 | 0 | 0 | 2 |
| 0 | 1 | 0 | 0 | 0 | 2 |
| 0 | 1 | 0 | 0 | 0 | 2 |
| 0 | 1 | 0 | 0 | 0 | 2 |
| 0 | 1 | 0 | 0 | 0 | 2 |
| 0 | 1 | 0 | 0 | 0 | 2 |
| 0 | 1 | 0 | 0 | 0 | 2 |
| 0 | 1 | 0 | 0 | 0 | 2 |
| 0 | 1 | 0 | 0 | 0 | 2 |
| 0 | 1 | 0 | 0 | 0 | 2 |
| 0 | 1 | 0 | 0 | 0 | 2 |

| | | | | | |
|---|---|---|---|---|---|
| 0 | 1 | 0 | 0 | 0 | 2 |
| 0 | 1 | 0 | 0 | 0 | 2 |
| 0 | 1 | 0 | 0 | 0 | 2 |
| 0 | 1 | 0 | 0 | 0 | 2 |
| 0 | 1 | 0 | 0 | 0 | 2 |
| 0 | 1 | 0 | 0 | 0 | 2 |
| 1 | 1 | 1 | 1 | 1 | 8 |
| 1 | 1 | 1 | 1 | 1 | 8 |
| 1 | 1 | 1 | 1 | 1 | 8 |
| 1 | 1 | 1 | 1 | 1 | 8 |
| 1 | 1 | 1 | 1 | 1 | 8 |
| 1 | 1 | 1 | 1 | 1 | 8 |
| 1 | 1 | 1 | 1 | 1 | 8 |
| 1 | 1 | 1 | 1 | 1 | 8 |
| 1 | 1 | 1 | 1 | 1 | 8 |
| 1 | 1 | 1 | 1 | 1 | 8 |
| 1 | 1 | 1 | 1 | 1 | 8 |
| 1 | 1 | 1 | 1 | 1 | 8 |
| 1 | 1 | 1 | 1 | 1 | 8 |
| 1 | 1 | 1 | 1 | 1 | 8 |
| 1 | 1 | 1 | 1 | 1 | 8 |
| 1 | 1 | 1 | 1 | 1 | 8 |
| 1 | 1 | 1 | 1 | 1 | 8 |
| 1 | 1 | 1 | 1 | 1 | 8 |
| 1 | 1 | 1 | 1 | 1 | 8 |
| 1 | 1 | 1 | 1 | 1 | 8 |
| 1 | 1 | 1 | 1 | 1 | 8 |
| 1 | 1 | 1 | 1 | 1 | 8 |
| 1 | 1 | 1 | 1 | 1 | 8 |
| 1 | 1 | 1 | 1 | 1 | 8 |
| 1 | 1 | 1 | 1 | 1 | 8 |
| 1 | 1 | 1 | 1 | 1 | 8 |
| 1 | 1 | 1 | 1 | 1 | 8 |
| 1 | 1 | 1 | 1 | 1 | 8 |
| 1 | 1 | 1 | 1 | 1 | 8 |
| 1 | 1 | 1 | 1 | 1 | 8 |
| 1 | 1 | 1 | 1 | 1 | 8 |
| 1 | 1 | 1 | 1 | 1 | 8 |
| 1 | 1 | 1 | 1 | 1 | 8 |
| 1 | 1 | 1 | 1 | 1 | 8 |
| 1 | 1 | 1 | 1 | 1 | 8 |
| 1 | 1 | 1 | 1 | 1 | 8 |
| 1 | 1 | 1 | 1 | 1 | 8 |
| 1 | 1 | 1 | 1 | 1 | 8 |

| | | | | | |
|---|---|---|---|---|---|
| 1 | 1 | 1 | 1 | 1 | 8 |
| 1 | 1 | 1 | 1 | 1 | 8 |
| 1 | 1 | 1 | 1 | 1 | 8 |
| 1 | 1 | 1 | 1 | 1 | 8 |
| 1 | 1 | 1 | 1 | 1 | 8 |
| 1 | 1 | 1 | 1 | 1 | 8 |
| 1 | 1 | 1 | 1 | 1 | 8 |
| 1 | 1 | 1 | 1 | 1 | 8 |
| 1 | 1 | 1 | 1 | 1 | 8 |
| 1 | 1 | 1 | 1 | 1 | 8 |
| 1 | 1 | 1 | 1 | 1 | 8 |
| 1 | 1 | 1 | 1 | 1 | 8 |
| 1 | 1 | 1 | 1 | 1 | 8 |
| 1 | 1 | 1 | 1 | 1 | 8 |
| 1 | 1 | 1 | 1 | 1 | 8 |
| 1 | 1 | 1 | 1 | 1 | 8 |
| 1 | 1 | 1 | 1 | 1 | 8 |
| 1 | 1 | 1 | 1 | 1 | 8 |
| 1 | 1 | 1 | 1 | 1 | 8 |
| 1 | 1 | 1 | 1 | 1 | 8 |
| 1 | 1 | 1 | 1 | 1 | 8 |
| 1 | 1 | 1 | 1 | 1 | 8 |
| 1 | 1 | 1 | 1 | 1 | 8 |
| 1 | 1 | 1 | 1 | 1 | 8 |
| 1 | 1 | 1 | 1 | 1 | 8 |
| 1 | 1 | 1 | 1 | 1 | 8 |
| 1 | 1 | 1 | 1 | 1 | 8 |
| 1 | 1 | 1 | 1 | 1 | 8 |
| 1 | 1 | 1 | 1 | 1 | 8 |
| 1 | 1 | 1 | 1 | 1 | 8 |
| 1 | 1 | 1 | 1 | 1 | 8 |
| 1 | 1 | 1 | 1 | 1 | 8 |
| 1 | 1 | 1 | 1 | 1 | 8 |
| 1 | 1 | 1 | 1 | 1 | 8 |
| 1 | 1 | 1 | 1 | 1 | 8 |
| 1 | 1 | 1 | 1 | 1 | 8 |
| 1 | 1 | 1 | 1 | 1 | 8 |
| 1 | 1 | 1 | 1 | 1 | 8 |
| 1 | 1 | 1 | 1 | 1 | 8 |
| 1 | 1 | 1 | 1 | 1 | 8 |
| 1 | 1 | 1 | 1 | 1 | 8 |
| 1 | 1 | 1 | 1 | 1 | 8 |
| 1 | 1 | 1 | 1 | 1 | 8 |
| 1 | 1 | 1 | 1 | 1 | 8 |

| | | | | | |
|---|---|---|---|---|---|
| 1 | 1 | 1 | 1 | 1 | 8 |
| 1 | 1 | 1 | 1 | 1 | 8 |
| 1 | 1 | 1 | 1 | 1 | 8 |
| 1 | 1 | 1 | 1 | 1 | 8 |
| 1 | 1 | 1 | 1 | 1 | 8 |
| 1 | 1 | 1 | 1 | 1 | 8 |
| 1 | 1 | 1 | 1 | 1 | 8 |
| 1 | 1 | 1 | 1 | 1 | 8 |
| 1 | 1 | 1 | 1 | 1 | 8 |
| 1 | 1 | 1 | 1 | 1 | 8 |
| 1 | 1 | 1 | 1 | 1 | 8 |
| 1 | 1 | 1 | 1 | 1 | 8 |
| 1 | 1 | 1 | 1 | 1 | 8 |
| 1 | 1 | 1 | 1 | 1 | 8 |
| 1 | 1 | 1 | 1 | 1 | 8 |
| 1 | 1 | 1 | 1 | 1 | 8 |
| 1 | 1 | 1 | 1 | 1 | 8 |
| 1 | 1 | 1 | 1 | 1 | 8 |
| 1 | 1 | 1 | 1 | 1 | 8 |
| 1 | 1 | 1 | 1 | 1 | 8 |
| 1 | 1 | 1 | 1 | 1 | 8 |
| 1 | 1 | 1 | 1 | 1 | 8 |
| 1 | 1 | 1 | 1 | 1 | 8 |
| 1 | 1 | 1 | 1 | 1 | 8 |
| 1 | 1 | 1 | 1 | 1 | 8 |
| 1 | 1 | 1 | 1 | 1 | 8 |
| 1 | 1 | 1 | 1 | 1 | 8 |
| 1 | 1 | 1 | 1 | 1 | 8 |
| 1 | 1 | 1 | 1 | 1 | 8 |
| 1 | 1 | 1 | 1 | 1 | 8 |
| 1 | 1 | 1 | 1 | 1 | 8 |
| 1 | 1 | 1 | 1 | 1 | 8 |
| 1 | 1 | 1 | 1 | 1 | 8 |
| 1 | 1 | 1 | 1 | 1 | 8 |
| 1 | 1 | 1 | 1 | 1 | 8 |
| 1 | 1 | 1 | 1 | 1 | 8 |
| 1 | 1 | 1 | 1 | 1 | 8 |
| 1 | 1 | 1 | 1 | 1 | 8 |
| 1 | 1 | 1 | 1 | 1 | 8 |
| 1 | 1 | 1 | 1 | 1 | 8 |
| 1 | 1 | 1 | 1 | 1 | 8 |
| 1 | 1 | 1 | 1 | 1 | 8 |
| 1 | 1 | 1 | 1 | 1 | 8 |
| 1 | 1 | 1 | 1 | 1 | 8 |

| | | | | | |
|---|---|---|---|---|---|
| 1 | 1 | 1 | 1 | 1 | 8 |
| 1 | 1 | 1 | 1 | 1 | 8 |
| 1 | 1 | 1 | 1 | 1 | 8 |
| 1 | 1 | 1 | 1 | 1 | 8 |
| 1 | 1 | 1 | 1 | 1 | 8 |
| 1 | 1 | 1 | 1 | 1 | 8 |
| 1 | 1 | 1 | 1 | 1 | 8 |
| 1 | 1 | 1 | 1 | 1 | 8 |
| 1 | 1 | 1 | 1 | 1 | 8 |
| 1 | 1 | 1 | 1 | 1 | 8 |
| 1 | 1 | 1 | 1 | 1 | 8 |
| 1 | 1 | 1 | 1 | 1 | 8 |
| 1 | 1 | 1 | 1 | 1 | 8 |
| 1 | 1 | 1 | 1 | 1 | 8 |
| 1 | 1 | 1 | 1 | 1 | 8 |
| 1 | 1 | 1 | 1 | 1 | 8 |
| 1 | 1 | 1 | 1 | 1 | 8 |
| 1 | 1 | 1 | 1 | 1 | 8 |
| 1 | 1 | 1 | 1 | 1 | 8 |
| 1 | 1 | 1 | 1 | 1 | 8 |
| 1 | 1 | 1 | 1 | 1 | 8 |
| 1 | 1 | 1 | 1 | 1 | 8 |
| 1 | 1 | 1 | 1 | 1 | 8 |
| 1 | 1 | 1 | 1 | 1 | 8 |
| 1 | 1 | 1 | 1 | 1 | 8 |
| 1 | 1 | 1 | 1 | 1 | 8 |
| 1 | 1 | 1 | 1 | 1 | 8 |
| 1 | 1 | 1 | 1 | 1 | 8 |
| 1 | 1 | 1 | 1 | 1 | 8 |
| 1 | 1 | 1 | 1 | 1 | 8 |
| 1 | 1 | 1 | 1 | 1 | 8 |
| 1 | 1 | 1 | 1 | 1 | 8 |
| 1 | 1 | 1 | 1 | 1 | 8 |
| 1 | 1 | 1 | 1 | 1 | 8 |
| 1 | 1 | 1 | 1 | 1 | 8 |
| 1 | 1 | 1 | 1 | 1 | 8 |
| 1 | 1 | 1 | 1 | 1 | 8 |
| 1 | 1 | 1 | 1 | 1 | 8 |
| 1 | 1 | 1 | 1 | 1 | 8 |
| 1 | 1 | 1 | 1 | 1 | 8 |
| 1 | 1 | 1 | 1 | 1 | 8 |
| 1 | 1 | 1 | 1 | 1 | 8 |
| 1 | 1 | 1 | 1 | 1 | 8 |
| 1 | 1 | 1 | 1 | 1 | 8 |

| | | | | | |
|---|---|---|---|---|---|
| 1 | 1 | 1 | 1 | 1 | 8 |
| 1 | 1 | 1 | 1 | 1 | 8 |
| 1 | 1 | 1 | 1 | 1 | 8 |
| 1 | 1 | 1 | 1 | 1 | 8 |
| 1 | 1 | 1 | 1 | 1 | 8 |
| 1 | 1 | 1 | 1 | 1 | 8 |
| 1 | 1 | 1 | 1 | 1 | 8 |
| 1 | 1 | 1 | 1 | 1 | 8 |
| 1 | 1 | 1 | 1 | 1 | 8 |
| 1 | 1 | 1 | 1 | 1 | 8 |
| 1 | 1 | 1 | 1 | 1 | 8 |
| 1 | 1 | 1 | 1 | 1 | 8 |
| 1 | 1 | 1 | 1 | 1 | 8 |
| 1 | 1 | 1 | 1 | 1 | 8 |
| 1 | 1 | 1 | 1 | 1 | 8 |
| 1 | 1 | 1 | 1 | 1 | 8 |
| 1 | 1 | 1 | 1 | 1 | 8 |
| 1 | 1 | 1 | 1 | 1 | 8 |
| 1 | 1 | 1 | 1 | 1 | 8 |
| 1 | 1 | 1 | 1 | 1 | 8 |
| 1 | 1 | 1 | 1 | 1 | 8 |
| 1 | 1 | 1 | 1 | 1 | 8 |
| 1 | 1 | 1 | 1 | 1 | 8 |
| 1 | 1 | 1 | 1 | 1 | 8 |
| 1 | 1 | 1 | 1 | 1 | 8 |
| 1 | 1 | 1 | 1 | 1 | 8 |
| 1 | 1 | 1 | 1 | 1 | 8 |
| 1 | 1 | 1 | 1 | 1 | 8 |
| 1 | 1 | 1 | 1 | 1 | 8 |
| 1 | 1 | 1 | 1 | 1 | 8 |
| 1 | 1 | 1 | 1 | 1 | 8 |
| 1 | 1 | 1 | 1 | 1 | 8 |
| 1 | 1 | 1 | 1 | 1 | 8 |
| 1 | 1 | 1 | 1 | 1 | 8 |
| 1 | 1 | 1 | 1 | 1 | 8 |
| 1 | 1 | 1 | 1 | 1 | 8 |
| 1 | 1 | 1 | 1 | 1 | 8 |
| 1 | 1 | 1 | 1 | 1 | 8 |
| 1 | 1 | 1 | 1 | 1 | 8 |
| 1 | 1 | 1 | 1 | 1 | 8 |
| 1 | 1 | 1 | 1 | 1 | 8 |
| 1 | 1 | 1 | 1 | 1 | 8 |
| 1 | 1 | 1 | 1 | 1 | 8 |
| 1 | 1 | 1 | 1 | 1 | 8 |

| | | | | | |
|---|---|---|---|---|---|
| 1 | 1 | 1 | 1 | 1 | 8 |
| 1 | 1 | 1 | 1 | 1 | 8 |
| 1 | 1 | 1 | 1 | 1 | 8 |
| 1 | 1 | 1 | 1 | 1 | 8 |
| 1 | 1 | 1 | 1 | 1 | 8 |
| 1 | 1 | 1 | 1 | 1 | 8 |
| 1 | 1 | 1 | 1 | 1 | 8 |
| 1 | 1 | 1 | 1 | 1 | 8 |
| 1 | 1 | 1 | 1 | 1 | 8 |
| 1 | 1 | 1 | 1 | 1 | 8 |
| 1 | 1 | 1 | 1 | 1 | 8 |
| 1 | 1 | 1 | 1 | 1 | 8 |
| 1 | 1 | 1 | 1 | 1 | 8 |
| 1 | 1 | 1 | 1 | 1 | 8 |
| 1 | 1 | 1 | 1 | 1 | 8 |
| 1 | 1 | 1 | 1 | 1 | 8 |
| 1 | 1 | 1 | 1 | 1 | 8 |
| 1 | 1 | 1 | 1 | 1 | 8 |
| 1 | 1 | 1 | 1 | 1 | 8 |
| 1 | 1 | 1 | 1 | 1 | 8 |
| 1 | 1 | 1 | 1 | 1 | 8 |
| 1 | 1 | 1 | 1 | 1 | 8 |
| 1 | 1 | 1 | 1 | 1 | 8 |
| 1 | 1 | 1 | 1 | 1 | 8 |
| 1 | 1 | 1 | 1 | 1 | 8 |
| 1 | 1 | 1 | 1 | 1 | 8 |
| 1 | 1 | 1 | 1 | 1 | 8 |
| 1 | 1 | 1 | 1 | 1 | 8 |
| 1 | 1 | 1 | 1 | 1 | 8 |
| 1 | 1 | 1 | 1 | 1 | 8 |
| 1 | 1 | 1 | 1 | 1 | 8 |
| 1 | 1 | 1 | 1 | 1 | 8 |
| 1 | 1 | 1 | 1 | 1 | 8 |
| 1 | 1 | 1 | 1 | 1 | 8 |
| 1 | 1 | 1 | 1 | 1 | 8 |
| 1 | 1 | 1 | 1 | 1 | 8 |
| 1 | 1 | 1 | 1 | 1 | 8 |
| 1 | 1 | 1 | 1 | 1 | 8 |
| 1 | 1 | 1 | 1 | 1 | 8 |
| 1 | 1 | 1 | 1 | 1 | 8 |
| 1 | 1 | 1 | 1 | 1 | 8 |
| 1 | 1 | 1 | 1 | 1 | 8 |
| 1 | 1 | 1 | 1 | 1 | 8 |
| 1 | 1 | 1 | 1 | 1 | 8 |

| | | | | | |
|---|---|---|---|---|---|
| 1 | 1 | 1 | 1 | 1 | 8 |
| 1 | 1 | 1 | 1 | 1 | 8 |
| 1 | 1 | 1 | 1 | 1 | 8 |
| 1 | 1 | 1 | 1 | 1 | 8 |
| 1 | 1 | 1 | 1 | 1 | 8 |
| 1 | 1 | 1 | 1 | 1 | 8 |
| 1 | 1 | 1 | 1 | 1 | 8 |
| 1 | 1 | 1 | 1 | 1 | 8 |
| 1 | 1 | 1 | 1 | 1 | 8 |
| 1 | 1 | 1 | 1 | 1 | 8 |
| 1 | 1 | 1 | 1 | 1 | 8 |
| 1 | 1 | 1 | 1 | 1 | 8 |
| 1 | 1 | 1 | 1 | 1 | 8 |
| 1 | 1 | 1 | 1 | 1 | 8 |
| 1 | 1 | 1 | 1 | 1 | 8 |
| 1 | 1 | 1 | 1 | 1 | 8 |
| 1 | 1 | 1 | 1 | 1 | 8 |
| 1 | 1 | 1 | 1 | 1 | 8 |
| 1 | 1 | 1 | 1 | 1 | 8 |
| 1 | 1 | 1 | 1 | 1 | 8 |
| 1 | 1 | 1 | 1 | 1 | 8 |
| 1 | 1 | 1 | 1 | 1 | 8 |
| 1 | 1 | 1 | 1 | 1 | 8 |
| 1 | 1 | 1 | 1 | 1 | 8 |
| 1 | 1 | 1 | 1 | 1 | 8 |
| 1 | 1 | 1 | 1 | 1 | 8 |
| 1 | 1 | 1 | 1 | 1 | 8 |
| 1 | 1 | 1 | 1 | 1 | 8 |
| 1 | 1 | 1 | 1 | 1 | 8 |
| 1 | 1 | 1 | 1 | 1 | 8 |
| 1 | 1 | 1 | 1 | 1 | 8 |
| 1 | 1 | 1 | 1 | 1 | 8 |
| 1 | 1 | 1 | 1 | 1 | 8 |
| 1 | 1 | 1 | 1 | 1 | 8 |
| 1 | 1 | 1 | 1 | 1 | 8 |
| 1 | 1 | 1 | 1 | 1 | 8 |
| 1 | 1 | 1 | 1 | 1 | 8 |
| 1 | 1 | 1 | 1 | 1 | 8 |
| 1 | 1 | 1 | 1 | 1 | 8 |
| 1 | 1 | 1 | 1 | 1 | 8 |
| 1 | 1 | 1 | 1 | 1 | 8 |
| 1 | 1 | 1 | 1 | 1 | 8 |
| 1 | 1 | 1 | 1 | 1 | 8 |
| 1 | 1 | 1 | 1 | 1 | 8 |

| | | | | | |
|---|---|---|---|---|---|
| 1 | 1 | 1 | 1 | 1 | 8 |
| 1 | 1 | 1 | 1 | 1 | 8 |
| 1 | 1 | 1 | 1 | 1 | 8 |
| 1 | 1 | 1 | 1 | 1 | 8 |
| 1 | 1 | 1 | 1 | 1 | 8 |
| 1 | 1 | 1 | 1 | 1 | 8 |
| 1 | 1 | 1 | 1 | 1 | 8 |
| 1 | 1 | 1 | 1 | 1 | 8 |
| 1 | 1 | 1 | 1 | 1 | 8 |
| 1 | 1 | 1 | 1 | 1 | 8 |
| 1 | 1 | 1 | 1 | 1 | 8 |
| 1 | 1 | 1 | 1 | 1 | 8 |
| 1 | 1 | 1 | 1 | 1 | 8 |
| 1 | 1 | 1 | 1 | 1 | 8 |
| 1 | 1 | 1 | 1 | 1 | 8 |
| 1 | 1 | 1 | 1 | 1 | 8 |
| 1 | 1 | 1 | 1 | 1 | 8 |
| 1 | 1 | 1 | 1 | 1 | 8 |
| 1 | 1 | 1 | 1 | 1 | 8 |
| 1 | 1 | 1 | 1 | 1 | 8 |
| 1 | 1 | 1 | 1 | 1 | 8 |
| 1 | 1 | 1 | 1 | 1 | 8 |
| 1 | 1 | 1 | 1 | 1 | 8 |
| 1 | 1 | 1 | 1 | 1 | 8 |
| 1 | 1 | 1 | 1 | 1 | 8 |
| 1 | 1 | 1 | 1 | 1 | 8 |
| 1 | 1 | 1 | 1 | 1 | 8 |
| 1 | 1 | 1 | 1 | 1 | 8 |
| 1 | 1 | 1 | 1 | 1 | 8 |
| 1 | 1 | 1 | 1 | 1 | 8 |
| 1 | 1 | 1 | 1 | 1 | 8 |
| 1 | 1 | 1 | 1 | 1 | 8 |
| 1 | 1 | 1 | 1 | 1 | 8 |
| 1 | 1 | 1 | 1 | 1 | 8 |
| 1 | 1 | 1 | 1 | 1 | 8 |
| 1 | 1 | 1 | 1 | 1 | 8 |
| 1 | 1 | 1 | 1 | 1 | 8 |
| 1 | 1 | 1 | 1 | 1 | 8 |
| 1 | 1 | 1 | 1 | 1 | 8 |
| 1 | 1 | 1 | 1 | 1 | 8 |
| 1 | 1 | 1 | 1 | 1 | 8 |
| 1 | 1 | 1 | 1 | 1 | 8 |
| 1 | 1 | 1 | 1 | 1 | 8 |
| 1 | 1 | 1 | 1 | 1 | 8 |

| | | | | | |
|---|---|---|---|---|---|
| 1 | 1 | 1 | 1 | 1 | 8 |
| 1 | 1 | 1 | 1 | 1 | 8 |
| 1 | 1 | 1 | 1 | 1 | 8 |
| 1 | 1 | 1 | 1 | 1 | 8 |
| 1 | 1 | 1 | 1 | 1 | 8 |
| 1 | 1 | 1 | 1 | 1 | 8 |
| 1 | 1 | 1 | 1 | 1 | 8 |
| 1 | 1 | 1 | 1 | 1 | 8 |
| 1 | 1 | 1 | 1 | 1 | 8 |
| 1 | 1 | 1 | 1 | 1 | 8 |
| 1 | 1 | 1 | 1 | 1 | 8 |
| 1 | 1 | 1 | 1 | 1 | 8 |
| 1 | 1 | 1 | 1 | 1 | 8 |
| 1 | 1 | 1 | 1 | 1 | 8 |
| 1 | 1 | 1 | 1 | 1 | 8 |
| 1 | 1 | 1 | 1 | 1 | 8 |
| 1 | 1 | 1 | 1 | 1 | 8 |
| 1 | 1 | 1 | 1 | 1 | 8 |
| 1 | 1 | 1 | 1 | 1 | 8 |
| 1 | 1 | 1 | 1 | 1 | 8 |
| 1 | 1 | 1 | 1 | 1 | 8 |
| 1 | 1 | 1 | 1 | 1 | 8 |
| 1 | 1 | 1 | 1 | 1 | 8 |
| 1 | 1 | 1 | 1 | 1 | 8 |
| 1 | 1 | 1 | 1 | 1 | 8 |
| 1 | 1 | 1 | 1 | 1 | 8 |
| 1 | 1 | 1 | 1 | 1 | 8 |
| 1 | 1 | 1 | 1 | 1 | 8 |
| 1 | 1 | 1 | 1 | 1 | 8 |
| 1 | 1 | 1 | 1 | 1 | 8 |
| 1 | 1 | 1 | 1 | 1 | 8 |
| 1 | 1 | 1 | 1 | 1 | 8 |
| 1 | 1 | 1 | 1 | 1 | 8 |
| 1 | 1 | 1 | 1 | 1 | 8 |
| 1 | 1 | 1 | 1 | 1 | 8 |
| 1 | 1 | 1 | 1 | 1 | 8 |
| 1 | 1 | 1 | 1 | 1 | 8 |
| 1 | 1 | 1 | 1 | 1 | 8 |
| 1 | 1 | 1 | 1 | 1 | 8 |
| 1 | 1 | 1 | 1 | 1 | 8 |
| 1 | 1 | 1 | 1 | 1 | 8 |
| 1 | 1 | 1 | 1 | 1 | 8 |
| 1 | 1 | 1 | 1 | 1 | 8 |
| 1 | 1 | 1 | 1 | 1 | 8 |

| | | | | | |
|---|---|---|---|---|---|
| 1 | 1 | 1 | 1 | 1 | 8 |
| 1 | 1 | 1 | 1 | 1 | 8 |
| 1 | 1 | 1 | 1 | 1 | 8 |
| 1 | 1 | 1 | 1 | 1 | 8 |
| 1 | 1 | 1 | 1 | 1 | 8 |
| 1 | 1 | 1 | 1 | 1 | 8 |
| 1 | 1 | 1 | 1 | 1 | 8 |
| 1 | 1 | 1 | 1 | 1 | 8 |
| 1 | 1 | 1 | 1 | 1 | 8 |
| 1 | 1 | 1 | 1 | 1 | 8 |
| 1 | 1 | 1 | 1 | 1 | 8 |
| 1 | 1 | 1 | 1 | 1 | 8 |
| 1 | 1 | 1 | 1 | 1 | 8 |
| 1 | 1 | 1 | 1 | 1 | 8 |
| 1 | 1 | 1 | 1 | 1 | 8 |
| 1 | 1 | 1 | 1 | 1 | 8 |
| 1 | 1 | 1 | 1 | 1 | 8 |
| 1 | 1 | 1 | 1 | 1 | 8 |
| 1 | 1 | 1 | 1 | 1 | 8 |
| 1 | 1 | 1 | 1 | 1 | 8 |
| 1 | 1 | 1 | 1 | 1 | 8 |
| 1 | 1 | 1 | 1 | 1 | 8 |
| 1 | 1 | 1 | 1 | 1 | 8 |
| 1 | 1 | 1 | 1 | 1 | 8 |
| 1 | 1 | 1 | 1 | 1 | 8 |
| 1 | 1 | 1 | 1 | 1 | 8 |
| 1 | 1 | 1 | 1 | 1 | 8 |
| 1 | 1 | 1 | 1 | 1 | 8 |
| 1 | 1 | 1 | 1 | 1 | 8 |
| 1 | 1 | 1 | 1 | 1 | 8 |
| 1 | 1 | 1 | 1 | 1 | 8 |
| 1 | 1 | 1 | 1 | 1 | 8 |
| 1 | 1 | 1 | 1 | 1 | 8 |
| 1 | 1 | 1 | 1 | 1 | 8 |
| 1 | 1 | 1 | 1 | 1 | 8 |
| 1 | 1 | 1 | 1 | 1 | 8 |
| 1 | 1 | 1 | 1 | 1 | 8 |
| 1 | 1 | 1 | 1 | 1 | 8 |
| 1 | 1 | 1 | 1 | 1 | 8 |
| 1 | 1 | 1 | 1 | 1 | 8 |
| 1 | 1 | 1 | 1 | 1 | 8 |
| 1 | 1 | 1 | 1 | 1 | 8 |
| 1 | 1 | 1 | 1 | 1 | 8 |
| 1 | 1 | 1 | 1 | 1 | 8 |

| | | | | | |
|---|---|---|---|---|---|
| 1 | 1 | 1 | 1 | 1 | 8 |
| 1 | 1 | 1 | 1 | 1 | 8 |
| 1 | 1 | 1 | 1 | 1 | 8 |
| 1 | 1 | 1 | 1 | 1 | 8 |
| 1 | 1 | 1 | 1 | 1 | 8 |
| 1 | 1 | 1 | 1 | 1 | 8 |
| 1 | 1 | 1 | 1 | 1 | 8 |
| 1 | 1 | 1 | 1 | 1 | 8 |
| 1 | 1 | 1 | 1 | 1 | 8 |
| 1 | 1 | 1 | 1 | 1 | 8 |
| 1 | 1 | 1 | 1 | 1 | 8 |
| 1 | 1 | 1 | 1 | 1 | 8 |
| 1 | 1 | 1 | 1 | 1 | 8 |
| 1 | 1 | 1 | 1 | 1 | 8 |
| 1 | 1 | 1 | 1 | 1 | 8 |
| 1 | 1 | 1 | 1 | 1 | 8 |
| 1 | 1 | 1 | 1 | 1 | 8 |
| 1 | 1 | 1 | 1 | 1 | 8 |
| 1 | 1 | 1 | 1 | 1 | 8 |
| 1 | 1 | 1 | 1 | 1 | 8 |
| 1 | 1 | 1 | 1 | 1 | 8 |
| 1 | 1 | 1 | 1 | 1 | 8 |
| 1 | 1 | 1 | 1 | 1 | 8 |
| 1 | 1 | 1 | 1 | 1 | 8 |
| 1 | 1 | 1 | 1 | 1 | 8 |
| 1 | 1 | 1 | 1 | 1 | 8 |
| 1 | 1 | 1 | 1 | 1 | 8 |
| 1 | 1 | 1 | 1 | 1 | 8 |
| 1 | 1 | 1 | 1 | 1 | 8 |
| 1 | 1 | 1 | 1 | 1 | 8 |
| 1 | 1 | 1 | 1 | 1 | 8 |
| 1 | 1 | 1 | 1 | 1 | 8 |
| 1 | 1 | 1 | 1 | 1 | 8 |
| 1 | 1 | 1 | 1 | 1 | 8 |
| 1 | 1 | 1 | 1 | 1 | 8 |
| 1 | 1 | 1 | 1 | 1 | 8 |
| 1 | 1 | 1 | 1 | 1 | 8 |
| 1 | 1 | 1 | 1 | 1 | 8 |
| 1 | 1 | 1 | 1 | 1 | 8 |
| 1 | 1 | 1 | 1 | 1 | 8 |
| 1 | 1 | 1 | 1 | 1 | 8 |
| 1 | 1 | 1 | 1 | 1 | 8 |
| 1 | 1 | 1 | 1 | 1 | 8 |
| 1 | 1 | 1 | 1 | 1 | 8 |

| | | | | | |
|---|---|---|---|---|---|
| 1 | 1 | 1 | 1 | 1 | 8 |
| 1 | 1 | 1 | 1 | 1 | 8 |
| 1 | 1 | 1 | 1 | 1 | 8 |
| 1 | 1 | 1 | 1 | 1 | 8 |
| 1 | 1 | 1 | 1 | 1 | 8 |
| 1 | 1 | 1 | 1 | 1 | 8 |
| 1 | 1 | 1 | 1 | 1 | 8 |
| 1 | 1 | 1 | 1 | 1 | 8 |
| 1 | 1 | 1 | 1 | 1 | 8 |
| 1 | 1 | 1 | 1 | 1 | 8 |
| 1 | 1 | 1 | 1 | 1 | 8 |
| 1 | 1 | 1 | 1 | 1 | 8 |
| 1 | 1 | 1 | 1 | 1 | 8 |
| 1 | 1 | 1 | 1 | 1 | 8 |
| 1 | 1 | 1 | 1 | 1 | 8 |
| 1 | 1 | 1 | 1 | 1 | 8 |
| 1 | 1 | 1 | 1 | 1 | 8 |
| 1 | 1 | 1 | 1 | 1 | 8 |
| 1 | 1 | 1 | 1 | 1 | 8 |
| 1 | 1 | 1 | 1 | 1 | 8 |
| 1 | 1 | 1 | 1 | 1 | 8 |
| 1 | 1 | 1 | 1 | 1 | 8 |
| 1 | 1 | 1 | 1 | 1 | 8 |
| 1 | 1 | 1 | 1 | 1 | 8 |
| 1 | 1 | 1 | 1 | 1 | 8 |
| 1 | 1 | 1 | 1 | 1 | 8 |
| 1 | 1 | 1 | 1 | 1 | 8 |
| 1 | 1 | 1 | 1 | 1 | 8 |
| 1 | 1 | 1 | 1 | 1 | 8 |
| 1 | 1 | 1 | 1 | 1 | 8 |
| 1 | 1 | 1 | 1 | 1 | 8 |
| 1 | 1 | 1 | 1 | 1 | 8 |
| 1 | 1 | 1 | 1 | 1 | 8 |
| 1 | 1 | 1 | 1 | 1 | 8 |
| 1 | 1 | 1 | 1 | 1 | 8 |
| 1 | 1 | 1 | 1 | 1 | 8 |
| 1 | 1 | 1 | 1 | 1 | 8 |
| 1 | 1 | 1 | 1 | 1 | 8 |
| 1 | 1 | 1 | 1 | 1 | 8 |
| 1 | 1 | 1 | 1 | 1 | 8 |
| 1 | 1 | 1 | 1 | 1 | 8 |
| 1 | 1 | 1 | 1 | 1 | 8 |
| 1 | 1 | 1 | 1 | 1 | 8 |
| 1 | 1 | 1 | 1 | 1 | 8 |

| | | | | | |
|---|---|---|---|---|---|
| 1 | 1 | 1 | 1 | 1 | 8 |
| 1 | 1 | 1 | 1 | 1 | 8 |
| 1 | 1 | 1 | 1 | 1 | 8 |
| 1 | 1 | 1 | 1 | 1 | 8 |
| 1 | 1 | 1 | 1 | 1 | 8 |
| 1 | 1 | 1 | 1 | 1 | 8 |
| 1 | 1 | 1 | 1 | 1 | 8 |
| 1 | 1 | 1 | 1 | 1 | 8 |
| 1 | 1 | 1 | 1 | 1 | 8 |
| 1 | 1 | 1 | 1 | 1 | 8 |
| 1 | 1 | 1 | 1 | 1 | 8 |
| 1 | 1 | 1 | 1 | 1 | 8 |
| 1 | 1 | 1 | 1 | 1 | 8 |
| 1 | 1 | 1 | 1 | 1 | 8 |
| 1 | 1 | 1 | 1 | 1 | 8 |
| 1 | 1 | 1 | 1 | 1 | 8 |
| 1 | 1 | 1 | 1 | 1 | 8 |
| 1 | 1 | 1 | 1 | 1 | 8 |
| 1 | 1 | 1 | 1 | 1 | 8 |
| 1 | 1 | 1 | 1 | 1 | 8 |
| 1 | 1 | 1 | 1 | 1 | 8 |
| 1 | 1 | 1 | 1 | 1 | 8 |
| 1 | 1 | 1 | 1 | 1 | 8 |
| 1 | 1 | 1 | 1 | 1 | 8 |
| 1 | 1 | 1 | 1 | 1 | 8 |
| 1 | 1 | 1 | 1 | 1 | 8 |
| 1 | 1 | 1 | 1 | 1 | 8 |
| 1 | 1 | 1 | 1 | 1 | 8 |
| 1 | 1 | 1 | 1 | 1 | 8 |
| 1 | 1 | 1 | 1 | 1 | 8 |
| 1 | 1 | 1 | 1 | 1 | 8 |
| 1 | 1 | 1 | 1 | 1 | 8 |
| 1 | 1 | 1 | 1 | 1 | 8 |
| 1 | 1 | 1 | 1 | 1 | 8 |
| 1 | 1 | 1 | 1 | 1 | 8 |
| 1 | 1 | 1 | 1 | 1 | 8 |
| 1 | 1 | 1 | 1 | 1 | 8 |
| 1 | 1 | 1 | 1 | 1 | 8 |
| 1 | 1 | 1 | 1 | 1 | 8 |
| 1 | 1 | 1 | 1 | 1 | 8 |
| 1 | 1 | 1 | 1 | 1 | 8 |
| 1 | 1 | 1 | 1 | 1 | 8 |
| 0 | 1 | 1 | 1 | 0 | 4 |
| 0 | 1 | 1 | 1 | 0 | 4 |

| | | | | | |
|---|---|---|---|---|---|
| 0 | 1 | 1 | 1 | 0 | 4 |
| 0 | 1 | 1 | 1 | 0 | 4 |
| 0 | 1 | 1 | 1 | 0 | 4 |
| 0 | 1 | 1 | 1 | 0 | 4 |
| 0 | 1 | 1 | 1 | 0 | 4 |
| 0 | 1 | 1 | 1 | 0 | 4 |
| 0 | 1 | 1 | 1 | 0 | 4 |
| 0 | 1 | 1 | 1 | 0 | 4 |
| 0 | 1 | 1 | 1 | 0 | 4 |
| 0 | 1 | 1 | 1 | 0 | 4 |
| 0 | 1 | 1 | 1 | 0 | 4 |
| 0 | 1 | 1 | 1 | 0 | 4 |
| 0 | 1 | 1 | 1 | 0 | 4 |
| 0 | 1 | 1 | 1 | 0 | 4 |
| 0 | 1 | 1 | 1 | 0 | 4 |
| 0 | 1 | 1 | 1 | 0 | 4 |
| 0 | 1 | 1 | 1 | 0 | 4 |
| 0 | 1 | 1 | 1 | 0 | 4 |
| 0 | 1 | 1 | 1 | 0 | 4 |
| 0 | 1 | 1 | 1 | 0 | 4 |
| 0 | 1 | 1 | 1 | 0 | 4 |
| 0 | 1 | 1 | 1 | 0 | 4 |
| 0 | 1 | 1 | 1 | 0 | 4 |
| 0 | 1 | 1 | 1 | 0 | 4 |
| 0 | 1 | 1 | 1 | 0 | 4 |
| 0 | 1 | 1 | 1 | 0 | 4 |
| 0 | 1 | 1 | 1 | 0 | 4 |
| 0 | 1 | 1 | 1 | 0 | 4 |
| 0 | 1 | 1 | 1 | 0 | 4 |
| 0 | 1 | 1 | 1 | 0 | 4 |
| 0 | 1 | 1 | 1 | 0 | 4 |
| 0 | 1 | 1 | 1 | 0 | 4 |
| 0 | 1 | 1 | 1 | 0 | 4 |
| 0 | 1 | 1 | 1 | 0 | 4 |
| 0 | 1 | 1 | 1 | 0 | 4 |
| 0 | 1 | 1 | 1 | 0 | 4 |
| 0 | 1 | 1 | 1 | 0 | 4 |
| 0 | 1 | 1 | 1 | 0 | 4 |
| 0 | 1 | 1 | 1 | 0 | 4 |
| 0 | 1 | 1 | 1 | 0 | 4 |
| 0 | 1 | 1 | 1 | 0 | 4 |
| 0 | 1 | 1 | 1 | 0 | 4 |
| 0 | 1 | 1 | 1 | 0 | 4 |
| 0 | 1 | 1 | 1 | 0 | 4 |

| | | | | | |
|---|---|---|---|---|---|
| 0 | 1 | 1 | 1 | 0 | 4 |
| 0 | 1 | 1 | 1 | 0 | 4 |
| 0 | 1 | 1 | 1 | 0 | 4 |
| 0 | 1 | 1 | 1 | 0 | 4 |
| 0 | 1 | 1 | 1 | 0 | 4 |
| 0 | 1 | 1 | 1 | 0 | 4 |
| 0 | 1 | 1 | 1 | 0 | 4 |
| 0 | 1 | 1 | 1 | 0 | 4 |
| 0 | 1 | 1 | 1 | 0 | 4 |
| 0 | 1 | 1 | 1 | 0 | 4 |
| 0 | 1 | 1 | 1 | 1 | 5 |
| 0 | 1 | 1 | 1 | 1 | 5 |
| 0 | 1 | 1 | 1 | 1 | 5 |
| 0 | 1 | 1 | 1 | 1 | 5 |
| 0 | 1 | 1 | 1 | 1 | 5 |
| 0 | 1 | 1 | 1 | 1 | 5 |
| 0 | 1 | 1 | 1 | 1 | 5 |
| 0 | 1 | 1 | 1 | 1 | 5 |
| 0 | 1 | 1 | 1 | 1 | 5 |
| 0 | 1 | 1 | 1 | 1 | 5 |
| 0 | 1 | 1 | 1 | 1 | 5 |
| 0 | 1 | 1 | 1 | 1 | 5 |
| 0 | 1 | 1 | 1 | 1 | 5 |
| 0 | 1 | 1 | 1 | 1 | 5 |
| 0 | 1 | 1 | 1 | 1 | 5 |
| 0 | 1 | 1 | 1 | 1 | 5 |
| 0 | 1 | 1 | 1 | 1 | 5 |
| 0 | 1 | 1 | 1 | 1 | 5 |
| 0 | 1 | 1 | 1 | 1 | 5 |
| 0 | 1 | 1 | 1 | 1 | 5 |
| 0 | 1 | 1 | 1 | 1 | 5 |
| 0 | 1 | 1 | 1 | 1 | 5 |
| 0 | 1 | 1 | 1 | 1 | 5 |
| 0 | 1 | 1 | 1 | 1 | 5 |
| 0 | 1 | 1 | 1 | 1 | 5 |
| 0 | 1 | 1 | 1 | 1 | 5 |
| 0 | 1 | 1 | 1 | 1 | 5 |
| 0 | 1 | 1 | 1 | 1 | 5 |
| 0 | 1 | 1 | 1 | 1 | 5 |
| 0 | 1 | 1 | 1 | 1 | 5 |
| 0 | 1 | 1 | 1 | 1 | 5 |
| 0 | 1 | 1 | 1 | 1 | 5 |
| 0 | 1 | 1 | 1 | 1 | 5 |
| 0 | 1 | 1 | 1 | 1 | 5 |

| | | | | | |
|---|---|---|---|---|---|
| 0 | 1 | 1 | 1 | 1 | 5 |
| 0 | 1 | 1 | 1 | 1 | 5 |
| 0 | 1 | 1 | 1 | 1 | 5 |
| 0 | 1 | 1 | 1 | 1 | 5 |
| 0 | 1 | 1 | 1 | 1 | 5 |
| 0 | 1 | 1 | 1 | 1 | 5 |
| 0 | 1 | 1 | 1 | 1 | 5 |
| 0 | 1 | 1 | 1 | 1 | 5 |
| 0 | 1 | 1 | 1 | 1 | 5 |
| 0 | 1 | 1 | 1 | 1 | 5 |
| 0 | 1 | 1 | 1 | 1 | 5 |
| 0 | 1 | 1 | 1 | 1 | 5 |
| 0 | 1 | 1 | 1 | 1 | 5 |
| 0 | 1 | 1 | 1 | 1 | 5 |
| 0 | 1 | 1 | 1 | 1 | 5 |
| 0 | 1 | 1 | 1 | 1 | 5 |
| 0 | 1 | 1 | 1 | 1 | 5 |
| 0 | 1 | 1 | 1 | 1 | 5 |
| 0 | 1 | 1 | 1 | 1 | 5 |
| 0 | 1 | 1 | 1 | 1 | 5 |
| 0 | 1 | 1 | 1 | 1 | 5 |
| 0 | 1 | 1 | 1 | 1 | 5 |
| 1 | 1 | 1 | 1 | 1 | 8 |
| 1 | 1 | 1 | 1 | 1 | 8 |
| 1 | 1 | 1 | 1 | 1 | 8 |
| 1 | 1 | 1 | 1 | 1 | 8 |
| 1 | 1 | 1 | 1 | 1 | 8 |
| 1 | 1 | 1 | 1 | 1 | 8 |
| 1 | 1 | 1 | 1 | 1 | 8 |
| 1 | 1 | 1 | 1 | 1 | 8 |
| 0 | 1 | 0 | 0 | 0 | 2 |
| 0 | 1 | 0 | 0 | 0 | 2 |
| 0 | 1 | 0 | 0 | 0 | 2 |
| 0 | 1 | 0 | 0 | 0 | 2 |
| 0 | 1 | 0 | 0 | 0 | 2 |
| 0 | 1 | 0 | 0 | 0 | 2 |
| 0 | 1 | 0 | 0 | 0 | 2 |
| 0 | 1 | 0 | 0 | 0 | 2 |
| 1 | 1 | 1 | 1 | 1 | 8 |
| 1 | 1 | 1 | 1 | 1 | 8 |
| 1 | 1 | 1 | 1 | 1 | 8 |
| 1 | 1 | 1 | 1 | 1 | 8 |
| 1 | 1 | 1 | 1 | 1 | 8 |
| 1 | 1 | 1 | 1 | 1 | 8 |

| | | | | | |
|---|---|---|---|---|---|
| 1 | 1 | 1 | 1 | 1 | 8 |
| 1 | 1 | 1 | 1 | 1 | 8 |
| 1 | 1 | 1 | 1 | 1 | 8 |
| 1 | 1 | 1 | 1 | 1 | 8 |
| 1 | 1 | 1 | 1 | 1 | 8 |
| 1 | 1 | 1 | 1 | 1 | 8 |
| 1 | 1 | 1 | 1 | 1 | 8 |
| 1 | 1 | 1 | 1 | 1 | 8 |
| 1 | 1 | 1 | 1 | 1 | 8 |
| 1 | 1 | 1 | 1 | 1 | 8 |
| 1 | 1 | 1 | 1 | 1 | 8 |
| 1 | 1 | 1 | 1 | 1 | 8 |
| 1 | 1 | 1 | 1 | 1 | 8 |
| 1 | 1 | 1 | 1 | 1 | 8 |
| 1 | 1 | 1 | 1 | 1 | 8 |
| 1 | 1 | 1 | 1 | 1 | 8 |
| 1 | 1 | 1 | 1 | 1 | 8 |
| 1 | 1 | 1 | 1 | 1 | 8 |
| 1 | 1 | 1 | 1 | 1 | 8 |
| 1 | 1 | 1 | 1 | 1 | 8 |
| 1 | 1 | 1 | 1 | 1 | 8 |
| 1 | 1 | 1 | 1 | 1 | 8 |
| 1 | 1 | 1 | 1 | 1 | 8 |
| 1 | 1 | 1 | 1 | 1 | 8 |
| 1 | 1 | 1 | 1 | 1 | 8 |
| 1 | 1 | 1 | 1 | 1 | 8 |
| 1 | 1 | 1 | 1 | 1 | 8 |
| 1 | 1 | 1 | 1 | 1 | 8 |
| 1 | 1 | 1 | 1 | 1 | 8 |
| 1 | 1 | 1 | 1 | 1 | 8 |
| 1 | 1 | 1 | 1 | 1 | 8 |
| 1 | 1 | 1 | 1 | 1 | 8 |
| 1 | 1 | 1 | 1 | 1 | 8 |
| 1 | 1 | 1 | 1 | 1 | 8 |
| 1 | 1 | 1 | 1 | 1 | 8 |
| 1 | 1 | 1 | 1 | 1 | 8 |
| 1 | 1 | 1 | 1 | 1 | 8 |
| 1 | 1 | 1 | 1 | 1 | 8 |
| 1 | 1 | 1 | 1 | 1 | 8 |
| 1 | 1 | 1 | 1 | 1 | 8 |
| 1 | 1 | 1 | 1 | 1 | 8 |
| 1 | 1 | 1 | 1 | 1 | 8 |
| 1 | 1 | 1 | 1 | 1 | 8 |
| 1 | 1 | 1 | 1 | 1 | 8 |

| | | | | | |
|---|---|---|---|---|---|
| 1 | 1 | 1 | 1 | 1 | 8 |
| 1 | 1 | 1 | 1 | 1 | 8 |
| 1 | 1 | 1 | 1 | 1 | 8 |
| 1 | 1 | 1 | 1 | 1 | 8 |
| 1 | 1 | 1 | 1 | 1 | 8 |
| 1 | 1 | 1 | 1 | 1 | 8 |
| 1 | 1 | 1 | 1 | 1 | 8 |
| 1 | 1 | 1 | 1 | 1 | 8 |
| 1 | 1 | 1 | 1 | 1 | 8 |
